# A normalized drug response metric improves accuracy and consistency of anticancer drug sensitivity quantification in cell-based screening

**DOI:** 10.1101/262568

**Authors:** Abhishekh Gupta, Prson Gautam, Krister Wennerberg, Tero Aittokallio

## Abstract

Accurate quantification of drug effects is crucial for identifying pharmaceutically actionable cancer vulnerabilities. Current cell viability-based measurements often lead to biased response estimates due to varying growth rates and experimental artifacts that explain part of the inconsistency in high-throughput screening results. We developed an improved drug scoring model, normalized drug response (NDR), which makes use of both positive and negative control conditions to account for differences in cell growth rates and experimental noise to better characterize drug-induced effects. We demonstrate an improved consistency and accuracy of NDR compared to existing metrics in assessing drug responses of cancer cells in various culture models and experimental setups. Notably, NDR reliably captures both toxicity and viability responses, and differentiates a wider spectrum of drug behavior, including lethal, growth-inhibitory and growth-stimulatory modes, based on a single viability readout. The method will therefore substantially reduce the time and resources required in cell-based drug sensitivity screening.

## INTRODUCTION

Cell-based compound profiling plays an important role both in basic biomedical research and in drug discovery. The availability of a wide range of approved and investigational compounds provides an exciting opportunity for systematic drug positioning and repurposing applications, where cellular screening based on phenotypic readouts have become crucial in establishing novel therapeutic strategies against cancers^1–4^. Quantitative assessment of drug efficacies in such large-scale screening efforts is often based on dose-response measurement datasets, where hundreds or thousands of compounds are profiled at several concentrations in a cohort of cancer samples or cell types.

Single parameters or summary metrics based on the end-point dose-response curves are commonly being used to score drug responses in high-throughput studies^2,5–8^. However, due to their dependence on the end-point measurement, these metrics are bound to have systematic differences when applied to different cell types. For instance, fast-growing cells exhibit different response patterns than slow-growing ones, and this difference may be driven by the cell state bias rather than the actual selective drug response. In addition, variations in culture conditions and seeding density also contribute to differences in drug sensitivity measurements^9^.

In the seminal NCI-60 tumor cell line screening project^10,11^, multiple parameters, such as half-growth inhibition (GI_50_), total growth inhibition (TGI), and half-lethal concentration (LC_50_), have been applied to control for the varying growth rates of cells under normal conditions. Recently, a growth rate-based metric (GR) was developed to take into account the variable rate of dividing cells^12^. These approaches are solely based on absorbance/fluorescence differences between drug-treated wells and negative controls, whereas they neglect the information about background noise that can be extracted from wells treated with a positive control (drug that is expected to have killed all cells by the endpoint).

Variability in background noise typically occurs due to artifacts in the assay or differences in the measurement system, in addition to the seeding differences, signal bleed-through, or other experimental factors, and therefore needs to be considered for accurate and consistent drug effect scoring. The normalized percent inhibition (PI) metric uses end-point readouts of the positive control as a proxy for background noise to quantify the variability between measurements^2,13,14^. This metric, however, does not model the dynamic changes that occur from the start of an experiment after treatment. Along with the drug-treated condition and negative control, the positive control readouts can also vary over time and across experimental conditions, which might partly explain the inconsistencies observed in large-scale drug response profiling^8,9,15–19^. Therefore, there is a need for a quantification model that normalizes for the effects of background noise that may vary between measurements and includes model parameters that can be easily interpreted in terms of the experimental and biological factors.

To address these limitations, we devised a normalized drug response (NDR) metric that models the growth rates not only in the drug-treated cells but also in both negative and positive control conditions to accurately capture a wide spectrum of drug effects. The metric makes use of both the start and end-point of a drug experiment to model the dynamics of experimental variability and background noise across various measurement setups. Compared to the other metrics, NDR significantly improves the consistency across measurements and it reliably captures both toxicity and viability phenotypes. Further, based on its improved drug-response curve fitting in various cell growth rates or tissue origins, NDR leads to better drug effect quantification than the existing metrics. We further introduced a summary score (DSS_NDR_), and show how it improves the accuracy of drug effect classification. The application of this metric to quantify drug-responses based on a single viability readout and using a relatively simple measurement setup should make it useful especially in large-scale drug screening efforts.

## RESULTS

### Development and benchmarking of the NDR metric in simulations

To tackle the experimental challenges posed by high throughput screening, including assay-dependent background noise and uneven cell seeding, we devised the normalized drug response (NDR) metric, which is based on the differences in signals measured at the start and the endpoint of an experiment (Fig. 1a). The unique aspect of the NDR is that it models also the dynamic behavior of the positive control, which reflects the sources of experimental variability.

**Figure 1:**
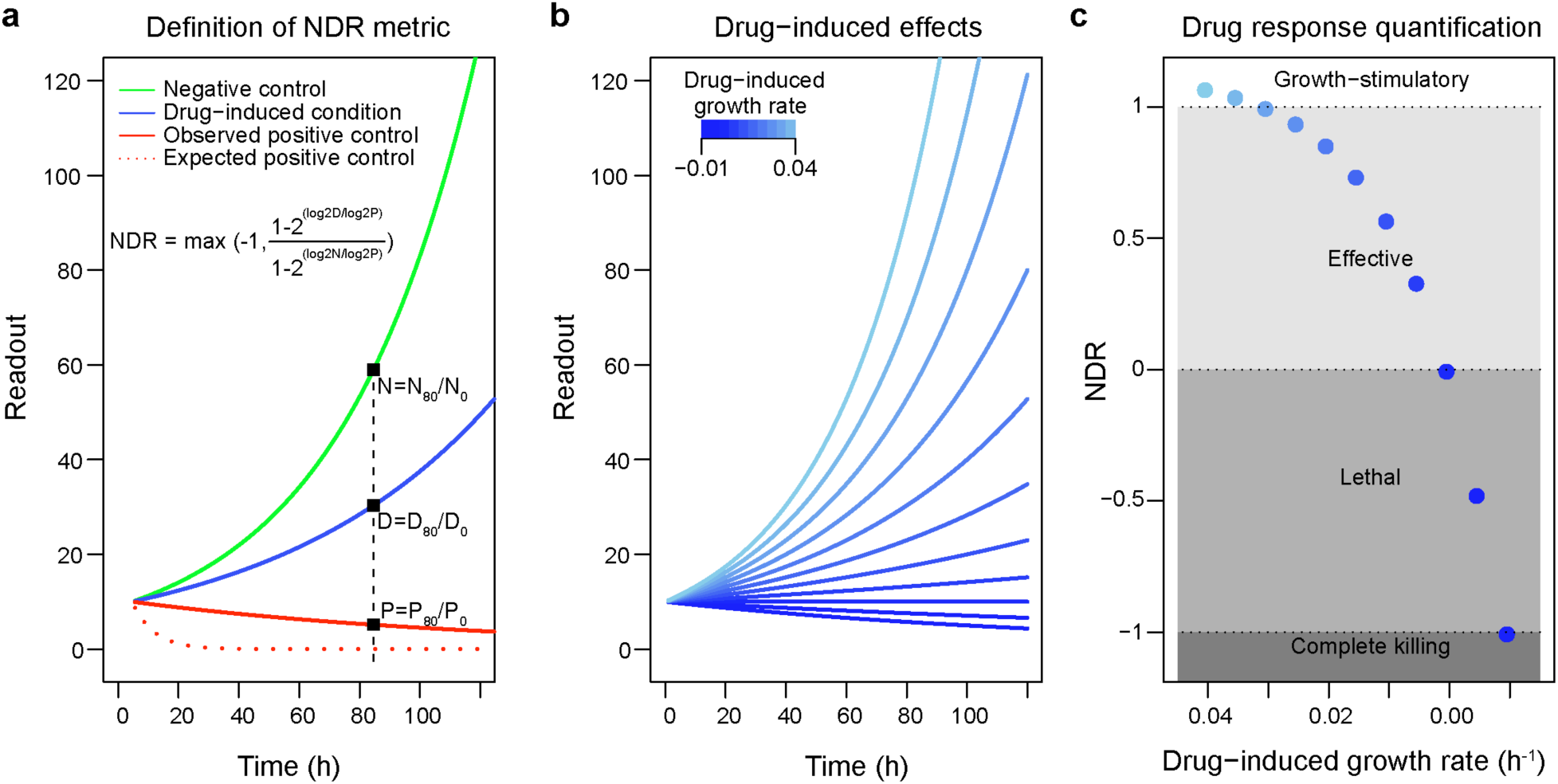
A schematic representation of the NDR metric in various drug-treated and control conditions, assuming that the negative control has no effect and the positive control is 100% lethal. (a) Dynamic change in readout under three simulated settings that reflect negative control condition (green), drug-treated condition (blue), and positive control condition (red). The expected positive control corresponds to the ideal scenario in which the readings of positive control stays at 0, whereas the observed positive control corresponds to the real scenarios in which the readings of positive control is often non-zero. Computation of NDR is demonstrated at a specific time point (*t* = 80 h). (b) Dynamic change in readout under simulation settings that reflect various drug-induced growth rates. The lower growth rates correspond to highly effective drugs or drug concentrations (dark shades of blue). The positive and negative controls are not shown in this panel. (c) NDR metric computed for different drug-induced growth rates under fixed positive and negative control conditions. The spectrum of drug-induced effects as captured by the NDR is illustrated in different shades of gray that spans from −1 to 1.

To systematically assess the performance of NDR, we simulated its outcomes under a fixed set of control conditions in various growth rates mimicking the drug-treated conditions (Fig. 1b). More specifically, we first calculated fold changes for drug-treated and control conditions at a specific time point (here, 80 h), and then used these fold changes to calculate the NDR-based drug response estimates. We found that the NDR metric captures a wider spectrum of possible drug-induced effects, ranging from complete cell death to growth-stimulatory effect (different shades of gray in Fig. 1c).

To further investigate how the NDR metric performs under multiple experimental drug-treated conditions (Fig. 1b), the growth rate of negative control was kept constant while the positive control background was varied to mimic differences in measurement setups (Fig. 2a). For comparison, we also calculated the PI-based and GR-based responses. We found that the PI and NDR responses vary accordingly (Fig. 2b and 2d), indicating that the same readouts in drug-treated condition can lead to different responses, depending on the readouts of the positive control. However, PI had narrower spectrum compared to that of NDR. On the other hand, since the GR metric does not account for the positive control condition, it could not capture these changes in the positive control (Fig. 2c), hence ignoring an important aspect of variability in drug profiling assays.

**Figure 2:**
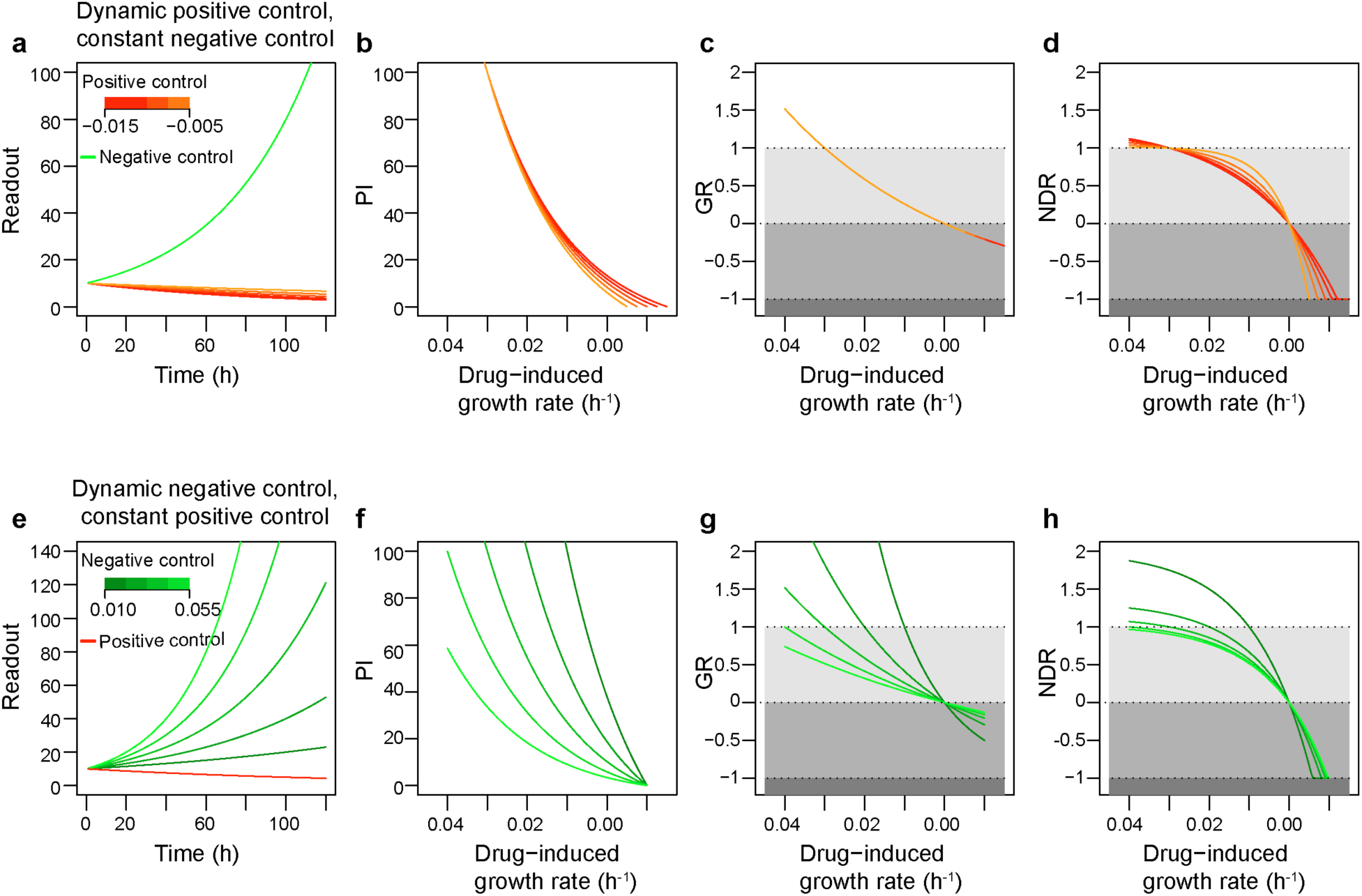
The performance of the NDR metric in various drug-treated and control conditions, assuming that the negative control has no effect and the positive control is 100% lethal. (a) Dynamic change in readout for different growth rates in positive control (shades of red) and constant growth in negative control (green). (b) Percent inhibition (PI) values computed for the conditions shown in Figure 1b and Figure 2a. (c) Growth rate (GR) values computed for the conditions shown in Figure 1b and Figure 2a. (d) NDR metric computed for the conditions shown in Figure 1b and Figure 2a. (e) Dynamic change in readout for different growth rates in negative control (shades of green) and constant growth rate in positive control (red). (f) PI metric computed for the condition shown in Figure 1b and Figure 2e. (g) GR metric computed for the condition shown in Figure 1b and Figure 2e. (h) NDR metric computed for the condition shown in Figure 1b and Figure 2e.

To study the performance of the NDR in cells with distinct growth characteristics, we next kept the positive control background constant while the negative control values were altered to mimic differently growing cells (Fig. 2e). We found that the PI responses were very sensitive to such changes in negative control (Fig. 2f). In contrast, even though both the GR and NDR reasonably accounted for the changes in negative control (Fig. 2g and 2h), NDR remained more stable, especially in the simulated slow growth conditions (Fig. 2h). This is due to the formulation of NDR that does not allow its values to spike up similarly as happens in GR (Fig. 2g). In both simulated conditions, the NDR metric captured a wide spectrum of drug effects, even in cells with slower division time. These improvements are due to its mathematical formulation as well as the capability of NDR to account for the differences in the positive control.

### NDR improves consistency in large-scale drug screening

To investigate the behavior of NDR in drug profiling experiments, we screened MCF-7 and MDA-MB-231 cells in two biological replicate experiments, each with two plates containing 131 oncology drugs in five different concentrations (Supplementary Table S2; see Methods). Since the preparation of single cell suspension for MCF-7 is technically challenging and often compromises its uniform seeding, there were marked differences in the distributions of luminescence intensity readings (RealTime-Glo, Promega) at the start of measurement both within and between the two biological replicate drug screens (Supplementary Fig. 1).

The NDR, GR and PI-based responses were computed for all the wells across the four plates in both cell lines separately. Fig. 3 shows the consistency of NDR between replicates for a plate containing the same drugs as an example in MCF-7 cells. To assess the consistency across replicates, we calculated the absolute difference between the response levels at the corresponding wells of each replicate. The distribution of such differences with the NDR metric is closer to zero compared to those of PI and GI (Fig. 3b; p< 0.005, Wilcoxon rank sum test), implying its improved consistency over replicates. Finally, based on the Z’-factor^20^ as a quality control measure (Fig. 3c), we conclude that the drug response quantification using NDR effectively reduces technical differences between the measurements, and thus improves the consistency over replicated measurements. The differences between the response levels of the corresponding wells of each replicate of MDA-MB-231 cells are shown in Supplementary Fig. 2.

**Figure 3:**
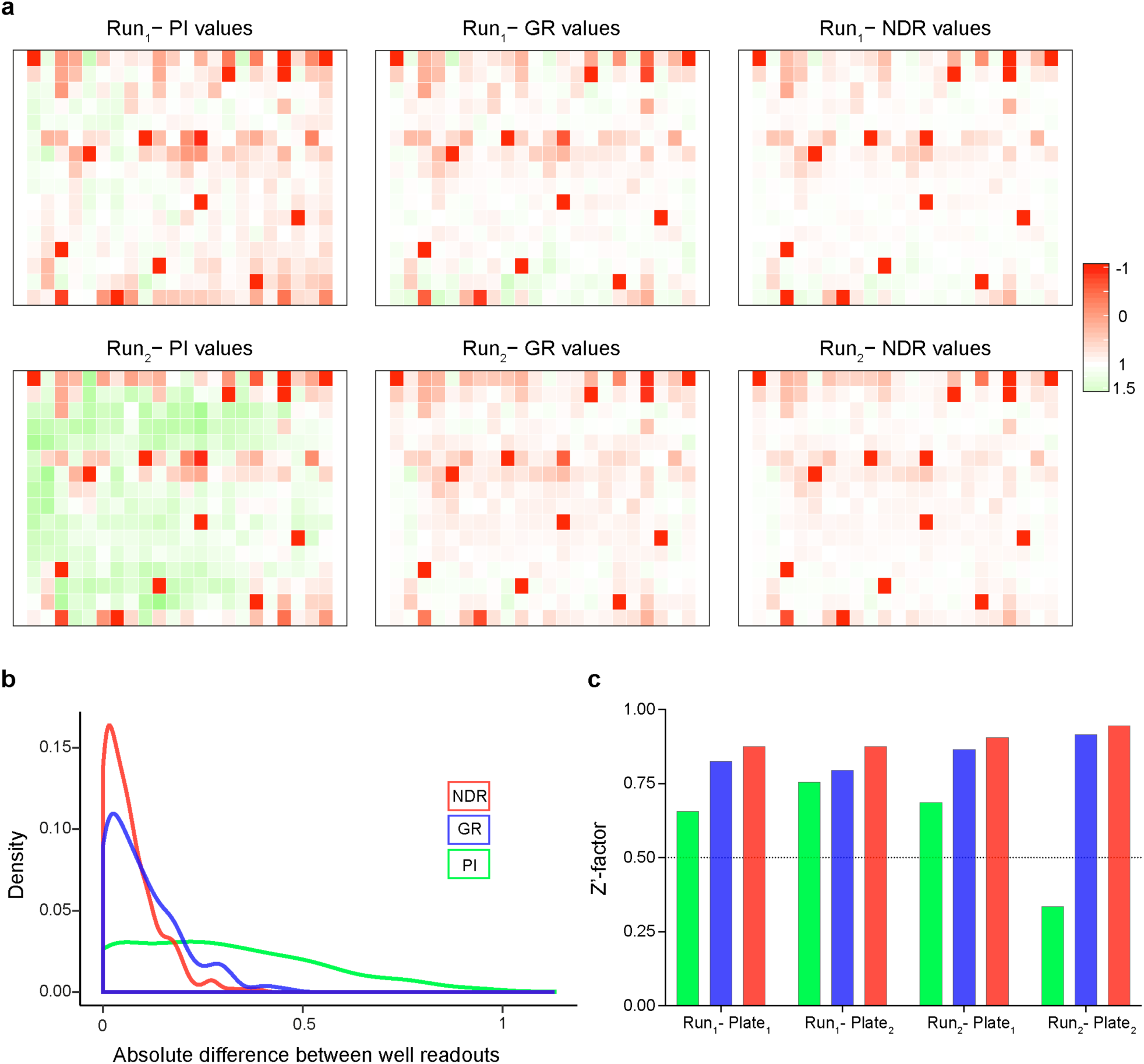
Consistency of the NDR results. (a) PI, GR and NDR values for each well illustrated in plate layouts of 2 replicate drug screens (Run_1_ and Run_2_) in MCF-7 cell line. For comparison purposes, the PI values have been normalized to range between −1 and 1.5. (b) Distribution of absolute difference of identical well positions of the replicate plates. A consistent replicate experiment is expected to result in an absolute difference close to zero. The NDR distribution is closer to zero compared to the PI and GR metrics. The NDR distribution differ significantly from the PI and GR distributions (p<0.005; Kolmogorov-Smirnov test of equality of distributions). (c) Z’-factor for each plate and each replicate experiment. A high-quality assay is expected have a Z’-factor above 0.5 (dotted horizontal line).

We next investigated the consistency of NDR for different cell seeding densities, using Mia-PaCa-2 cells seeded differently at the baseline (before treatments). Notably, we found that the NDR responses were more consistent between two drug profiling experiments, in which 250 and 750 cells were seeded per well at the beginning of the experiment (see Supplementary Fig. 3). To further investigate the behavior of NDR in drug profiling experiments at various end time points, we calculated the difference between the response levels in Pa02C cells screened against 131 oncology drugs at 4 different time points, namely 20h, 28h, 51h and 72h (see Methods). We found the distribution of NDR-based differences was closer to zero, compared to those of PI and GR metrics (p<0.005, Wilcoxon rank sum test; Supplementary Fig. 4), implying an improved consistency of NDR over multiple time points.

**Figure 4:**
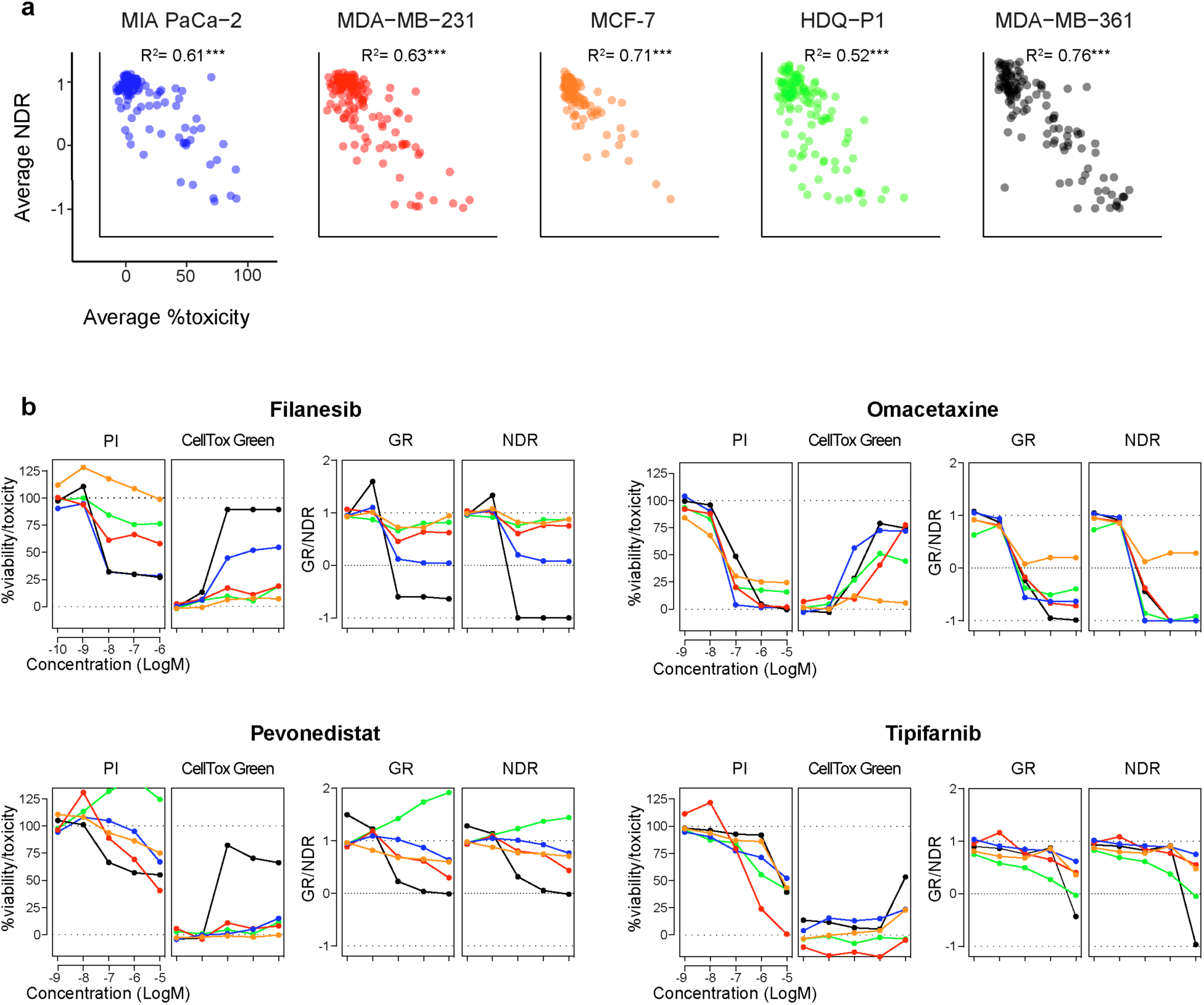
Relationship of NDR with independent toxicity readout (CellTox Green). (a) Scatter plots of the average viability values and independent toxicity readouts of drugs across 5 cell lines (MIA PaCa-2, MDA_MB-231, MCF-1, HDQ-P1, MDA-MB-361) in ascending order of their doubling times (Supplementary Table S1). Coefficient of determination (R^2^) was calculated using Pearson’s correlation. ***p < 0.005, Fisher’s z-transformation. (b) Examples of viability-based PI, GR and NDR results, and percent toxicity responses based on CellTox Green readout across the 5 concentrations of 4 representative drugs. The colors correspond to the different cell lines in panel (a). Similar agreement between the viability and toxicity readouts for all the other drugs are illustrated in Supplementary File 1.

### NDR captures both the toxicity and viability readouts

To further examine the broader behavior of the NDR in drug profiling experiment, we screened 4 additional cancer cell lines against the same set of 131 oncology drugs. The 3 breast cancer cell lines, MDA-MB-231, MDA-MB-361, and HDQ-P1, are known to have different metabolic activity that mimics their doubling times^7,21^. The pancreatic cancer cell line, MIA PaCa-2, was chosen to represent a different tissue type^22^. Furthermore, all these cell lines have been extensively profiled as disease models in chemo-sensitivity studies^23–27^, providing additional information to validate our findings. In agreement with the reported metabolic activity of these cell lines, we observed marked differences in the fold changes of the readouts in the control conditions (Supplementary Fig. 5).

**Figure 5:**
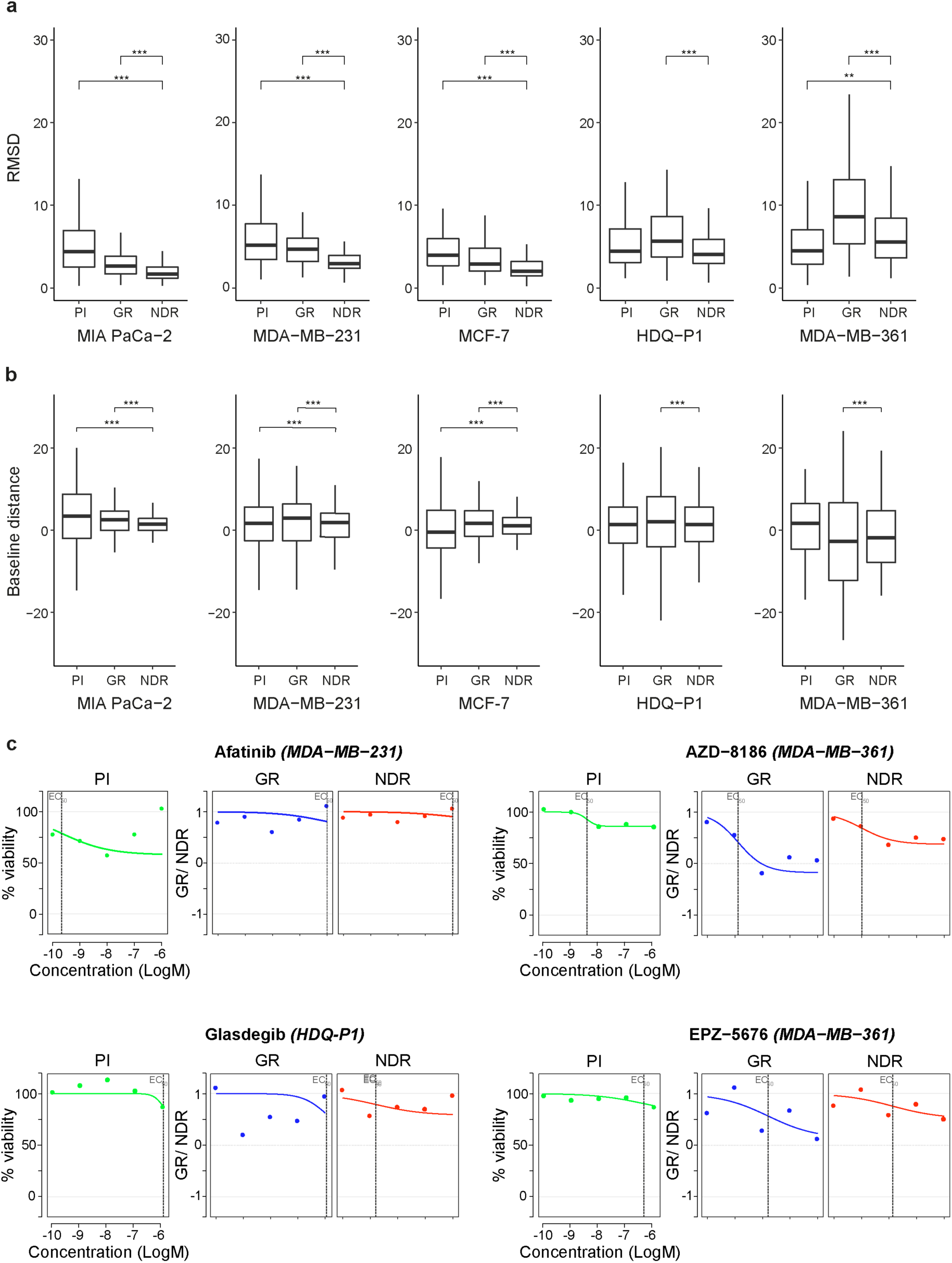
Improved curve fitting with NDR. (a) RMSD error computed between the estimated and observed dose-response curves obtained using the PI, GR and NDR metric in the 5 cell lines (MIA PaCa-2, MDA_MB-231, MCF-1, HDQ-P1, MDA-MB-361). Only drugs that showed non-zero values at least in 1 of the 5 concentrations for all the metrics were considered. **p< 0.05 and ***p< 0.005; Wilcoxon rank sum test for the difference in location. (b) Baseline distance from zero response computed at the lowest drug concentration using the PI, GR and NDR metric in the 5 cell lines. **p< 0.05 and ***p< 0.005; F-test for the difference in variance. (c) Dose-response curves obtained using PI, GR and NDR metric for 4 representative drugs that show extreme differences in curve fittings. The representative drugs illustrate both the improvement in curve-fittings as well as decrease in baseline distances in the respective cell lines.

To validate the reliability of the drug response results, we also measured an independent cytotoxicity end-point readout (CellTox Green, Promega). For all the cell lines tested, NDR at each drug concentration decreased with the increasing toxicity readout for most of the drugs, suggesting that the NDR relates closely to the toxicity measurements. As expected, the average NDR-based viability was negatively correlated with the average toxicity readout (p < 0.005; Fig. 4a).

The results from 4 representative compounds with different mechanism of action and differential response across the 5 cell lines illustrate the ability of NDR to capture not only the viability but also the toxicity readouts (Fig. 4b). Filanesib, a kinesin spindle protein inhibitor induced a selective toxic response towards MIA-PaCa-2 and MDA-MB-361. The toxicity reading of filanesib in MIA PaCa-2 and MDA-MB-361 cell lines were well-differentiated both with NDR and GR but not with the PI-based response. Although the GR values are similar to those of NDR, the NDR metric is more predictive of the toxicity responses, as was evident especially in the slowly-growing MDA-MB-361 cells (Fig. 4b). Based on NDR, filanesib treatment seems to only induce cytostatic effect on MIA-PaCa-2 cells, whereas cell death readout suggests that it induces cell death as well. Even though filanesib treatment halted the cell division of MIA-PaCa-2 cells, there remained some degree of cell division, which was balanced out by the cell death, hence imitating the cytostatic effect. This was further confirmed by imaging-based drug effect assessment (see Methods section and Supplementary Fig. 6).

**Figure 6:**
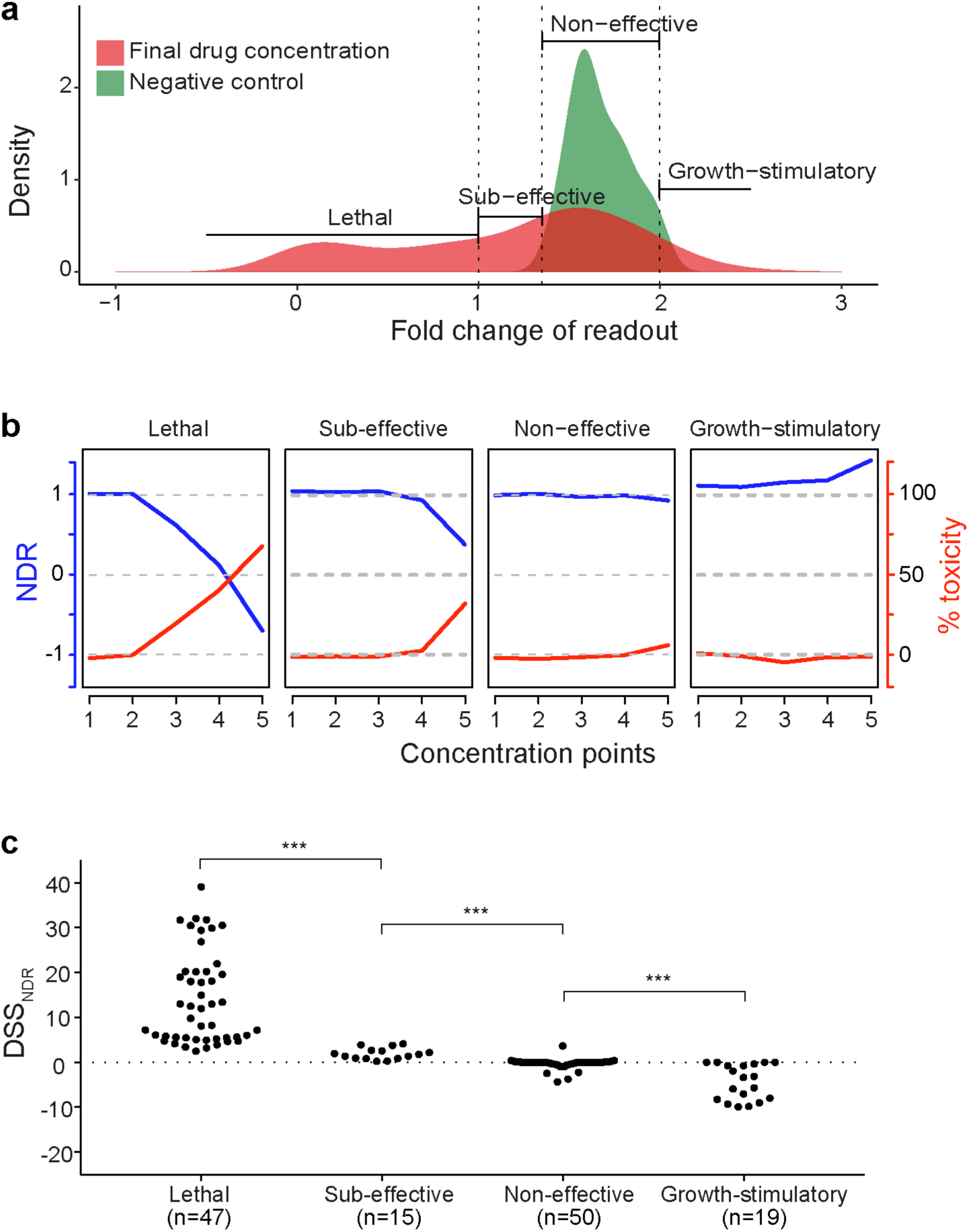
DSS_NDR_ categorizes drugs based on their potency. (a) Categorization of drugs based on the difference between the fold change at final concentration (red) and the fold change in the negative control (green) in MDA-MB-361. The drugs with fold change lower than 1 in the final drug concentration are classified as “Lethal”. The drugs with fold change between 1 and 2 standard deviation (SD) below the growth rate in the negative control (DMSO) are categorized as “Sub-effective”. The drugs with fold change greater than 2 SD of growth rate in DMSO are categorized as “Growth-stimulatory”. The remaining drugs, with fold change similar to that in DMSO, are categorized as “Non-effective”. (b) The class-specific drug behavior quantified by the average NDR-based viability readout (blue) and average toxicity readout (red) in MDA-MB-361. (c) DSS_NDR_ levels of drugs in the four classes. ***p<0.005, Wilcoxon rank sum test.

Omacetaxine, a protein synthesis inhibitor, was toxic in all cell lines except for MCF-7. This selective behavior was missed by the PI-based readout. Furthermore, NDR and GR were also able to capture the cytostatic behavior of omacetaxine against MCF-7 cells, which could neither be inferred from PI-based viability or toxicity measurements. Tipifarnib, a farnesyltransferase inhibitor, was largely non-toxic to all the cell lines, which was clearly reflected with the NDR but not in the PI readout. Finally, pevonedistat, a NEDD8 activating enzyme inhibitor was cytotoxic only to the MDA-MB-361 cells, which was reflected both in the NDR and GR responses. NDR and GR additionally revealed a growth-stimulatory (enhanced metabolic-activity) effect in the HDQP1 cell line. Based on these results, we conclude that NDR better captures the viability and cytotoxic behavior of different drugs, even though the overall performance of GR is relatively similar.

### NDR improves drug response quantification in large scale screening

To quantify the drug response for each drug, we generated dose-response curves based on NDR at 5 concentrations using drc R-package^28^. Based on visual inspection of the dose-response curves, we found a consistent improvement in the curve fitting across all 5 cell lines when compared to the curves obtained by GR and PI using the same drc package (exemplified in Fig. 5c). To quantify the curve-fitting behavior, we calculated the root mean squared distance (RMSD) between the observed and estimated dose-response curves for all the drugs with non-zero response in at least 1 of the 5 concentrations (illustrated in Supplementary Fig. 7).

The average RMSD calculated using NDR was lower in the fast-growing cell lines (MIA PaCa-2, MDA-MB-231 and MCF-7), when compared to both the GR and PI-based responses (p < 0.005, Wilcoxon rank sum test; Fig. 5a). Notably, in the slow growing cell lines (HDQP1 and MDA-MB-361), the GR normalization led to increased RMSD values compared to those obtained from the NDR and PI. The simple PI performs better than NDR in the MDA-MB-361 cells, which is the slowest growing cell line among the cell lines tested here. This suggests that in case of slow growing or non-dividing cells (during experimental time), even PI provides adequate responses provided there is not considerable differences in cell seeding uniformity. However, the PI normalization for these cells is bound to be less effective in detecting cytostatic effects (see Supplementary File S4).

In dose-response curves, the lowest drug concentration is usually expected to have minimal or no effect, and any deviation from this baseline behavior can eventually bias the drug sensitivity parameters, such as IC_50_ or EC_50_ values. We observed that the NDR responses at the lowest drug concentration consistently was closer to the negative control level when compared to the GR and PI-based responses. To quantify this, we computed the distance of lowest concentration response from negative control viability value (100 for PI, 1 for GR and NDR), termed as baseline distance (illustrated in Supplementary Fig. 7). The variability of the baseline distance with NDR in the fast-growing cell lines was significantly lower than that obtained using GR and PI (p < 0.005, F-test for difference in variances; Fig. 5b). In the slow-growing cells, GR led to an increased variance of the baseline distances when compared to NDR or PI (see Fig. 5c for representative examples). We note that there were also drugs with low baseline distance but high RMSD values, implying that the lower baseline distance does not necessarily result in a lower RMSD values (see Supplementary Fig. 8).

Similar NDR-driven improvements in the RMSD values and baseline distances were also found when analyzing dose-response curves of MDA-MB-231 in two external datasets, one from Cancer Therapeutics Response Portal (CTRPv2)^29,30^ and the other from Genomics of Drug Sensitivity in Cancer (GDSC1000)^31^ (Supplementary Fig. 9). Furthermore, the NDR metric improved the dose-response curve fittings of 131 drugs screened against freshly extracted mononuclear cells from the bone marrow of an AML patient (Supplementary Fig. 10), demonstrating its benefits also for functional profiling-based precision medicine.

### NDR-based DSS distinguishes a wide spectrum of drug effects

After confirming that the NDR enables reliable quantification of different drug effects, we computed the NDR-based DSS (DSS_NDR_; see Methods) that summarizes the dose-response relationships over the whole concentration range into a single response score. As expected, the distributions of DSS_NDR_ showed selective efficacy of only a few drugs in a particular cell line (Supplementary Fig. 11). We next investigated whether DSS_NDR_ values could be used to uniquely differentiate the drug-potency of all drugs in the screening panel.

As a ground-truth, we first classified the effects of the drugs into four drug response categories, namely, lethal, sub-effective, non-effective and growth-stimulatory, based on their fold change of the viability readouts at their highest drug concentration in the 5 cell lines (see Fig. 6a for MDA-MB-361; Methods). The viability readout of this classification showed a good overall agreement with the independent toxicity readout (Fig. 6b). More specifically, at higher concentrations of lethal drugs, the viability readout dropped while the toxicity readout increased. This behavior of the two readouts was weaker for sub-effective drugs that comprise either less toxic (not lethal) or cytostatic drugs. As expected, the toxicity readout barely changed for non-effective drugs and growth-stimulatory drugs. The viability readout, on the other hand, changed negligibly in response to non-effective drugs, but increased at higher concentrations of growth-stimulatory drugs. We further confirmed that the lethal class included drugs that are known to be toxic in these cell lines^23,24,32^, for example, proteasome and HDAC inhibitors (see Supplementary Table S3). Furthermore, most of the anti-mitotic and kinase inhibitors were classified as sub-effective (Supplementary Table S3).

The four drug categories showed distinct DSS_NDR_ distributions in MDA-MB-361 cells (p <0.005, Wilcoxon rank sum test; Fig. 6c). The lethal drugs were well-separated from the sub-effective drugs based on their DSS_NDR_ values. Furthermore, most of the non-effective drugs had a DSS_NDR_ close to zero, whereas the growth-stimulatory drugs tend to have a negative DSS_NDR_. Similar distributions for the 4 drug response categories were observed when we merged all the drugs across all the cell lines (Supplementary Fig. 12a). When comparing the NDR-based findings with those computed using the GR and PI metric, we found that even though the distribution of DSS looks similar, the overlap between the distributions of adjacent drug categories is smallest when using NDR (Supplementary Fig. 12). This suggests that it is possible to reliably infer the effect of a drug solely based on its DSS_NDR_ value, reducing the requirement of other validation experiments.

## DISCUSSION

*In vitro* cell-based drug screening is commonly carried out as an end-point cell count surrogate assay, and such end-point drug response profiling is being widely applied to quantify the sensitivity of drugs in cancer cell lines or patient-derived samples. In these screening efforts, robust dose-response curve fitting is pivotal for defining accurate drug vulnerabilities. Due to the experimental limitations and noise inherent to high throughput settings, however, it is often difficult to obtain smooth dose-response curves using the existing measures, which results in significant number of false positive and negative hits. The experimental errors typically originate from inconsistent seeding of cells, or their differing growth rates, as well as from different readouts, among other technical issues^16^. Due to the scale of these profiling experiments and their running costs, it is undesirable and many times even impossible to repeat the whole experiment, hence calling for a response metric that effectively normalizes for such experimental errors and decreases the false hit rates, with the aim to reduce the time and resources required in cell-based drug sensitivity screening and to make their results more reproducible and accurate.

In this study, we developed and carefully tested a novel NDR metric that reduces the effects of experimental inconsistencies, leads to more accurate dose-response curves, and therefore improves the consistency and accuracy of drug profiling results. The metric is based on the comparison of the end point readout with that of the initial state of an experiment in the drug-treated condition, as well as accounting for both negative and positive control conditions. By means of systematic simulations, we first demonstrated how the NDR reliably captures the wide spectrum of drug responses under different control conditions. The other metrics, such as GR, do not account for the positive control condition, and therefore it fails to capture drug responses in slow-dividing cells. This is particularly relevant in experimental models based on primary cells or patient samples, which generally grow slower than established cell lines. In studying such responses, the NDR was able to capture the various drug effects more accurately, as demonstrated by the simulations (Figs. 1 and 2), and in a proof of concept experiment of AML patient sample *ex vivo* drug screening (Supplementary Fig. 10). Such improvements are especially important in personalized oncology applications that rely on accurate and consistent drug effect quantification for optimal treatment selection based on primary cell-based compound screening ^1,2,4,5^.

The availability of real-time viability measurement reagents made it possible to test the NDR metric in large-scale drug screening setups. MCF-7 replicate drug screening results suggested that NDR effectively reduces the experimental variability, and thus significantly improves the consistency between drug response measurements between biological replicates, different end time-points and different seeding densities (Fig. 3 and Supplementary Figs. 2-4). This will offer improved solutions to the ongoing debate on the inconsistencies in drug response profiling^9,15–19^.

While the existing drug response calculations are prone to the variability between measurements, NDR metric is likely to lead to more consistent comparison of drug response quantifications across different samples as well as across different measurement assays. The NDR might also become valuable in 3D-culture models or clonogenic drug screening assays, where uniform cell seeding is crucial. The reliability of NDR responses computed for 131 drugs across 5 cell lines with different doubling times was further validated by a parallel cell toxicity screen (Fig. 4).

The higher consistency of NDR over the other metrics was also evident in the improved drug-response curve fittings (Fig. 5). As error in a single data point of a dose-response curve fit can result in overestimation of drug response, these results demonstrate that the NDR metric does not only improve curve fitting and the baseline quantification of drug effects, but consequently also reduces false hit callings in large-scale screenings. False hits are a major practical problem in cell-based drug sensitivity screening, as they lead to both increased costs and times of the secondary confirmation experiments, or, when not controlled for, confusion and inconsistencies in the published literature. The benefits of NDR were also confirmed on the external CCLE and GDSC datasets (Supplementary Fig. 9), highlighting the wide-applicability of NDR in various large-scale screening assays. The improved results in slow dividing patient-derived primary cells further support the usage of NDR as an accurate metric in the functional profiling-based personalized medicine applications.

Viability/metabolic-activity measurements are classically used to assess the drug effect in large-scale screenings. Even though metabolic activity is considered as representative of the number of cells, reduction in viability does not always correspond to lethality^24^; rather, it may instead represent cytostatic, or anti-metabolic effects. We showed here that NDR-based DSS can be used to infer the drug behavior from a single viability measurement. More specifically, we showed that based on DSS_NDR_ values, one can reliably categorize the drugs according to their differential effects: lethal, sub-effective, non-effective and growth-stimulatory (Fig. 6). This has a significant impact on large-scale high throughput drug profiling efforts as it will notably reduce the cost and time of further validation for cytotoxicity. Moreover, detection of growth-stimulatory drugs is very important in precision medicine as it provides insights into the cellular mechanism of specific cells, tissues or diseases. As drug resistance against monotherapies has directed oncology research towards combinatorial approaches, identifying growth-stimulatory targets will be valuable for deciphering the disease specific resistance driving pathways, and thereby devising novel and effective drug combination strategies.

One of the main limitations of metabolic readout-based viability measurement is its inability to distinguish the concurrent cell growth and cell death since the estimated cell growth with metabolic readout is the sum of growing and dead cells. As a result, metrics implemented for high-throughput settings, such as NDR, capture only the beginning and end of a given treatment period, but not the complex treatment dynamics. This issue can be addressed utilizing time-lapse high-content image-based profiling techniques, such as drug-induced proliferation (DIP) metric^33^. Even though such imaging methods can accurately measure the drug–induced effects, however, their translation to higher throughput drug profiling settings still remain a major challenge because of need of continuous imaging. Furthermore, as the DIP approach involves genetically engineered fluorescently labeled cells, its applications to the primary cells or patient samples is not straightforward. More recently, a scalable time-lapse analysis of cell death kinetics (STACK) method was introduced to quantify the kinetics of compound-induced cell death at the cell population level^34^. However, this method is based on a single control only. In the future, it would be therefore interesting to combine the benefits of NDR with the STACK-based methodology.

Based on the present results, we conclude that NDR accurately portrays a widened spectrum of drug-induced effects, as well as results in improved consistency and accuracy across different measurement systems and culture conditions in high-throughput drug profiling setting. The calculation of NDR requires only a minor modification in the widely-used experimental setups for high-throughput drug profiling, making the NDR-based drug response quantification broadly feasible and beneficial in a wide range of applications with cell-based chemical screening.

## MATERIAL AND METHODS

### Cell lines

The cell lines used in this study were human breast cancer cell lines MDA-MB-231, MDA-MB-361, HDQ-P1, MCF-7 and pancreatic ductal adenocarcinoma MIA PaCa-2 (details in Supplementary Table S1). All breast cancer cell lines were purchased from ATCC and MIA PaCa-2 was a generous gift from Professor Channing Der (University of North Carolina at Chapel Hill, NC, USA). All cells were maintained in DMEM with 2.2 g/L NaHCO_3_ (Life Technologies) at 37°C with 5% CO_2_ in a humidified incubator, according to provider’s instruction.

### Drug Screening

131 oncology compounds library (Supplementary Table S2) was screened against the cell lines using Drug Sensitivity and Resistance Testing (DSRT) platform, as previously described^2,24^ with a slight modification. In brief, compounds were added in 384-well plates (Corning) in 5 different concentrations covering 10,000-fold concentration range using an Echo 550 Liquid Handler (Labcyte). 0.1% dimethyl sulfoxide (DMSO) and 100 μM benzethonium chloride were used as negative and positive controls respectively. The library screen was performed only once for each compound concentrations on each cell lines. The pre-dispensed compounds were first dissolved in 5 μl of complete medium per well containing RealTime-Glo (1:1000 final volume, Promega) and CellTox Green (1:2000 final volume, Promega) and then 20 μl of cell suspension per well was added maintaining the required final cell densities as listed in Supplementary Table S3. As initial point, after 1 h of seeding and as end point, after 72 h of incubation, toxicity (CellTox Green, fluorescence) and viability (RealTime-Glo, luminescence) of the treated cells were measured using a PHERAstar FS plate reader (BMG Labtech).

AML patient bone marrow sample was obtained after informed consent with ethical committee approval (No. 239/13/03/00/2010, 303/13/03/01/2011) and in accordance with the Declaration of Helsinki. Primary AML cells were freshly isolated from patient bone marrow sample, maintained in culture overnight and *ex vivo* DSRT was performed the next day against same 131 compound library similarly as described earlier.

### Drug effect assessment with imaged-based readout

MIA-PaCa-2 cells expressing nuclear mKate2 (Nuclight Red) were treated with different concentrations of filanesib in five replicates. The experiment was carried out in 384 well plate seeding 750 cells per well. CellTox Green (Promega) was used to monitor the number of dead cells. Changes in the numbers of live (red nuclei) and dead (green nuclei) cells were followed for 72h (every 6h) in IncuCyte (Sartorius).

### Drug response metrics

#### NDR metric

The normalized drug response (NDR) is formulated to account for the changes under drug-treated as well as the changes in both (positive and negative) control conditions. For that, NDR first calculates the change in drug-treated condition when compared to the positive control, which ideally is representative of the background noise as positive control is expected to kill all cells. Similarly, NDR also calculates the change in the negative control when compared to the positive control. Both these changes are then used to quantify the total effect of a drug, hence effectively eliminating background noise and other sources of experimental variability.

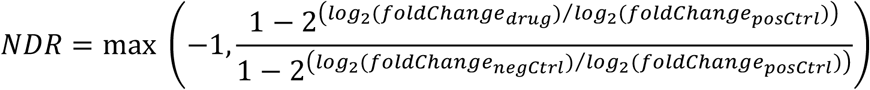

where the fold change between the readouts at start and end-point of the measurement is given as:

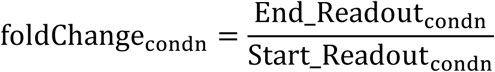

The log_2_ scaling is used merely for the visualization and comparison purposes. The NDR calculation is valid for any other logarithm base constants, which will result in quantitatively similar results. To make the NDR calculation less sensitive against the effect of outliers in the control conditions, we have used median values of the fold changes in our calculations. More specifically, the median of readings for BzCl were used as positive control and the median of readings for DMSO were used as negative control. From the raw data (in Supplementary Table S5), the readings of xBzCl and a drug combination (cytarabine/idarubicin) were ignored in the current analysis.

Based on the NDR values, the drug effects can be classified as (see Fig. 1):

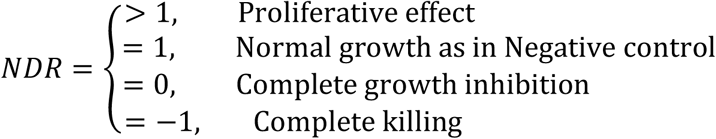

#### GR metric

The growth rate (GR) metric, based the work of Hafner and colleagues^12^, is computed as:

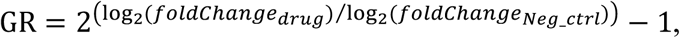

where the fold change between the readouts at start and end of the measurement is given as:

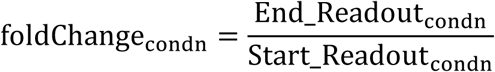

#### PI normalization

The percent inhibition (PI), based on the endpoint readouts, is computed as:

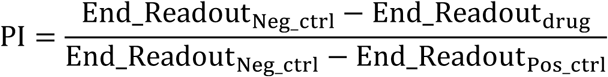

### DSS calculation

Drug sensitivity score (DSS) is a quantitative scoring approach based on the continuous model estimation and interpolation to effectively summarize the complex dose-response relationships^6^. More specifically, for a normalized drug-response *R*(*x*) at concentration *x*, the integral response *I* over the dose range that exceeds a given minimum activity level (*A*_min_) is calculated analytically as a continuous function of multiple parameters of the non-linear response model, including its slope at IC_50_ as well as the top and bottom asymptotes of the response (R_max_ and R_min_). Formally, the DSS is computed as

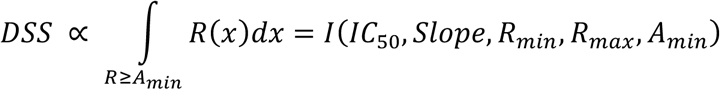

For the DSS-related analyses, we used the DSS R-package freely available at https://bitbucket.org/BhagwanYadav/drug-sensitivity-score-dss-calculation. As the input to DSS computation R-package, we scaled the metrics as:

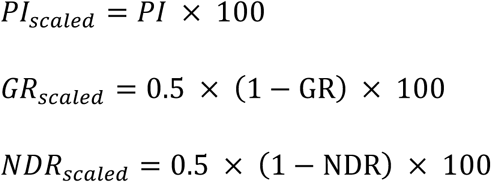

To compute the negative DSS for the drugs that have negative responses *R*(*x*) in all the five concentrations tested, we flipped the responses, using 1-*R*(*x*) scaling, so that the fitting of the drug-response curves was effectively mirrored. After the DSS values were computed based on the mirrored drug-response curves, we set the DSS value to be negative.

### Data analysis and statistical tests

All the data analysis and statistical test were performed in the R statistical programming environment (http://R-project.org). All raw data and summary results as well as R function to compute and reproduce the NDR calculations and the figures are available at: https://github.com/abishakGupta/NDR_results.

#### Statistical analysis

To evaluate the association between two response profiles, we used Pearson correlation coefficient^35^. The statistical significance (p-values) of the Pearson correlation coefficient values was computed using the Fisher’s z-transformation. Shapiro-Wilk test was used to test a normality of a distribution. If the normality was established, the difference in means or variances between two distributions was assessed using Welch two-sample t-test or F-test, respectively. To assess whether two non-normal distributions differ in their location, we used the non-parametric Wilcoxon rank sum test^36^. To compute the overlap between two distributions, we used the overlapping coefficient^37^ as a point estimate of the overlap between two normal densities.

#### Root mean squared distance (RMSD) calculation

To quantify the goodness of dose-response curve fits, we computed the root mean squared distance (RMSD) between the observed and estimated values of the response curves. We used the conventional formula of RMSD computation given as:

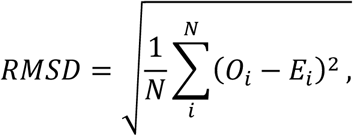

where *N* is the number of concentration points, and *O*_*i*_ and *E*_*i*_ are the observed and estimated drug response values at concentration *i*, respectively.

### Simulated drug response data

To systematically test the NDR metric performance in a fully-controlled ground-truth setup, we used simulated data of representative drugs, where the control conditions were varied at different realistic rates.

For the first simulation model, we set the growth rate of negative control to 0.03 h^-1^, such that the doubling time was ∼30 h and the change rate in positive control to −0.01 h^-1^. We set the growth rate of representative drugs to lie in between these rates of the controls. We also added growth rates higher than those in the negative control (with doubling time of ∼25 h) to emulate the growth stimulating effect. We then computed the NDR metric at a specific time point with *foldChange*_*negCtrl*_ = 4 folds, *foldChange*_*postCtrl*_ = 0.5 folds, and *foldChange*_*Drug*_ = 0.5 to 8 folds.

For the second simulation model, with the same representative growth rates of drugs, we set the growth rate of negative control to 0.03 h^-1^ and let the growth rate of positive control to vary from −0.015 to −0.005 h^-1^. We then computed the NDR metric at a specific time point with *foldChange*_*negCtrl*_ = 4 folds, *foldChange*_*postCtrl*_ = 0.4 to 0.8 folds, and *foldChange*_*Drug*_ = 0.5 to 8 folds.

For the third theoretical model, with the same representative growth rates of drugs, we let the growth rate of negative control to vary from 0.01 to 0.055 h^-1^ and set the growth rate in positive control to −0.01 h^-1^. We then computed the NDR metric at a specific time point with *foldChange*_*negCtrl*_ = 2 to 15 folds, *foldChange*_*postCtrl*_ = 0.5 folds, and *foldChange*_*Drug*_ = 0.5 to 8 folds.

### Drug classification

The 131 drugs used in the drug sensitivity and resistance testing (DSRT) assay were classified into four groups, based on the fold change of the viability readouts at the highest drug concentration from the start to the end-point of measurement. The first group of drugs included the ones with a fold change less than 1. The final readout for these drugs is smaller than the readout at start, and hence these drugs are labeled as “lethal”. As a second group, the drugs with fold change above 1 and lower than 1 standard deviation (SD) on the lower side of growth rate in the negative control (DMSO) were labeled as “sub-effective” (Supplementary Fig. 12). This group of drugs is expected to comprise of cytostatic as well as less toxic drugs. The third set of drugs is labeled “non-effective”, since their fold change was similar to the growth rate in the negative control condition. The final drug group consists of drugs that result in proliferation higher than in 1 SD on the higher side of the growth rate in the negative control, and are labelled as “growth-stimulatory”.

### NDR calculation on CCLE and GDSC datasets

To test the performance of NDR in independent datasets, we extracted two publicly available raw drug sensitivity screening data, namely Cancer Therapeutics Response Portal (CTRPv2)^29,30^ from the Broad Institute and Genomics of Drug Sensitivity in Cancer (GDSC1000)^31,38^ datasets from the Sanger Institute. We used MDA-MB-231 cell line data against all drugs and across all concentrations (9 concentrations in GDSC1000 and 16 in CTRPv2).

As measurements at the beginning of the experiments were not available in both datasets, we estimated the starting value based on the fold change (3.2) that was observed in our screens for MDA-MB-231 cells, which is also similar to growth rate reported by others ^7^. The estimated values were then used in the GR and NDR computation.

## AUTHOR CONTRIBUTIONS

A.G. and P.G. conceived this study and wrote the manuscript. A.G. devised the NDR metric and performed the computational analyses. P.G. designed and performed the experiments. K.W. and T.A. supervised the work and critically reviewed and revised the manuscript.

### ACKNOWLEDGEMENTS

We sincerely thank Sanna Timonen for her help with drug screening, Alok Jaiswal and Gopal Peddinti for their valuable inputs for the NDR metric. We thank Laura Turunen and other members of High Throughput Biomedicine unit of the FIMM Technology Centre (supported by University of Helsinki/Biocenter Finland research infrastructure funds) for technical assistance. The authors also thank the Individualized System Medicine (ISM) group at the Institute for Molecular Medicine for providing AML patient cells and Petr Smirnov (University of Toronto) for reproducing our results.

This project was supported by the Academy of Finland (grants 292611, 279163, 295504, 310507 to TA; 272577, 277293 to KW), the Cancer Society of Finland (PG, TA and KW), the Sigrid Jusélius Foundation (KW and TA) and the AACR/Pancreatic Cancer Action Network (KW).

