## Supplementary Figures for "A normalized drug response metric improves accuracy and consistency of anticancer drug sensitivity quantification in cell-based screening"

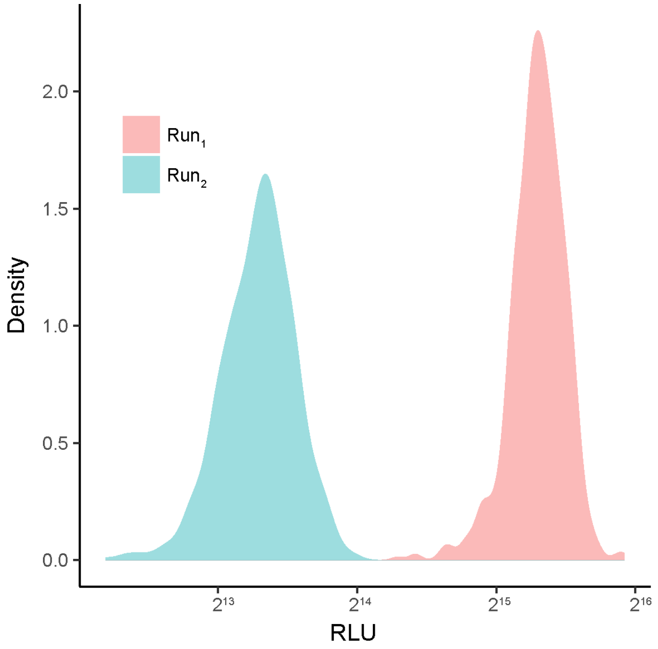


**Supplementary Figure 1**: Distribution of well readouts at the start of experiment, as measured with the relative luminescence unit (RLU) for two replicate drug screens ($Run_1$ and $Run_2$) in MCF-7 cells.

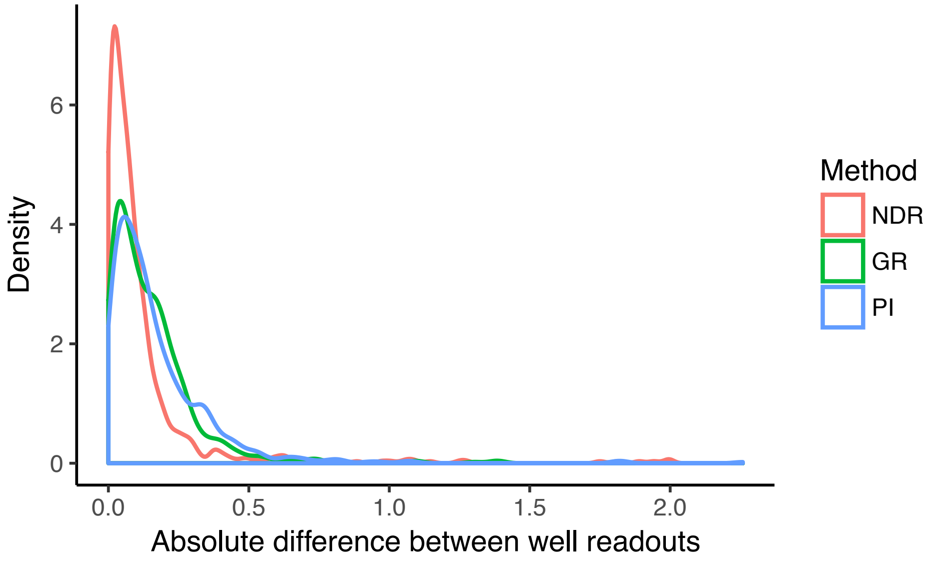


**Supplementary Figure 2**: Distribution of absolute difference of identical well positions of the replicate plates for MDA-MB-231 cells. A consistent replicate experiment is expected to result in an absolute difference close to zero. The NDR distribution is closer to zero compared to the PI and GR metrics (p< 0.005, Wilcoxon rank sum test).

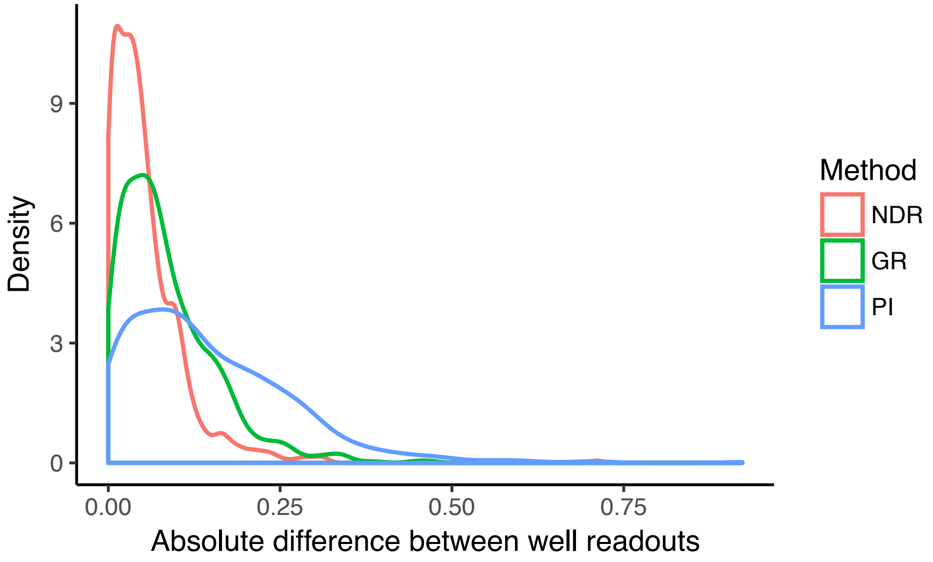


**Supplementary Figure 3**: Distribution of absolute difference of identical well positions of the plates for MIA-PaCa-2 cells seeded differently. Two experiments with initial seeding of 250 and 750 cells were performed for this. A seeding-consistent response is expected to result in an absolute difference close to zero. The NDR distribution is closer to zero compared to the PI and GR metrics (p< 0.005, Wilcoxon rank sum test).

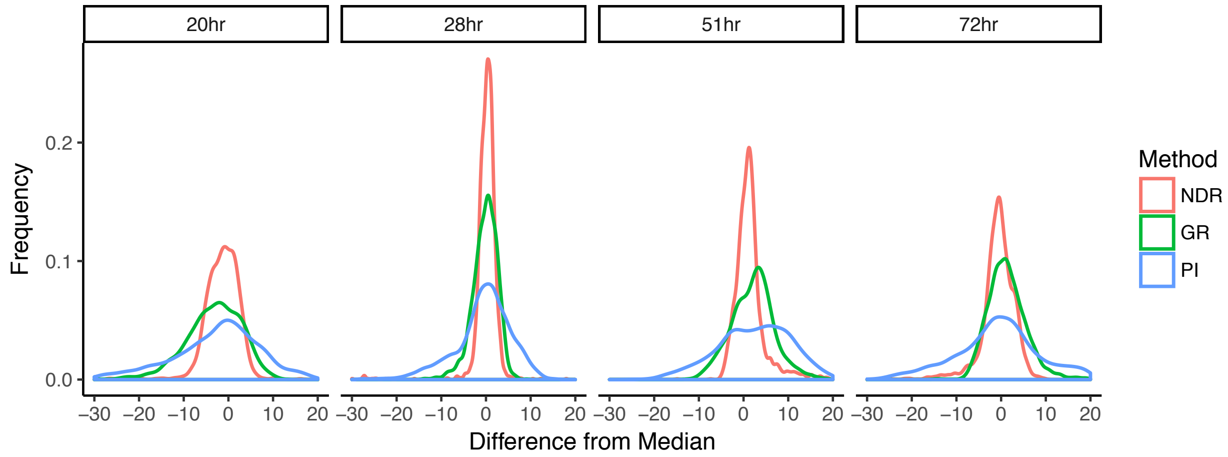


**Supplementary Figure 4:** Consistency of the NDR results across time. Distribution of the difference between the response at a given time and median response across all times. A time-consistent response is expected to result in an absolute difference close to zero. The NDR distribution is closer to zero compared to the PI and GR metrics (p< 0.005, Wilcoxon rank sum test).

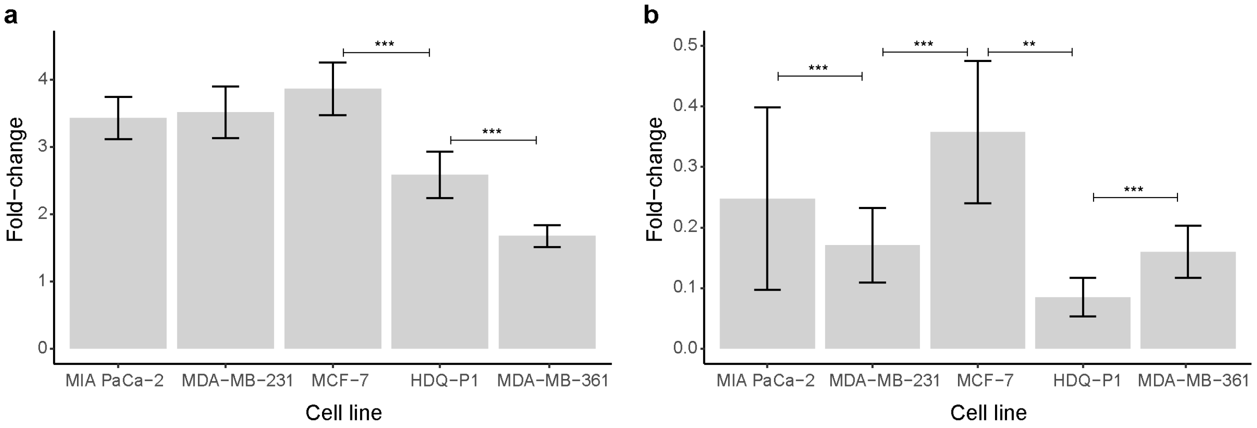


**Supplementary Figure 5**: (a) Mean fold change in luminescence readings between the start- and the end-point measurements for wells containing the negative control (DMSO). (b) Mean fold change in luminescence readings between the start-point and the end-point measurements for wells containing the positive control (BzCl). Also shown in both the plots are the standard deviations (error bars). **$p < 0.05$, ***$p < 0.005$; Welch Two Sample t-test after normality test using Sharpiro-Wilk test.

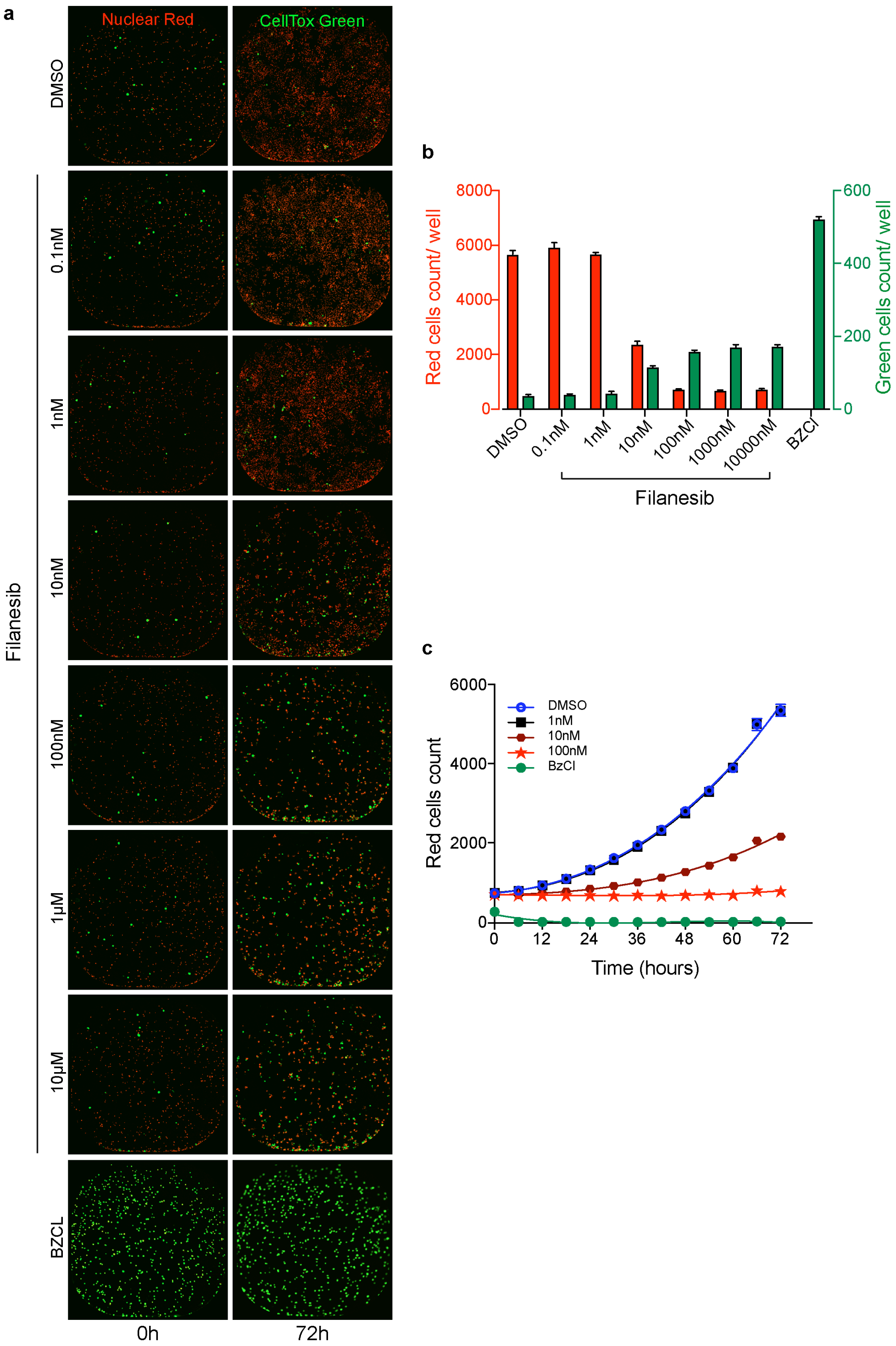


**Supplementary Figure 6**: The unique balance between cytostatic and cytotoxicity effect of filanesib on MIA-PaCa-2. (a) Images of filanesib treated cells expressing nuclear mKate2 (Nuclight Red) at two time point initial (0h) and endpoint (72h). Dead cells nuclei are stained green and red nuclei represents viable cells. (b) Quantification of live (red) and dead (green) cell count in different treatment conditions. (c) Change in number of viable cells (red count) over the experimental period in different treatment conditions.

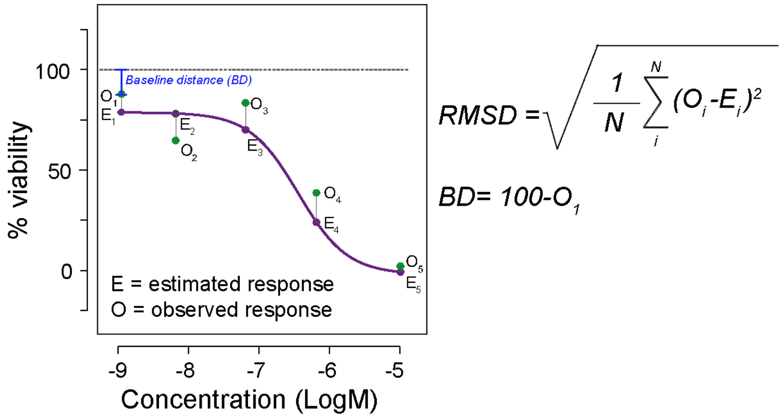


**Supplementary Figure 7**: Schematic illustration of the Root Mean Squared Distance (RMSD) and baseline distance (BD) calculation. $O_i$ and $E_i$ are the observed and estimated drug response values at concentration *i*.

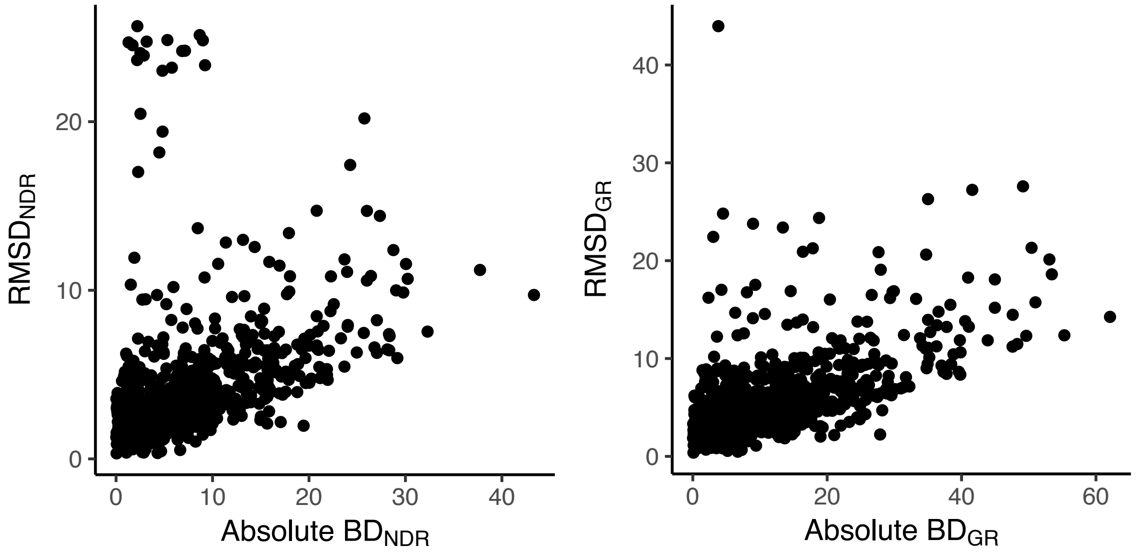


**Supplementary Figure 8**: Scatter plot showing the relationship between Baseline Distance (BD) and the root mean square distance (RMSD) between the observed and estimated drug response curves for the NDR and GR metrics. The Pearson correlation coefficient for GR metric was 0.58 and that for NDR was 0.36.

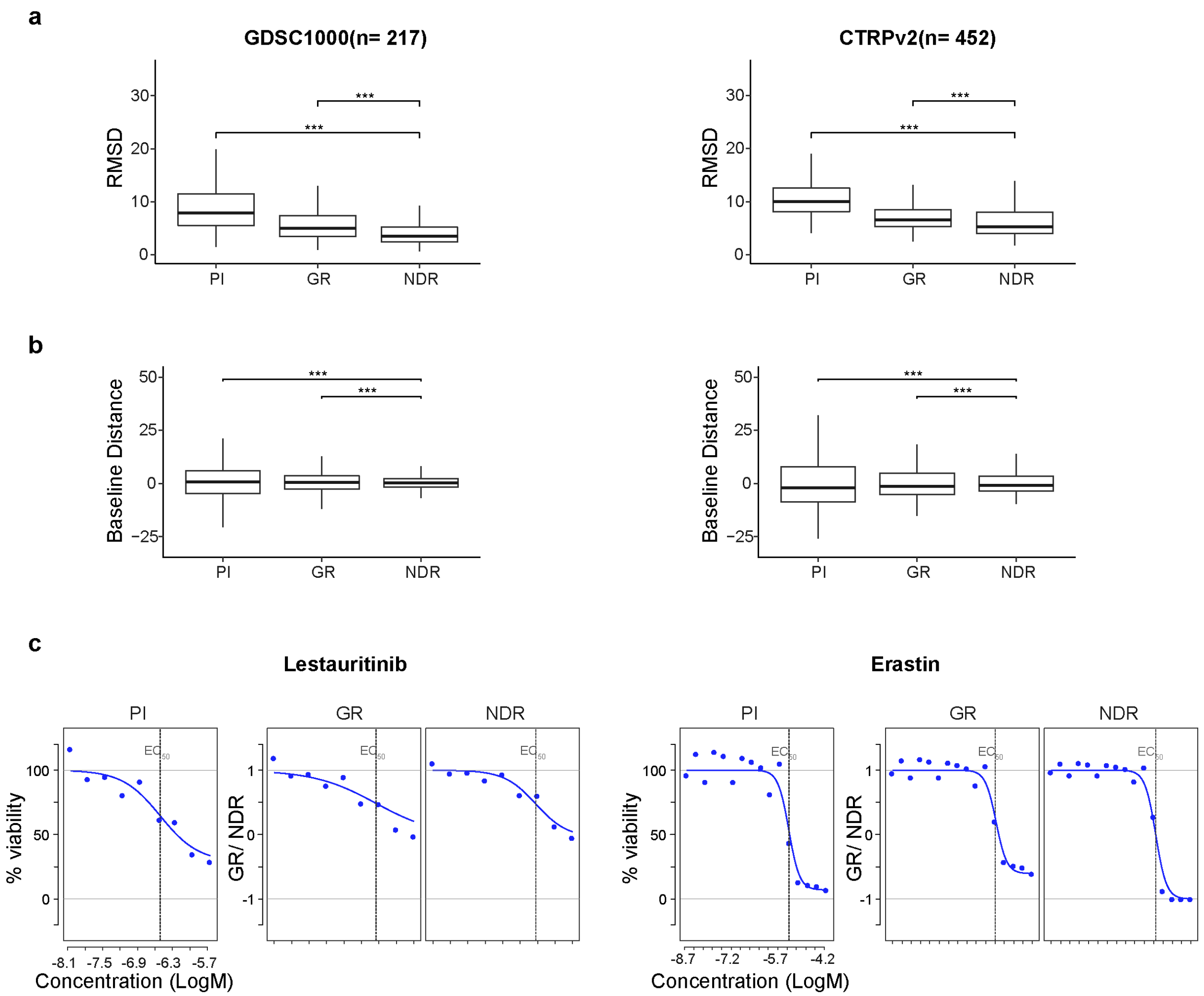


**Supplementary Figure 9** : Improved curve fitting using NDR in independent drug sensitivity datasets. (a) RMSD values computed between the estimated and observed dose-response curves obtained by applying the PI, GR and NDR metric in MDA-MB-231 cell line data from GDSC1000 (left) and CTRPv2 (right) datasets. Only drugs that showed non-zero values at all 5 concentrations for all the metrics were considered. ***p< 0.005; Wilcoxon rank sum test. (b) Baseline distance from zero computed at the lowest drug concentration using the PI, GR and NDR metric for the same set of samples. ***p< 0.005; F-test to compare the variances of two samples. (c) Dose-response curves obtained using PI, GR and NDR metric for a representative drug from each dataset showing the differences in curve fittings. These representative drugs illustrate both the improvement in curve-fittings as well as in baseline distances.

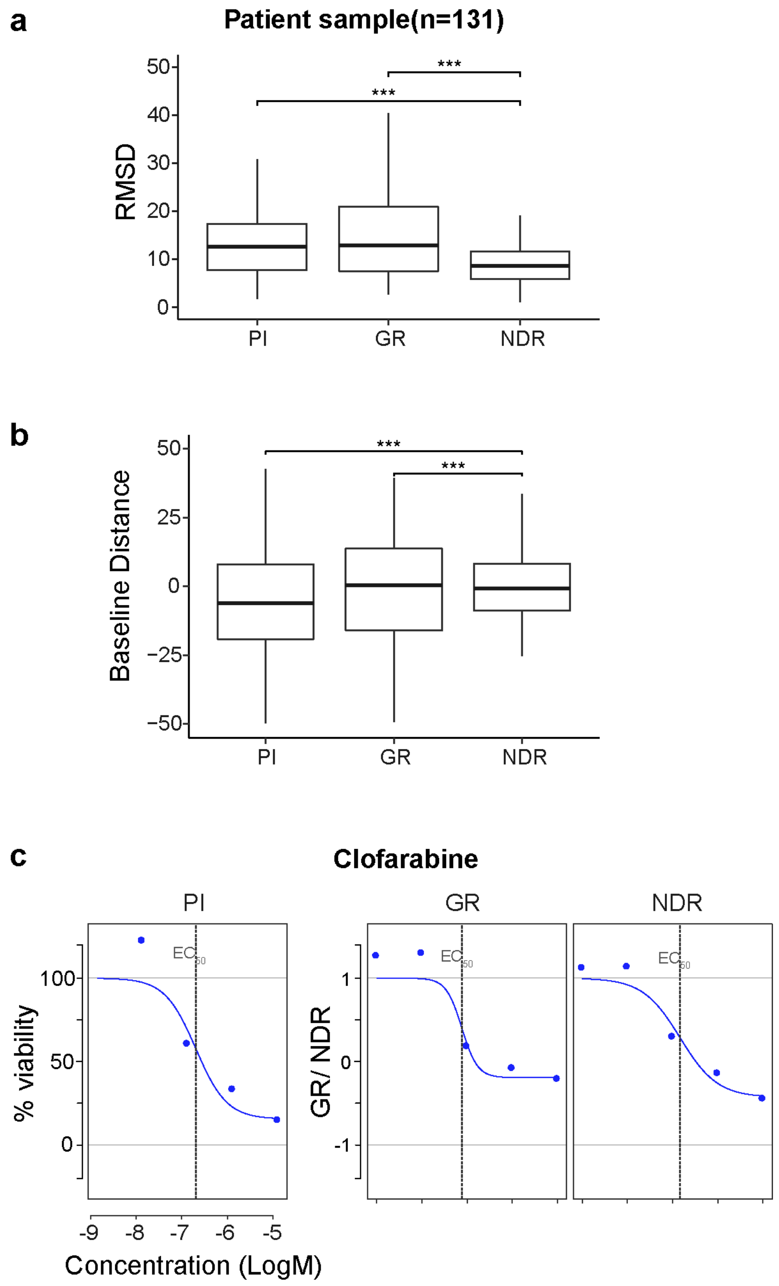


**Supplementary Figure 10**: Improved curve fitting using NDR in an AML patient sample. (a) RMSD values computed between the estimated and observed dose-response curves obtained by applying the PI, GR and NDR metric in a patient sample screened against 131 drugs. Only drugs that showed non-zero values at all 5 concentrations for all the metrics were considered. ***p< 0.005; Wilcoxon rank sum test. (b) Baseline distance from zero computed at the lowest drug concentration using the PI, GR and NDR metric for dose-response curves of the same samples. ***p< 0.005; F-test to compare the variances of two samples. (c) Dose-response curves obtained using PI, GR and NDR metric for a representative drug illustrating the improvement in curve-fittings as well as in baseline distances.

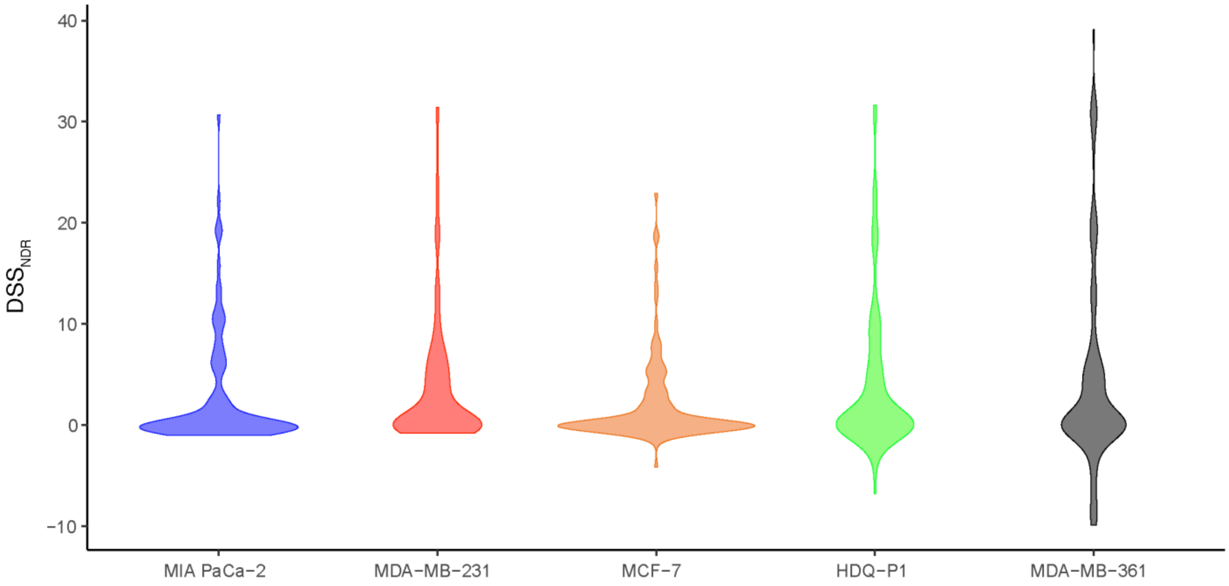


**Supplementary Figure 11**: Distribution of DSS$_{NDR}$ for 131 drugs in each of the 5 cell lines, in the ascending order of their doubling times.

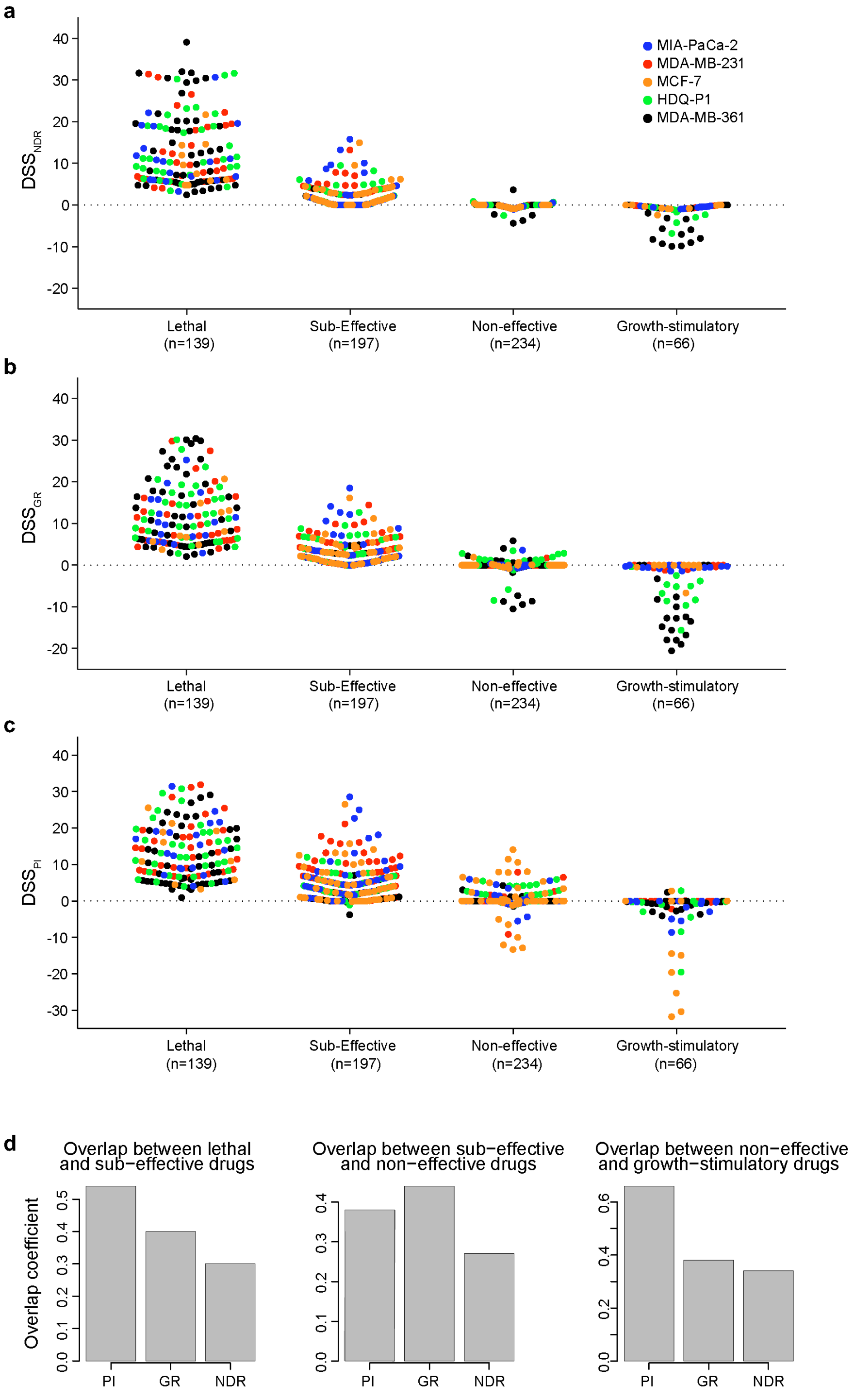


**Supplementary Figure 12**: Class-specific DSS values of drugs across all the cell lines. (a) $DSS_{NDR}$ distribution of drugs in the four classes. (b) $DSS_{GR}$ distribution of drugs in the four classes. (c) $DSS_{PI}$ distribution of drugs in the four classes. (d) Bar plot of the overlap coefficients between the distribution of DSS values for adjacent drug classes computed using NDR, GR and PI metric. The overlap is calculated in pairwise manner, namely, overlap coefficients between lethal and sub-effective drugs (left), overlap coefficients between sub-effective and non-effective drugs (middle), and overlap coefficients between non-effective and growth-proliferative drugs (right).
