## Supplementary Tables for "A normalized drug response metric improves accuracy and consistency of anticancer drug sensitivity quantification in cell-based screening": Supplementary_File_1.pdf

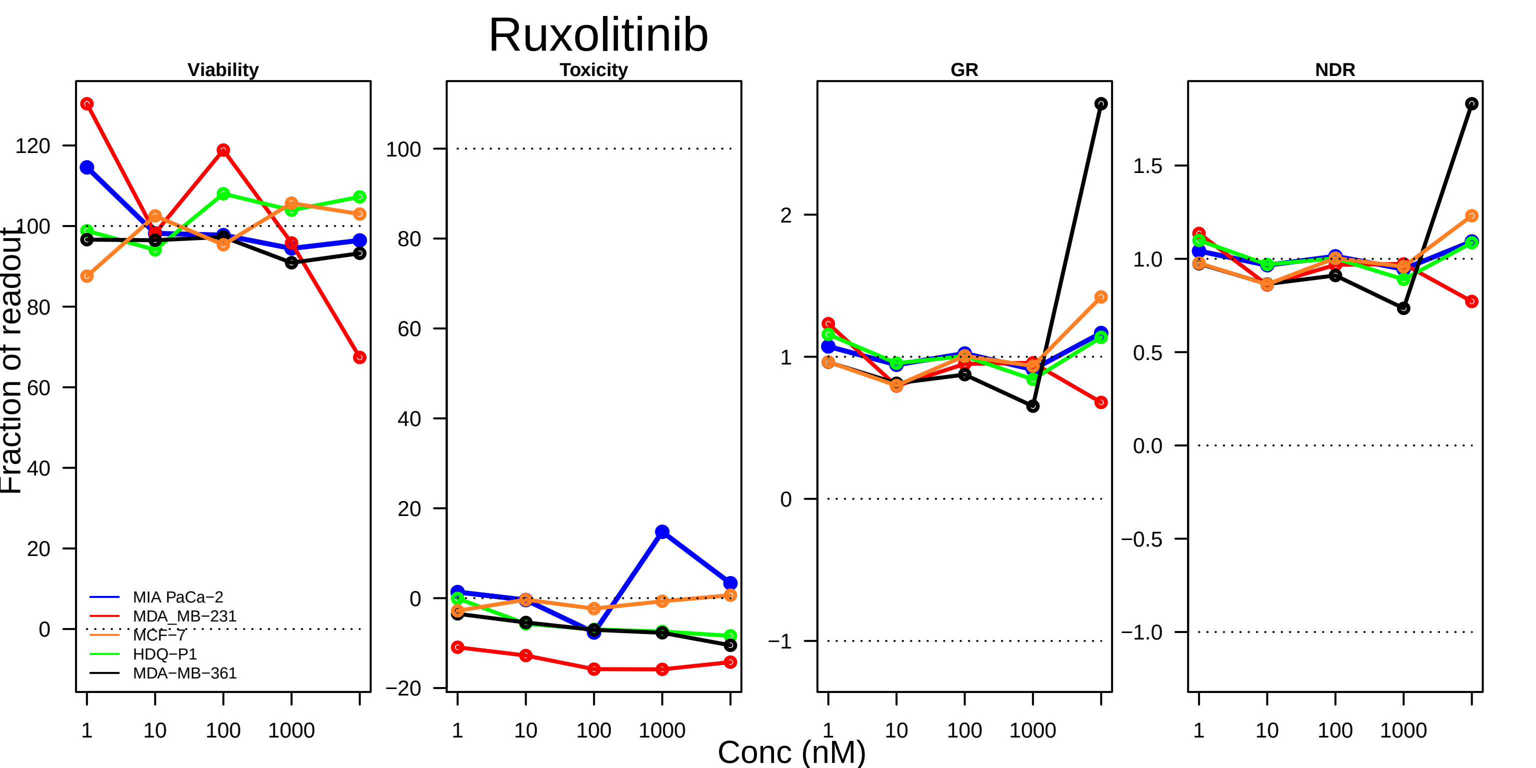
Ruxolitinib
Viability
Toxicity
GR
NDR
Fraction of readout
Conc (nM)
MIA PaCa−2
MDA_MB−231
MCF−7
HDQ−P1
MDA−MB−361
1
10
100
1000

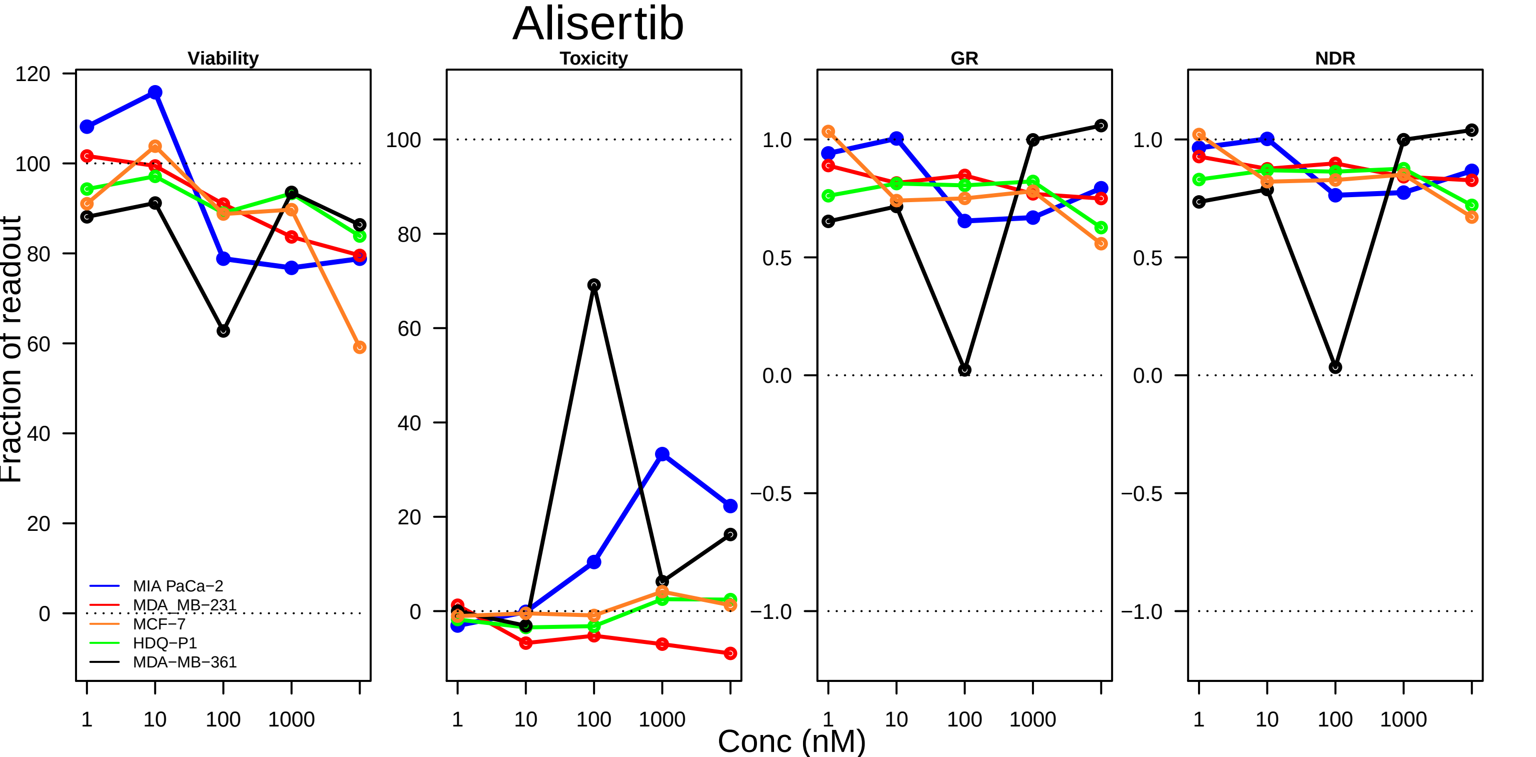

Alisertib
Viability
Toxicity
GR
NDR
Fraction of readout
Conc (nM)
1
10
100
1000
120
100
80
60
40
20
0
1.0
0.5
0.0
−0.5
−1.0
MIA PaCa−2
MDA_MB−231
MCF−7
HDQ−P1
MDA−MB−361

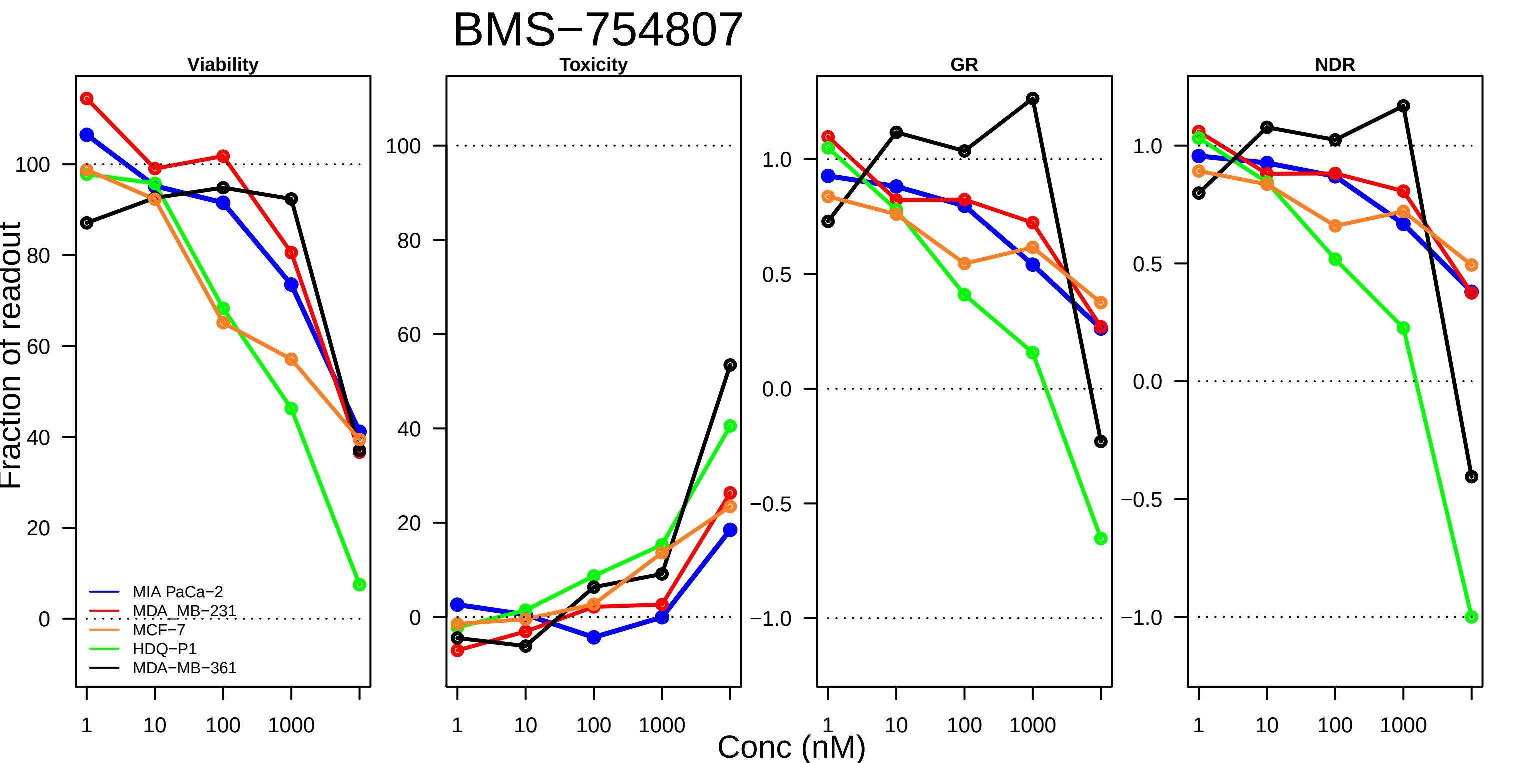

BMS−754807
Viability
Toxicity
GR
NDR
Fraction of readout
Conc (nM)
MIA PaCa−2
MDA_MB−231
MCF−7
HDQ−P1
MDA−MB−361

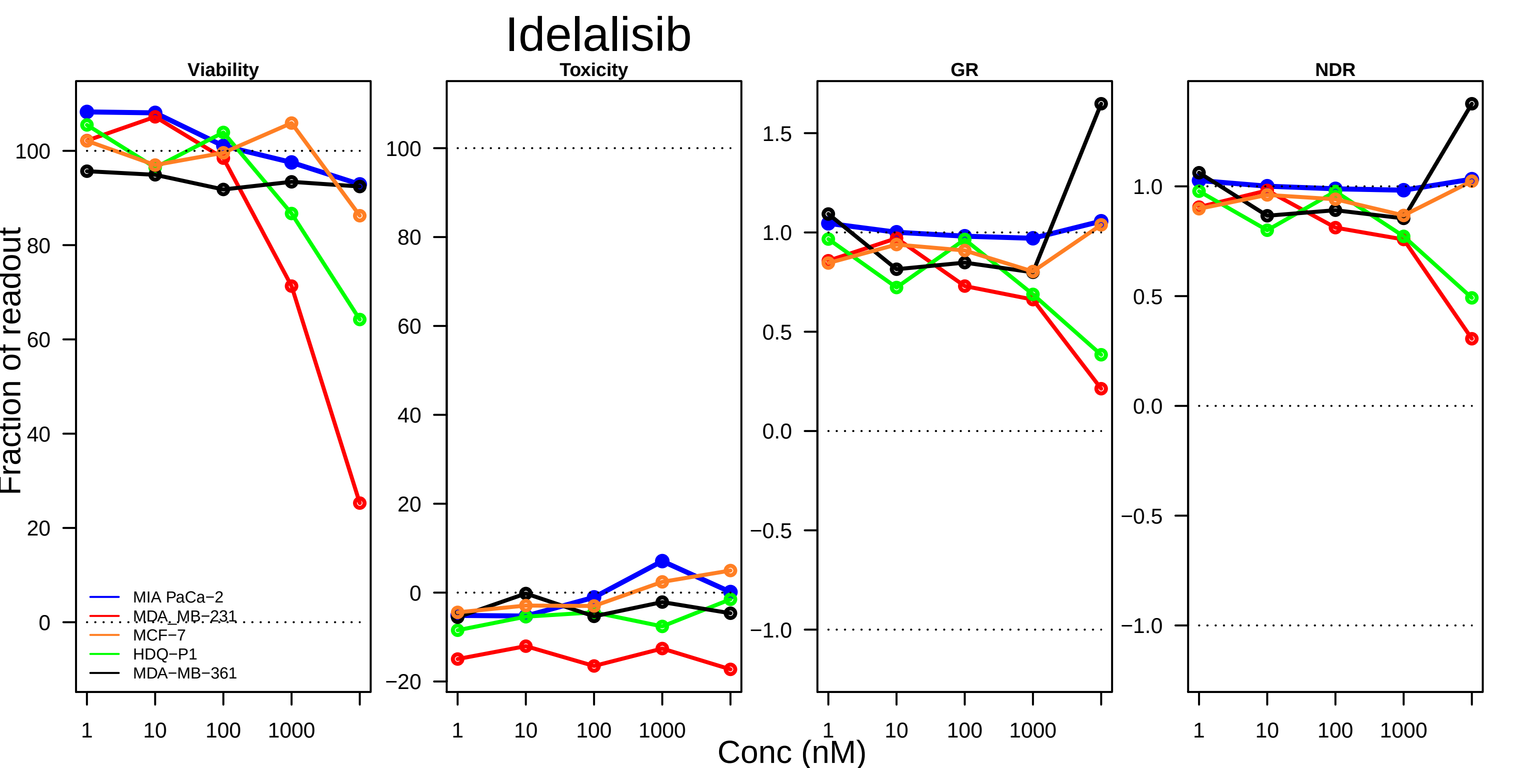

Idelalisib
Viability
Toxicity
GR
NDR
Fraction of readout
Conc (nM)
1
10
100
1000
MIA PaCa−2
MDA_MB−231
MCF−7
HDQ−P1
MDA−MB−361

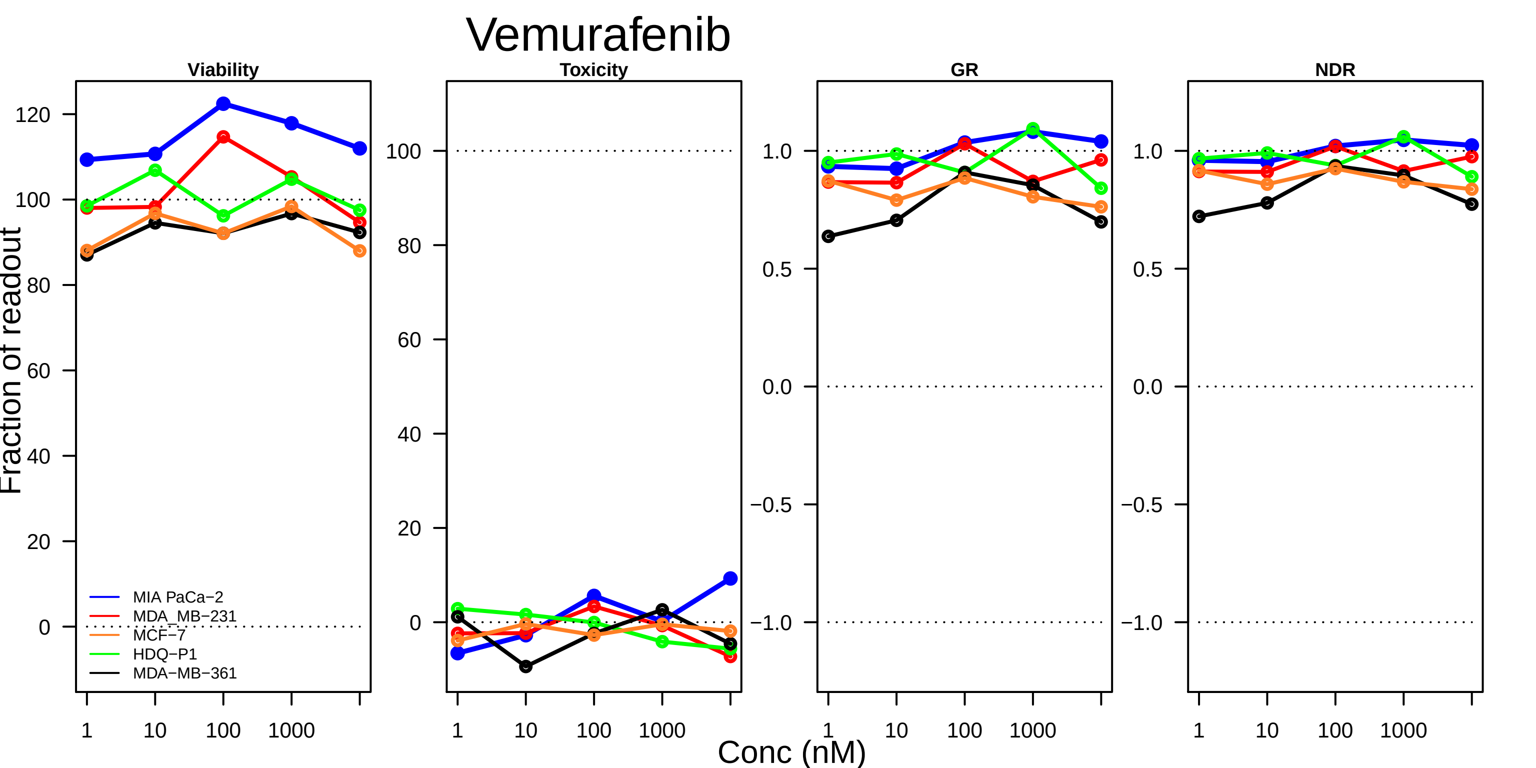
Vemurafenib
Viability
Toxicity
GR
NDR
Fraction of readout
120
100
80
60
40
20
0
1.0
0.5
0.0
−0.5
−1.0
MIA PaCa−2
MDA_MB−231
MCF−7
HDQ−P1
MDA−MB−361
1
10
100
1000
Conc (nM)

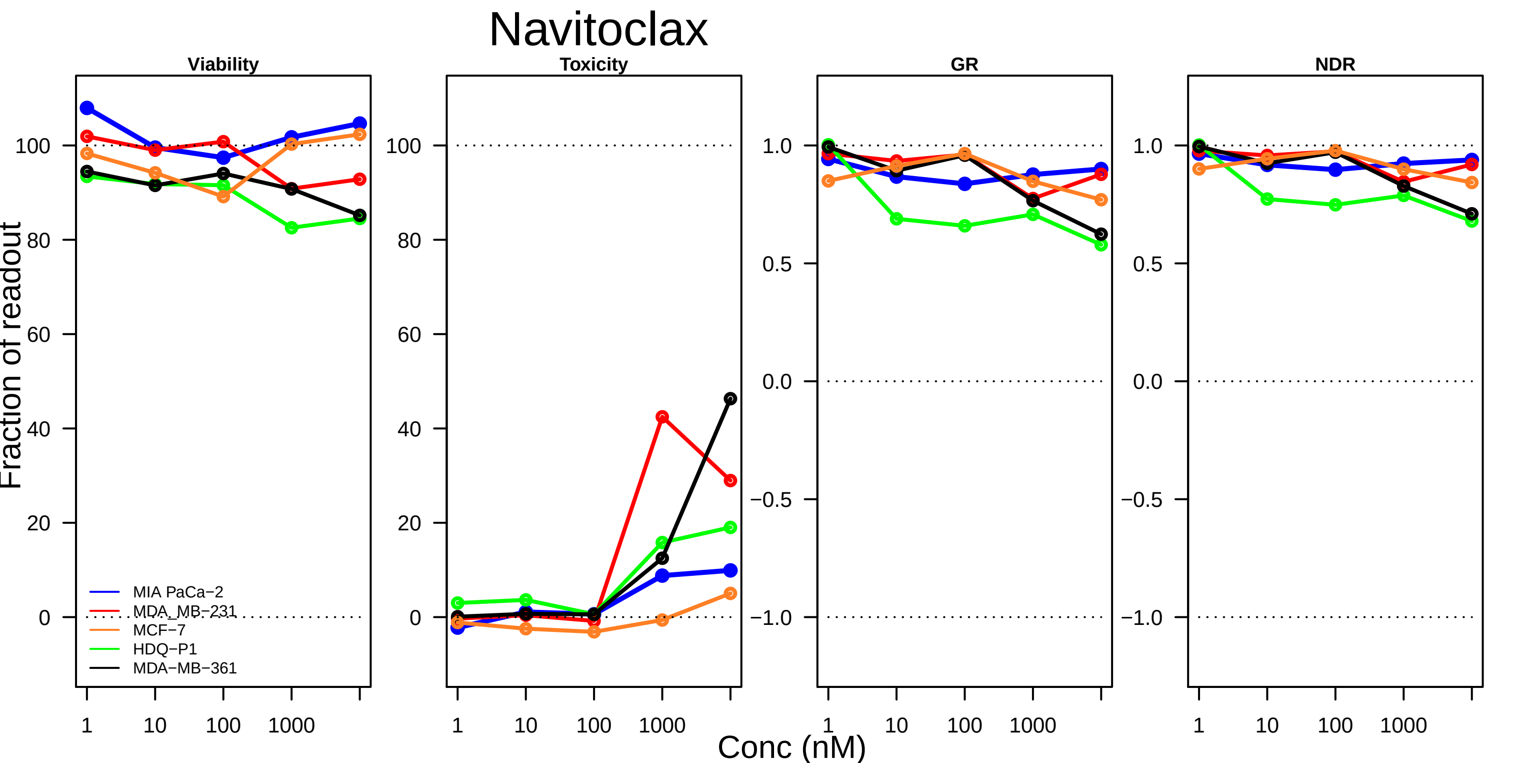
Navitoclax
Viability
Toxicity
GR
NDR
Fraction of readout
Conc (nM)
MIA PaCa−2
MDA_MB−231
MCF−7
HDQ−P1
MDA−MB−361
100
80
60
40
20
0
1.0
0.5
0.0
−0.5
−1.0
1
10
100
1000

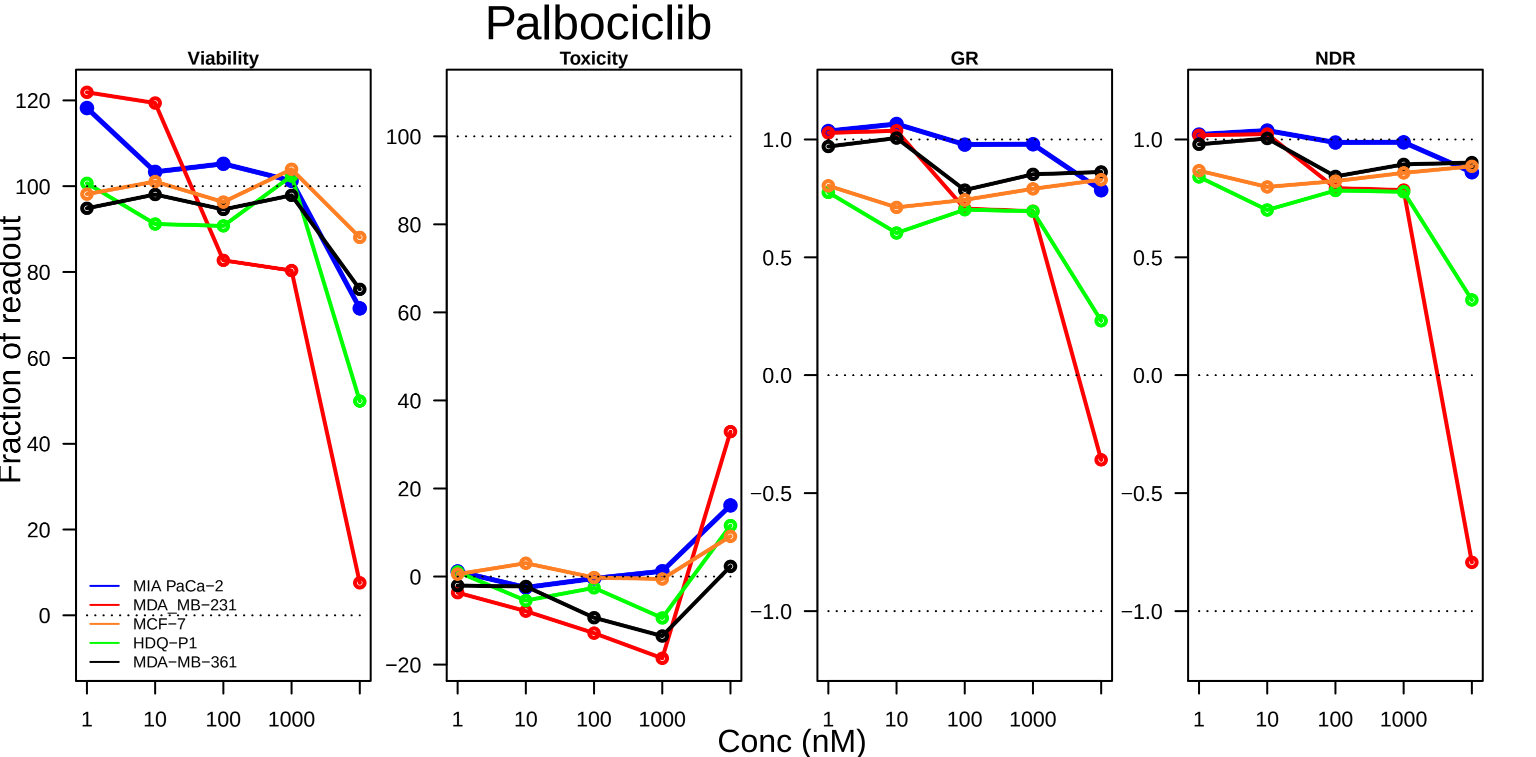
Palbociclib
Viability
Toxicity
GR
NDR
Fraction of readout
Conc (nM)
1
10
100
1000
120
100
80
60
40
20
0
−20
1.0
0.5
0.0
−0.5
−1.0
MIA PaCa−2
MDA_MB−231
MCF−7
HDQ−P1
MDA−MB−361

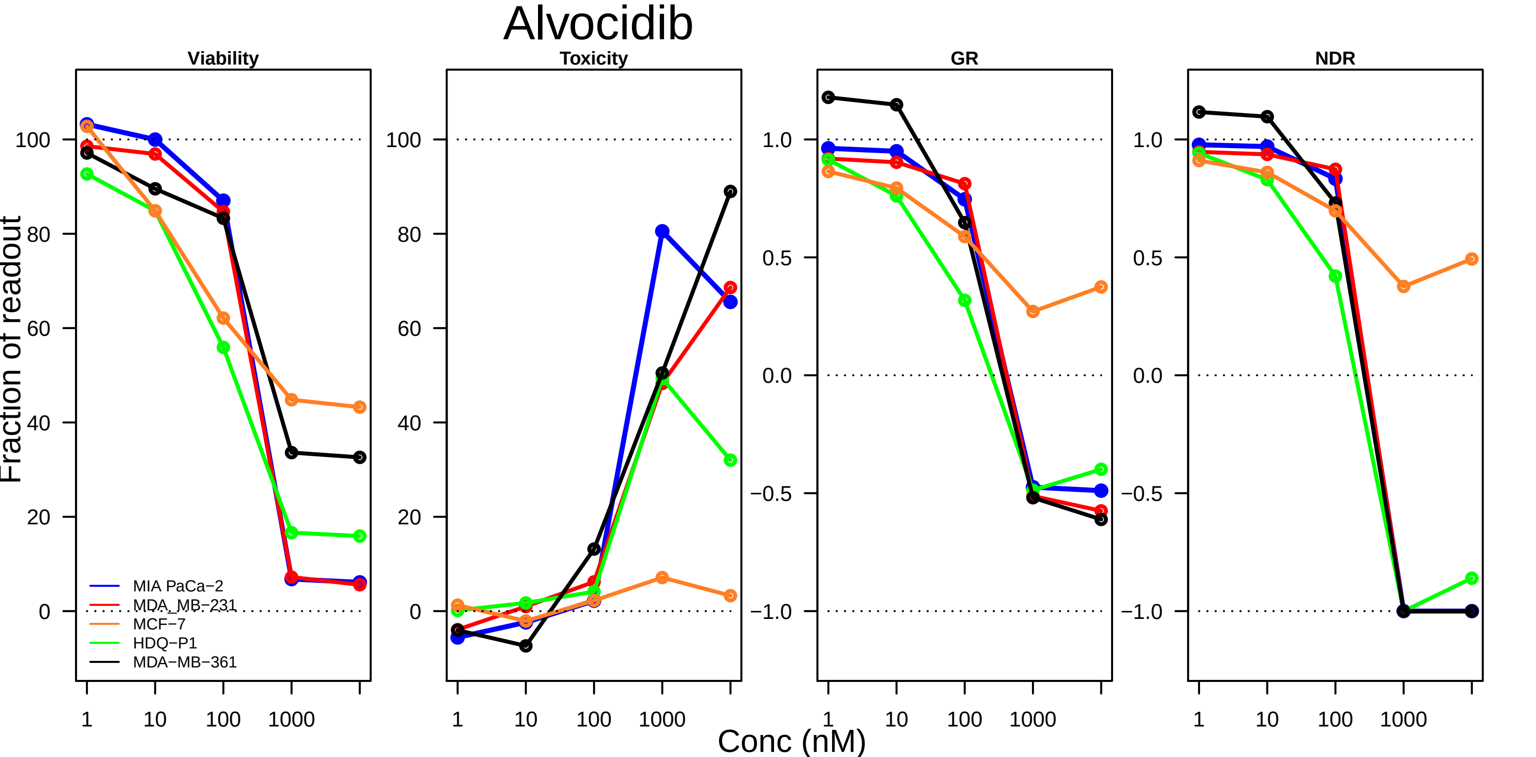

Alvocidib
Viability
Toxicity
GR
NDR
Fraction of readout
Conc (nM)
1
10
100
1000
0
20
40
60
80
100
−1.0
−0.5
0.0
0.5
1.0
MIA PaCa−2
MDA_MB−231
MCF−7
HDQ−P1
MDA−MB−361

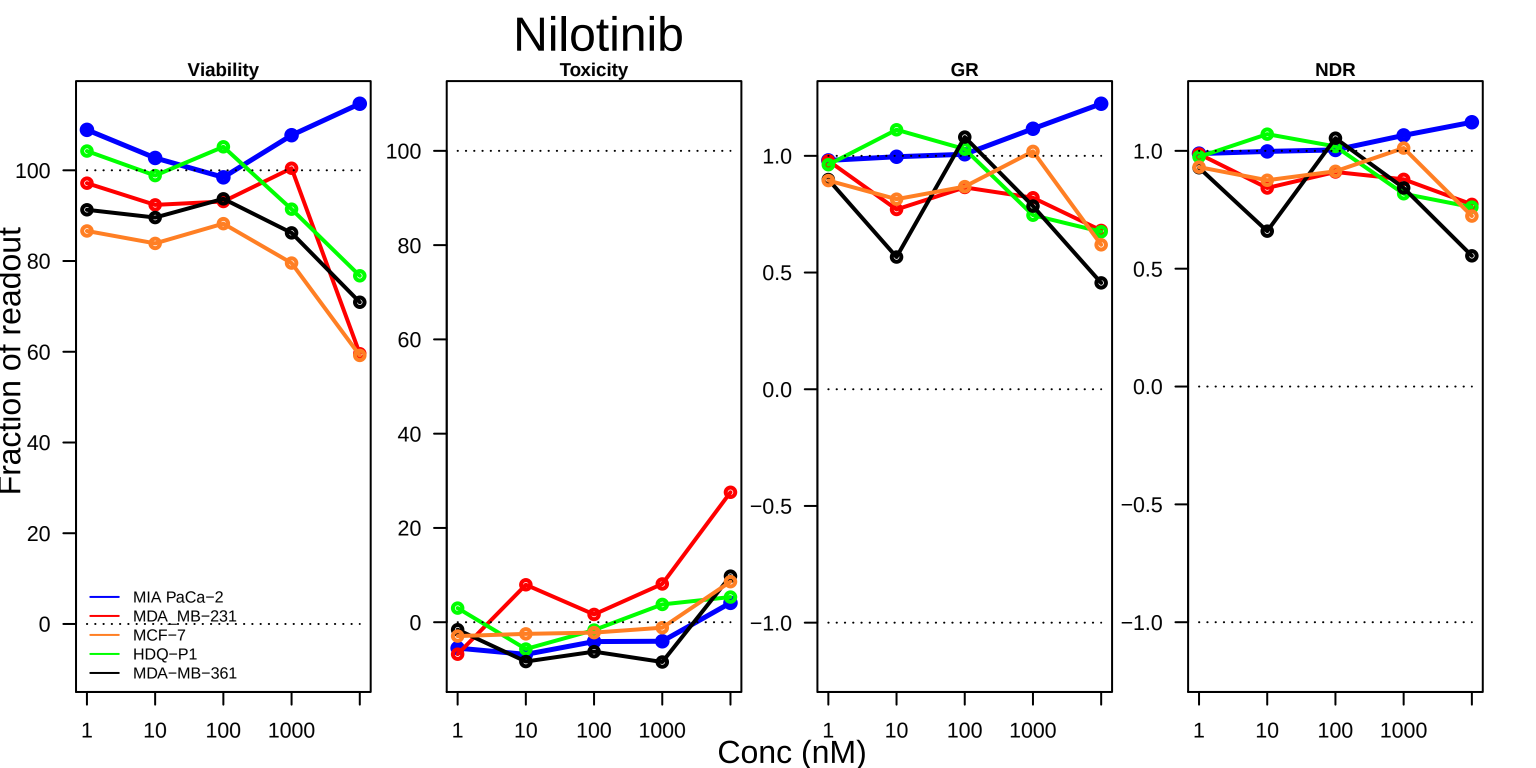

Nilotinib
Viability
Toxicity
GR
NDR
Fraction of readout
Conc (nM)
1
10
100
1000
MIA PaCa−2
MDA_MB−231
MCF−7
HDQ−P1
MDA−MB−361
0
20
40
60
80
100
0.0
0.5
1.0
−0.5
−1.0

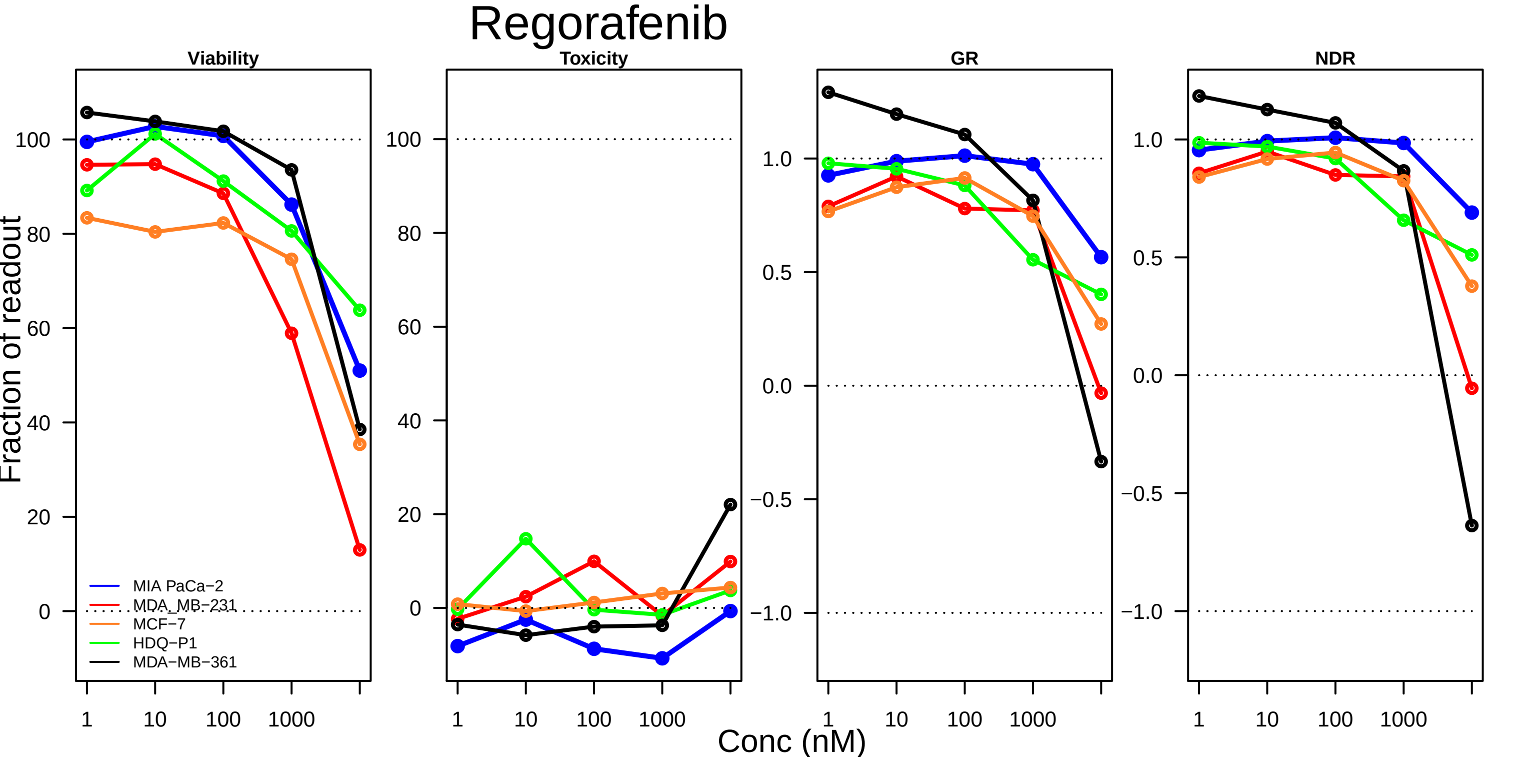
Regorafenib
Viability
Toxicity
GR
NDR
Fraction of readout
100
80
60
40
20
0
1.0
0.5
0.0
−0.5
−1.0
1
10
100
1000
MIA PaCa−2
MDA_MB−231
MCF−7
HDQ−P1
MDA−MB−361
Conc (nM)

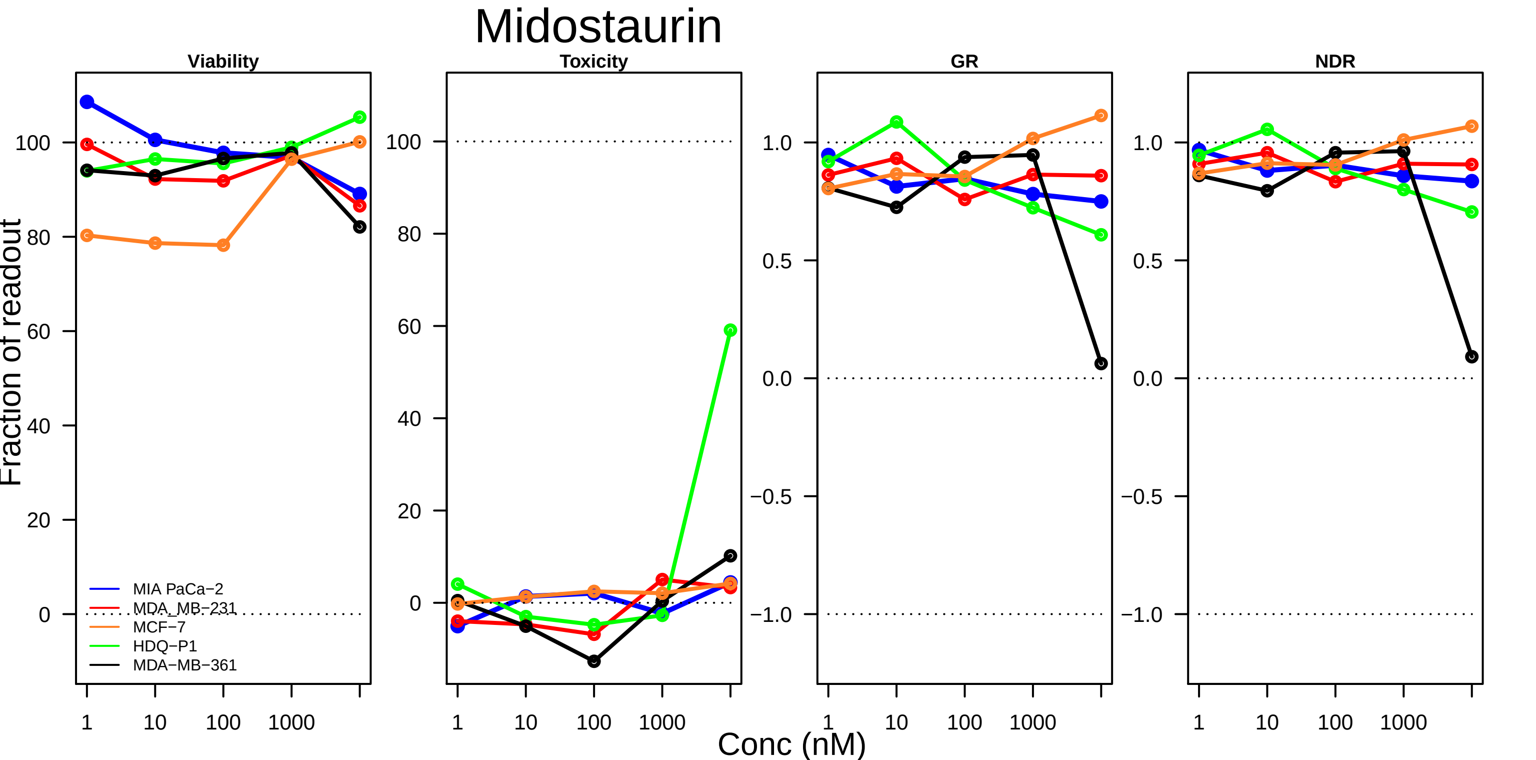
Midostaurin
Viability
Toxicity
GR
NDR
Fraction of readout
100
80
60
40
20
0
1.0
0.5
0.0
−0.5
−1.0
1
10
100
1000
MIA PaCa−2
MDA_MB−231
MCF−7
HDQ−P1
MDA−MB−361
Conc (nM)

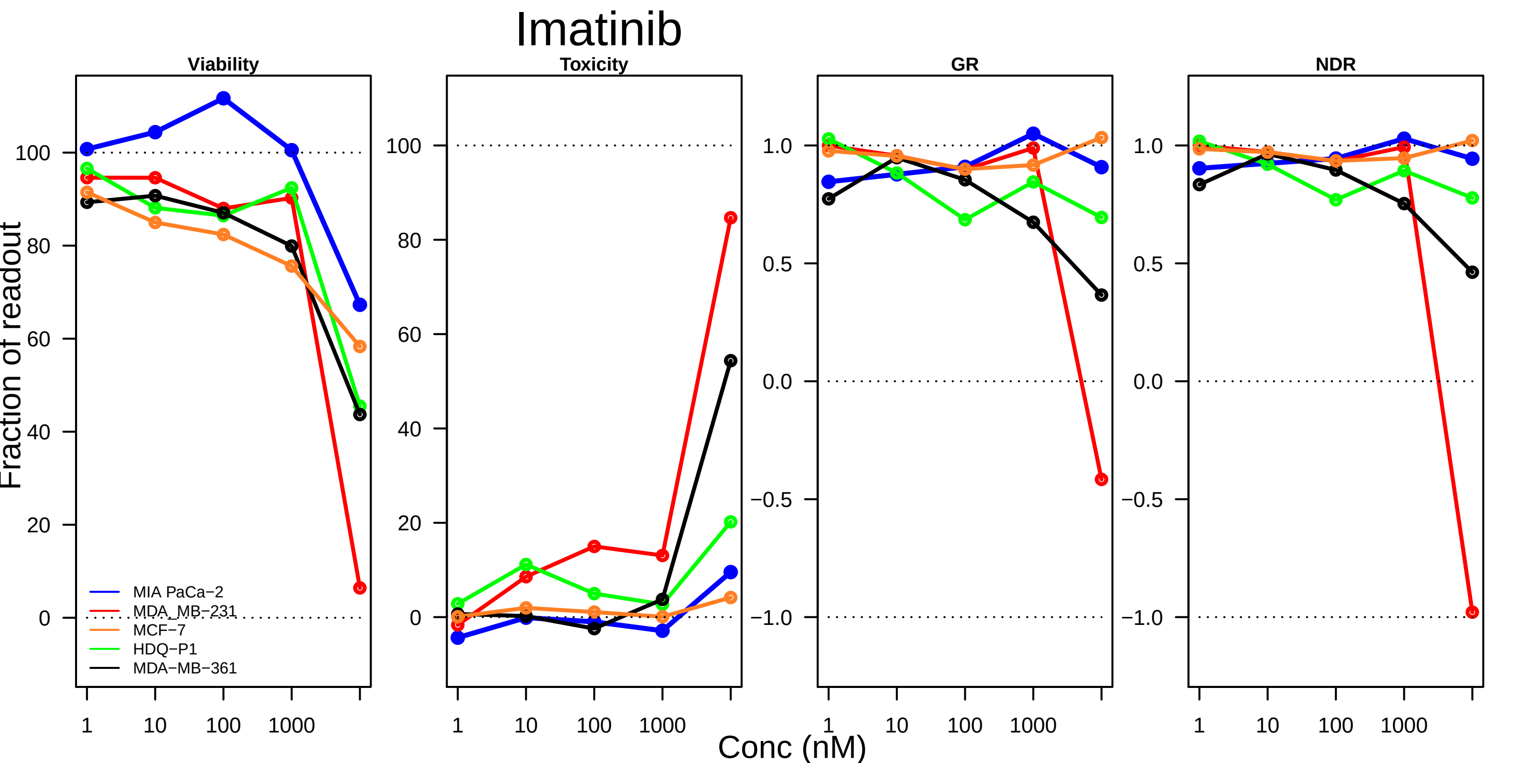

Imatinib
Viability
Toxicity
GR
NDR
Fraction of readout
Conc (nM)
MIA PaCa−2
MDA_MB−231
MCF−7
HDQ−P1
MDA−MB−361
1
10
100
1000

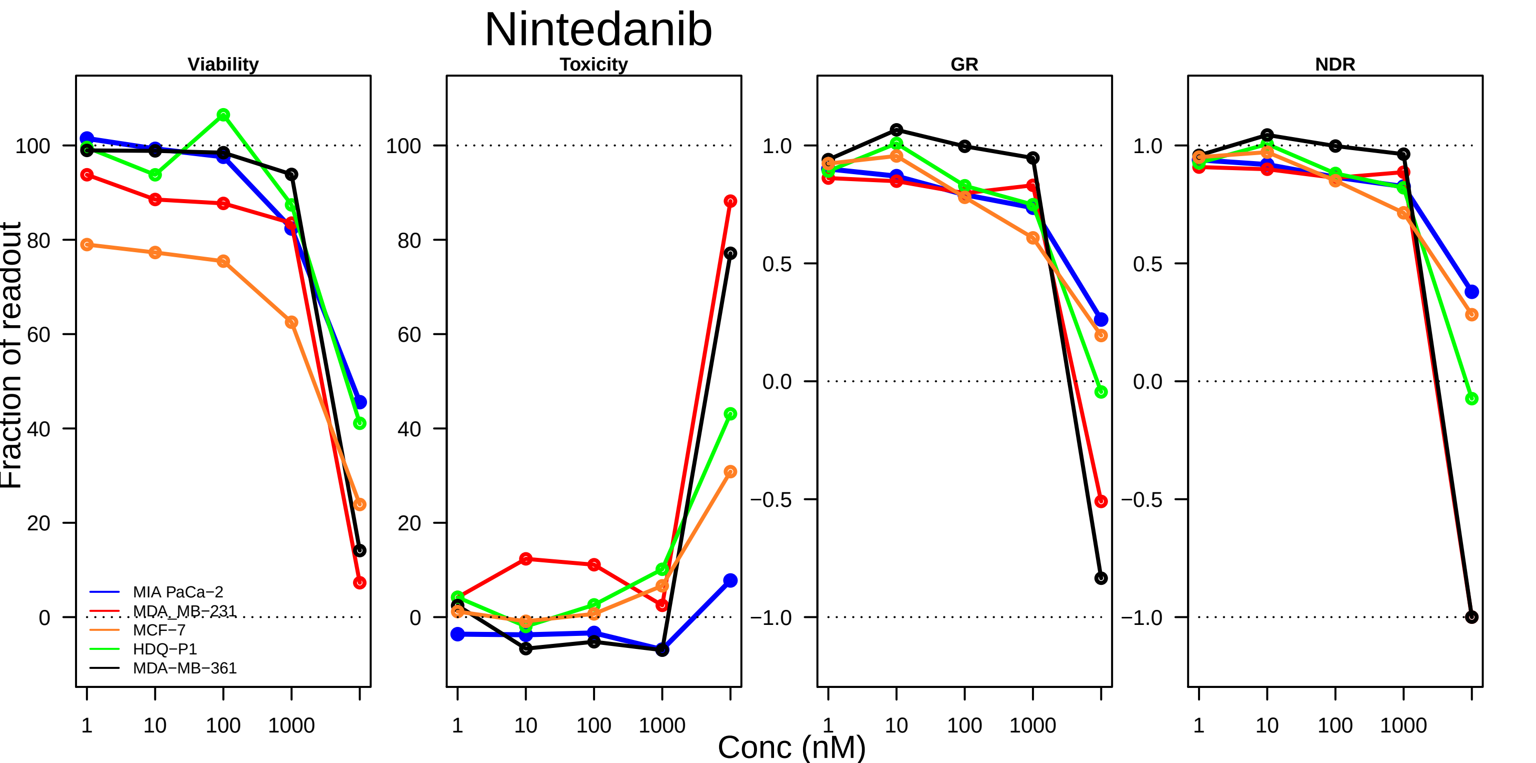
Nintedanib
Viability
Toxicity
GR
NDR
Fraction of readout
Conc (nM)
MIA PaCa−2
MDA_MB−231
MCF−7
HDQ−P1
MDA−MB−361
100
80
60
40
20
0
1.0
0.5
0.0
−0.5
−1.0
1
10
100
1000

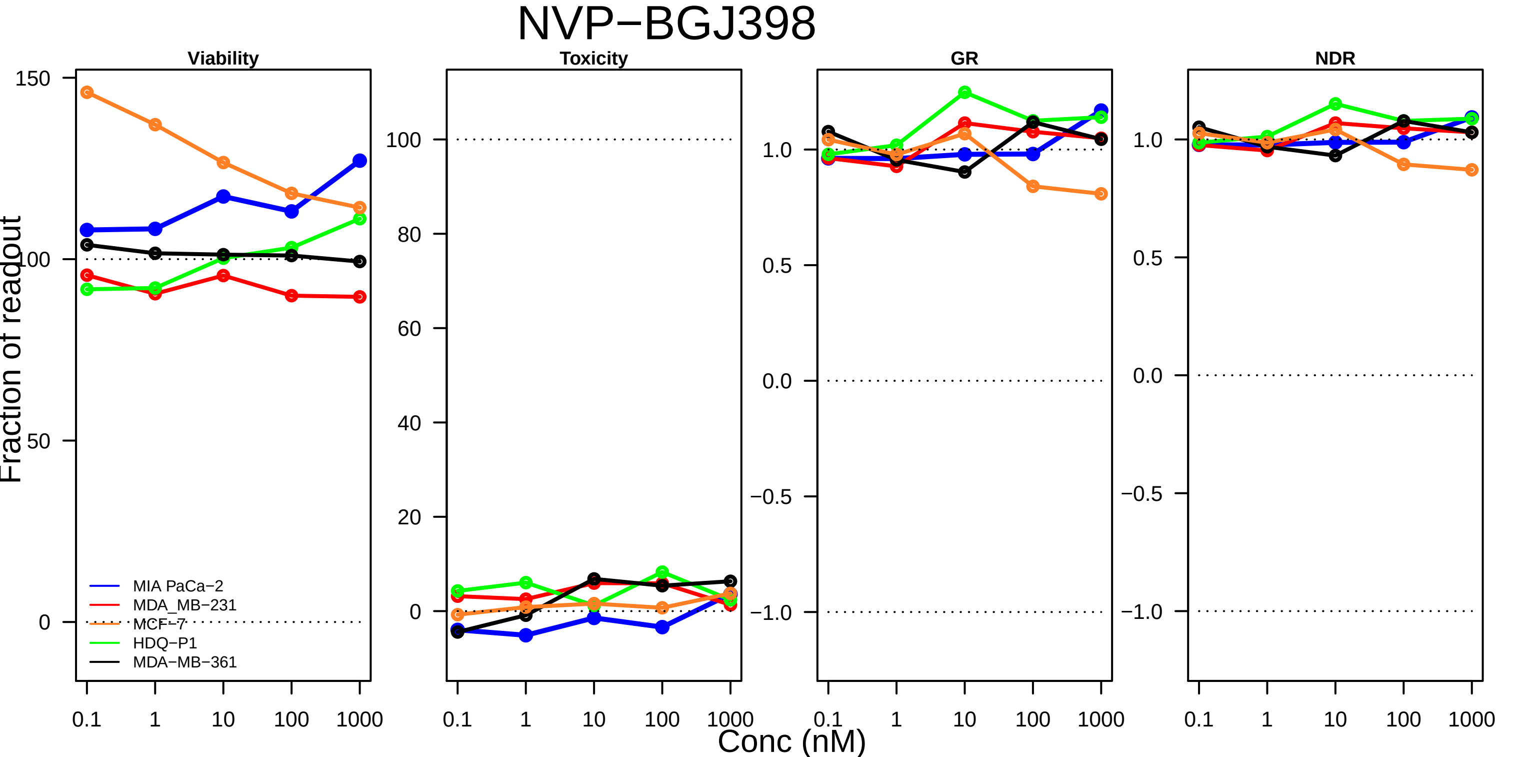

NVP−BGJ398
Viability
Toxicity
GR
NDR
Fraction of readout
Conc (nM)
150
100
50
0
80
60
40
20
1.0
0.5
0.0
−0.5
−1.0
0.1
1
10
1000
MIA PaCa−2
MDA_MB−231
MCF−7
HDQ−P1
MDA−MB−361

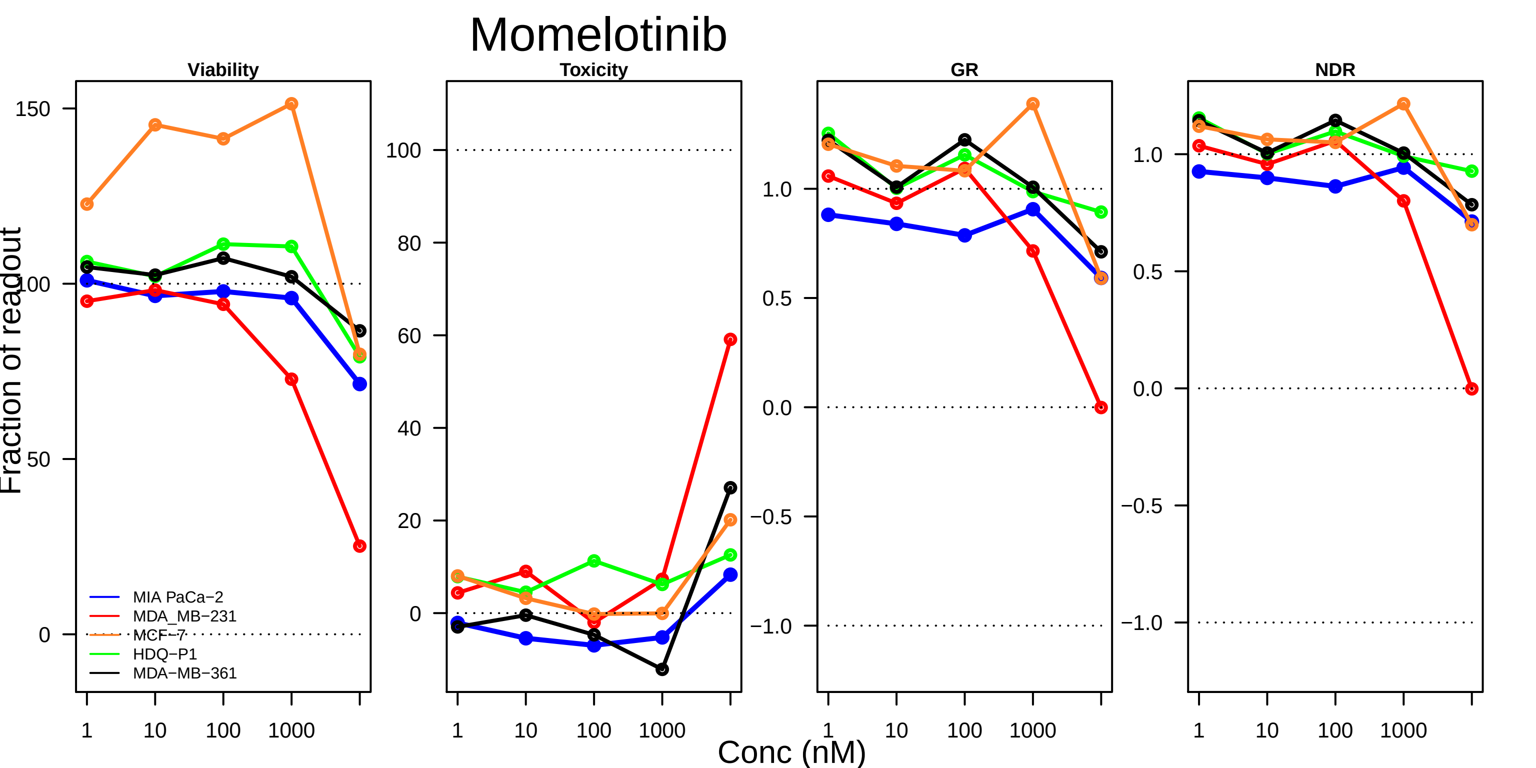

Momelotinib
Viability
Toxicity
GR
NDR
Fraction of readout
Conc (nM)
1
10
100
1000
150
100
50
0
80
60
40
20
1.0
0.5
0.0
−0.5
−1.0
MIA PaCa−2
MDA_MB−231
MCF−7
HDQ−P1
MDA−MB−361

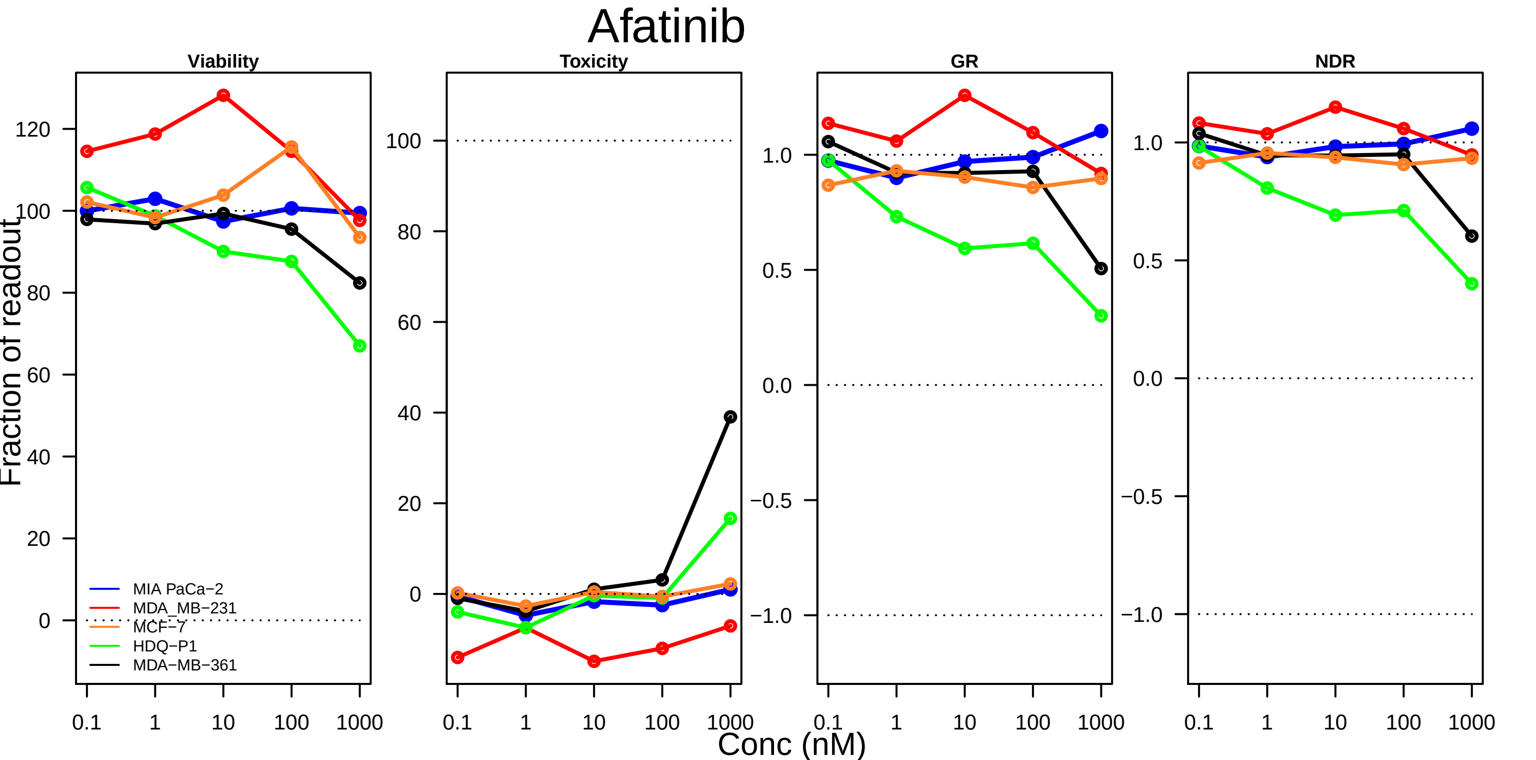

Afatinib
Viability
Toxicity
GR
NDR
Fraction of readout
Conc (nM)
0.1
1
10
100
1000
120
100
80
60
40
20
0
−0.5
−1.0
0.0
0.5
1.0
MIA PaCa−2
MDA_MB−231
MCF−7
HDQ−P1
MDA−MB−361

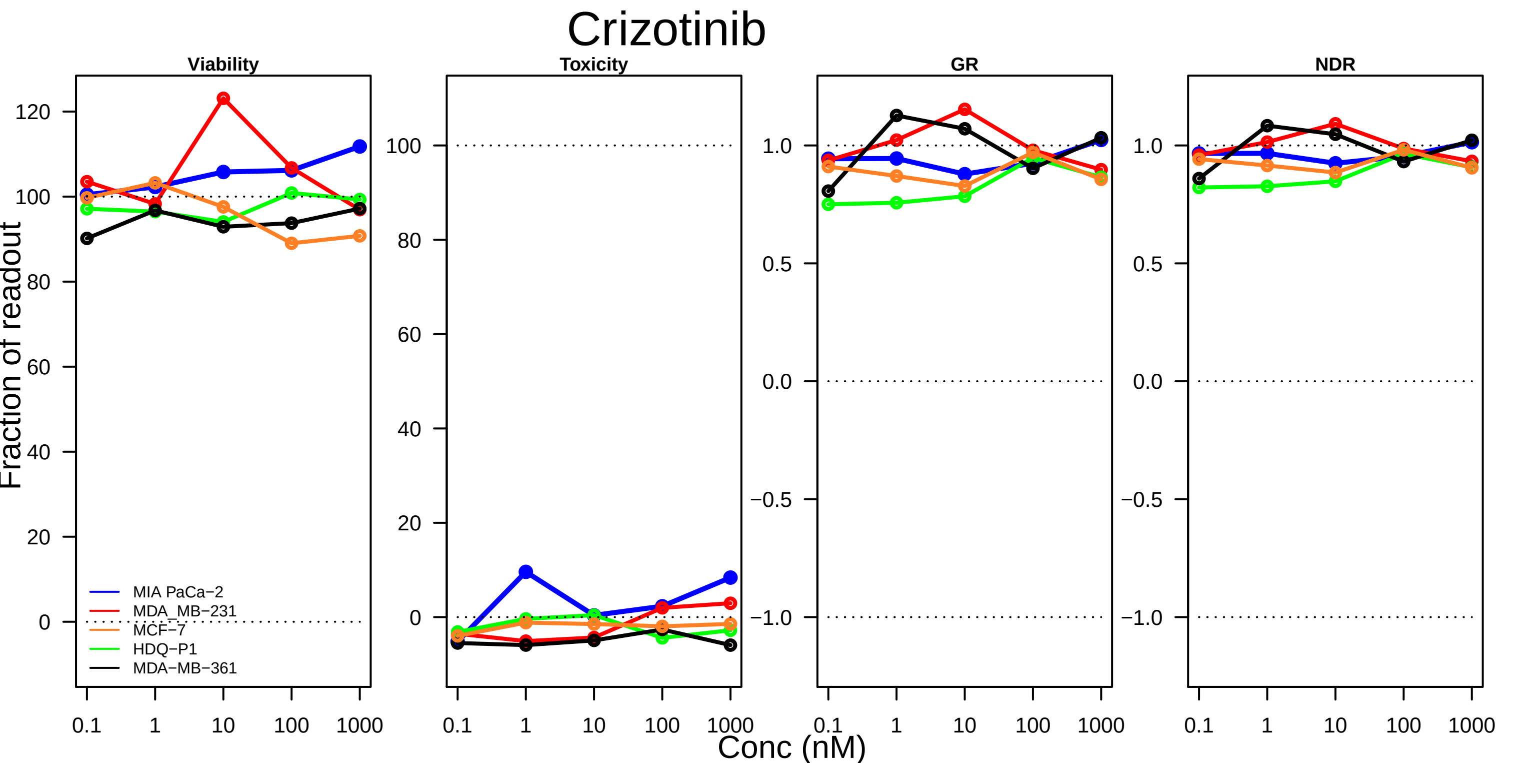
Crizotinib
Viability
Toxicity
GR
NDR
Fraction of readout
Conc (nM)
MIA PaCa−2
MDA_MB−231
MCF−7
HDQ−P1
MDA−MB−361
0.1
1
10
100
1000

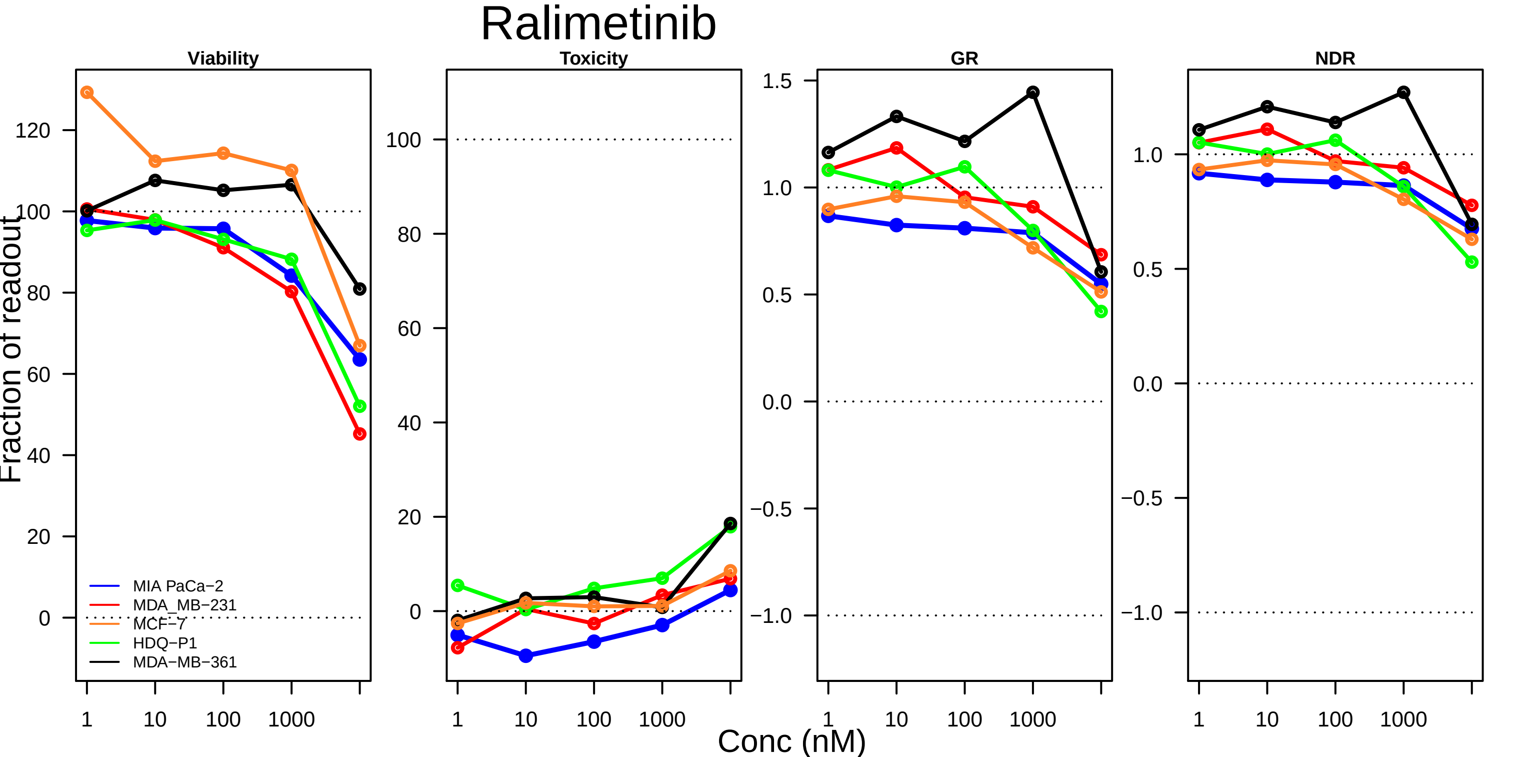

Ralimetinib
Viability
Toxicity
GR
NDR
Fraction of readout
Conc (nM)
1
10
100
1000
MIA PaCa−2
MDA_MB−231
MCF−7
HDQ−P1
MDA−MB−361

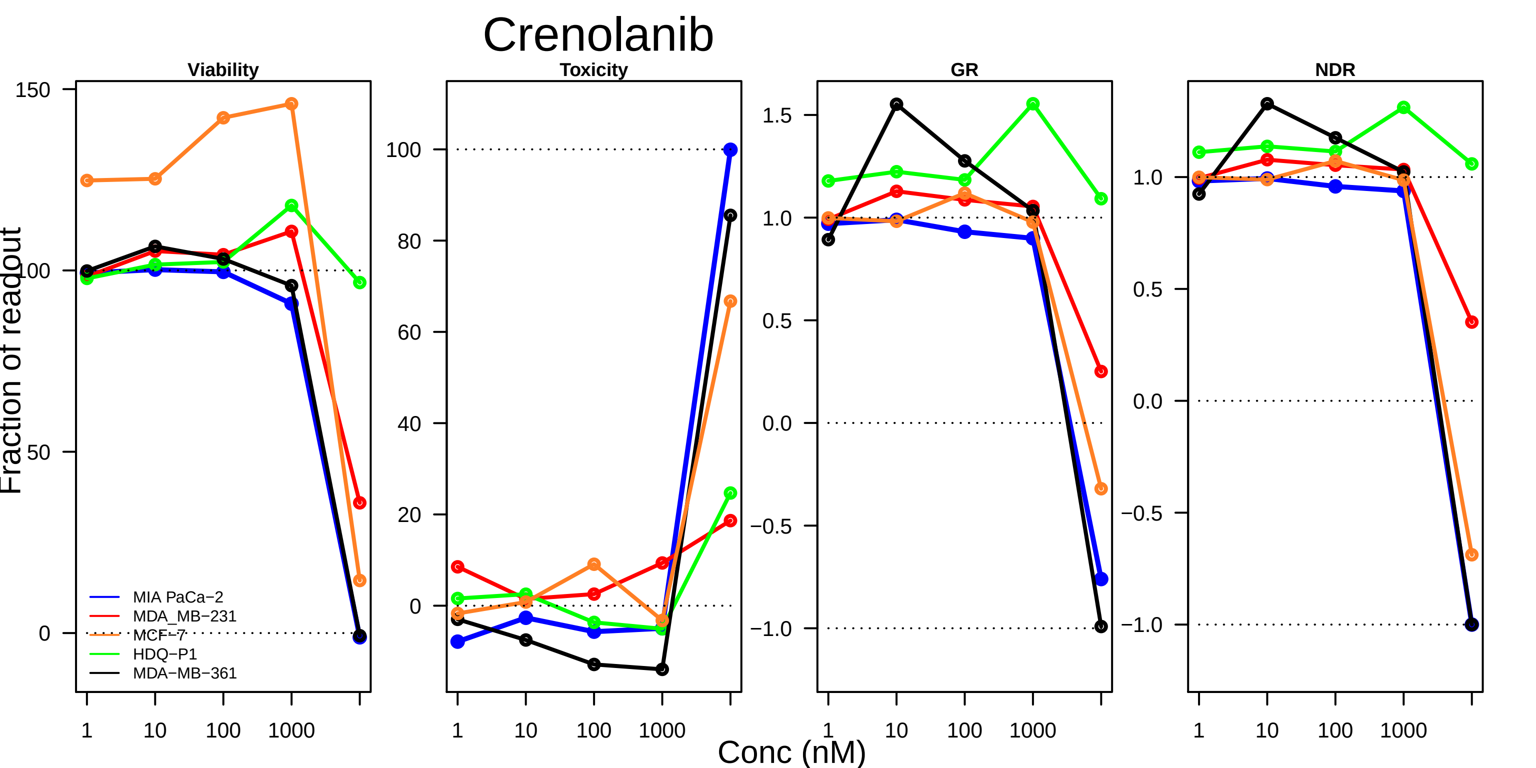

Crenolanib
Viability
Toxicity
GR
NDR
Fraction of readout
Conc (nM)
MIA PaCa−2
MDA_MB−231
MCF−7
HDQ−P1
MDA−MB−361
150
100
50
0
80
60
40
20
1.5
1.0
0.5
0.0
−0.5
−1.0
1
10
100
1000

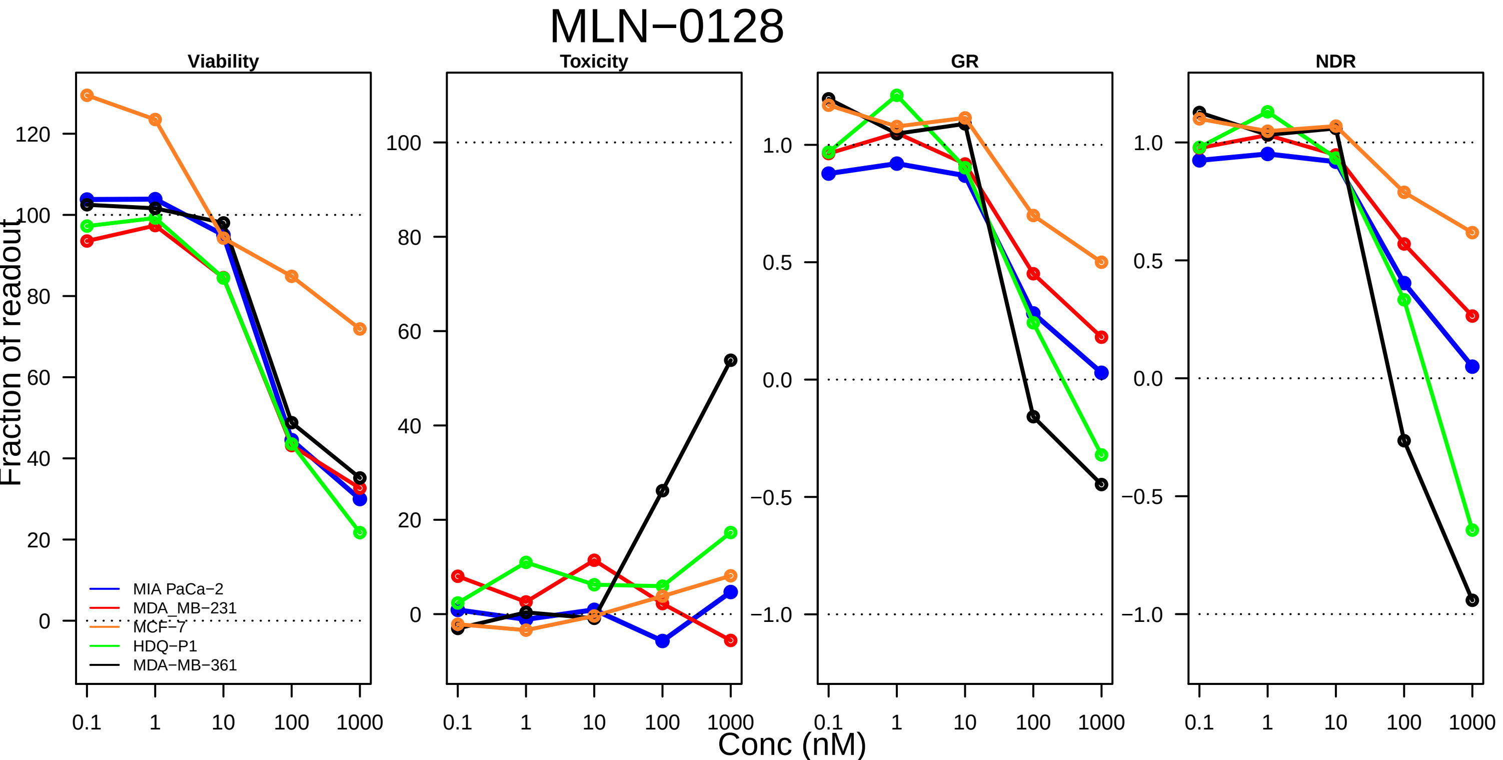

MLN−0128
Viability
Toxicity
GR
NDR
Fraction of readout
Conc (nM)
0.1
1
10
100
1000
MIA PaCa−2
MDA_MB−231
MCF−7
HDQ−P1
MDA−MB−361

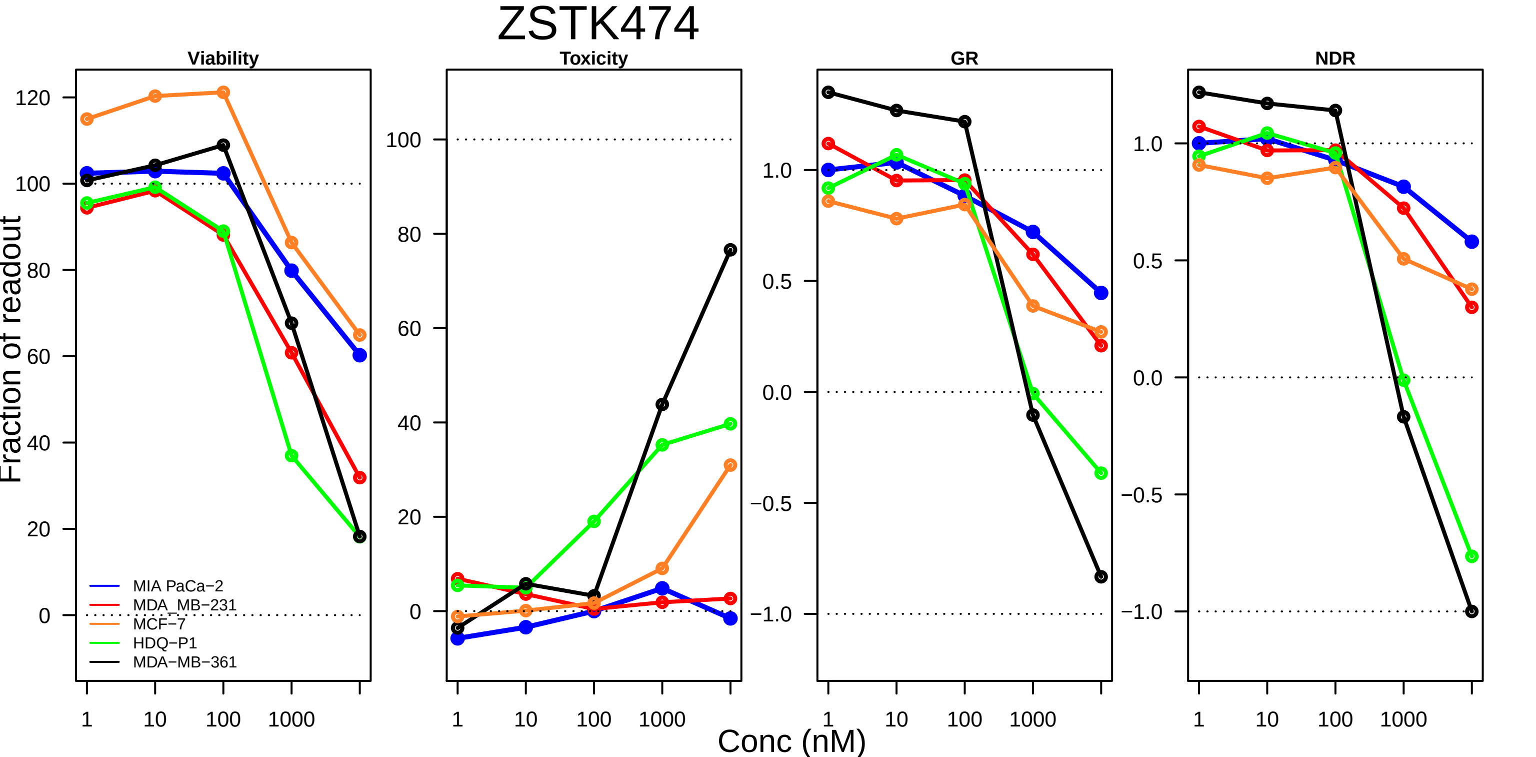

ZSTK474
Viability
Toxicity
GR
NDR
Fraction of readout
Conc (nM)
MIA PaCa−2
MDA_MB−231
MCF−7
HDQ−P1
MDA−MB−361
1
10
100
1000

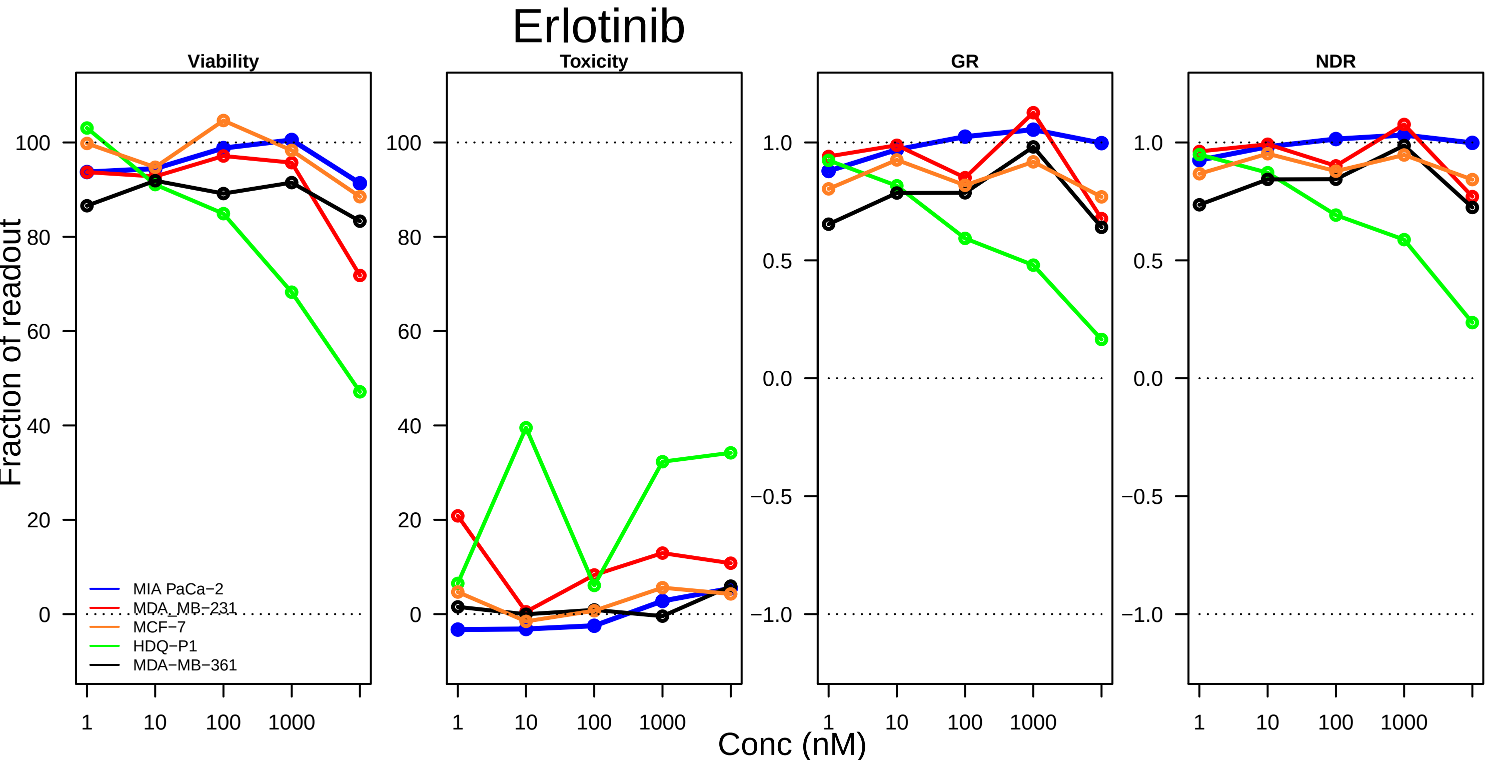

Erlotinib
Viability
Toxicity
GR
NDR
Fraction of readout
Conc (nM)
100
80
60
40
20
0
1.0
0.5
0.0
−0.5
−1.0
1
10
100
1000
MIA PaCa−2
MDA_MB−231
MCF−7
HDQ−P1
MDA−MB−361

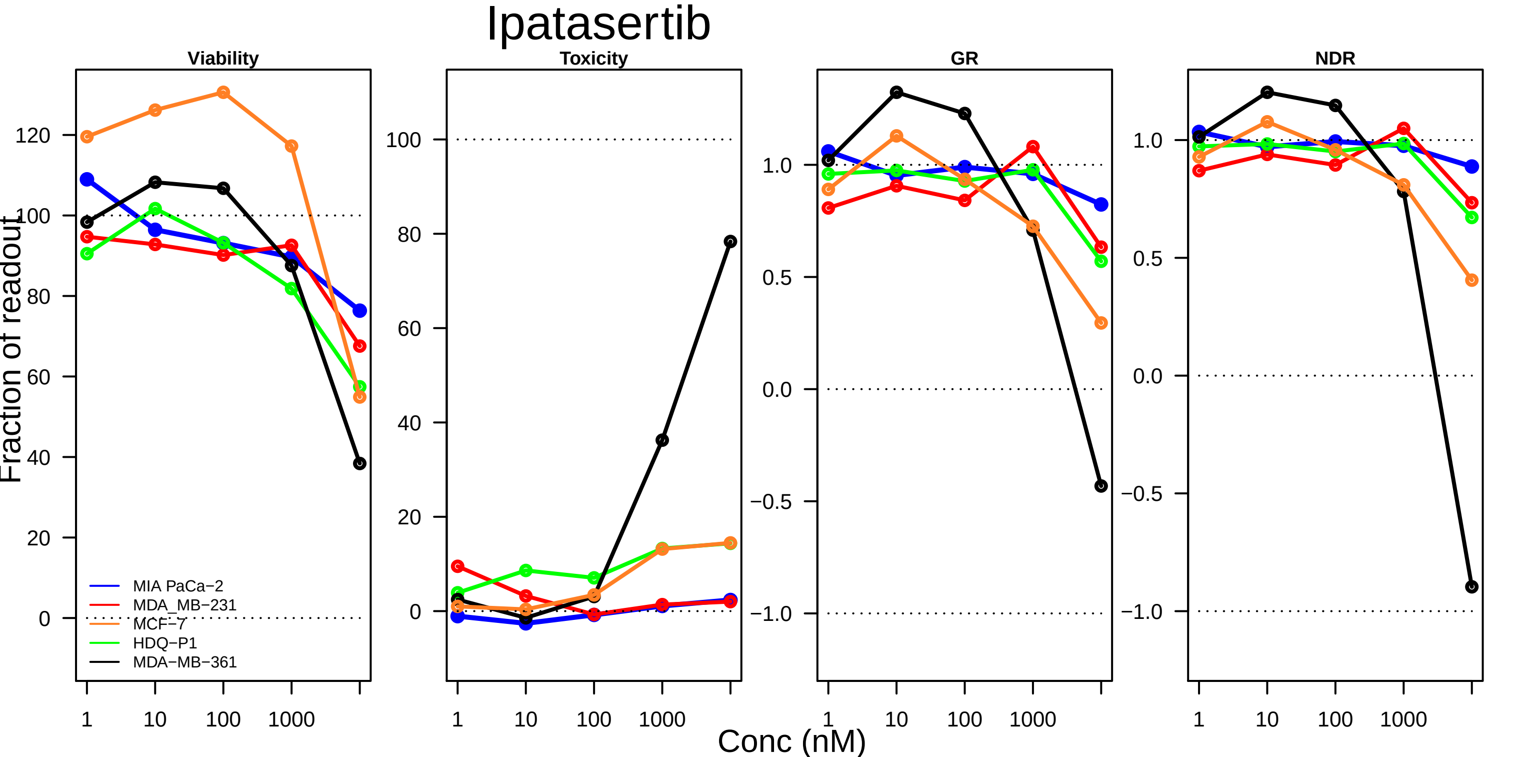

Ipatasertib
Viability
Toxicity
GR
NDR
Fraction of readout
Conc (nM)
1
10
100
1000
MIA PaCa−2
MDA_MB−231
MCF−7
HDQ−P1
MDA−MB−361

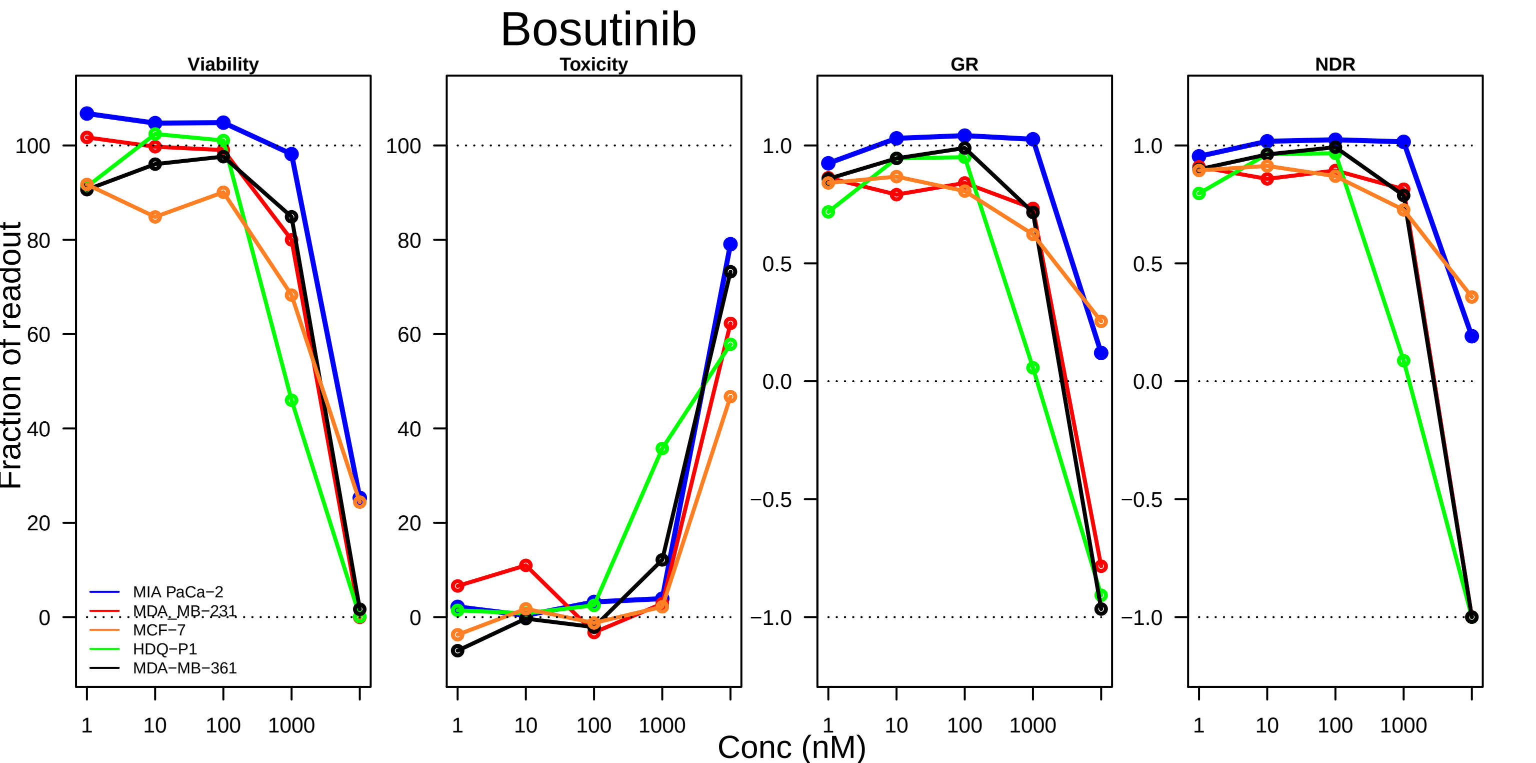

Bosutinib
Viability
Toxicity
GR
NDR
Fraction of readout
Conc (nM)
MIA PaCa−2
MDA_MB−231
MCF−7
HDQ−P1
MDA−MB−361
100
80
60
40
20
0
1.0
0.5
0.0
−0.5
−1.0
1
10
100
1000

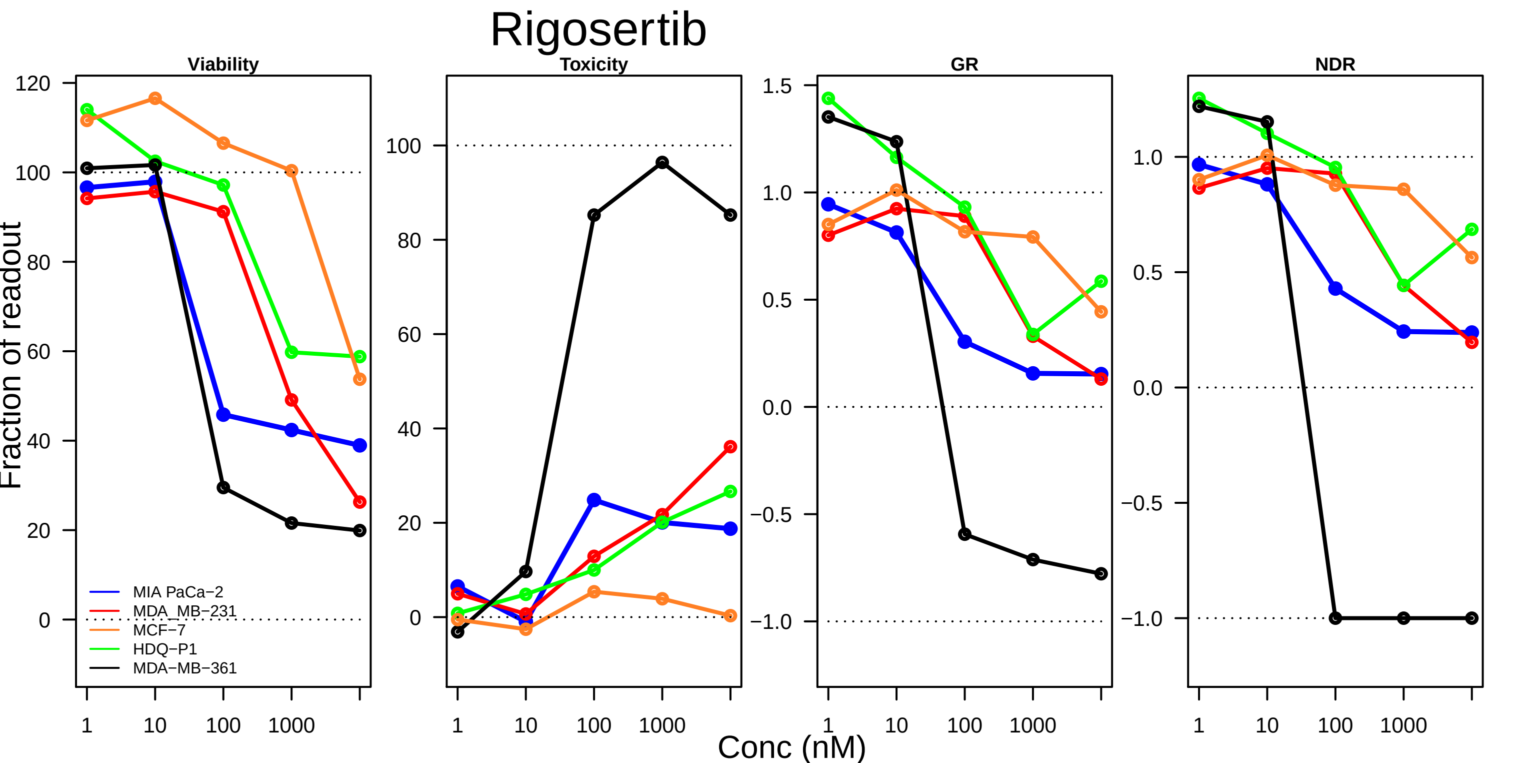

Rigosertib
Viability
Toxicity
GR
NDR
Fraction of readout
Conc (nM)
MIA PaCa−2
MDA_MB−231
MCF−7
HDQ−P1
MDA−MB−361

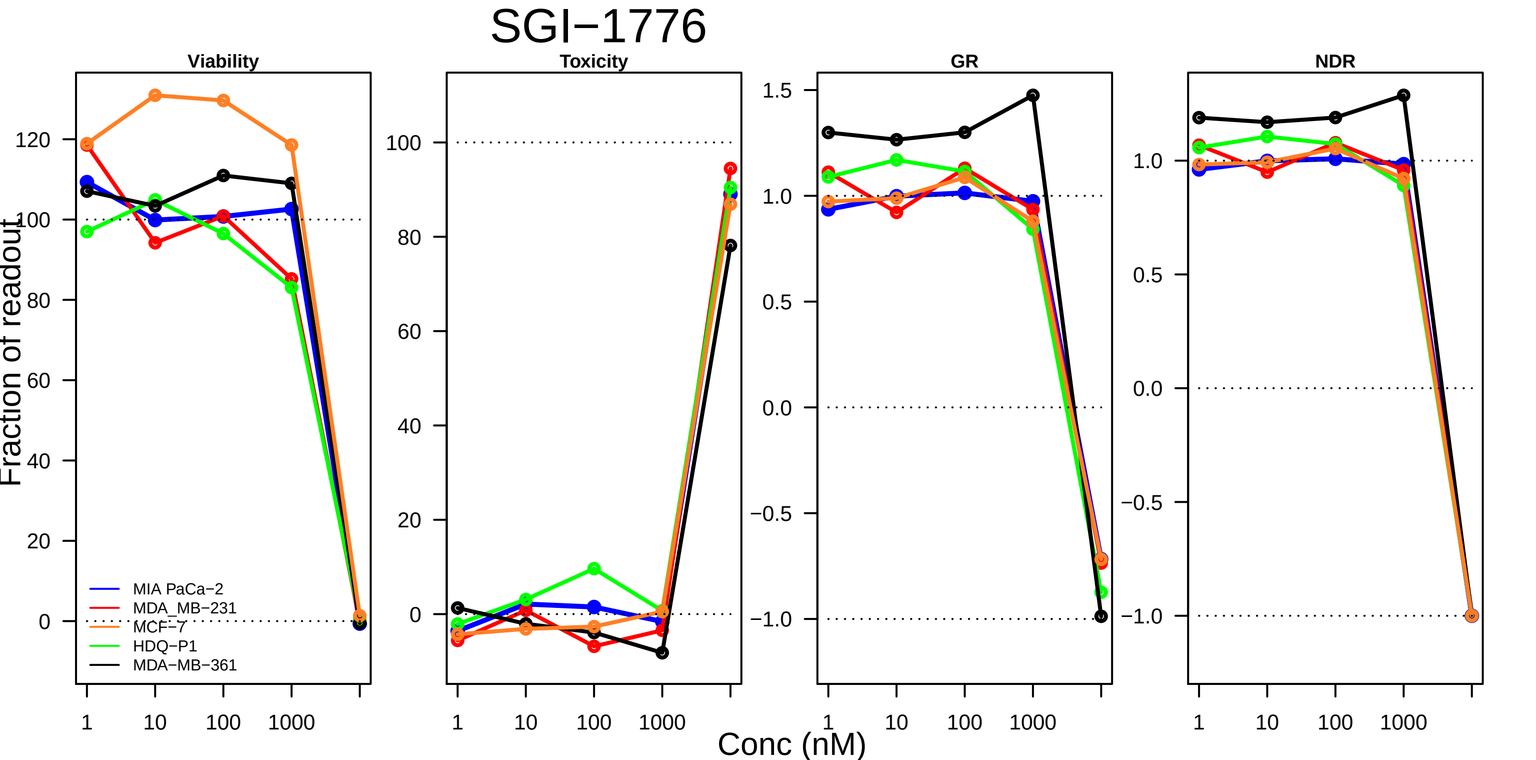
SGI−1776
Viability
Toxicity
GR
NDR
Fraction of readout
Conc (nM)
1
10
100
1000
120
100
80
60
40
20
0
1.5
1.0
0.5
0.0
−0.5
−1.0
MIA PaCa−2
MDA_MB−231
MCF−7
HDQ−P1
MDA−MB−361

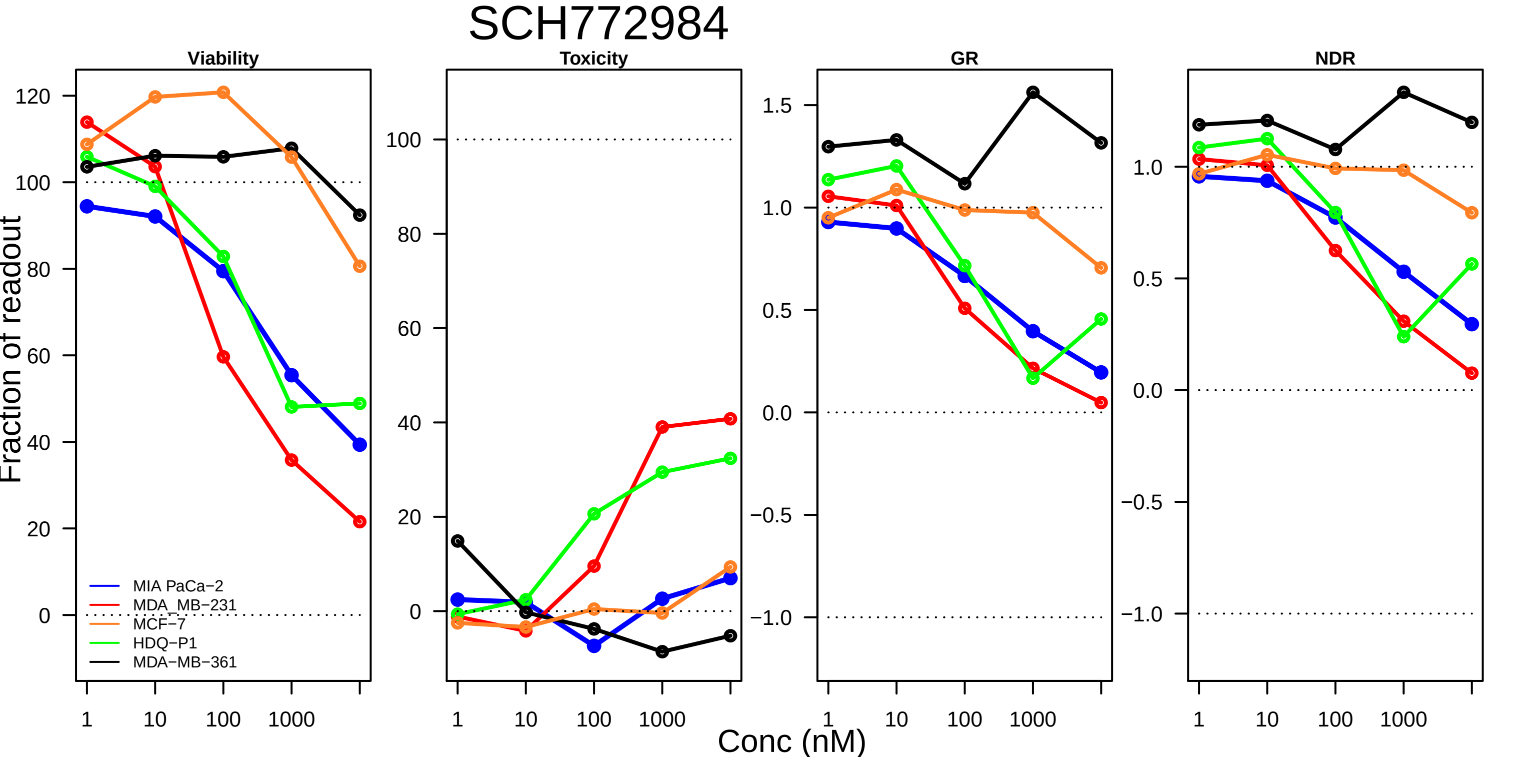

SCH772984
Viability
Toxicity
GR
NDR
Fraction of readout
Conc (nM)
MIA PaCa−2
MDA_MB−231
MCF−7
HDQ−P1
MDA−MB−361
120
100
80
60
40
20
0
1
10
100
1000
1.5
1.0
0.5
0.0
−0.5
−1.0

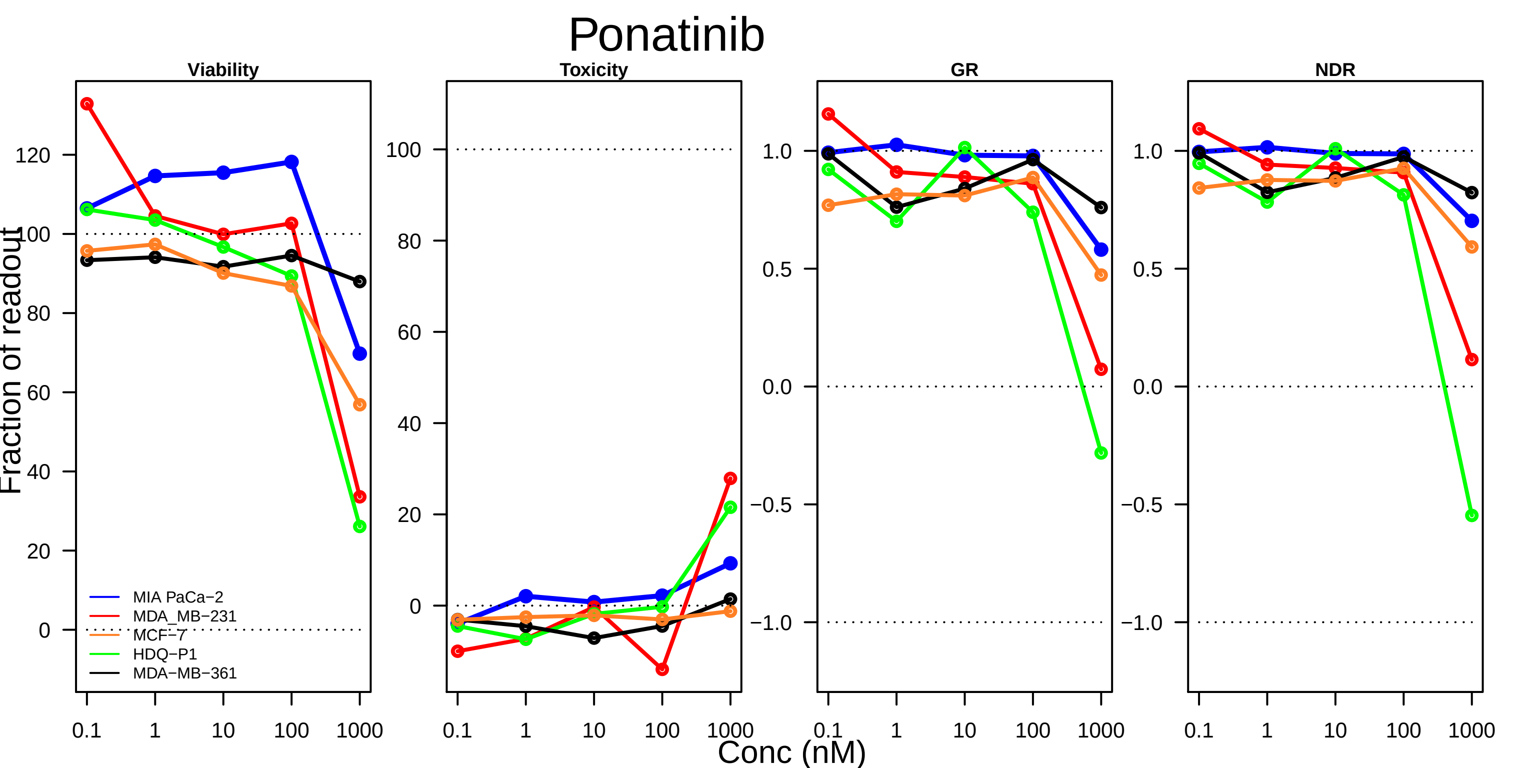

Ponatinib
Viability
Toxicity
GR
NDR
Fraction of readout
Conc (nM)
0.1
1
10
100
1000
0
20
40
60
80
100
120
-0.5
-1.0
0.0
0.5
1.0
MIA PaCa−2
MDA_MB−231
MCF−7
HDQ−P1
MDA−MB−361

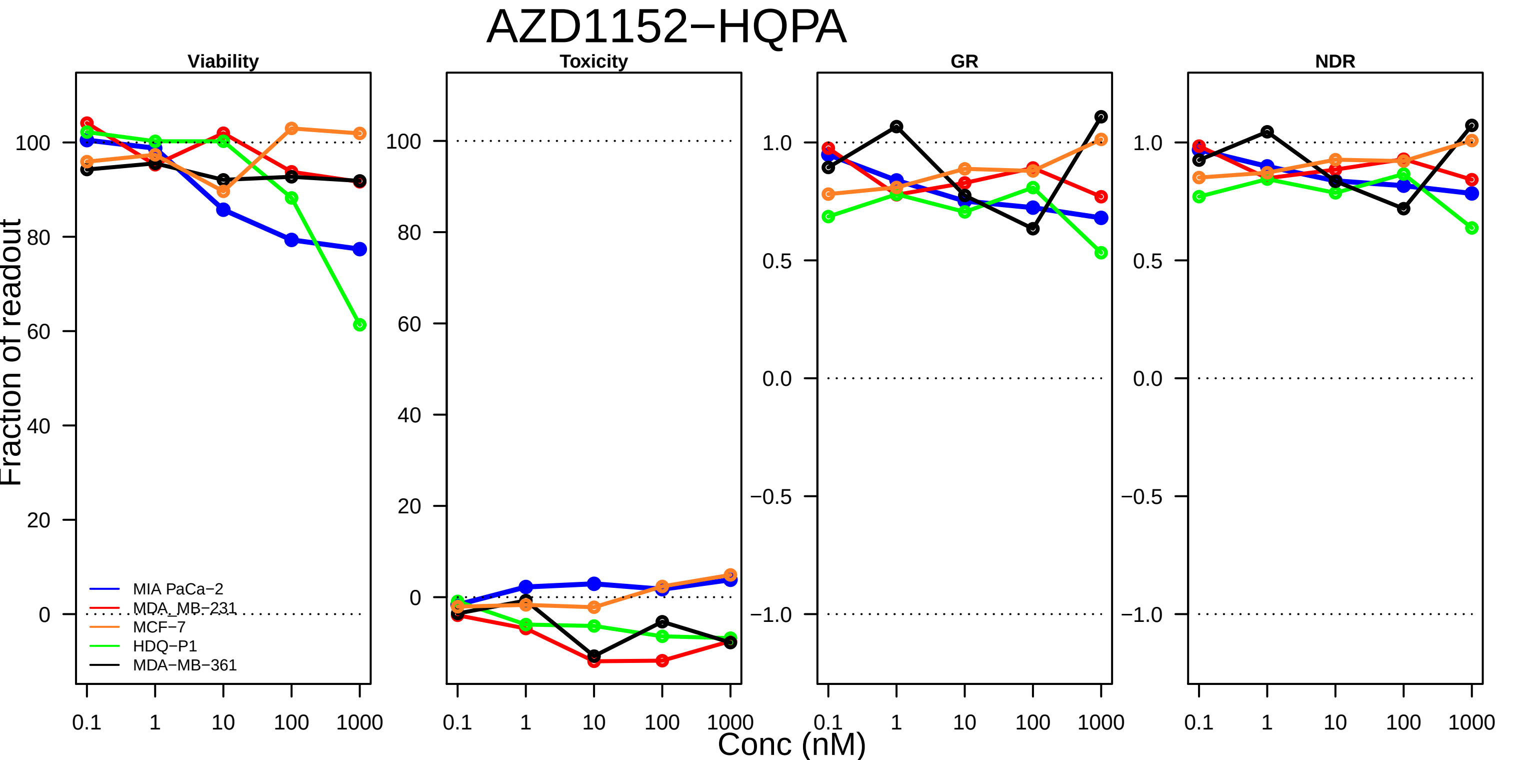
AZD1152−HQPA
Viability
Toxicity
GR
NDR
Fraction of readout
Conc (nM)
MIA PaCa−2
MDA_MB−231
MCF−7
HDQ−P1
MDA−MB−361
0.1
1
10
100
1000
0
20
40
60
80
100
−1.0
−0.5
0.0
0.5
1.0

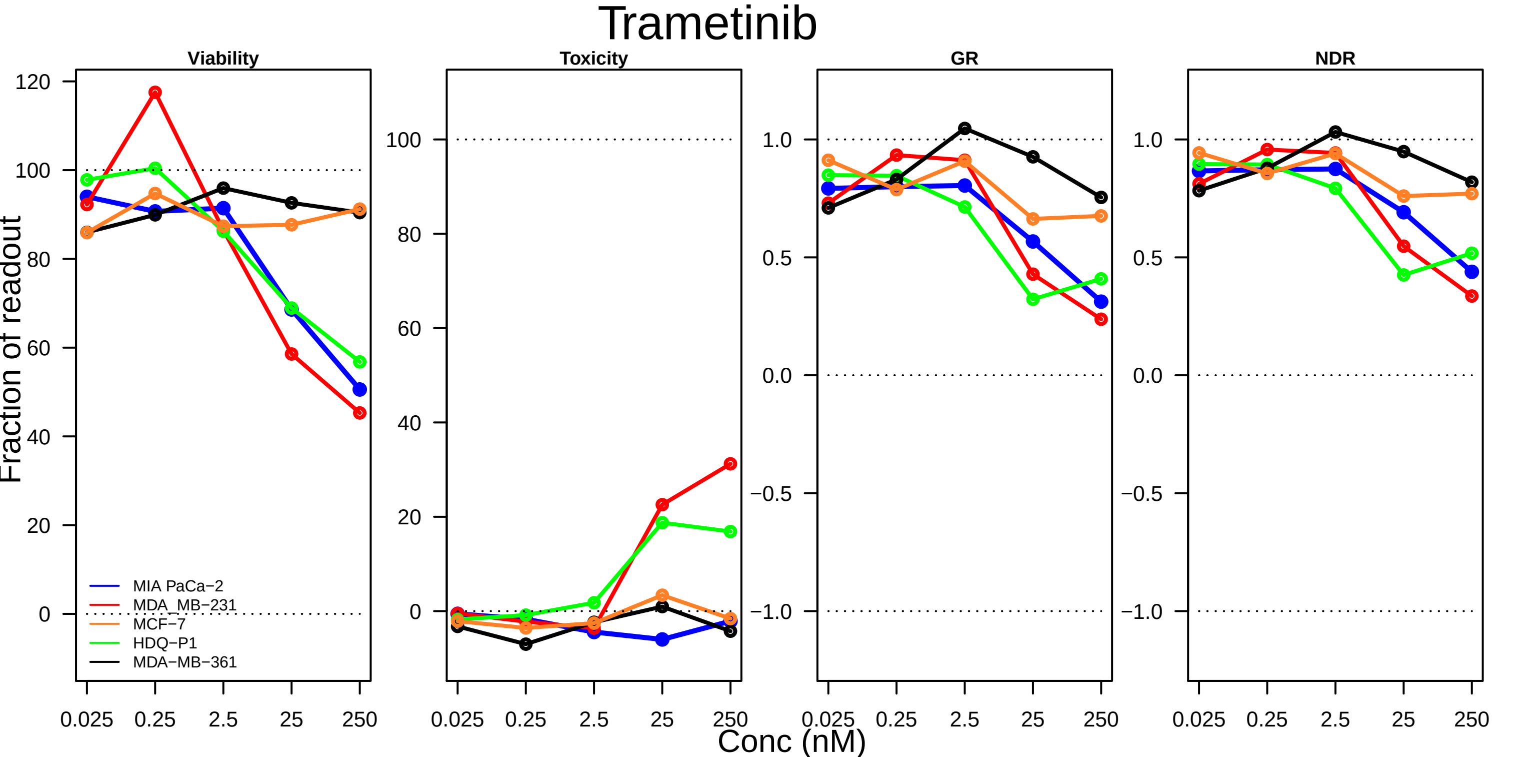
Trametinib
Viability
Toxicity
GR
NDR
Fraction of readout
Conc (nM)
MIA PaCa−2
MDA_MB−231
MCF−7
HDQ−P1
MDA−MB−361
0.025
0.25
2.5
25
250

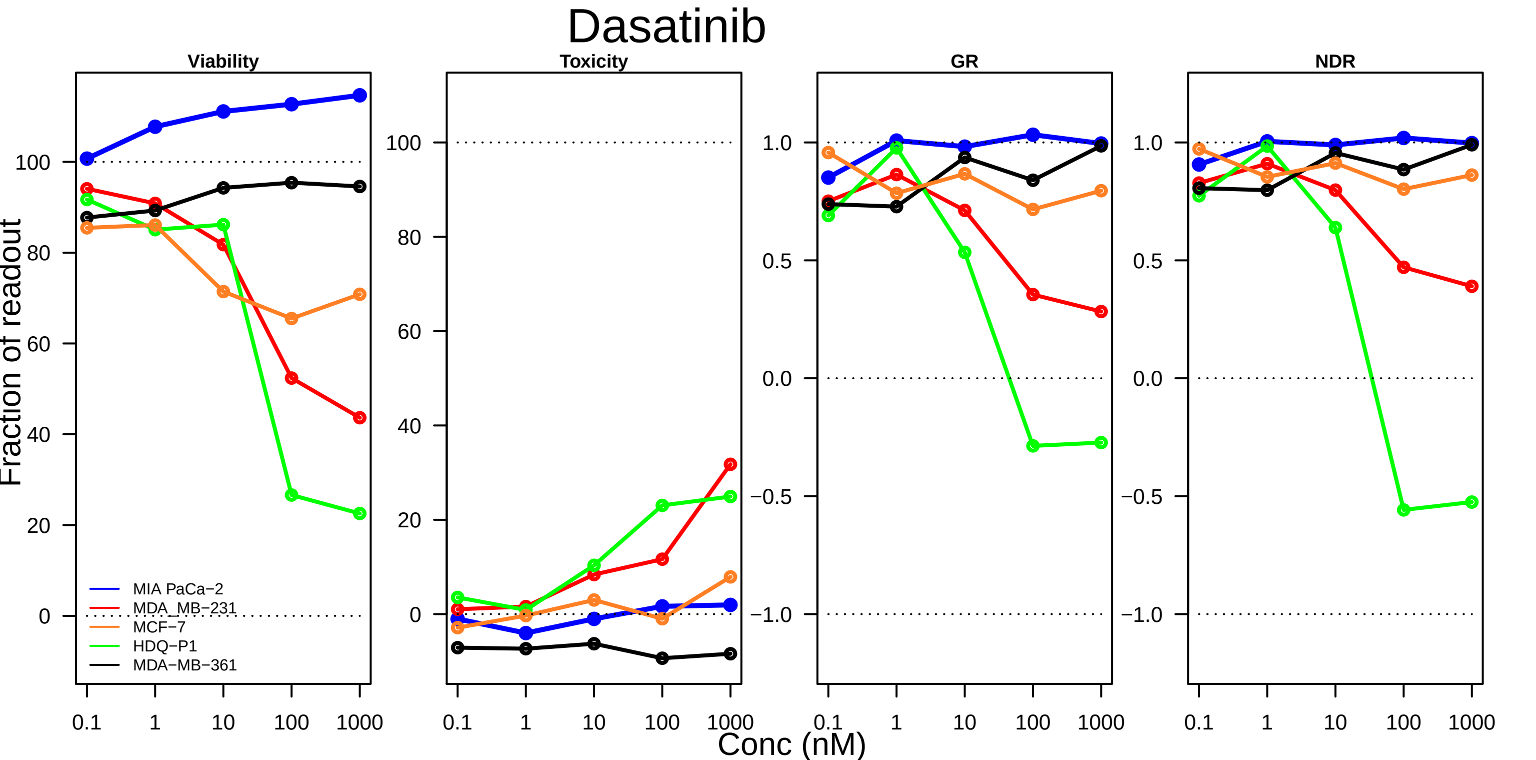

Dasatinib
Viability
Toxicity
GR
NDR
Fraction of readout
Conc (nM)
MIA PaCa−2
MDA_MB−231
MCF−7
HDQ−P1
MDA−MB−361
0.1
1
10
100
1000
0
20
40
60
80
100
−1.0
−0.5
0.0
0.5
1.0

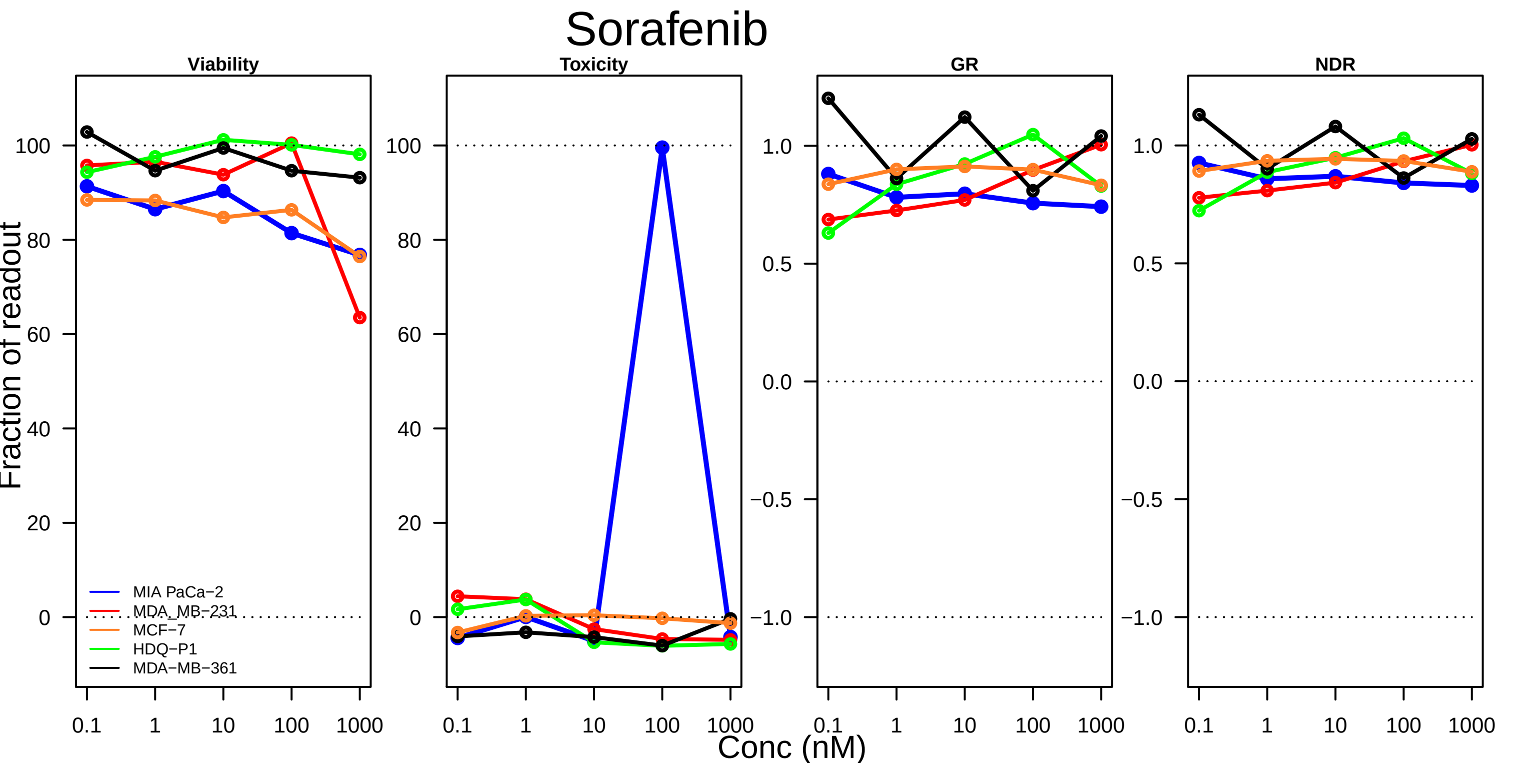
Sorafenib
Viability
Toxicity
GR
NDR
Fraction of readout
Conc (nM)
MIA PaCa−2
MDA_MB−231
MCF−7
HDQ−P1
MDA−MB−361
100
80
60
40
20
0
1.0
0.5
0.0
−0.5
−1.0
0.1
1
10
100
1000

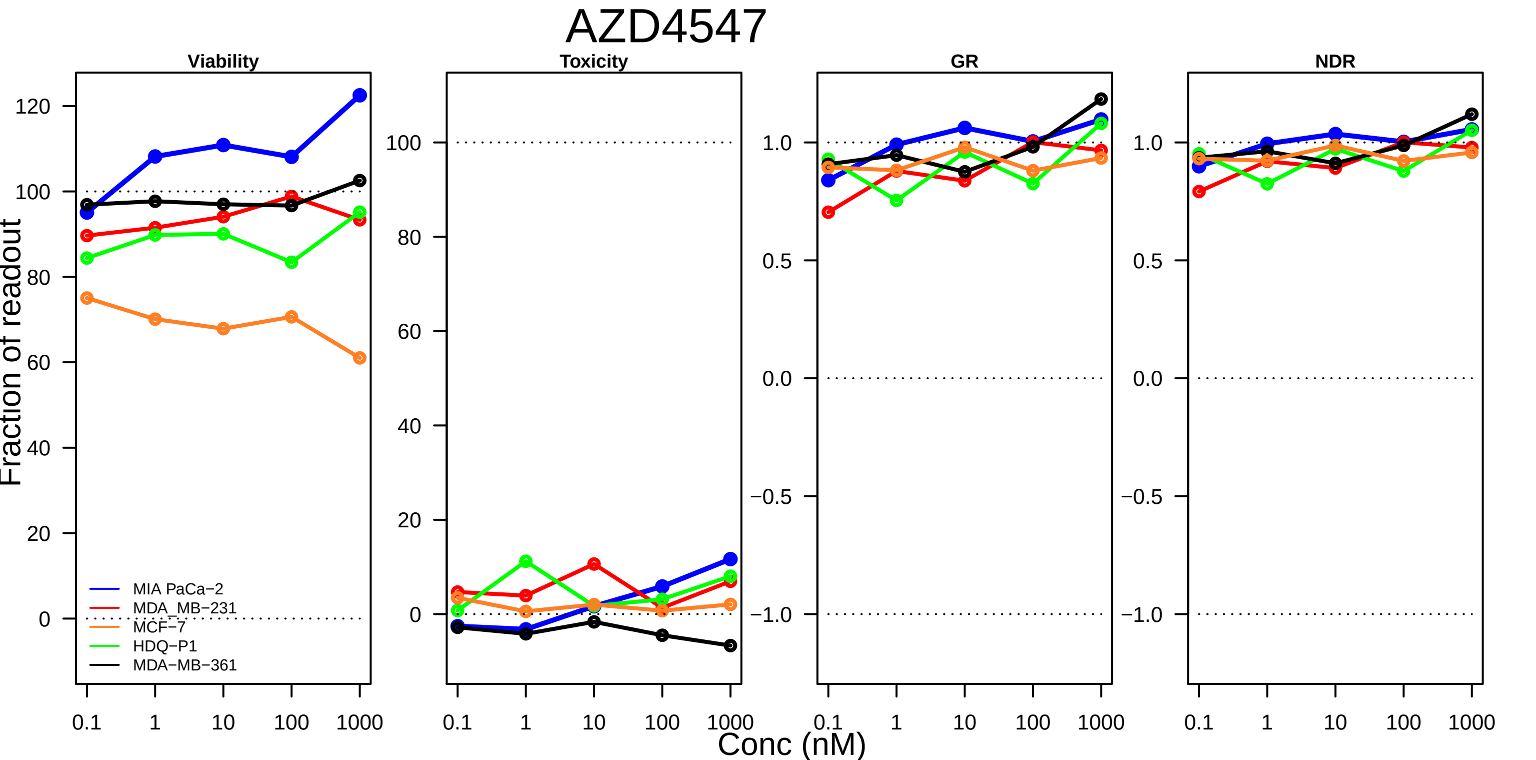

AZD4547
Viability
Toxicity
GR
NDR
Fraction of readout
MIA PaCa−2
MDA_MB−231
MCF−7
HDQ−P1
MDA−MB−361
Conc (nM)

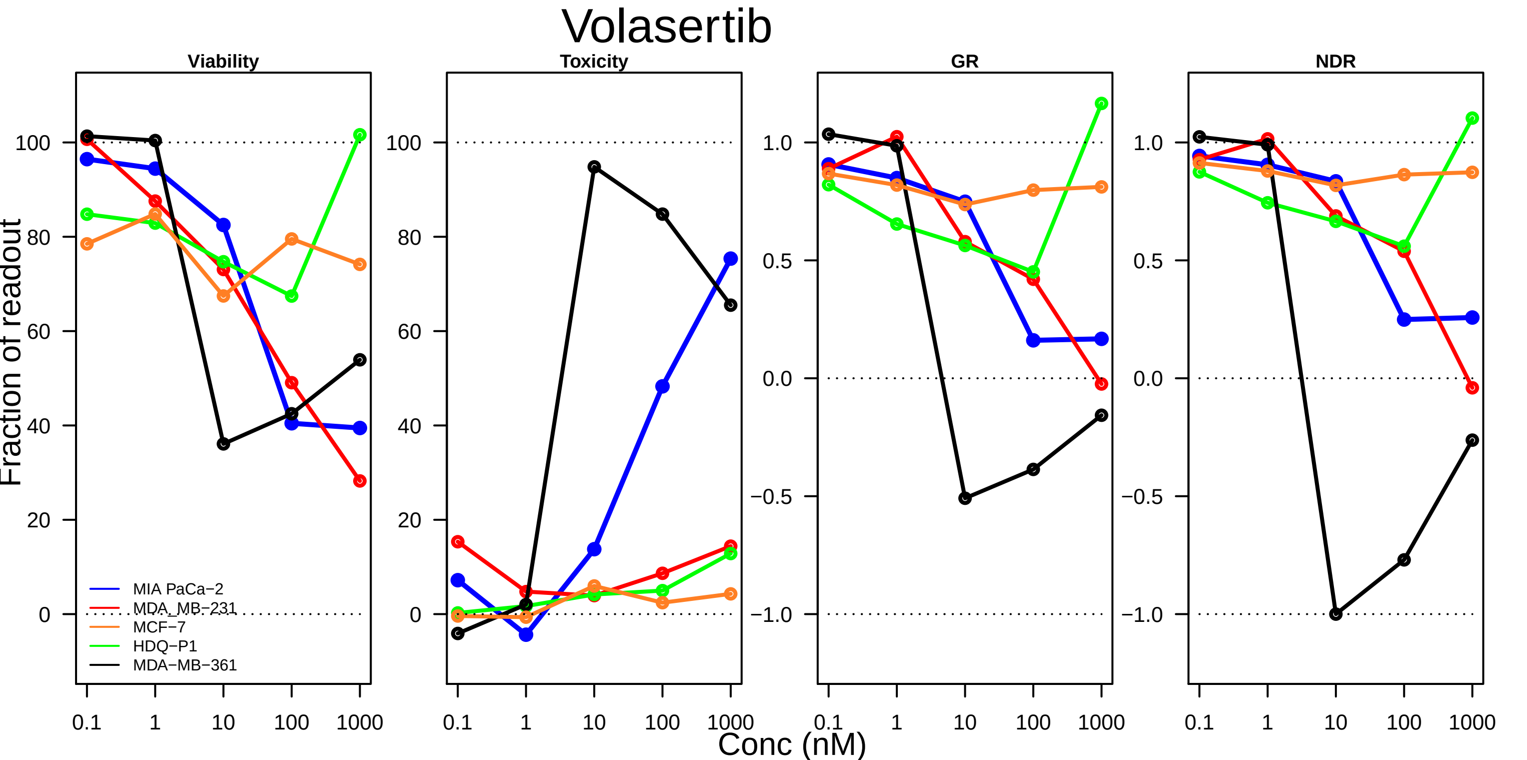

Volasertib
Viability
Toxicity
GR
NDR
Fraction of readout
Conc (nM)
100
80
60
40
20
0
1.0
0.5
0.0
−0.5
−1.0
0.1
1
10
100
1000
MIA PaCa−2
MDA_MB−231
MCF−7
HDQ−P1
MDA−MB−361

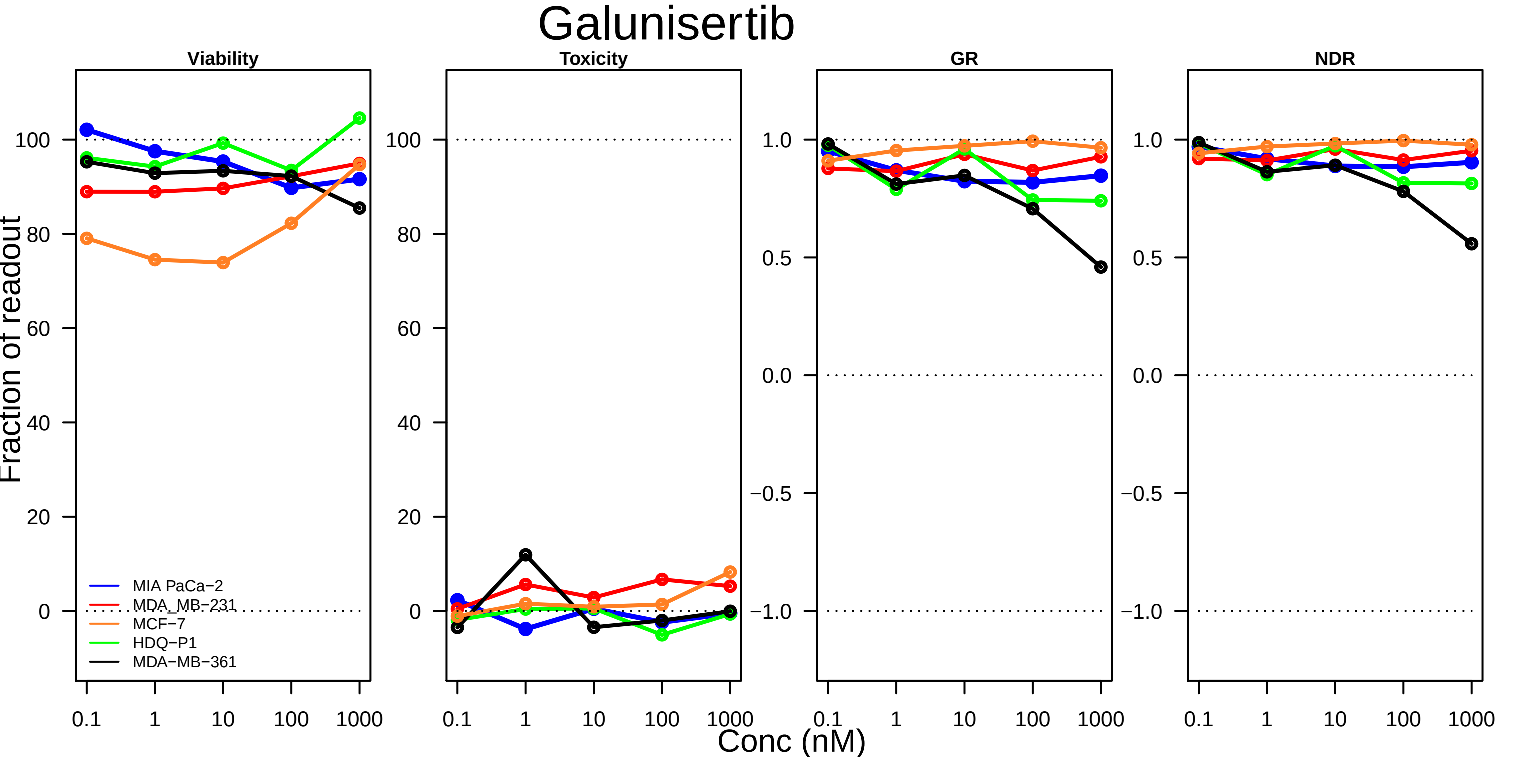
Galunisertib
Viability
Toxicity
GR
NDR
Fraction of readout
Conc (nM)
MIA PaCa−2
MDA_MB−231
MCF−7
HDQ−P1
MDA−MB−361
0.1
1
10
100
1000
100
80
60
40
20
0
1.0
0.5
0.0
−0.5
−1.0

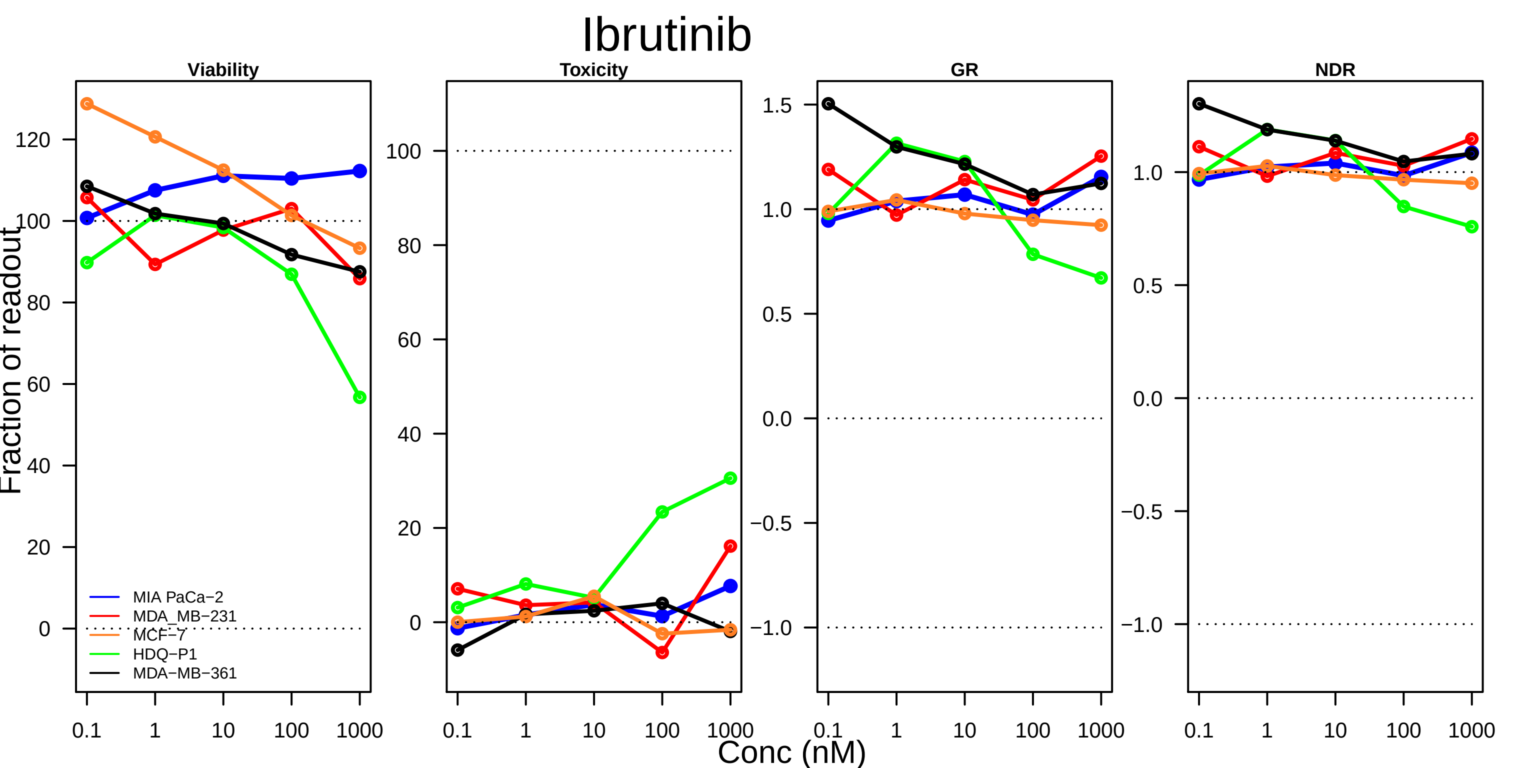

Ibrutinib
Viability
Toxicity
GR
NDR
Fraction of readout
Conc (nM)
0.1
1
10
100
1000
MIA PaCa−2
MDA_MB−231
MCF−7
HDQ−P1
MDA−MB−361

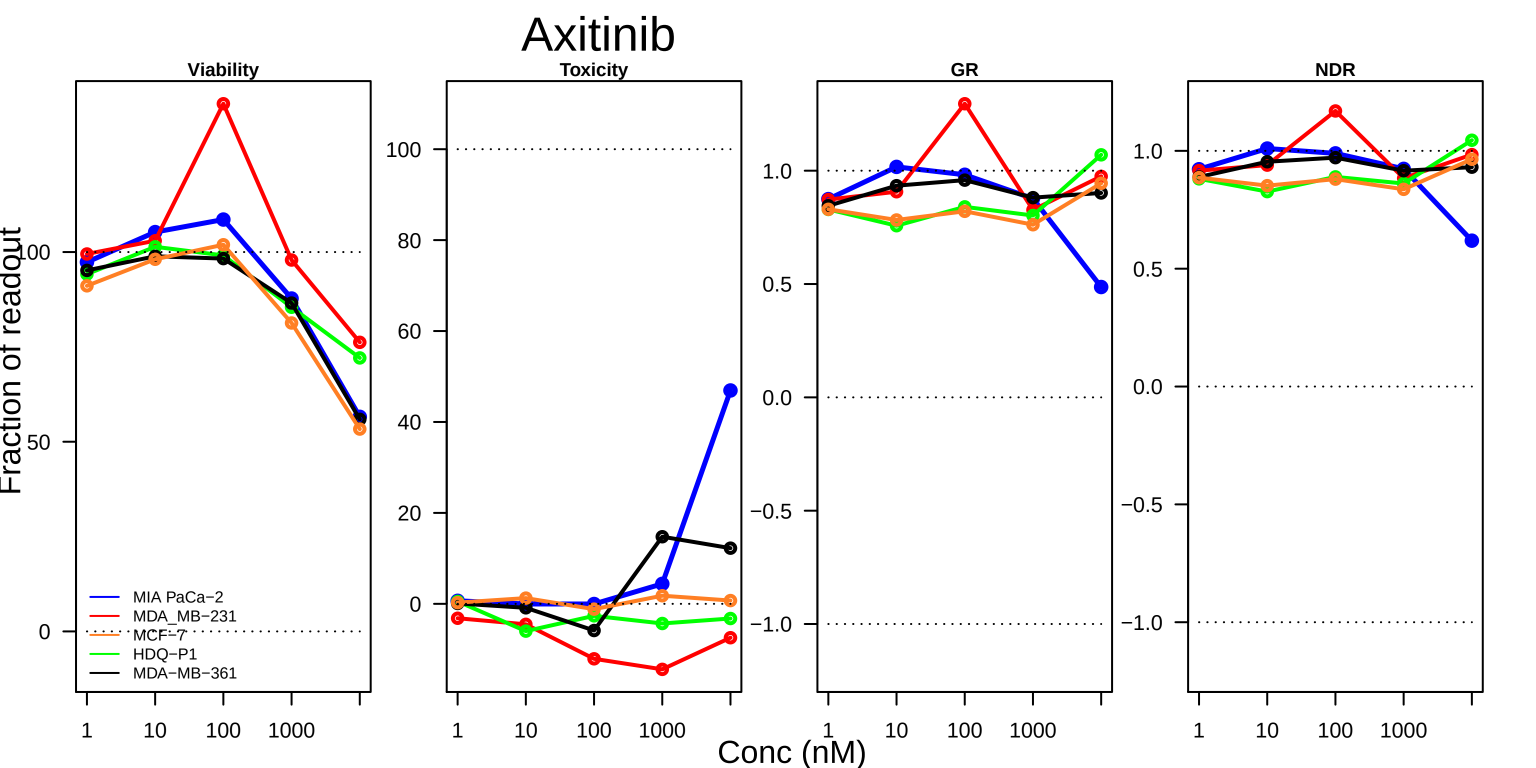

Axitinib
Viability
Toxicity
GR
NDR
Fraction of readout
Conc (nM)
MIA PaCa−2
MDA_MB−231
MCF−7
HDQ−P1
MDA−MB−361
0
50
100
20
40
60
80
−1.0
−0.5
0.0
0.5
1.0
1
10
100
1000

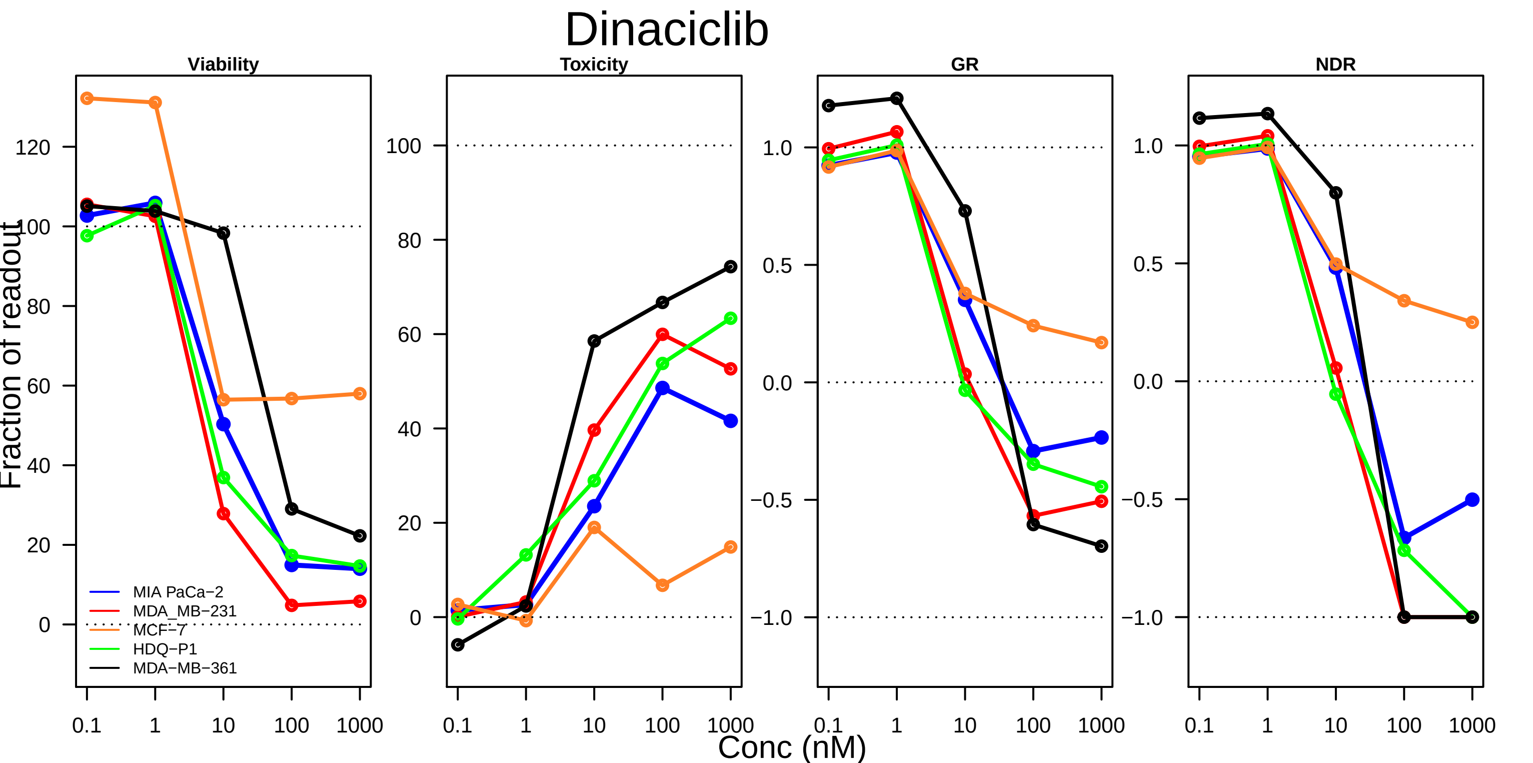
Dinaciclib
Viability
Toxicity
GR
NDR
Fraction of readout
Conc (nM)
MIA PaCa−2
MDA_MB−231
MCF−7
HDQ−P1
MDA−MB−361
0.1
1
10
100
1000

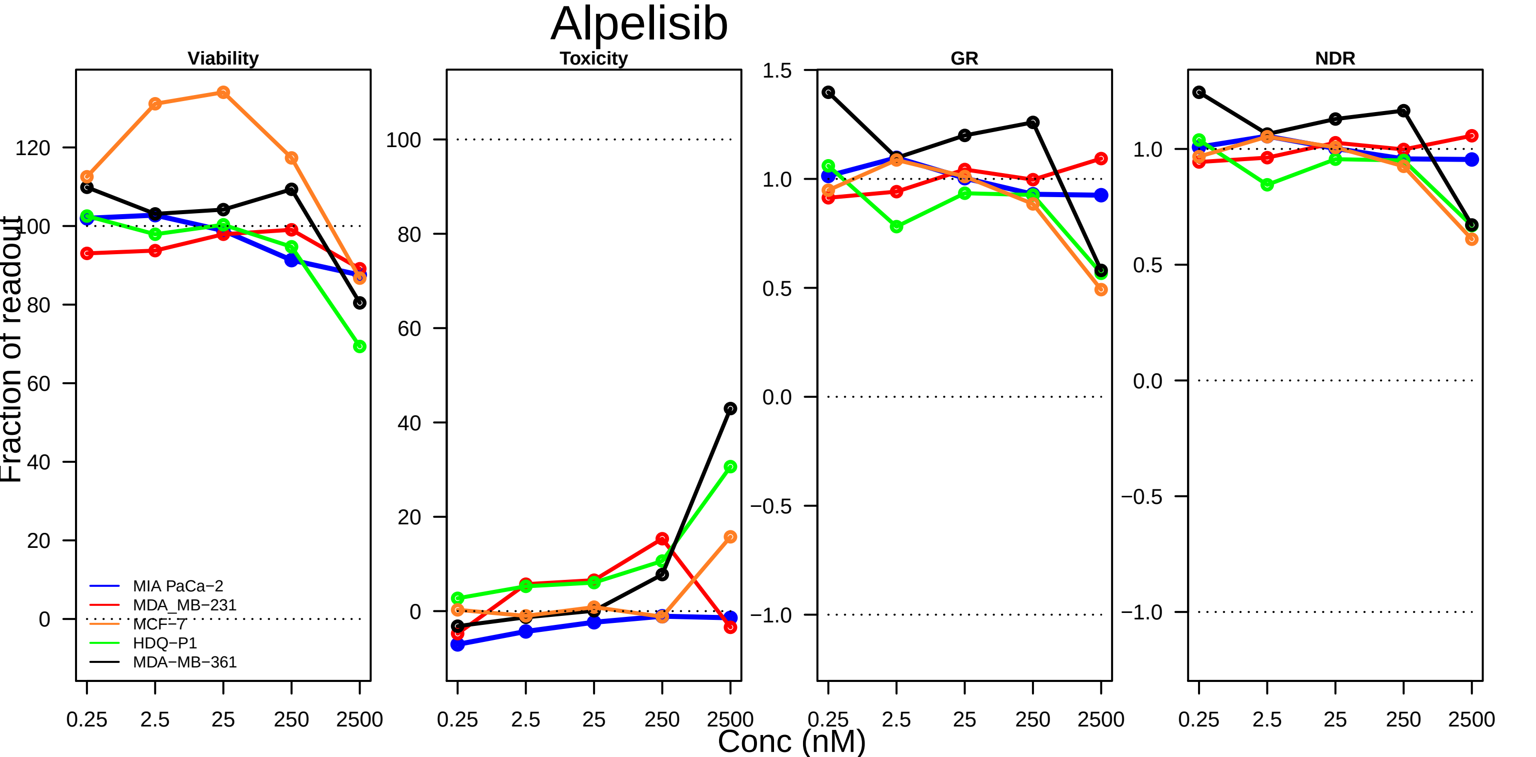

Alpelisib
Viability
Toxicity
GR
NDR
Fraction of readout
Conc (nM)
0.25
2.5
25
250
2500
MIA PaCa−2
MDA_MB−231
MCF−7
HDQ−P1
MDA−MB−361

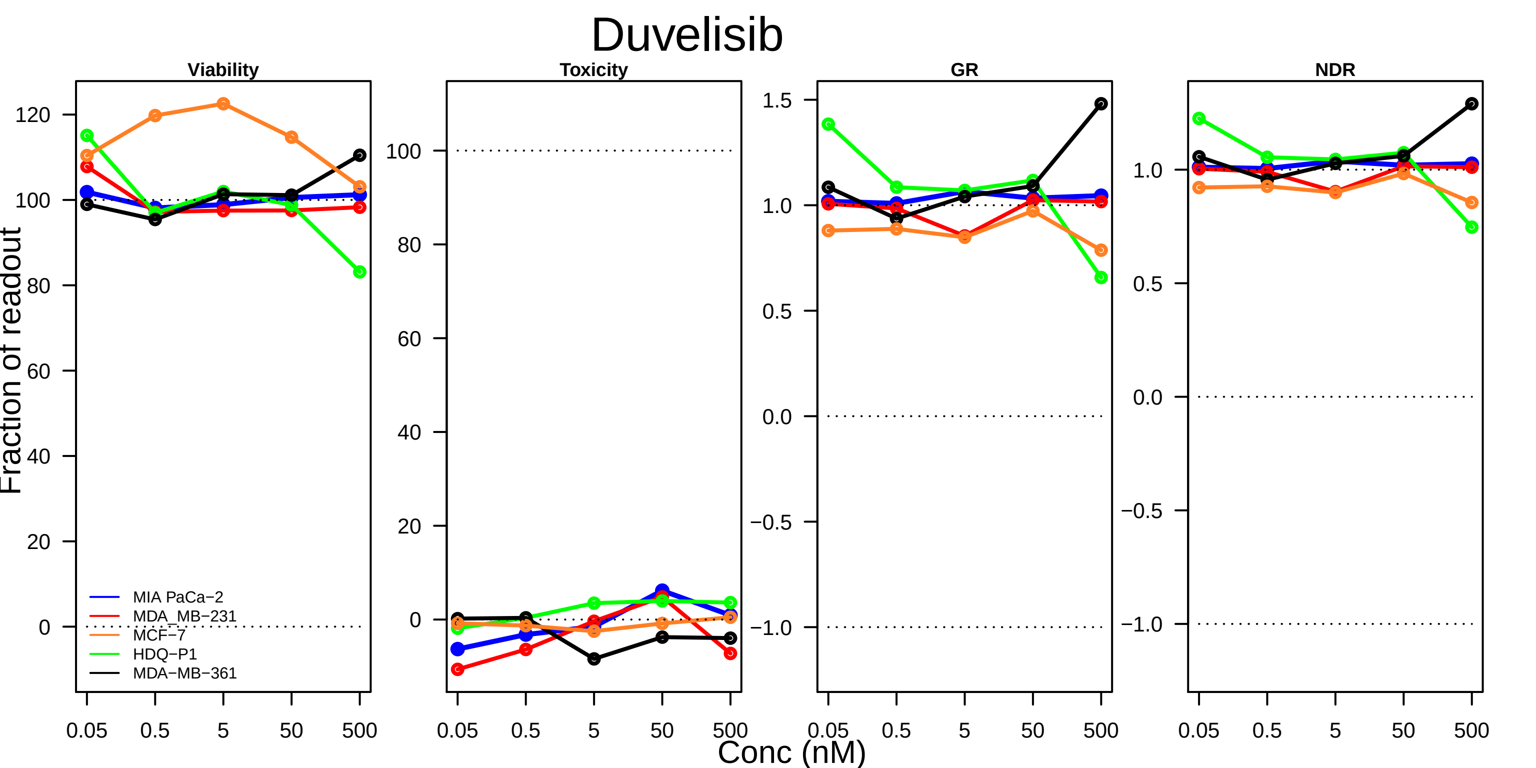
Duvelisib
Viability
Toxicity
GR
NDR
Fraction of readout
120
100
80
60
40
20
0
1.5
1.0
0.5
0.0
−0.5
−1.0
MIA PaCa−2
MDA_MB−231
MCF−7
HDQ−P1
MDA−MB−361
0.05
0.5
5
50
500
Conc (nM)

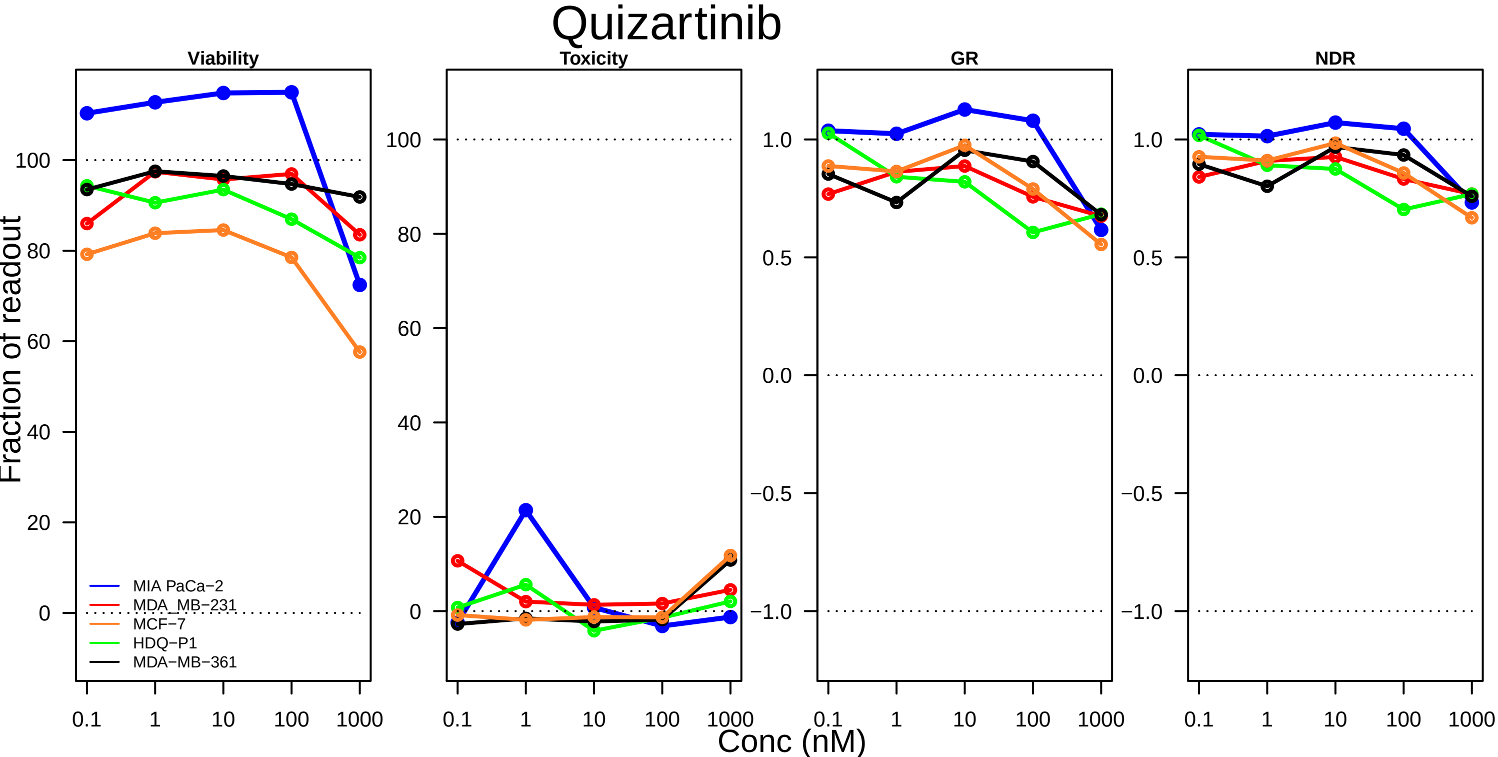

Quizartinib
Viability
Toxicity
GR
NDR
Fraction of readout
Conc (nM)
MIA PaCa−2
MDA_MB−231
MCF−7
HDQ−P1
MDA−MB−361
0.1
1
10
100
1000

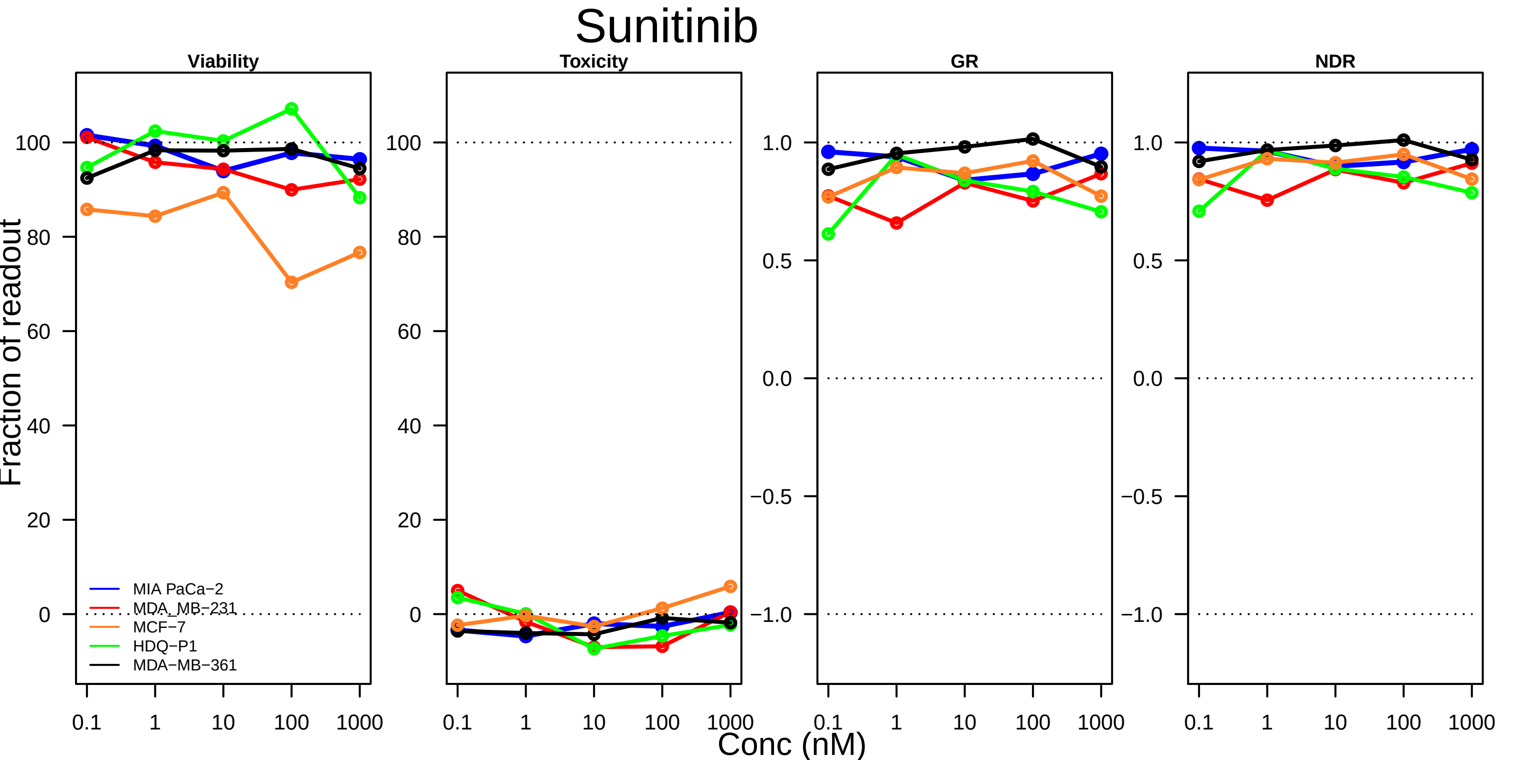
Sunitinib
Viability
Toxicity
GR
NDR
Fraction of readout
Conc (nM)
MIA PaCa−2
MDA_MB−231
MCF−7
HDQ−P1
MDA−MB−361
0
20
40
60
80
100
−1.0
−0.5
0.0
0.5
1.0
0.1
1
10
100
1000

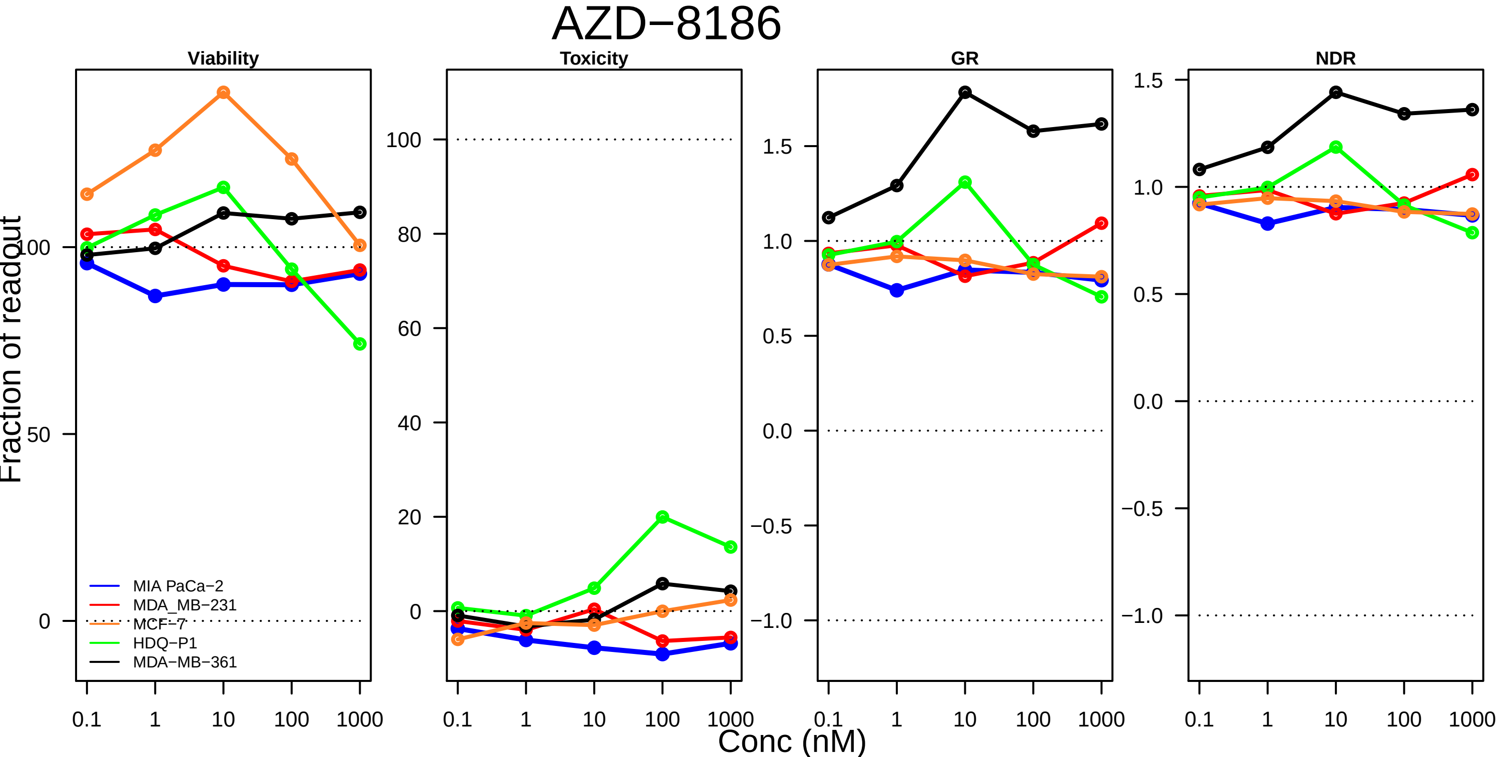

AZD−8186
Viability
Toxicity
GR
NDR
Fraction of readout
Conc (nM)
MIA PaCa−2
MDA_MB−231
MCF−7
HDQ−P1
MDA−MB−361
0.1
1
10
100
1000
0
50
100
20
40
60
80
−1.0
−0.5
0.0
0.5
1.0
1.5

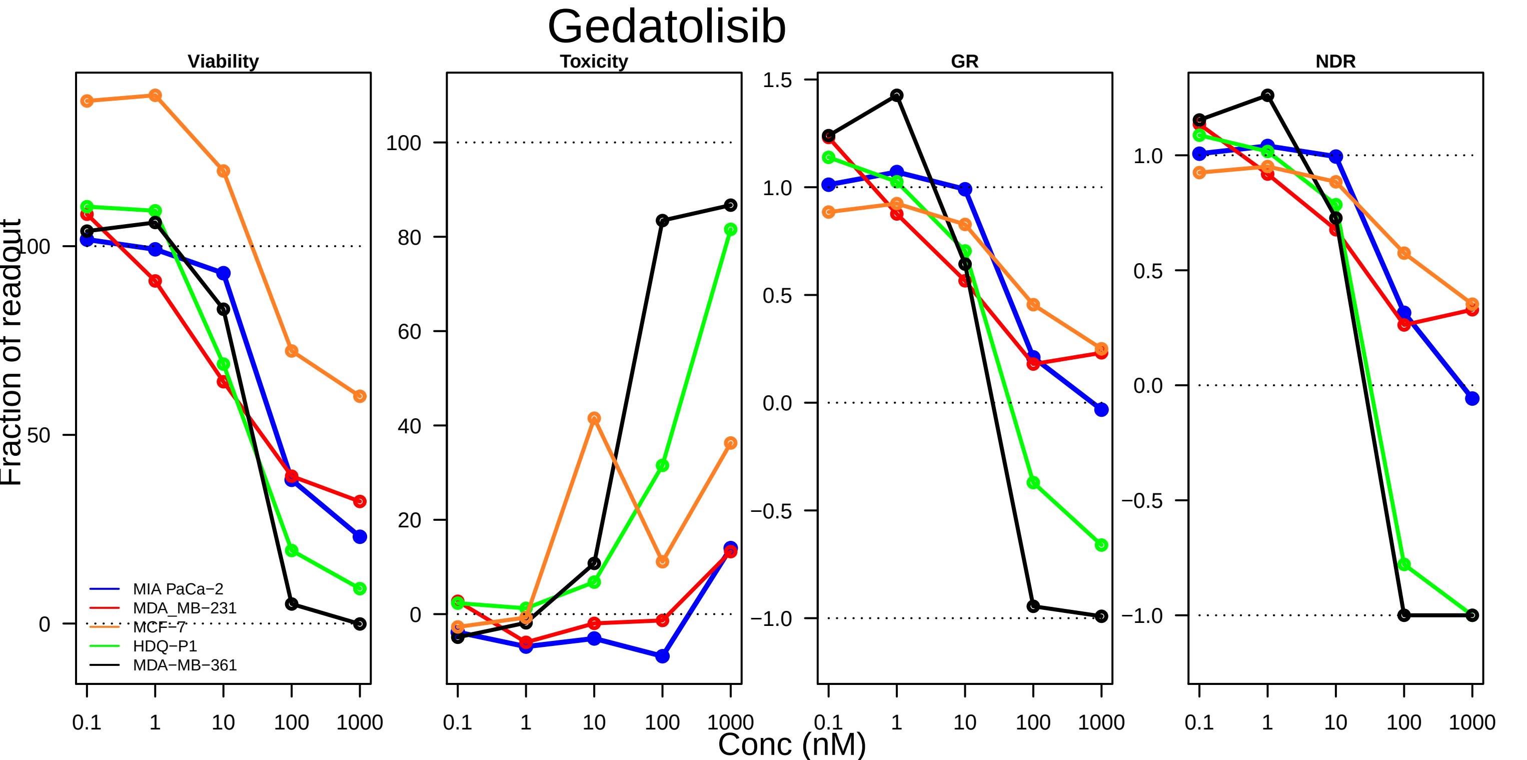

Gedatolisib
Viability
Toxicity
GR
NDR
Fraction of readout
100
50
0
80
60
40
20
1.5
1.0
0.5
0.0
−0.5
−1.0
MIA PaCa−2
MDA_MB−231
MCF−7
HDQ−P1
MDA−MB−361
0.1
1
10
100
1000
Conc (nM)

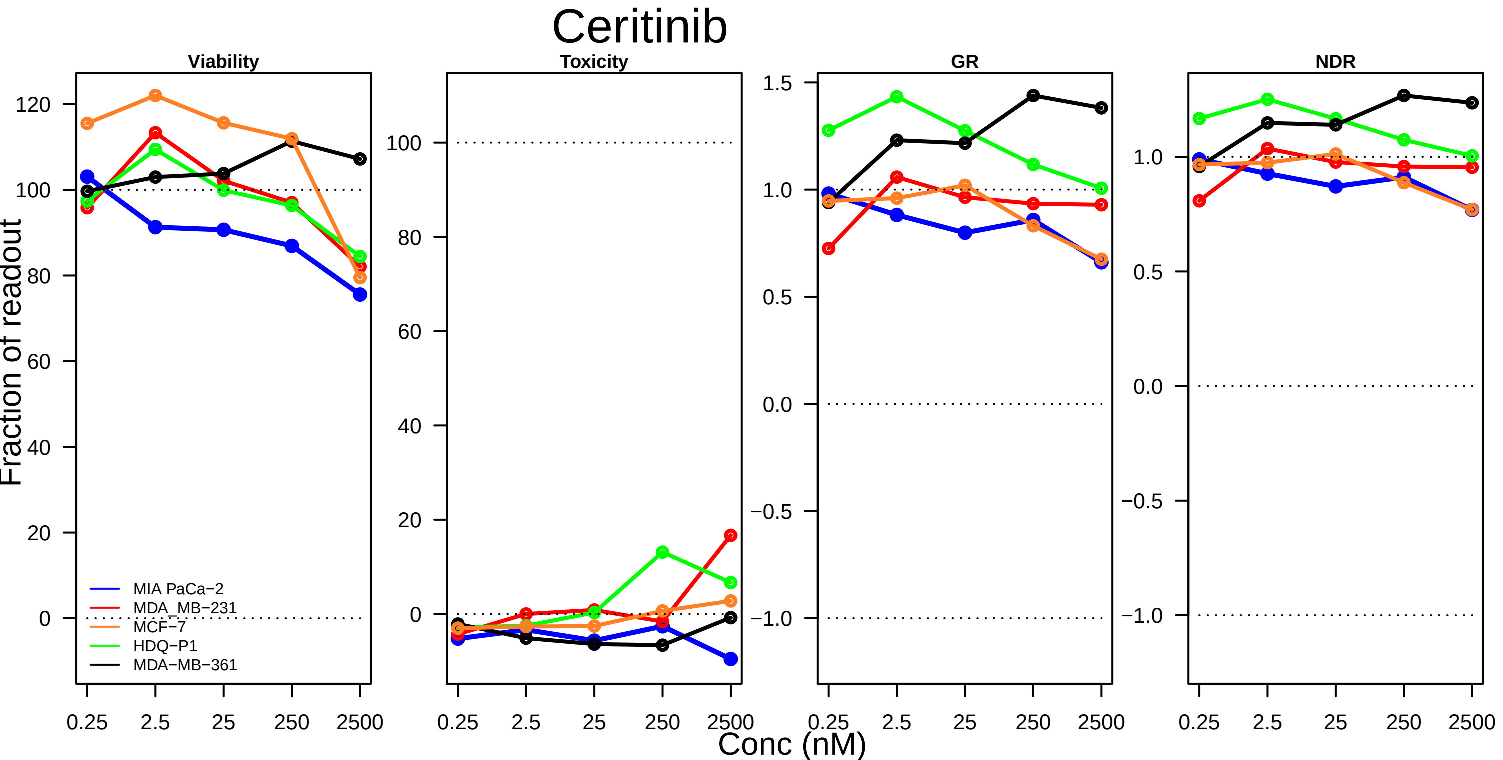

Ceritinib
Viability
Toxicity
GR
NDR
Fraction of readout
Conc (nM)
0.25
2.5
25
250
2500
MIA PaCa−2
MDA_MB−231
MCF−7
HDQ−P1
MDA−MB−361

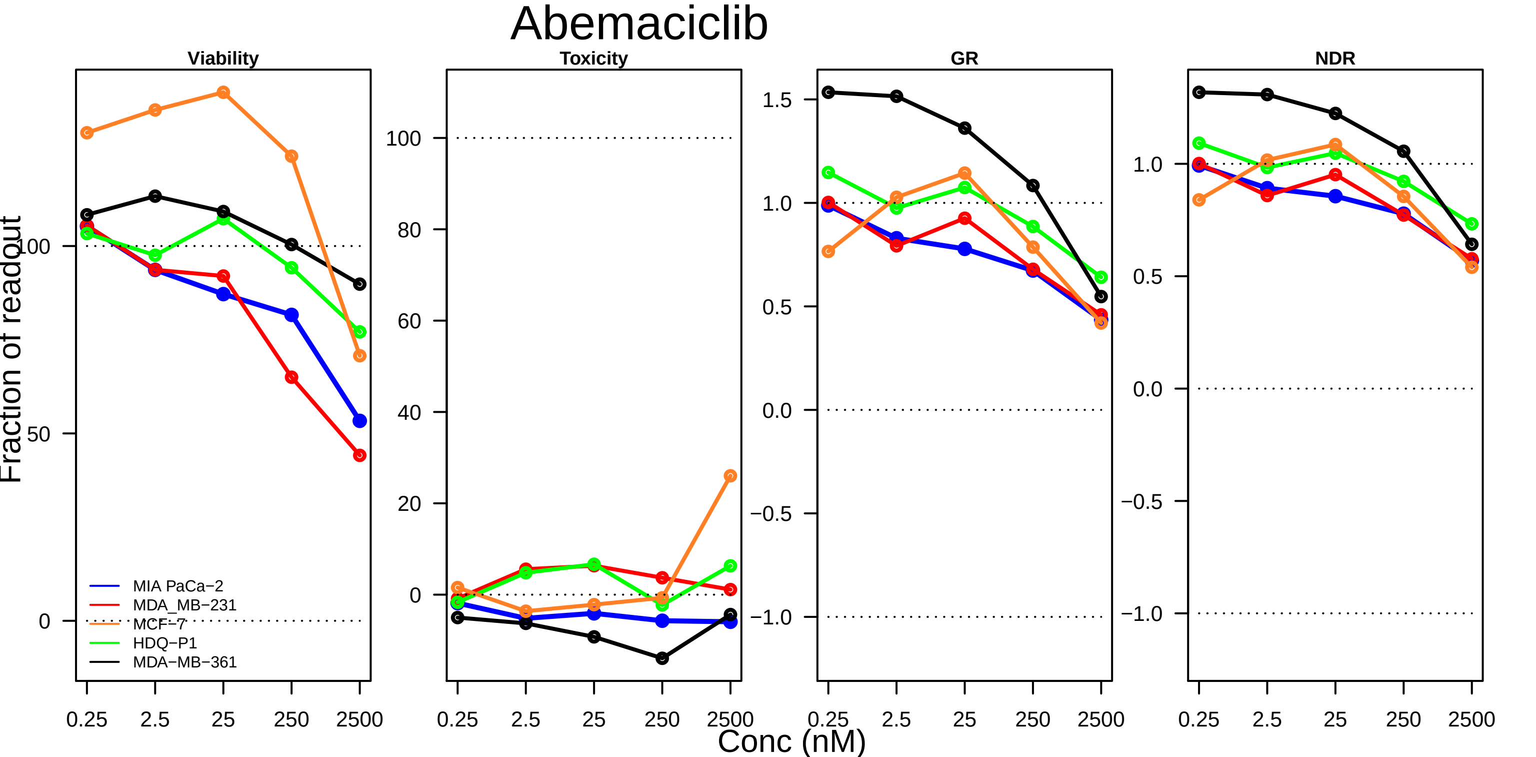

Abemaciclib
Viability
Toxicity
GR
NDR
Fraction of readout
Conc (nM)
0.25
2.5
25
250
2500
0
50
100
20
40
60
80
1.5
1.0
0.5
0.0
−0.5
−1.0
MIA PaCa−2
MDA_MB−231
MCF−7
HDQ−P1
MDA−MB−361

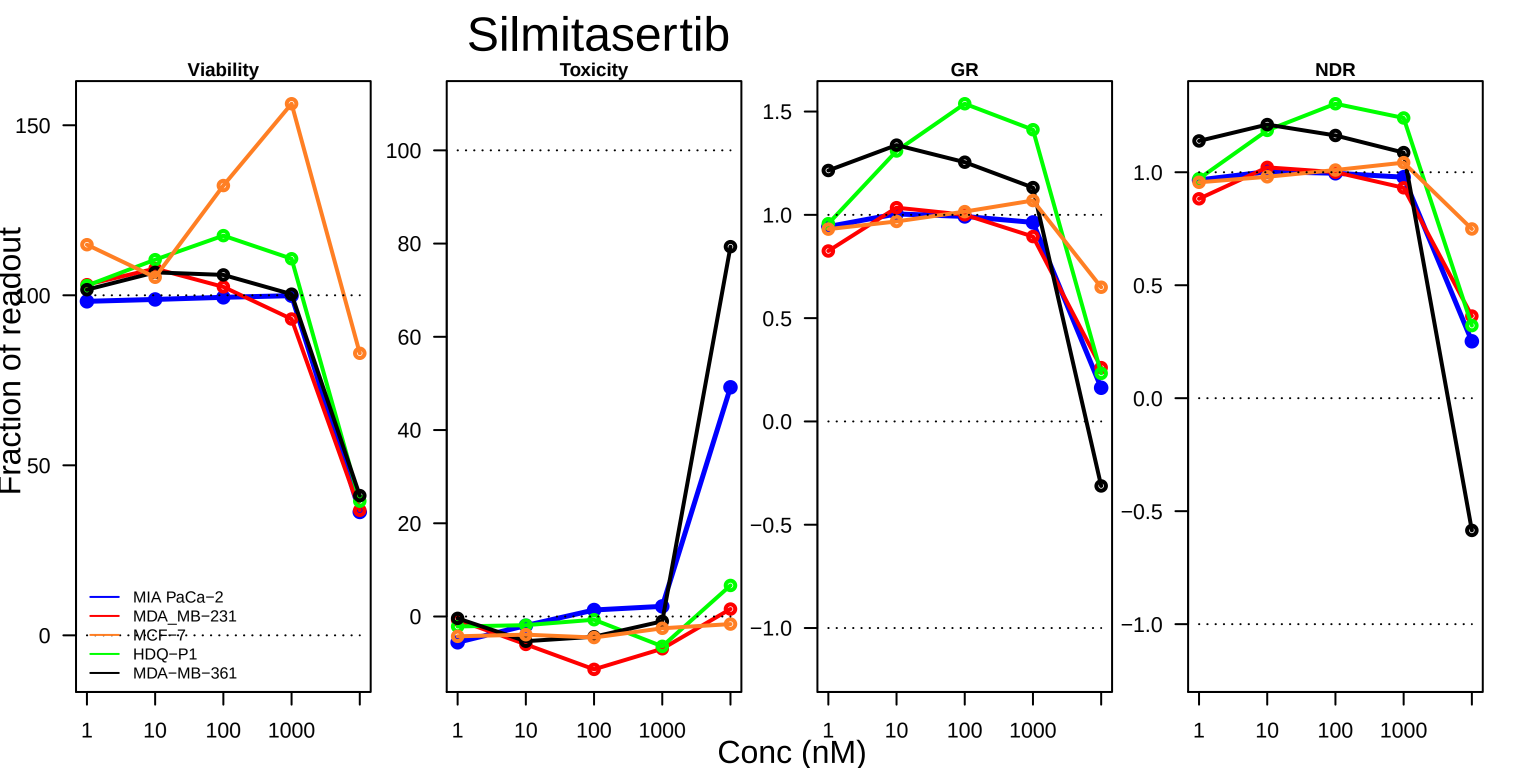

Silmitasertib
Viability
Toxicity
GR
NDR
Fraction of readout
Conc (nM)
1
10
100
1000
150
100
50
0
80
60
40
20
1.5
1.0
0.5
0.0
−0.5
−1.0
MIA PaCa−2
MDA_MB−231
MCF−7
HDQ−P1
MDA−MB−361

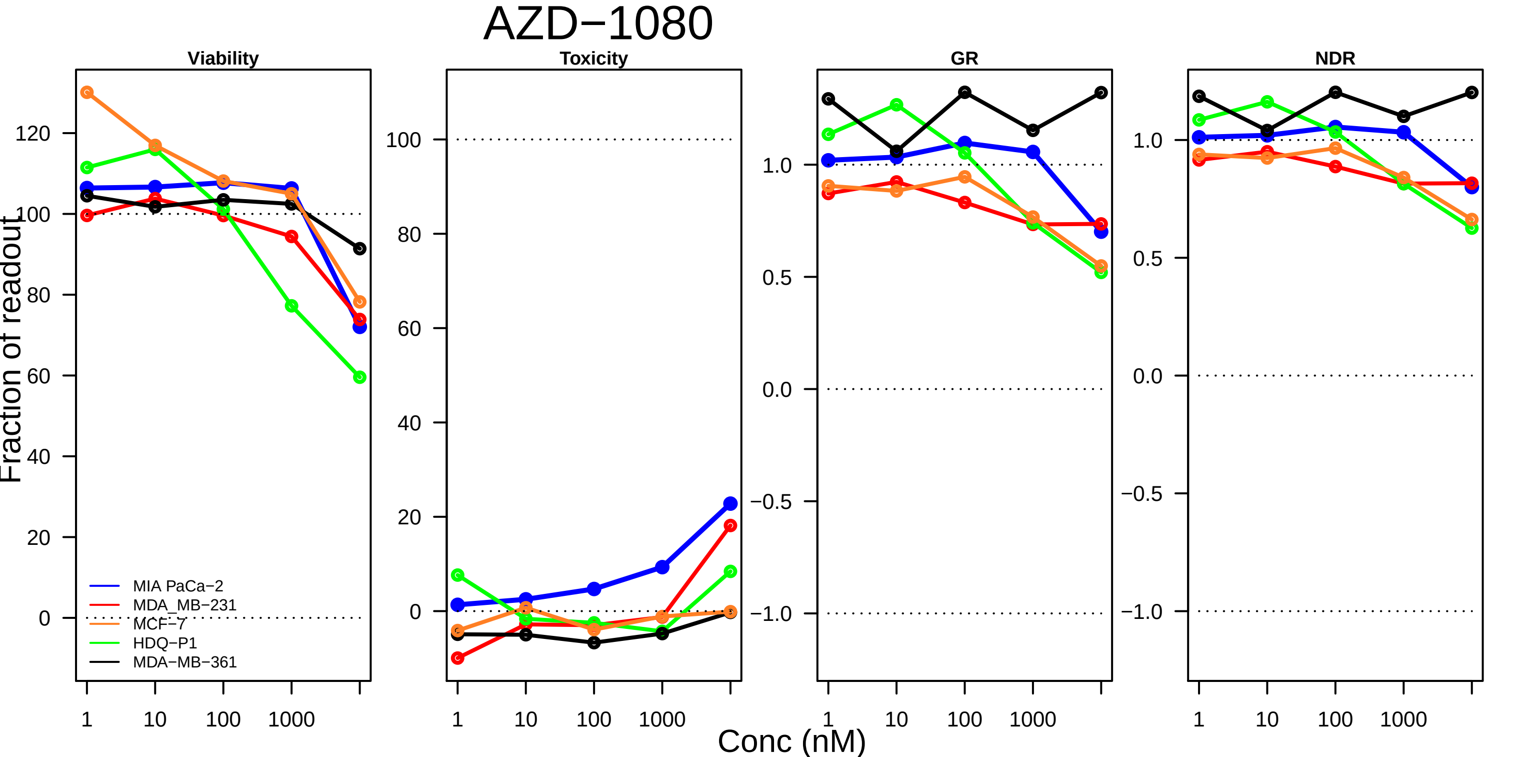

AZD−1080
Viability
Toxicity
GR
NDR
Fraction of readout
MIA PaCa−2
MDA_MB−231
MCF−7
HDQ−P1
MDA−MB−361
Conc (nM)
1
10
100
1000

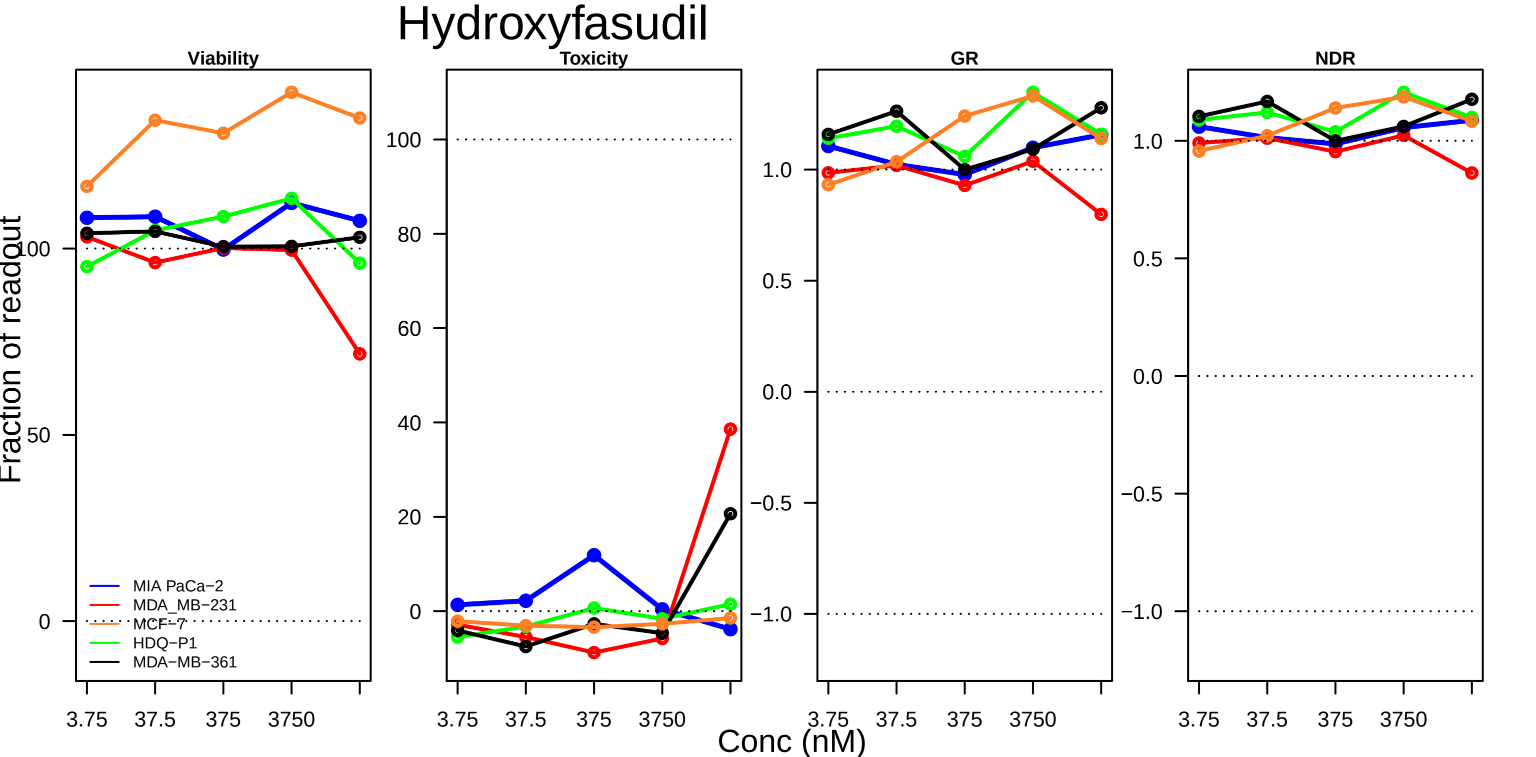

Hydroxyfasudil
Viability
Toxicity
GR
NDR
Fraction of readout
Conc (nM)
3.75
37.5
375
3750
MIA PaCa−2
MDA_MB−231
MCF−7
HDQ−P1
MDA−MB−361
100
50
0
80
60
40
20
1.0
0.5
0.0
−0.5
−1.0

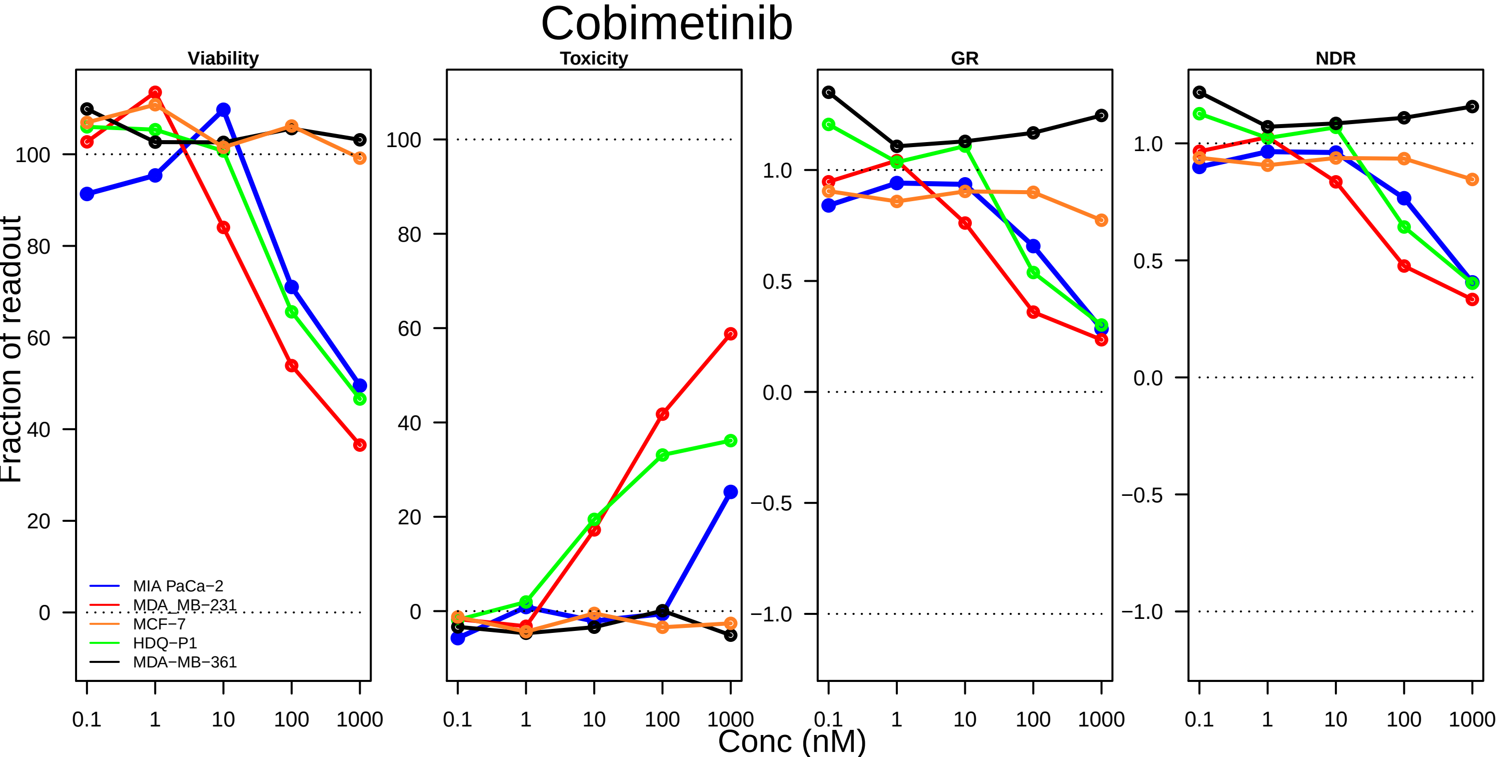

Cobimetinib
Viability
Toxicity
GR
NDR
Fraction of readout
Conc (nM)
0.1
1
10
100
1000
MIA PaCa−2
MDA_MB−231
MCF−7
HDQ−P1
MDA−MB−361

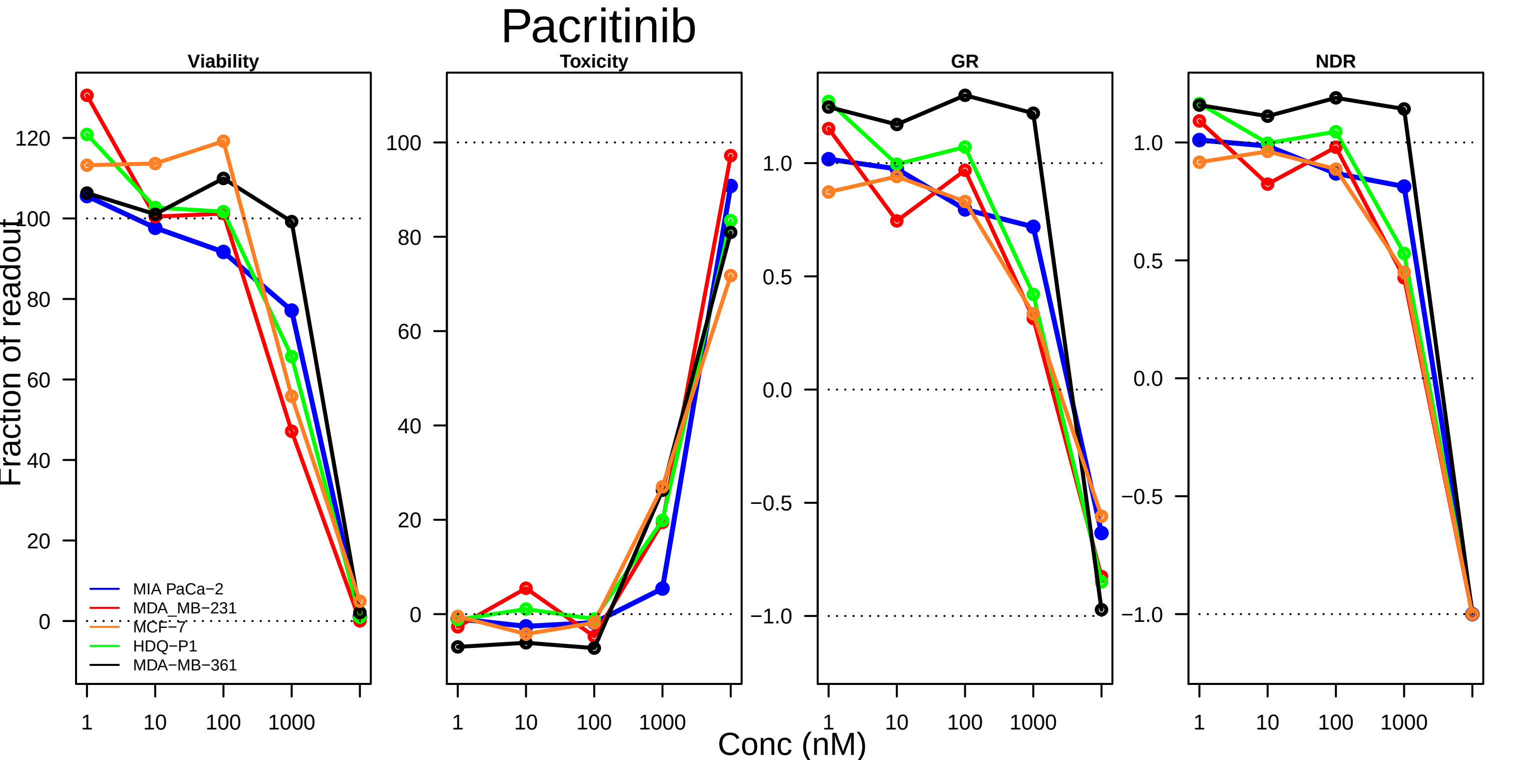

Pacritinib
Viability
Toxicity
GR
NDR
Fraction of readout
Conc (nM)
1
10
100
1000
120
100
80
60
40
20
0
1.0
0.5
0.0
−0.5
−1.0
MIA PaCa−2
MDA_MB−231
MCF−7
HDQ−P1
MDA−MB−361

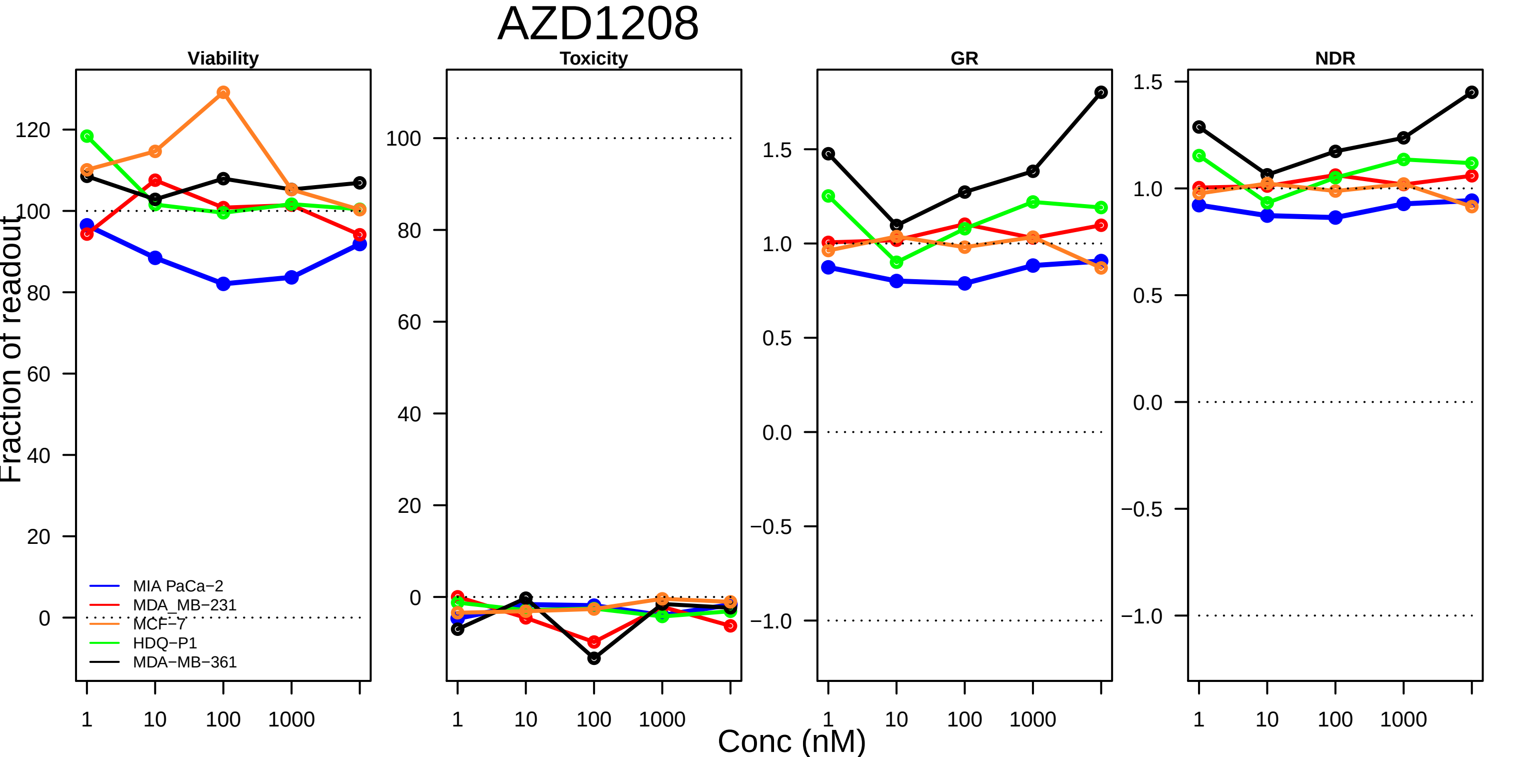

AZD1208
Viability
Toxicity
GR
NDR
Fraction of readout
Conc (nM)
1
10
100
1000
MIA PaCa−2
MDA_MB−231
MCF−7
HDQ−P1
MDA−MB−361

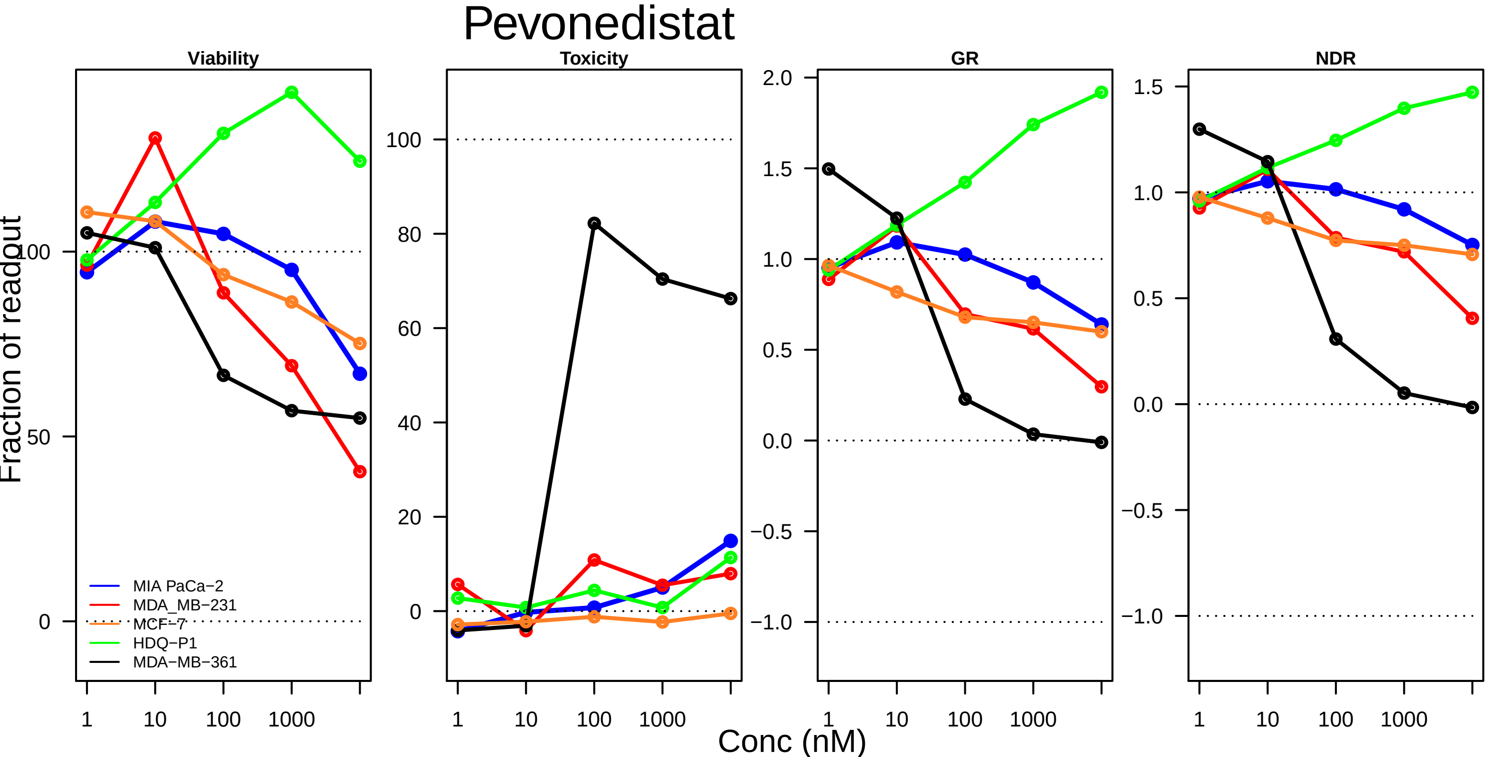

Pevonedistat
Viability
Toxicity
GR
NDR
Fraction of readout
Conc (nM)
MIA PaCa−2
MDA_MB−231
MCF−7
HDQ−P1
MDA−MB−361
0
50
100
20
40
60
80
−1.0
−0.5
0.0
0.5
1.0
1.5
2.0
1
10
100
1000

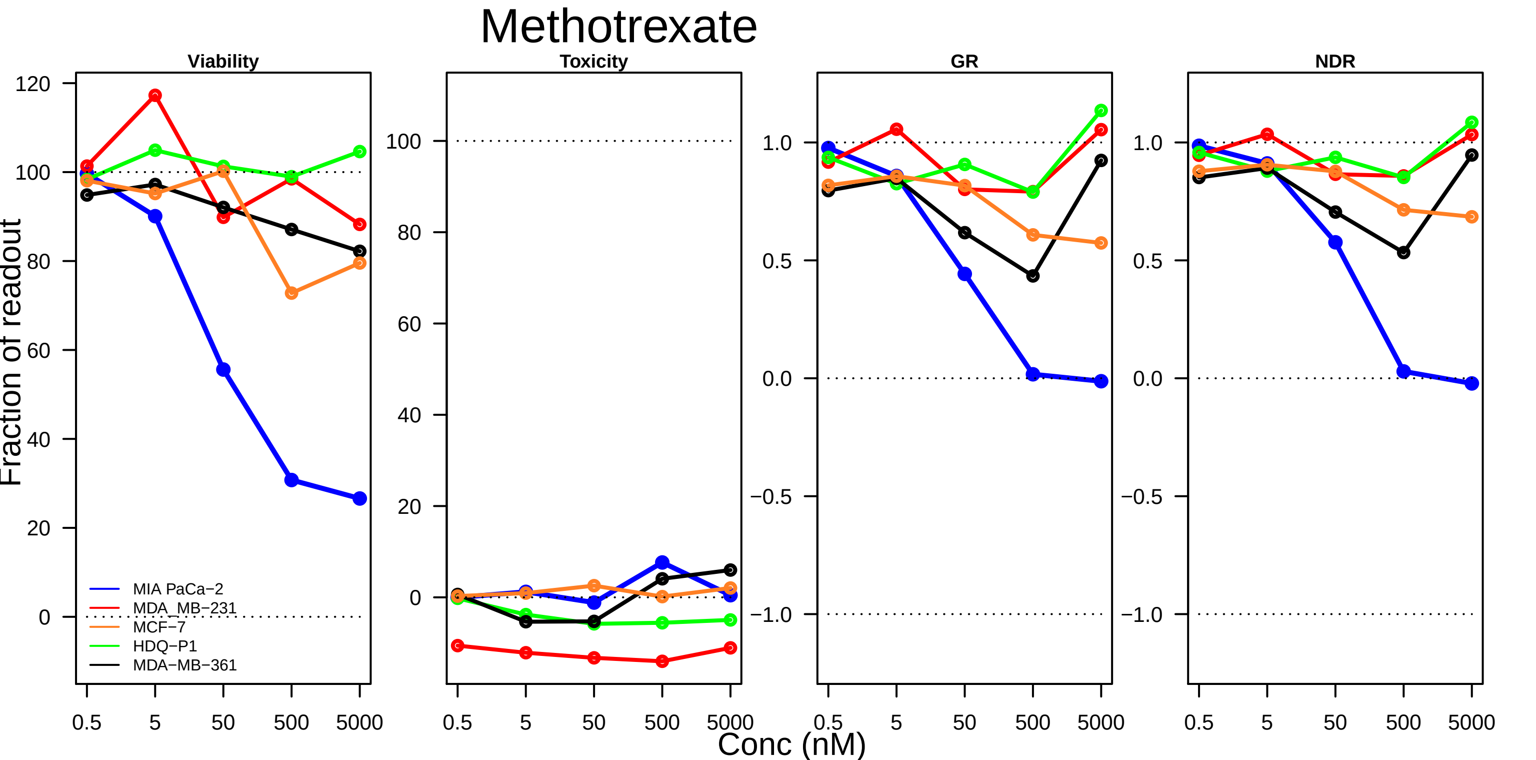

Methotrexate
Viability
Toxicity
GR
NDR
Fraction of readout
Conc (nM)
MIA PaCa−2
MDA_MB−231
MCF−7
HDQ−P1
MDA−MB−361
120
100
80
60
40
20
0
−0.5
−1.0
0.5
1.0
0.0
5
50
500
5000

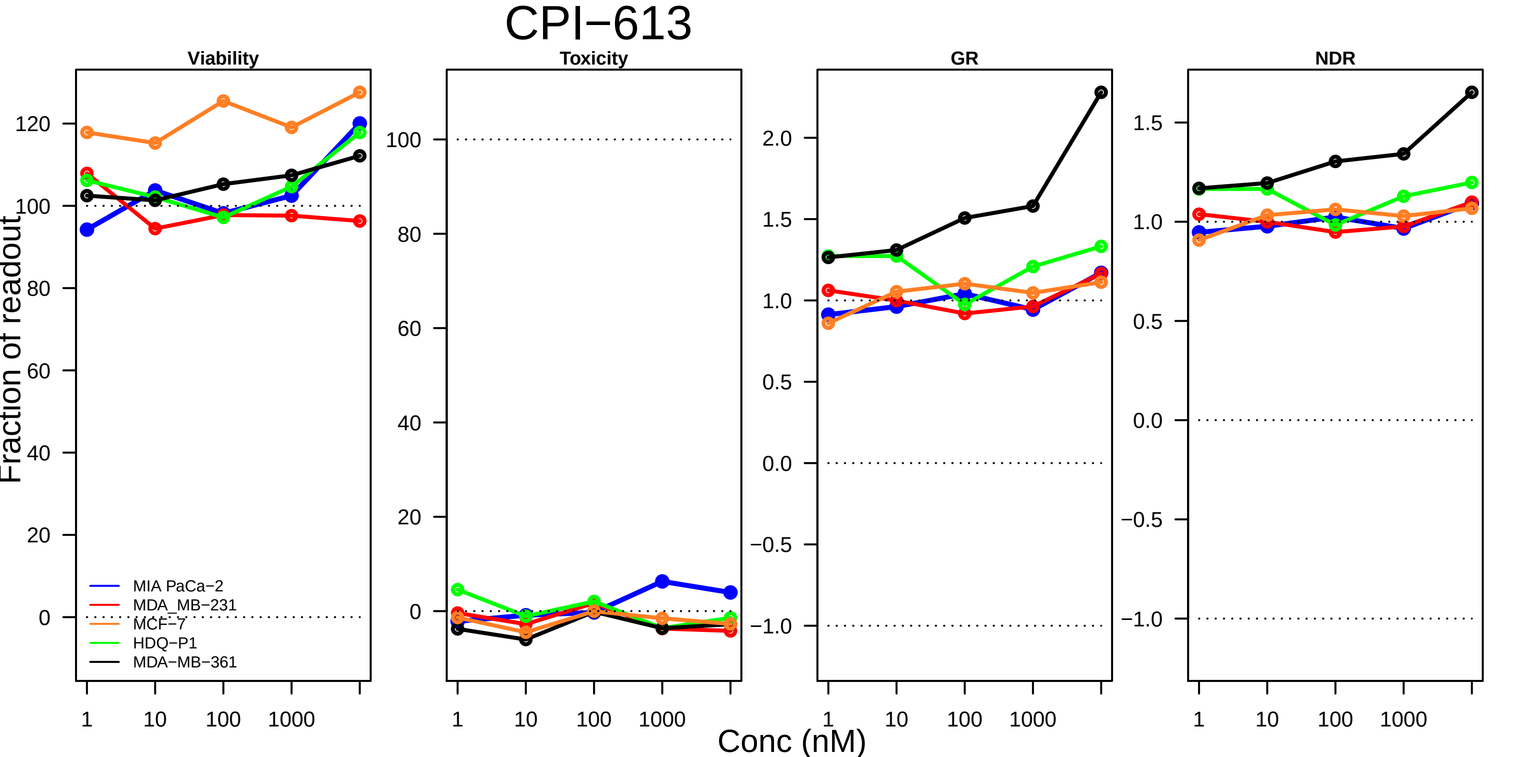

CPI−613
Viability
Toxicity
GR
NDR
Fraction of readout
Conc (nM)
MIA PaCa−2
MDA_MB−231
MCF−7
HDQ−P1
MDA−MB−361
120
100
80
60
40
20
0
2.0
1.5
1.0
0.5
0.0
−0.5
−1.0
1
10
100
1000

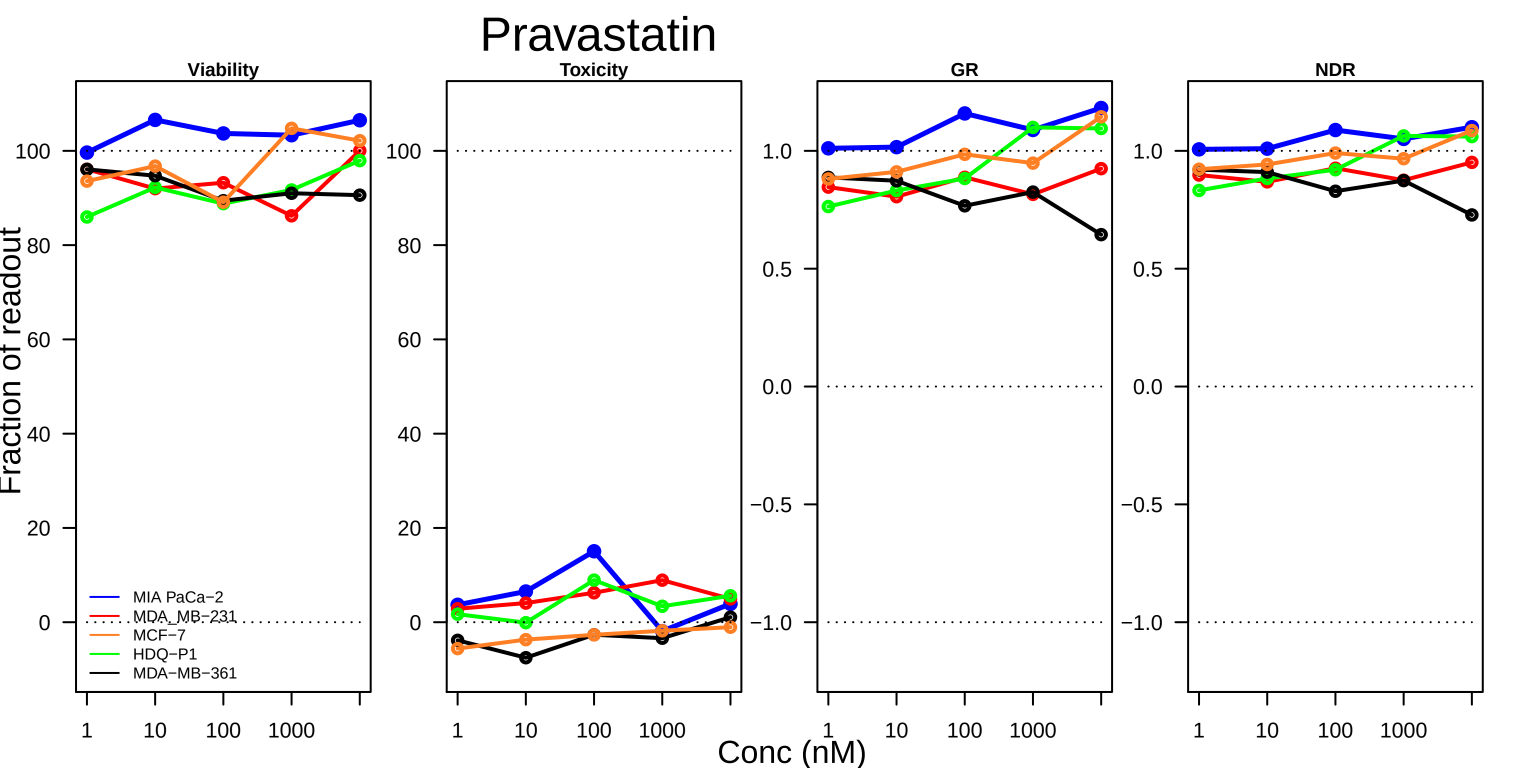

Pravastatin
Viability
Toxicity
GR
NDR
Fraction of readout
Conc (nM)
1
10
100
1000
0
20
40
60
80
100
−1.0
−0.5
0.0
0.5
1.0
MIA PaCa−2
MDA_MB−231
MCF−7
HDQ−P1
MDA−MB−361

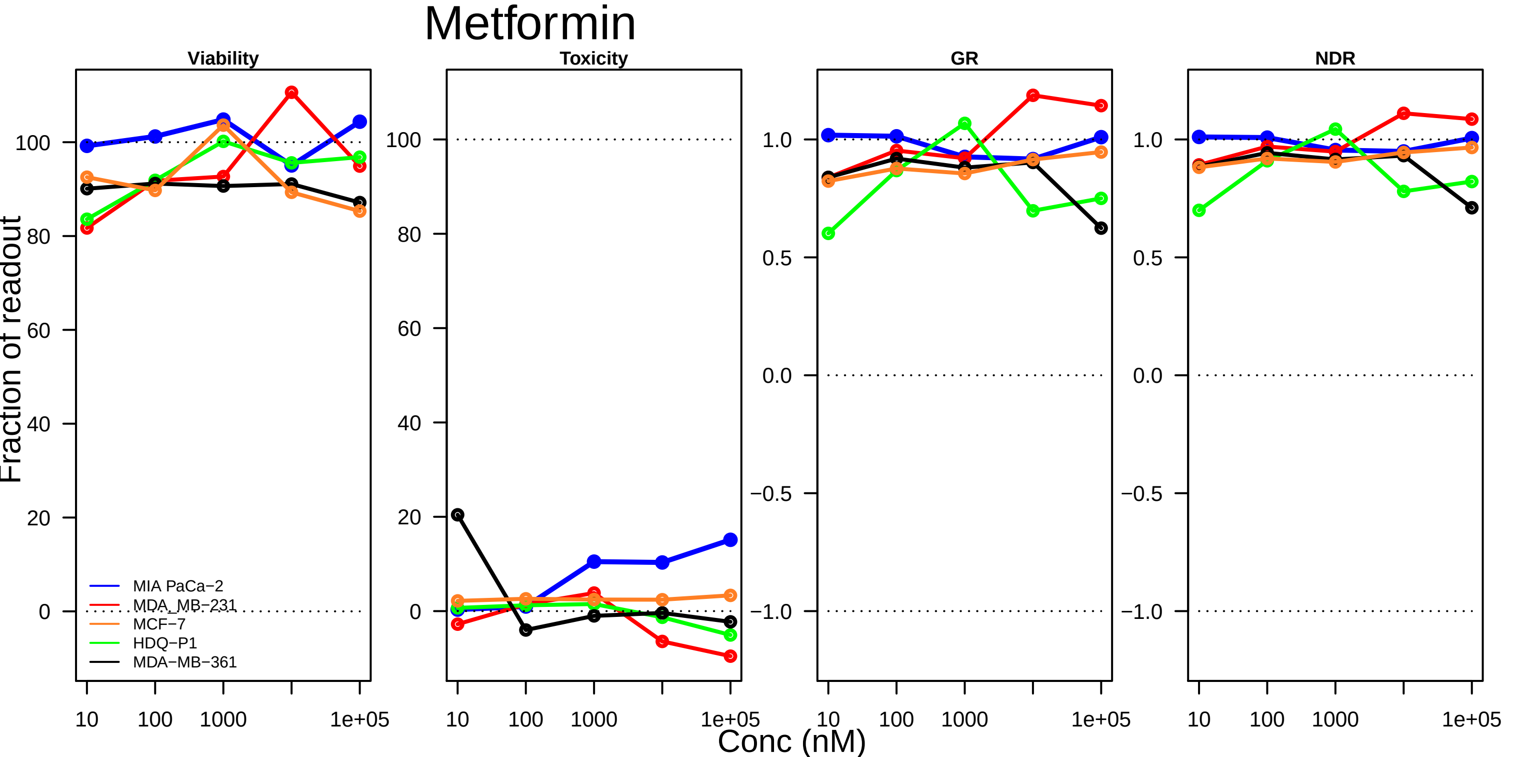

Metformin
Viability
Toxicity
GR
NDR
Fraction of readout
Conc (nM)
MIA PaCa−2
MDA_MB−231
MCF−7
HDQ−P1
MDA−MB−361
10
100
1000
1e+05

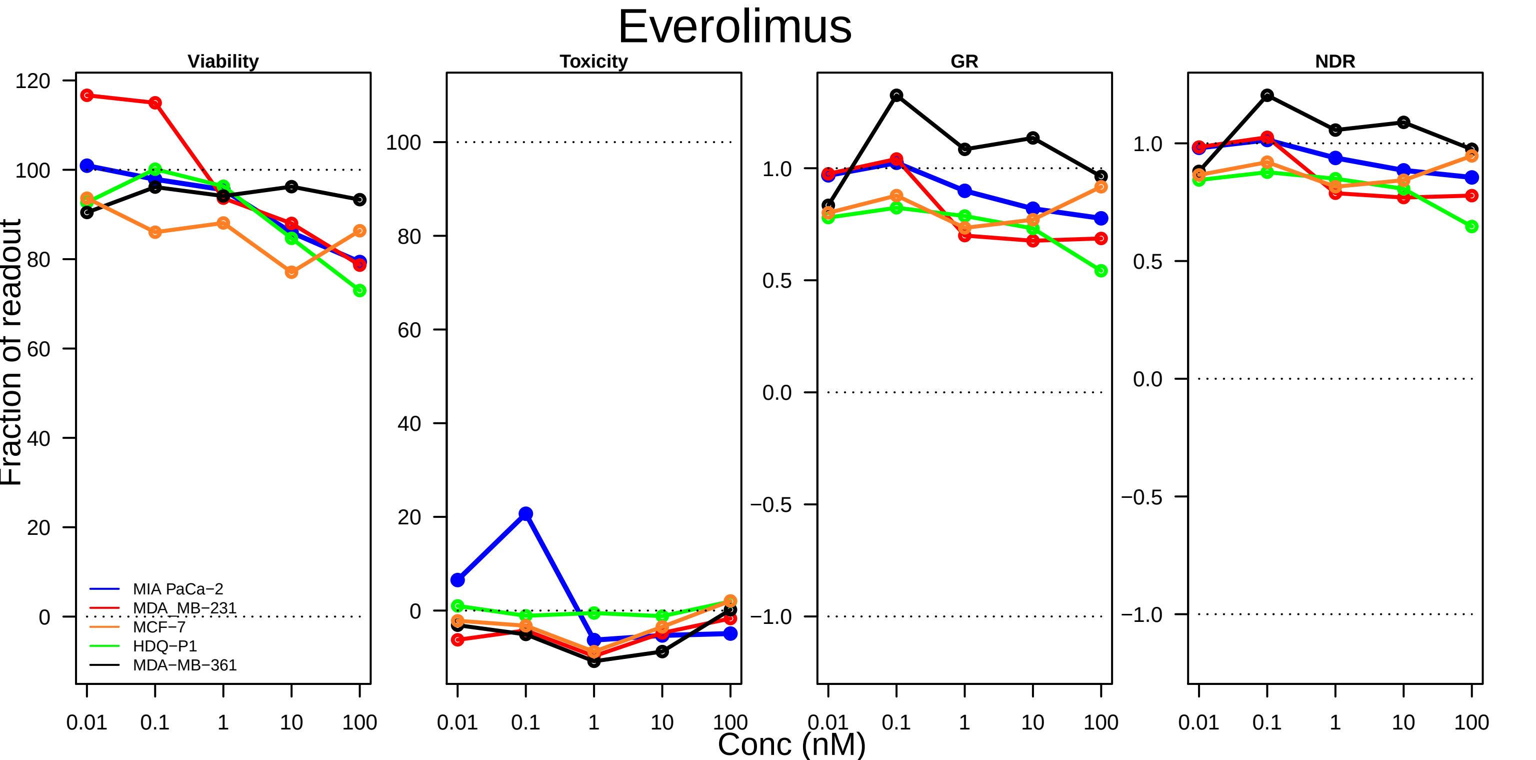

Everolimus
Viability
Toxicity
GR
NDR
Fraction of readout
Conc (nM)
MIA PaCa−2
MDA_MB−231
MCF−7
HDQ−P1
MDA−MB−361
120
100
80
60
40
20
0
1.0
0.5
0.0
−0.5
−1.0
0.01
0.1
1
10
100

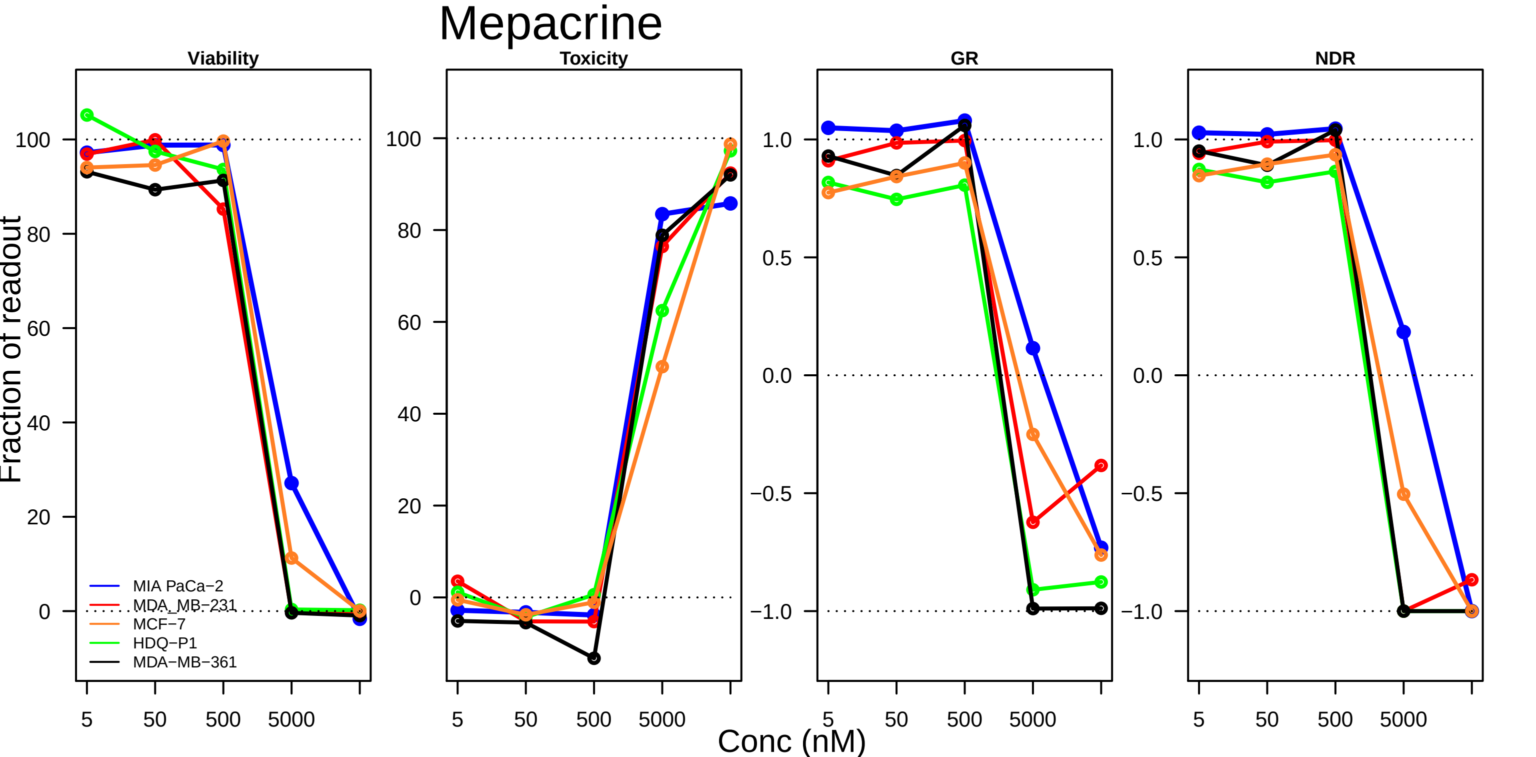
Mepacrine
Viability
Toxicity
GR
NDR
Fraction of readout
MIA PaCa−2
MDA_MB−231
MCF−7
HDQ−P1
MDA−MB−361
Conc (nM)
5
50
500
5000
100
80
60
40
20
0
1.0
0.5
0.0
−0.5
−1.0

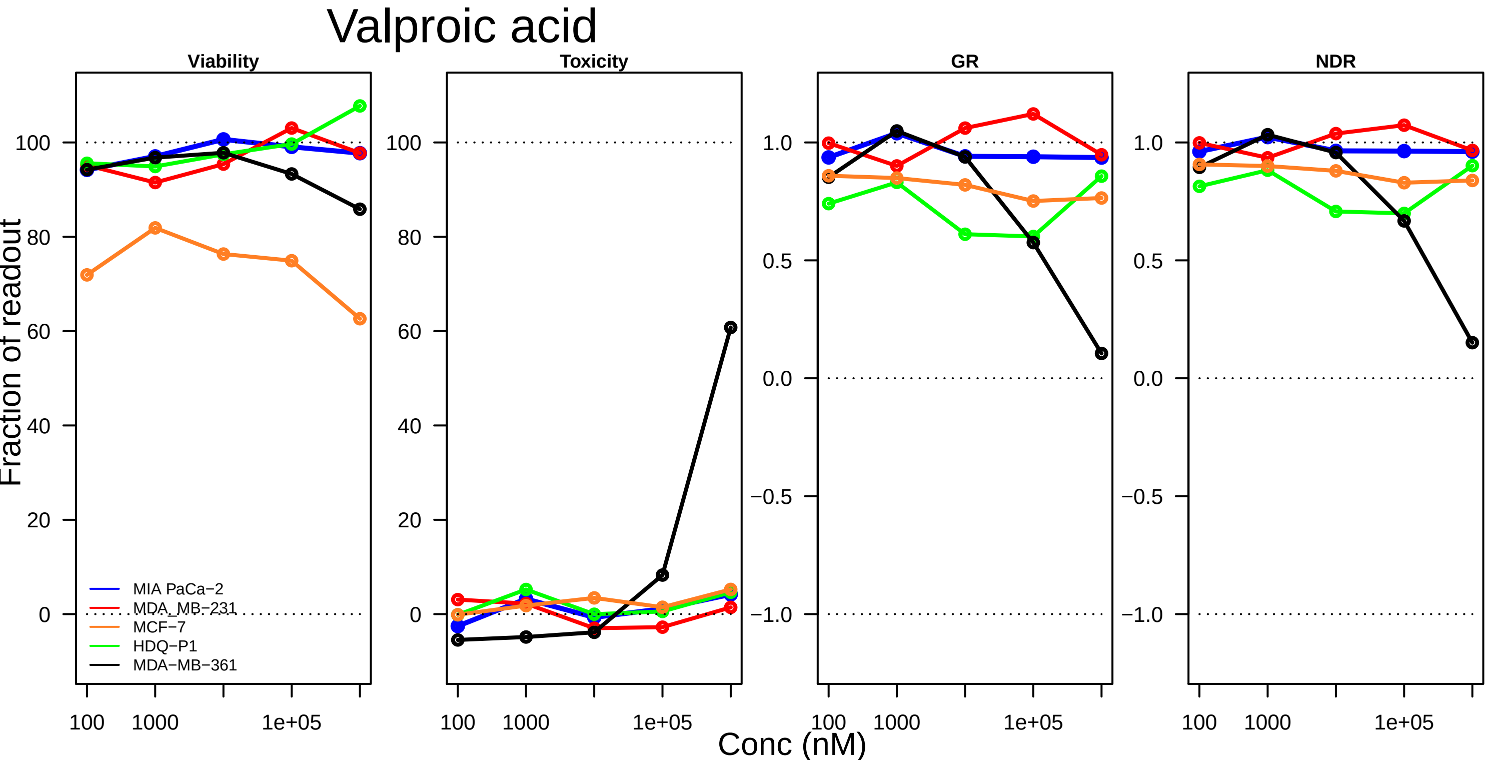

Valproic acid
Viability
Toxicity
GR
NDR
Fraction of readout
Conc (nM)
MIA PaCa−2
MDA_MB−231
MCF−7
HDQ−P1
MDA−MB−361
100
80
60
40
20
0
1.0
0.5
0.0
−0.5
−1.0
100
1000
1e+05

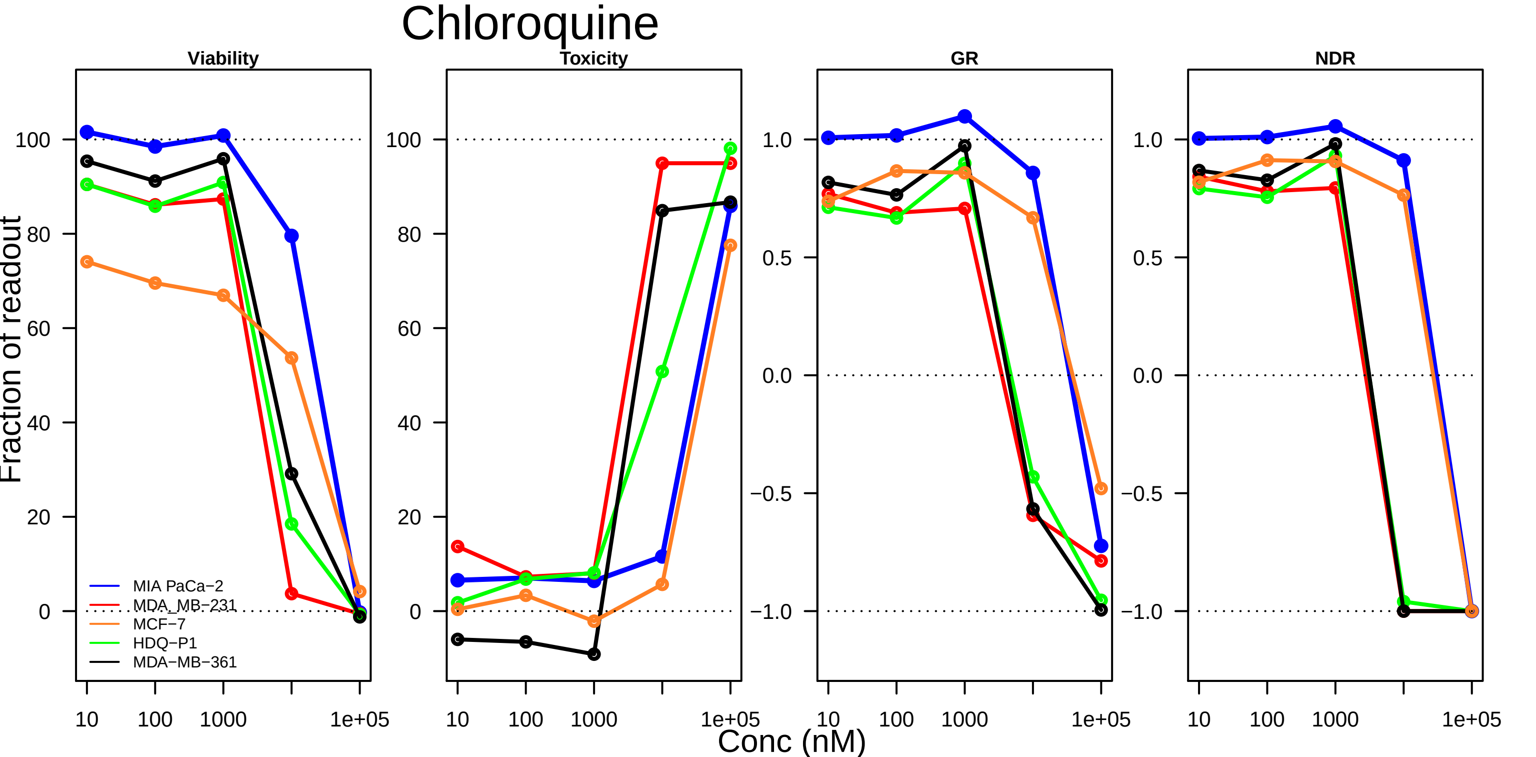

Chloroquine
Viability
Toxicity
GR
NDR
Fraction of readout
100
80
60
40
20
0
1.0
0.5
0.0
−0.5
−1.0
10
100
1000
1e+05
Conc (nM)
MIA PaCa−2
MDA_MB−231
MCF−7
HDQ−P1
MDA−MB−361

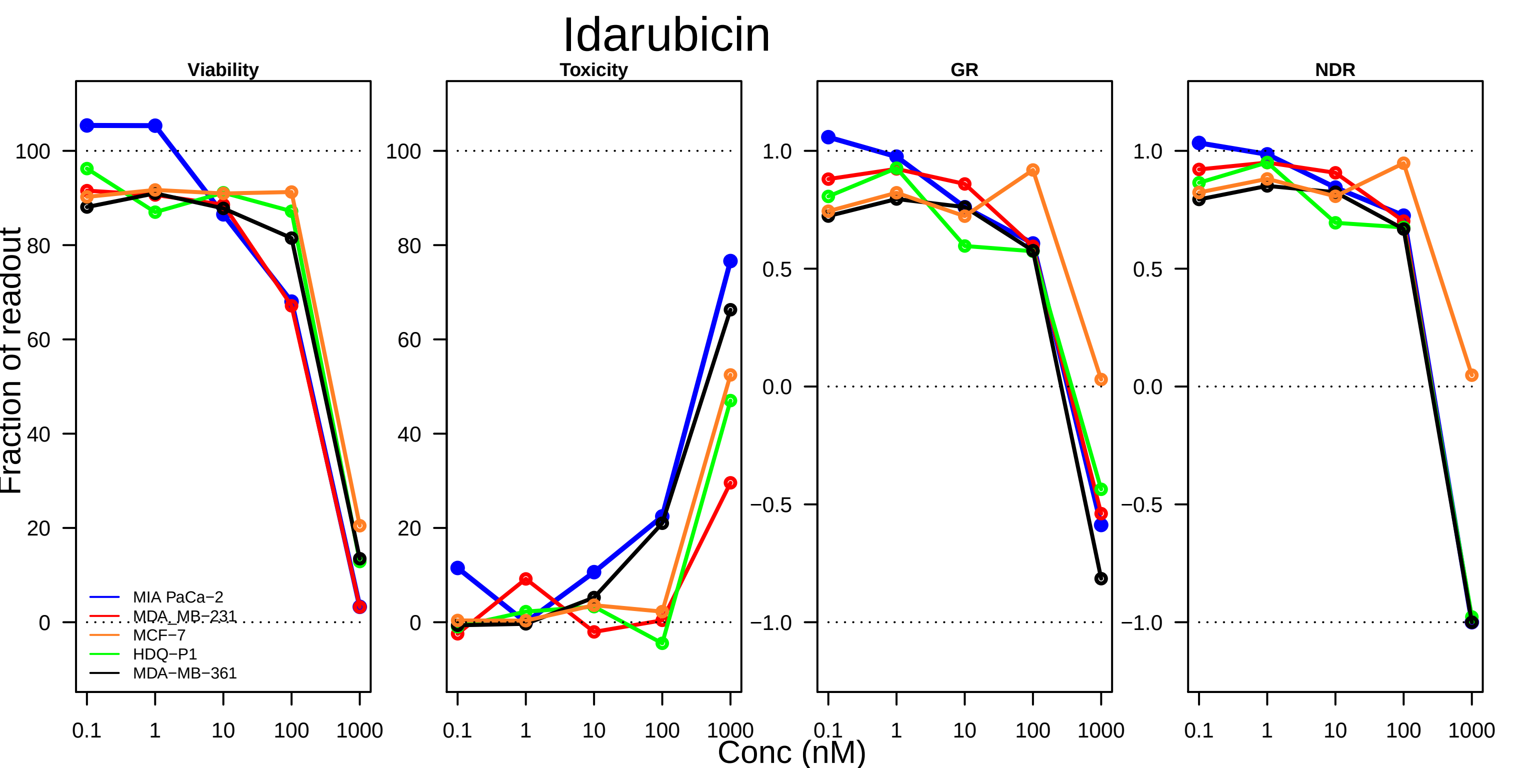

Idarubicin
Viability
Toxicity
GR
NDR
Fraction of readout
100
80
60
40
20
0
1.0
0.5
0.0
−0.5
−1.0
0.1
1
10
100
1000
Conc (nM)
MIA PaCa−2
MDA_MB−231
MCF−7
HDQ−P1
MDA−MB−361

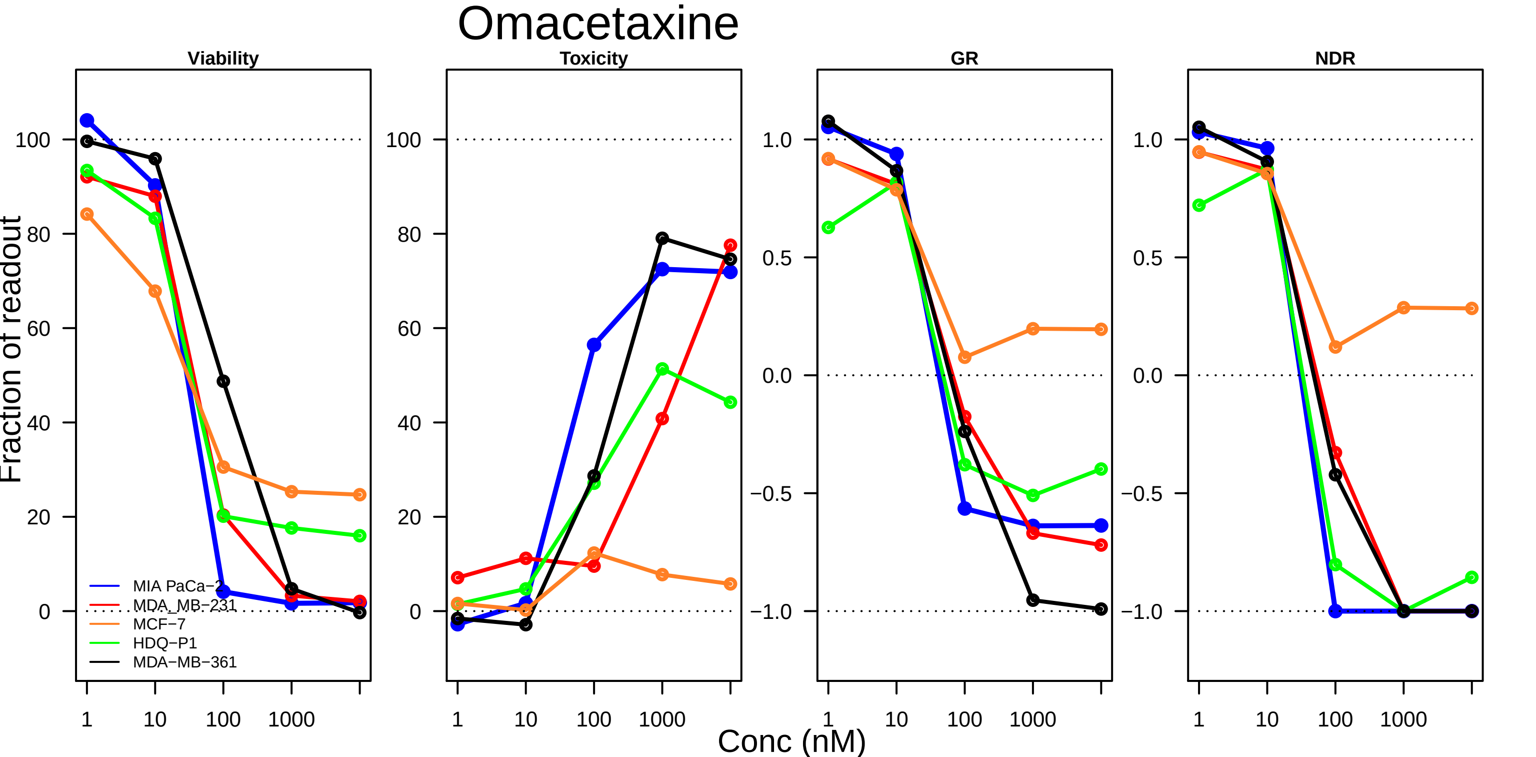

Omacetaxine
Viability
Toxicity
GR
NDR
Fraction of readout
Conc (nM)
MIA PaCa−2
MDA_MB−231
MCF−7
HDQ−P1
MDA−MB−361

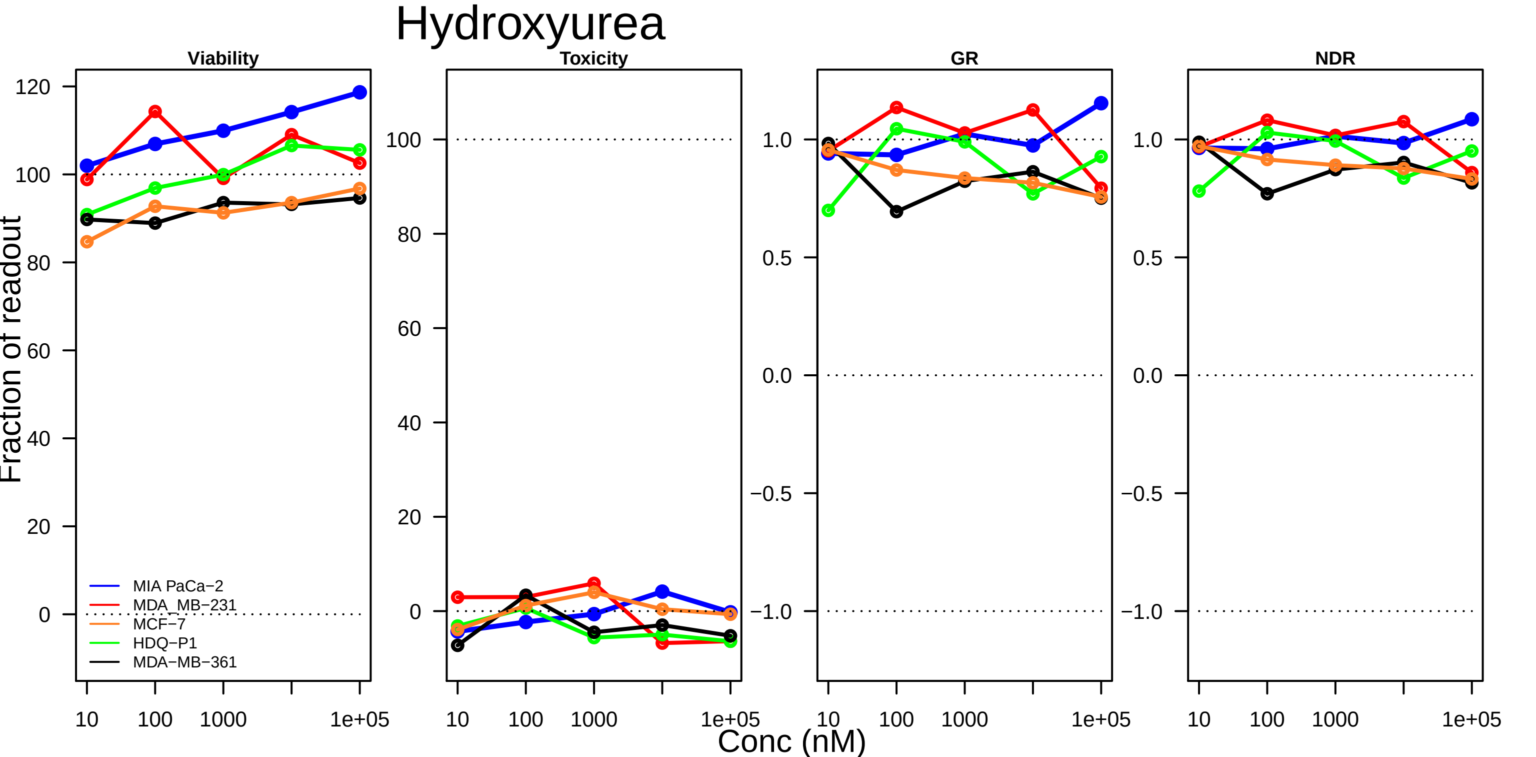

Hydroxyurea
Viability
Toxicity
GR
NDR
Fraction of readout
Conc (nM)
MIA PaCa−2
MDA_MB−231
MCF−7
HDQ−P1
MDA−MB−361
120
100
80
60
40
20
0
1.0
0.5
0.0
−0.5
−1.0
10
100
1000
1e+05

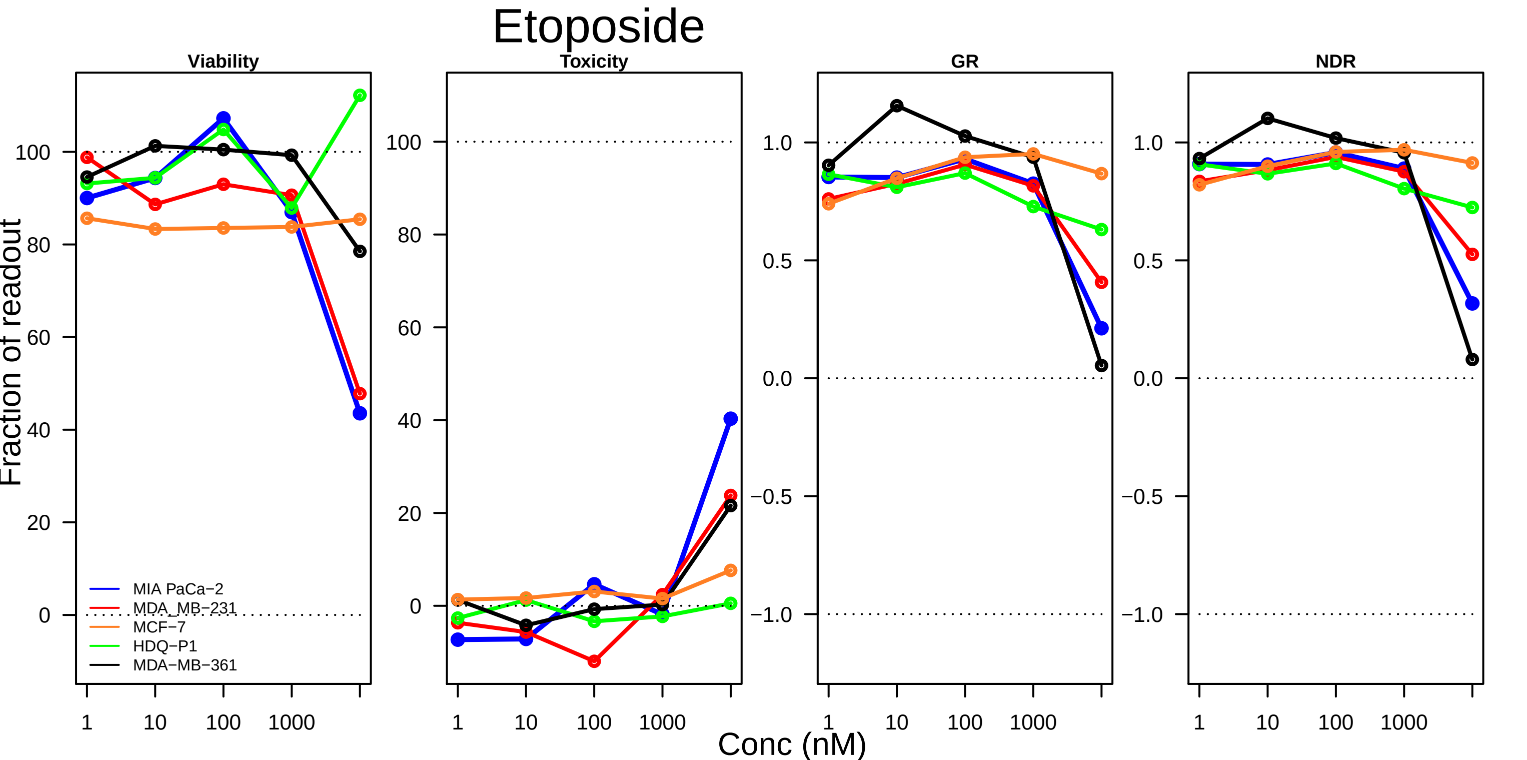

Etoposide
Viability
Toxicity
GR
NDR
Fraction of readout
Conc (nM)
1
10
100
1000
MIA PaCa−2
MDA_MB−231
MCF−7
HDQ−P1
MDA−MB−361
0
20
40
60
80
100
−1.0
−0.5
0.0
0.5
1.0

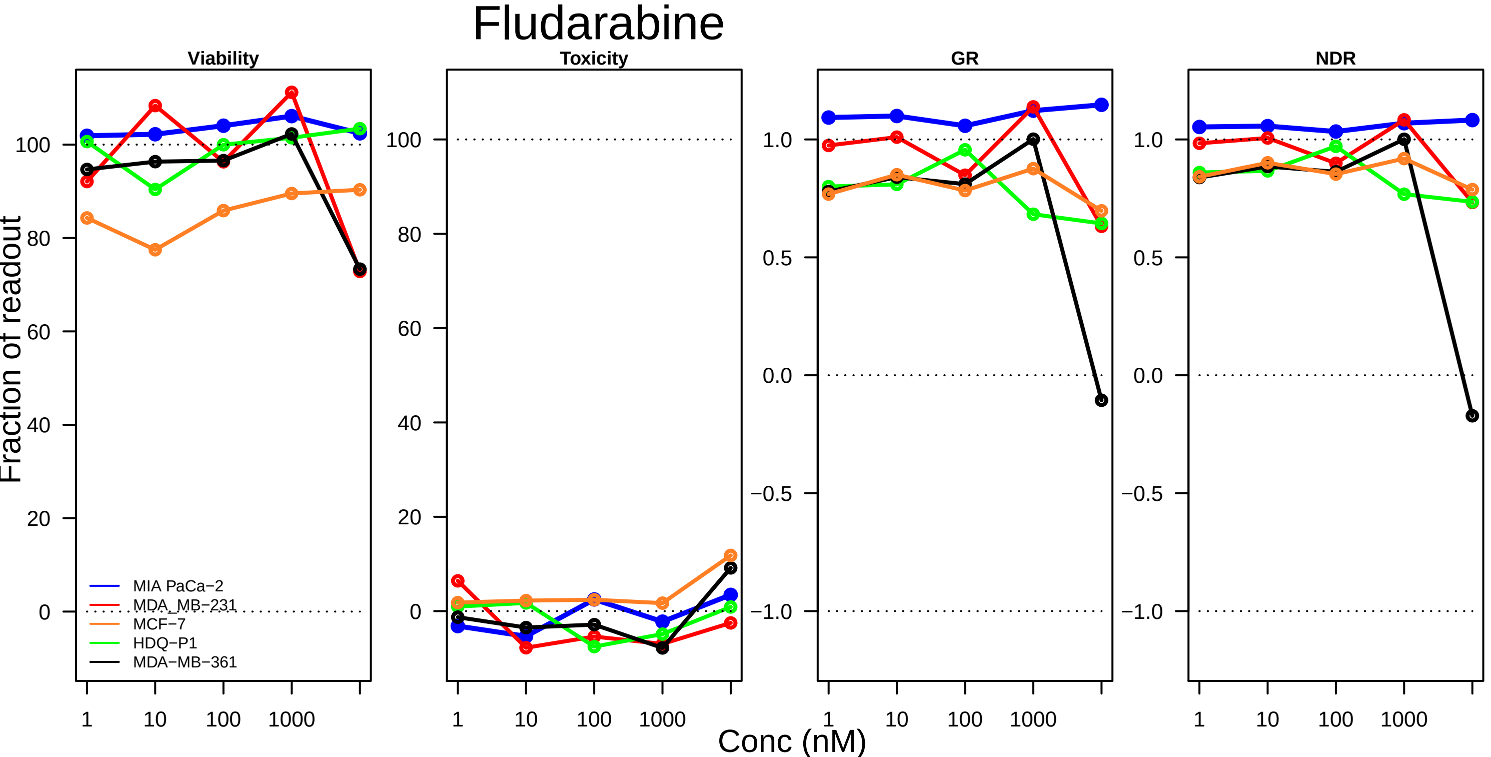

Fludarabine
Viability
Toxicity
GR
NDR
Fraction of readout
Conc (nM)
1
10
100
1000
100
80
60
40
20
0
1.0
0.5
0.0
−0.5
−1.0
MIA PaCa−2
MDA_MB−231
MCF−7
HDQ−P1
MDA−MB−361

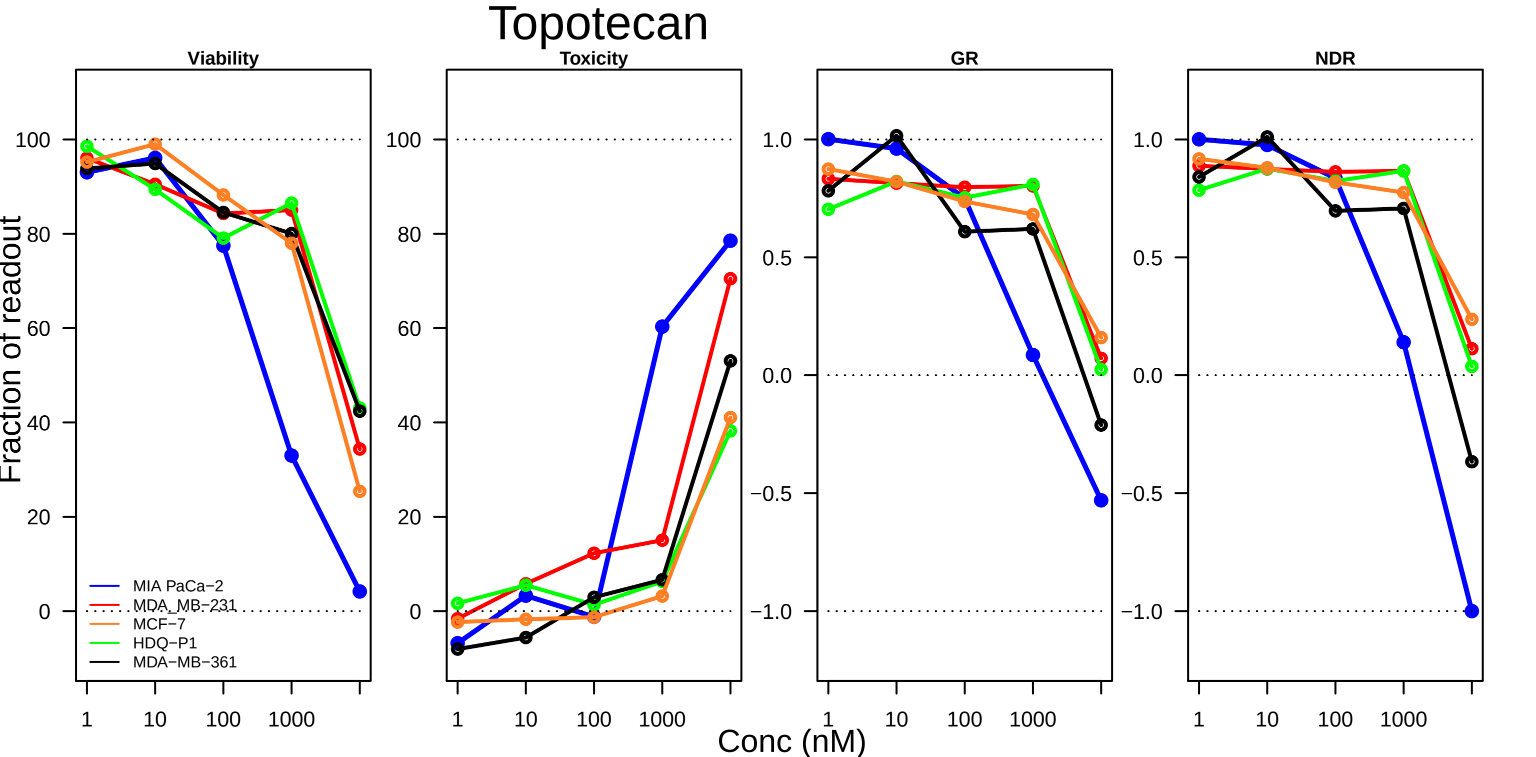

Topotecan
Viability
Toxicity
GR
NDR
Fraction of readout
Conc (nM)
1
10
100
1000
100
80
60
40
20
0
1.0
0.5
0.0
−0.5
−1.0
MIA PaCa−2
MDA_MB−231
MCF−7
HDQ−P1
MDA−MB−361

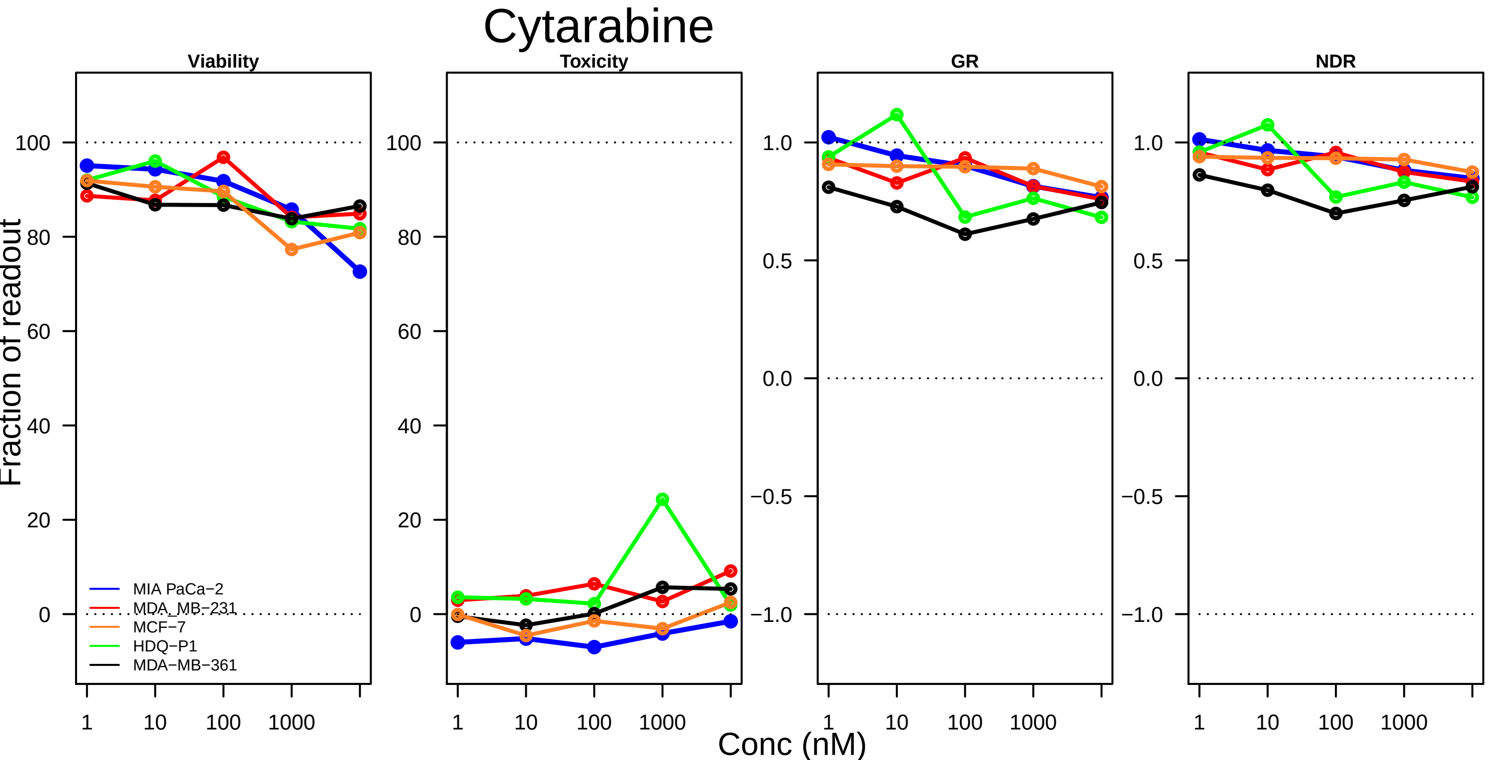
Cytarabine
Viability
Toxicity
GR
NDR
Fraction of readout
Conc (nM)
MIA PaCa−2
MDA_MB−231
MCF−7
HDQ−P1
MDA−MB−361
100
80
60
40
20
0
1.0
0.5
0.0
−0.5
−1.0
1
10
100
1000

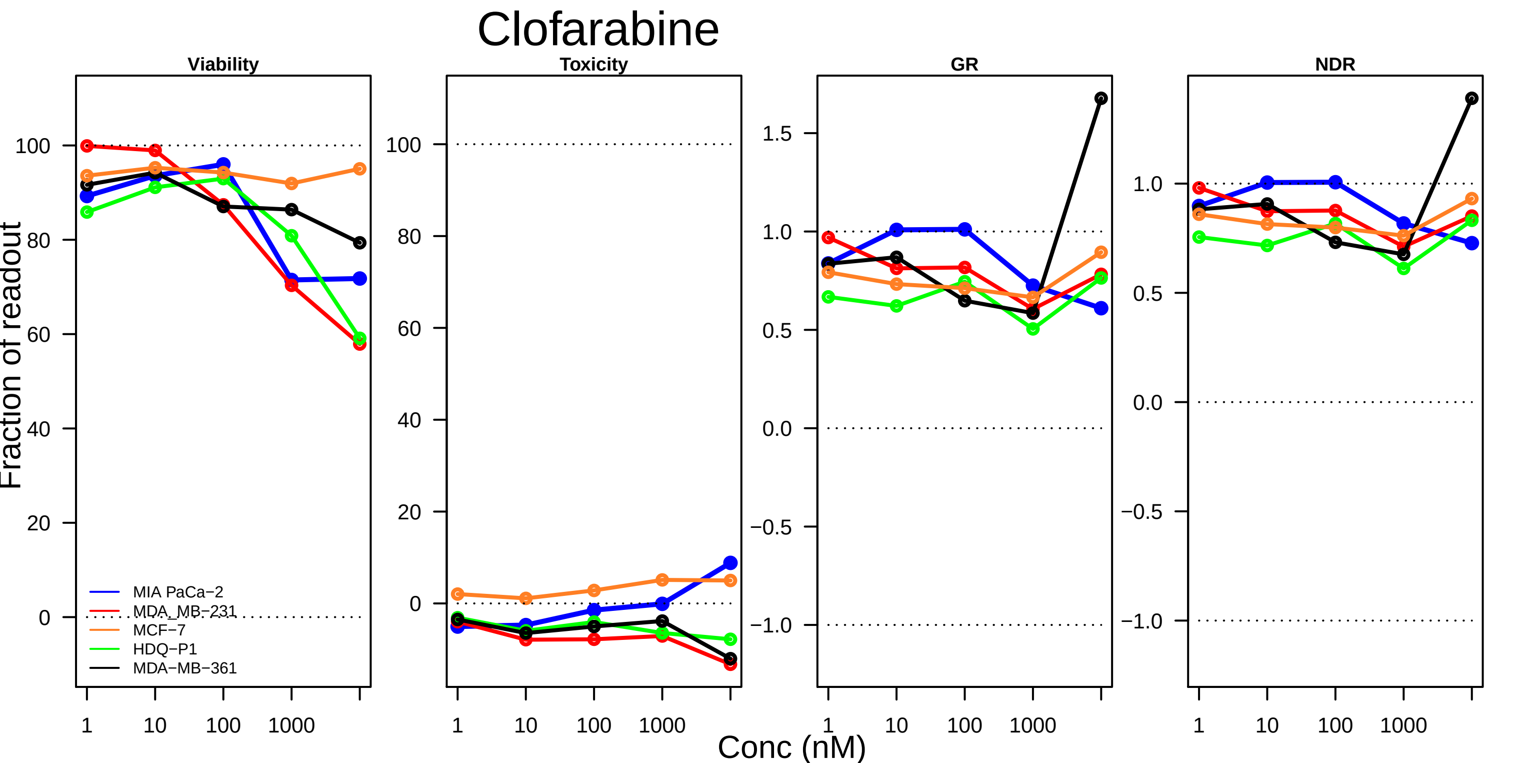

Clofarabine
Viability
Toxicity
GR
NDR
Fraction of readout
Conc (nM)
MIA PaCa−2
MDA_MB−231
MCF−7
HDQ−P1
MDA−MB−361
1
10
100
1000

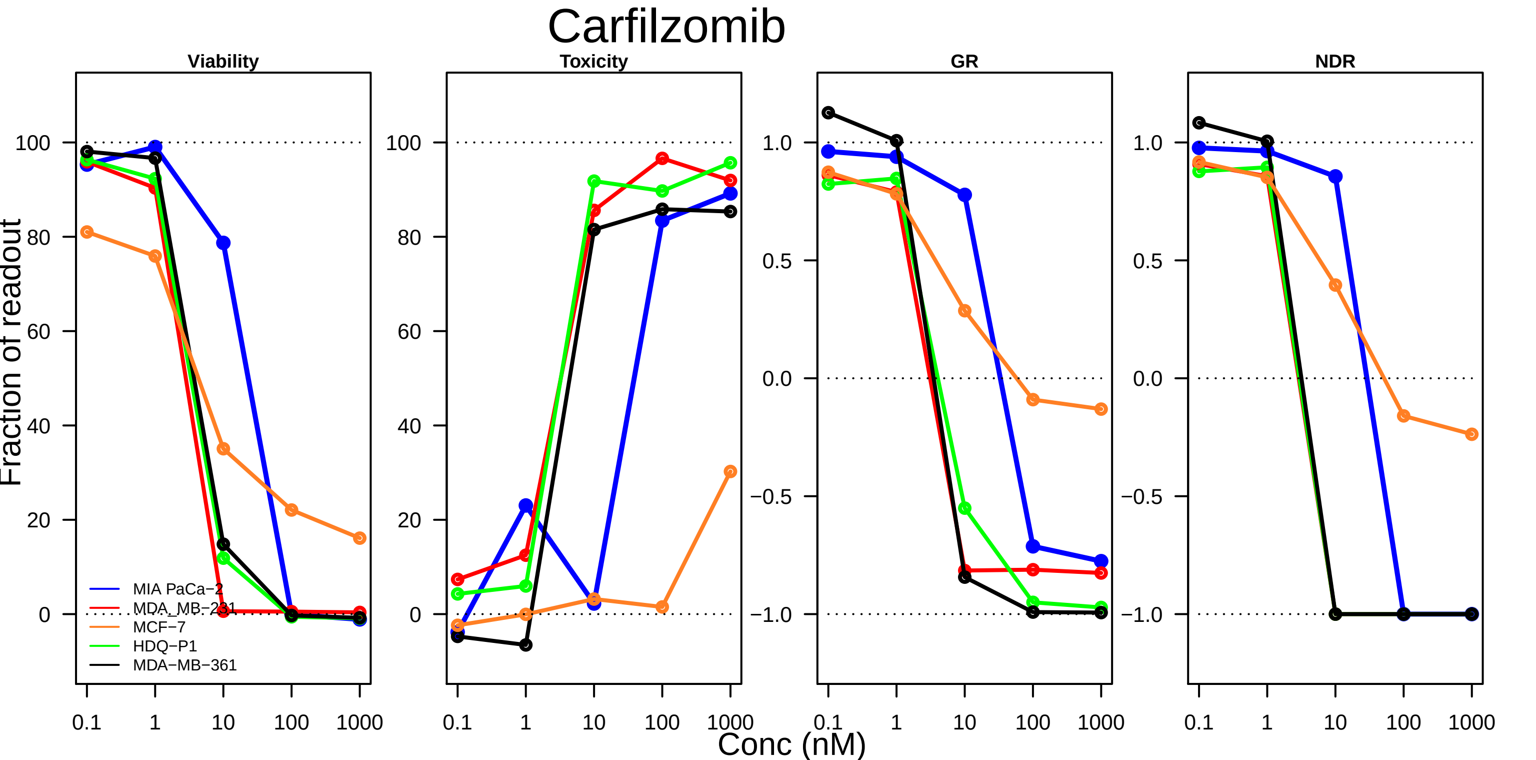

Carfilzomib
Viability
Toxicity
GR
NDR
Fraction of readout
Conc (nM)
0.1
1
10
100
1000
0
20
40
60
80
100
−1.0
−0.5
0.0
0.5
1.0
MIA PaCa−2
MDA_MB−231
MCF−7
HDQ−P1
MDA−MB−361

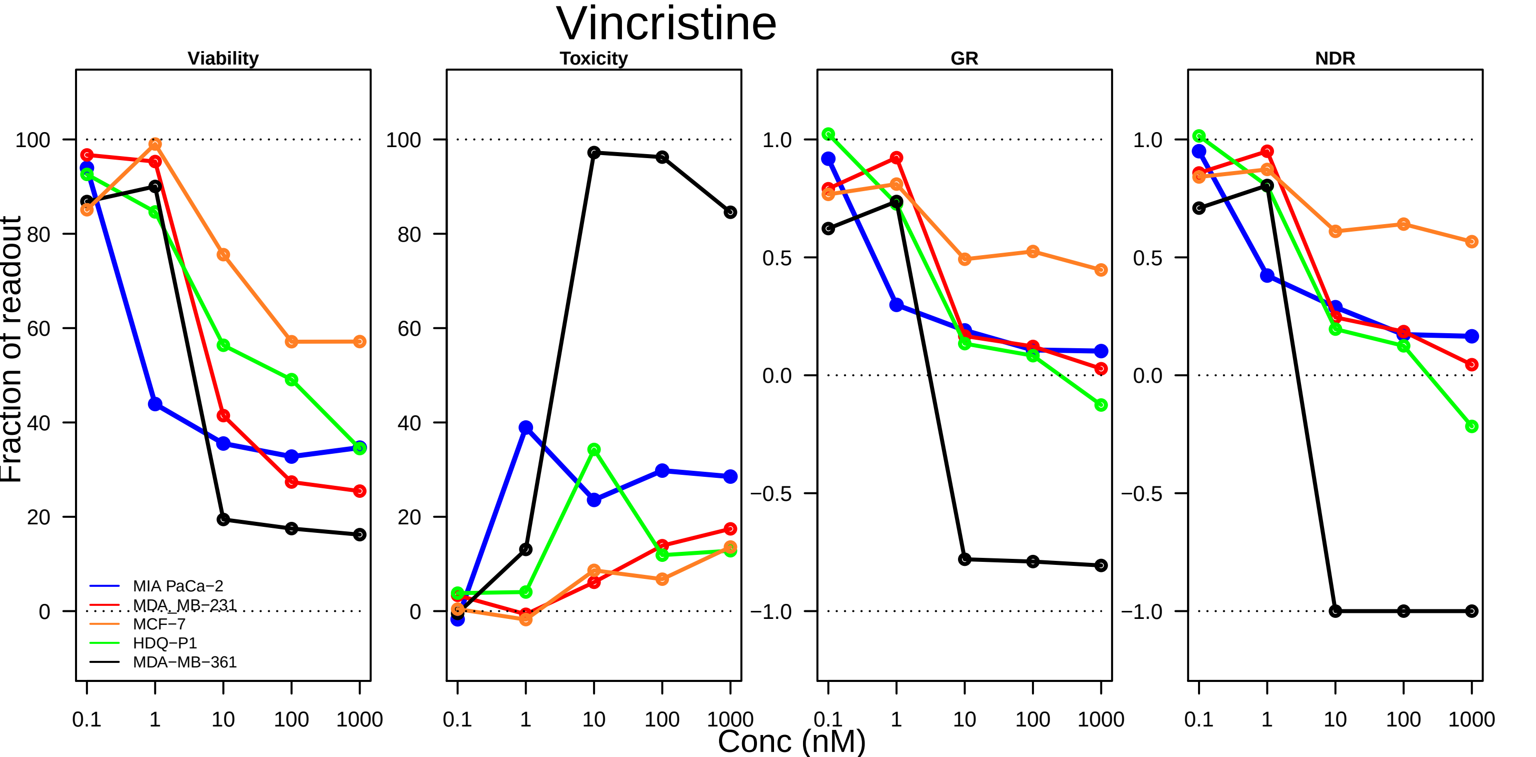

Vincristine
Viability
Toxicity
GR
NDR
Fraction of readout
100
80
60
40
20
0
1.0
0.5
0.0
−0.5
−1.0
0.1
1
10
100
1000
Conc (nM)
MIA PaCa−2
MDA_MB−231
MCF−7
HDQ−P1
MDA−MB−361

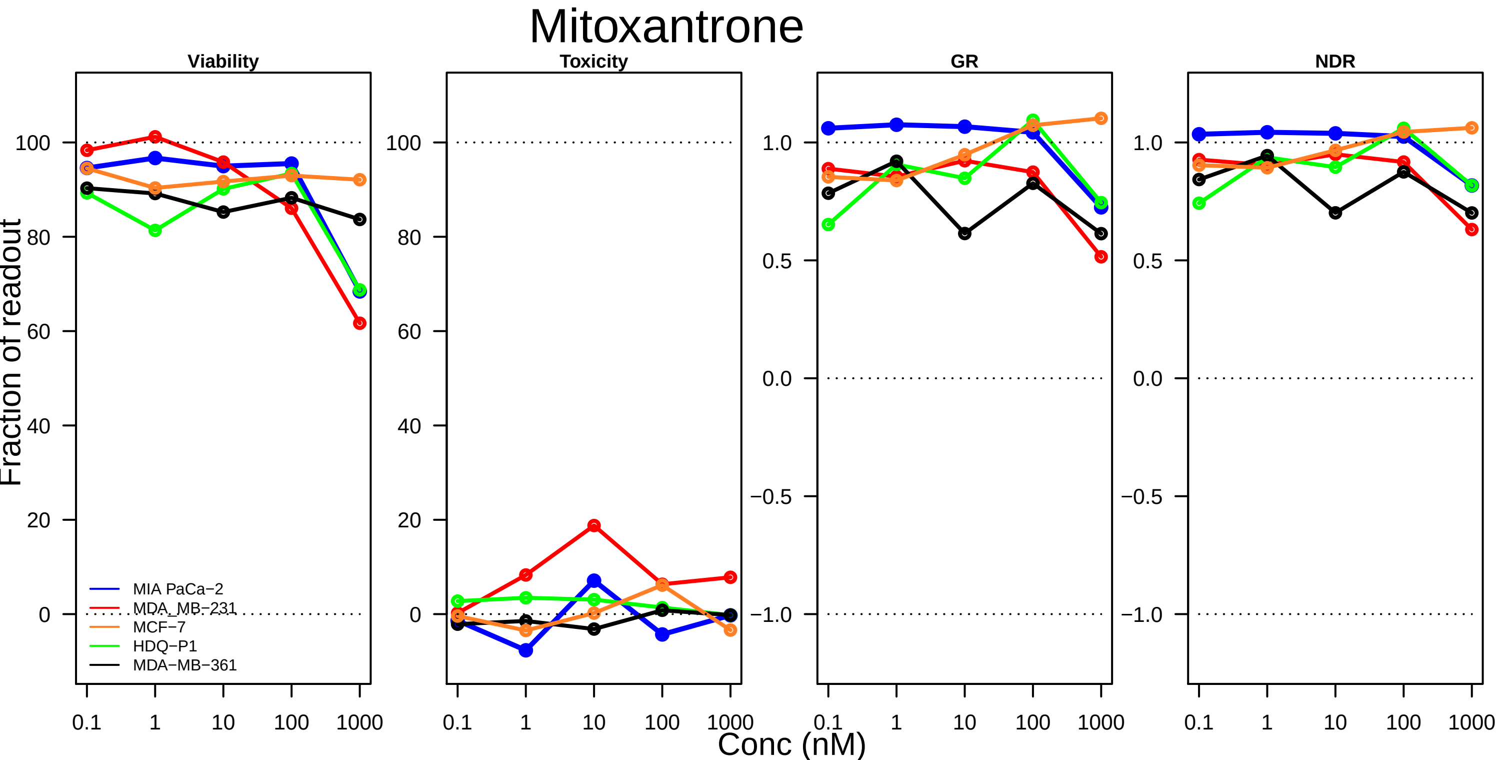

Mitoxantrone
Viability
Toxicity
GR
NDR
Fraction of readout
Conc (nM)
MIA PaCa−2
MDA_MB−231
MCF−7
HDQ−P1
MDA−MB−361
0.1
1
10
100
1000

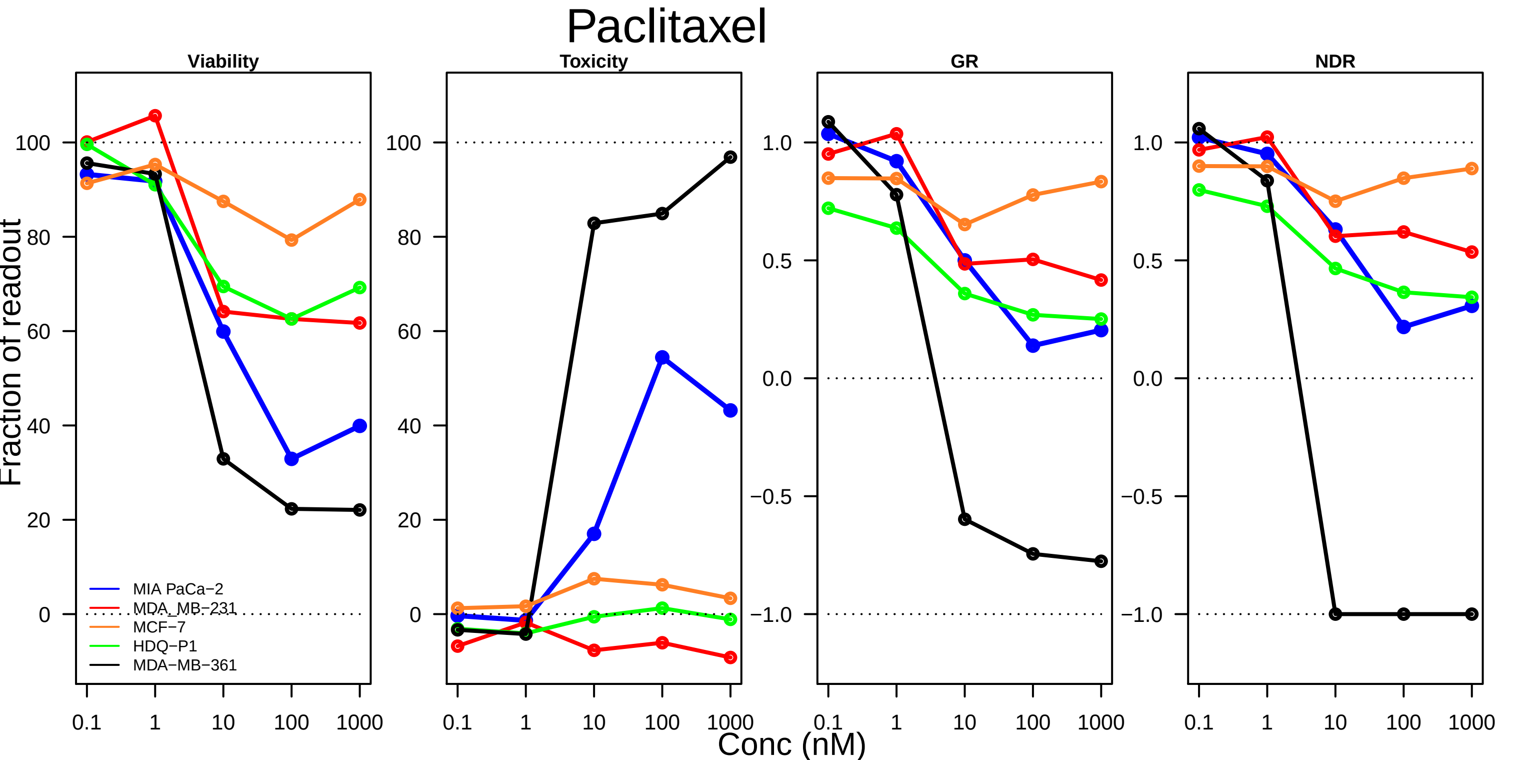

Paclitaxel
Viability
Toxicity
GR
NDR
Fraction of readout
Conc (nM)
0.1
1
10
100
1000
MIA PaCa−2
MDA_MB−231
MCF−7
HDQ−P1
MDA−MB−361

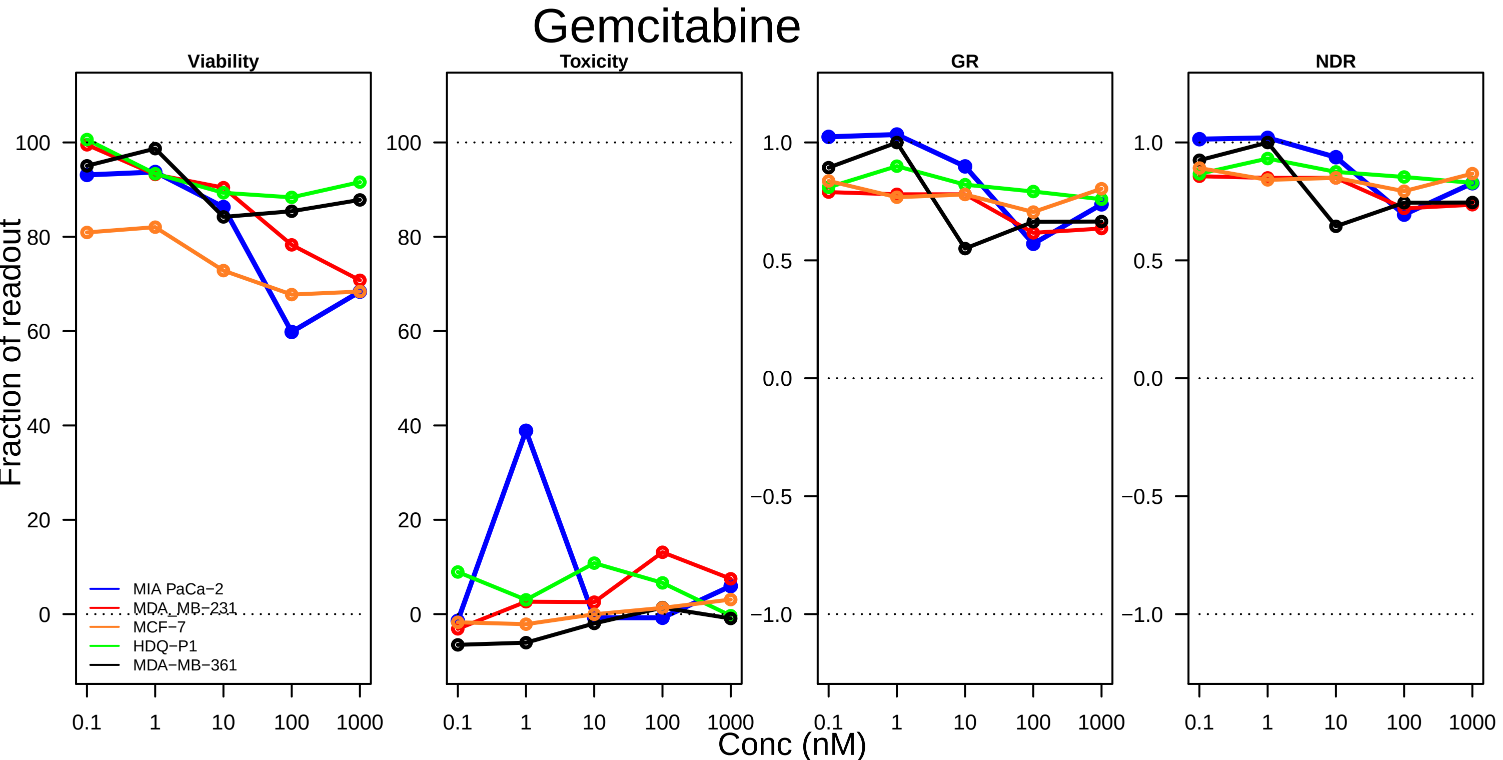
Gemcitabine
Viability
Toxicity
GR
NDR
Fraction of readout
Conc (nM)
MIA PaCa−2
MDA_MB−231
MCF−7
HDQ−P1
MDA−MB−361
0
20
40
60
80
100
1.0
0.5
0.0
−0.5
−1.0
0.1
1
10
100
1000

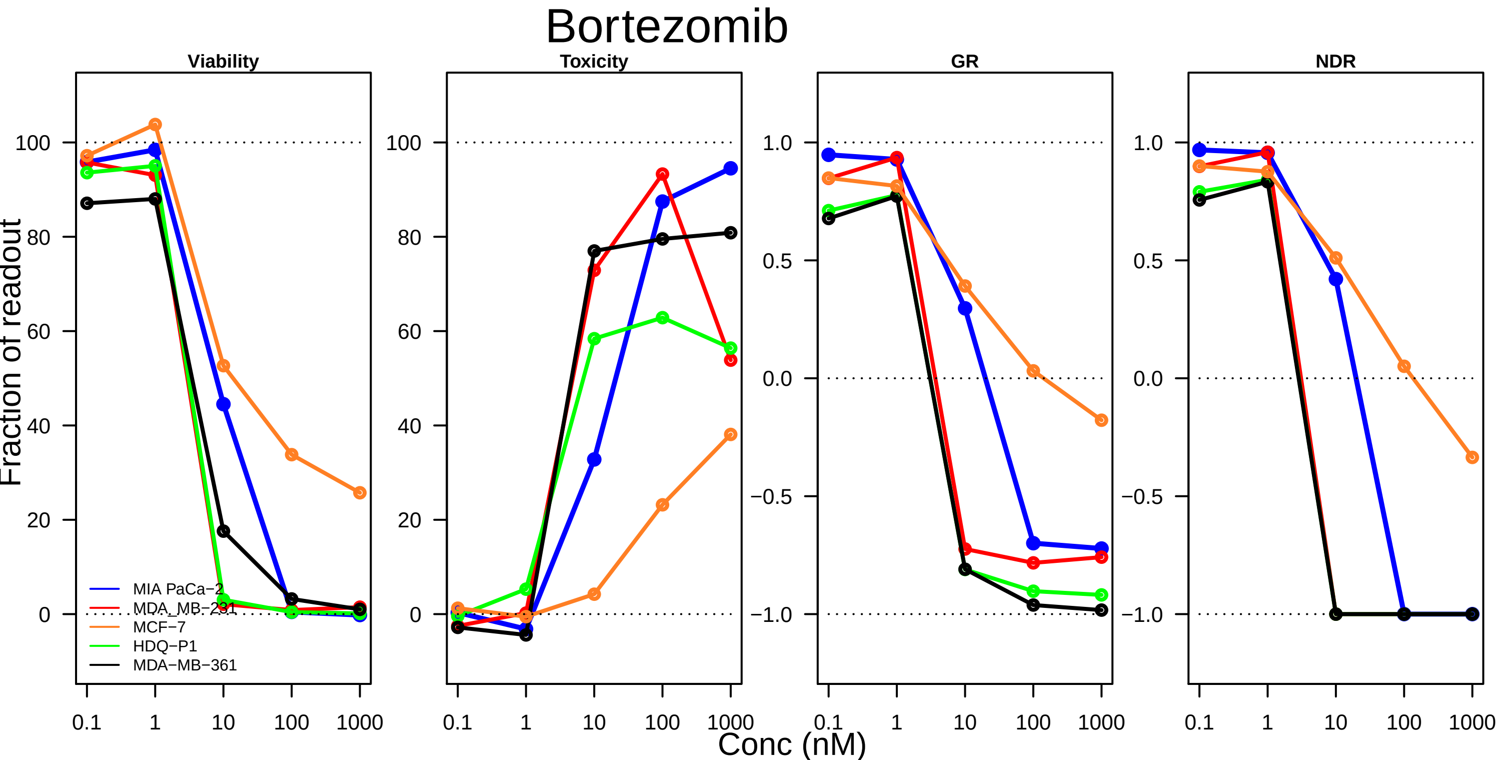

Bortezomib
Viability
Toxicity
GR
NDR
Fraction of readout
Conc (nM)
MIA PaCa−2
MDA_MB−231
MCF−7
HDQ−P1
MDA−MB−361
0.1
1
10
100
1000

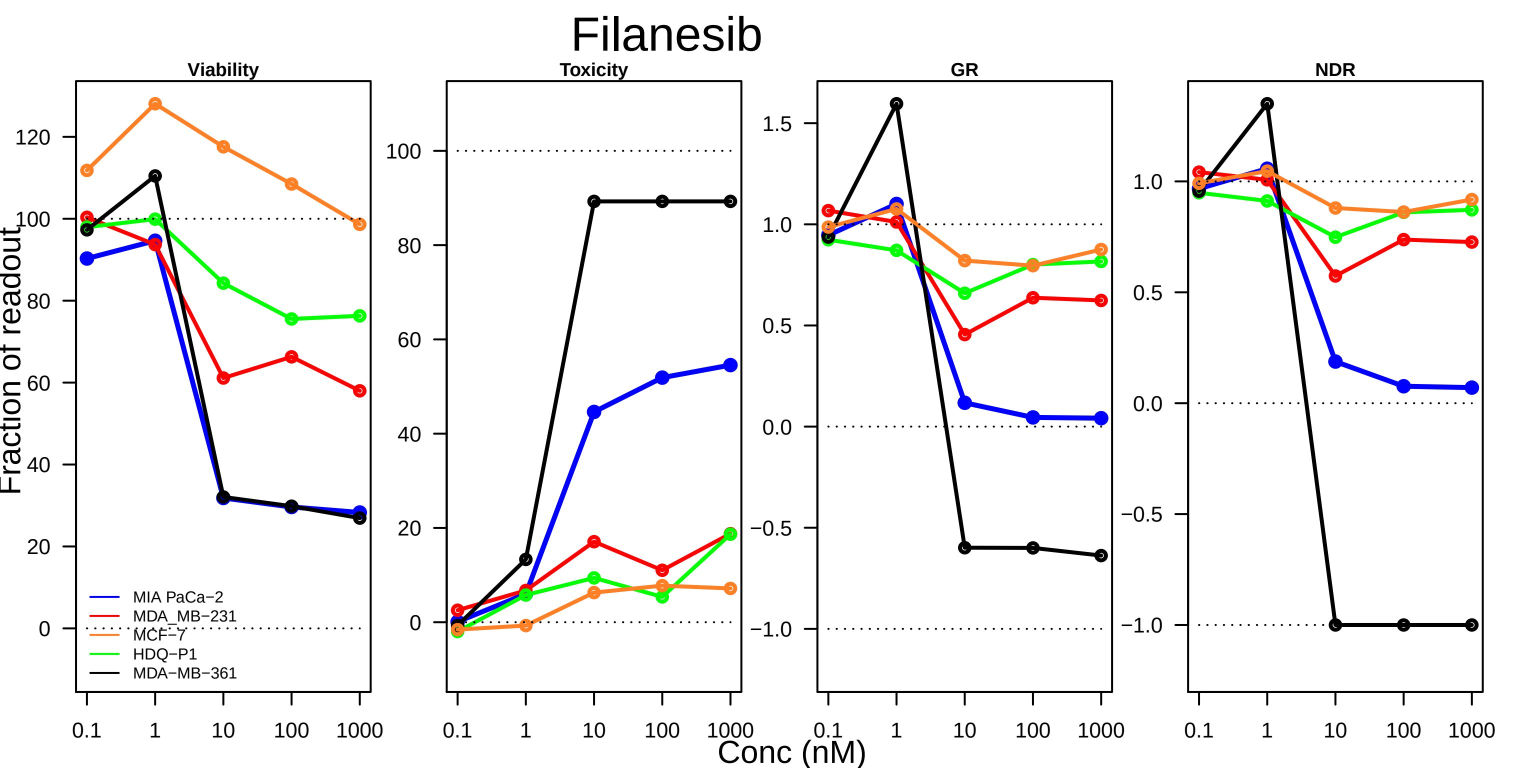

Filanesib
Viability
Toxicity
GR
NDR
Fraction of readout
Conc (nM)
MIA PaCa−2
MDA_MB−231
MCF−7
HDQ−P1
MDA−MB−361
0.1
1
10
100
1000

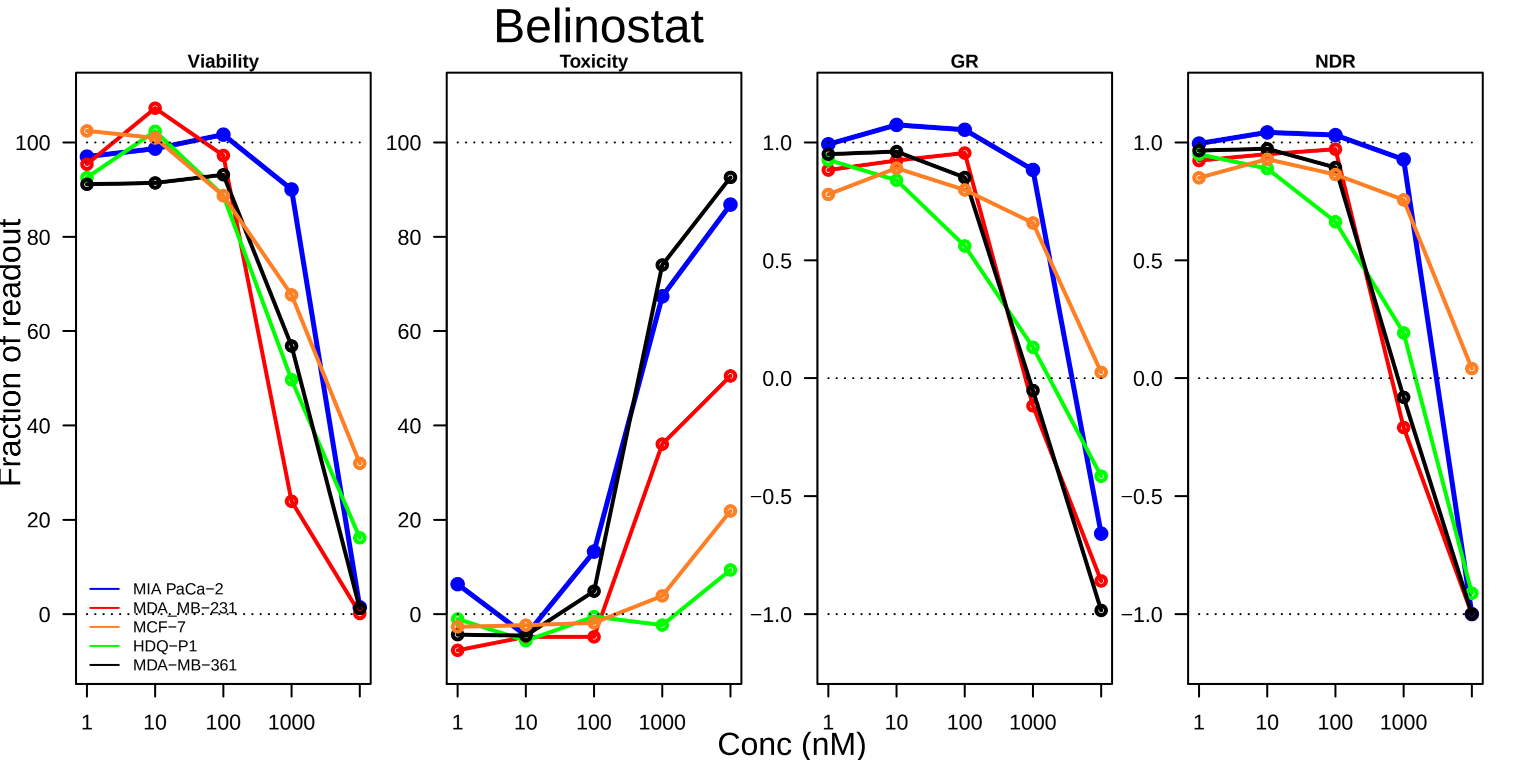
Belinostat
Viability
Toxicity
GR
NDR
Fraction of readout
Conc (nM)
100
80
60
40
20
0
1.0
0.5
0.0
−0.5
−1.0
1
10
100
1000
MIA PaCa−2
MDA_MB−231
MCF−7
HDQ−P1
MDA−MB−361

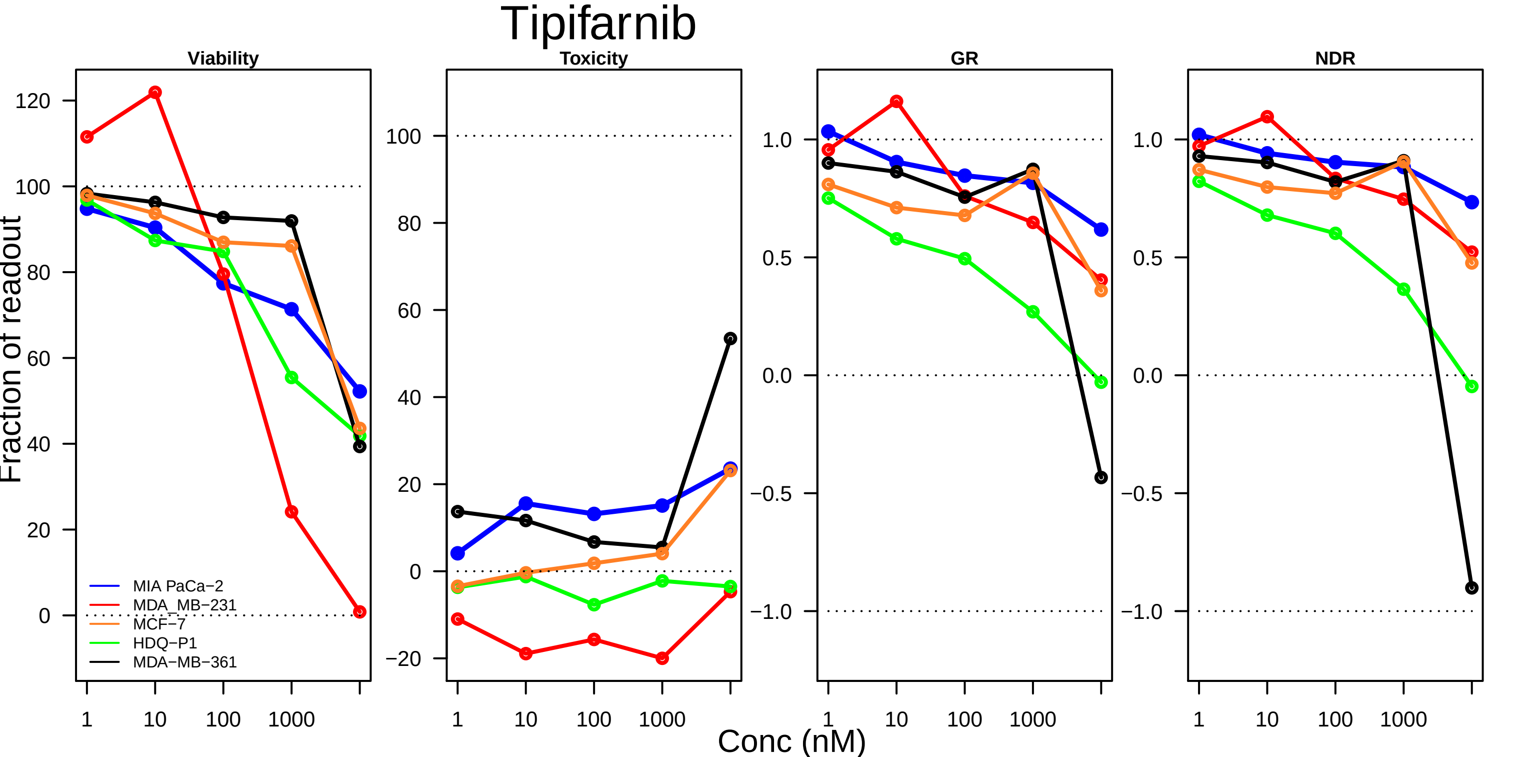

Tipifarnib
Viability
Toxicity
GR
NDR
Fraction of readout
Conc (nM)
MIA PaCa−2
MDA_MB−231
MCF−7
HDQ−P1
MDA−MB−361
1
10
100
1000

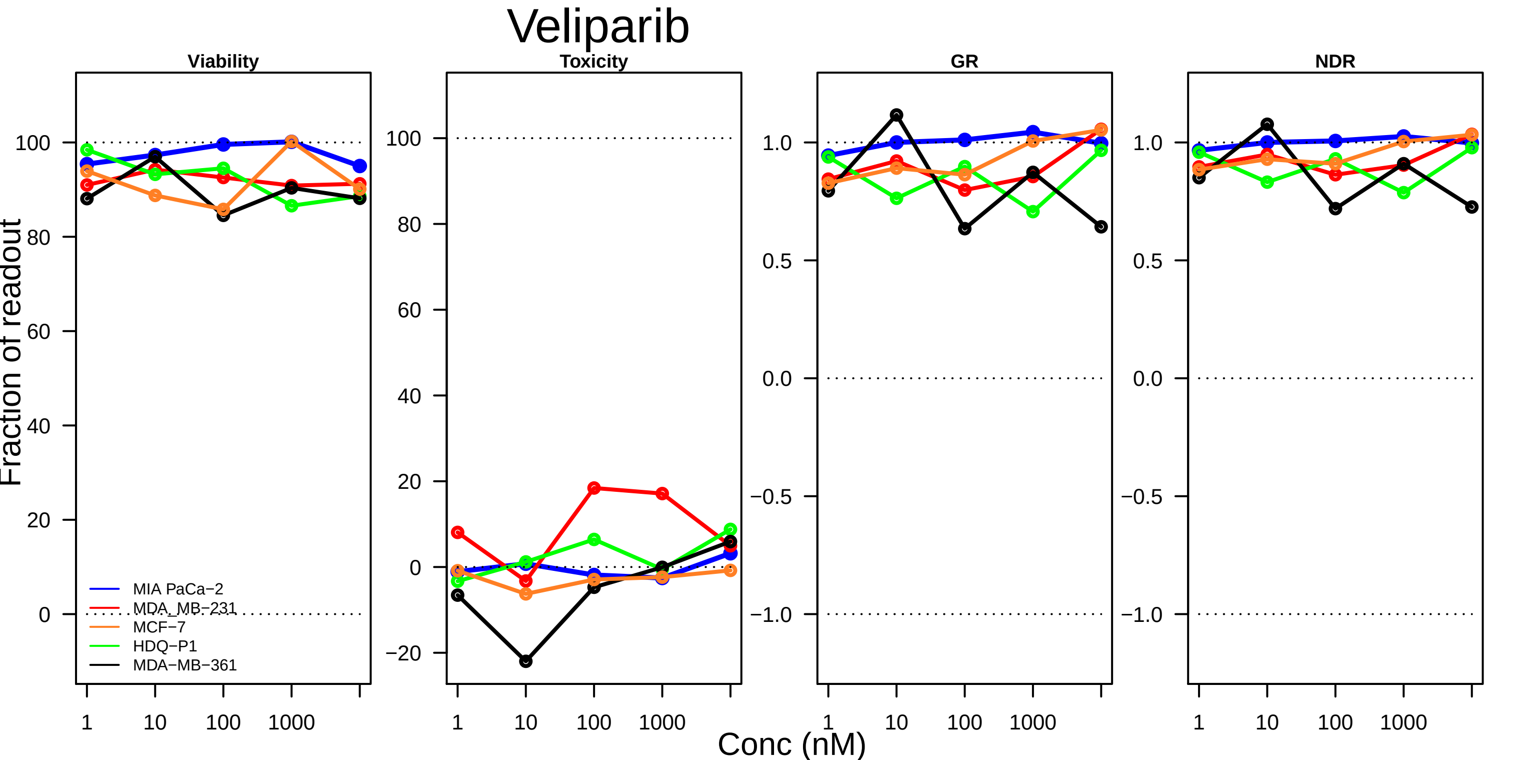

Veliparib
Viability
Toxicity
GR
NDR
Fraction of readout
Conc (nM)
1
10
100
1000
100
80
60
40
20
0
−20
1.0
0.5
0.0
−0.5
−1.0
MIA PaCa−2
MDA_MB−231
MCF−7
HDQ−P1
MDA−MB−361

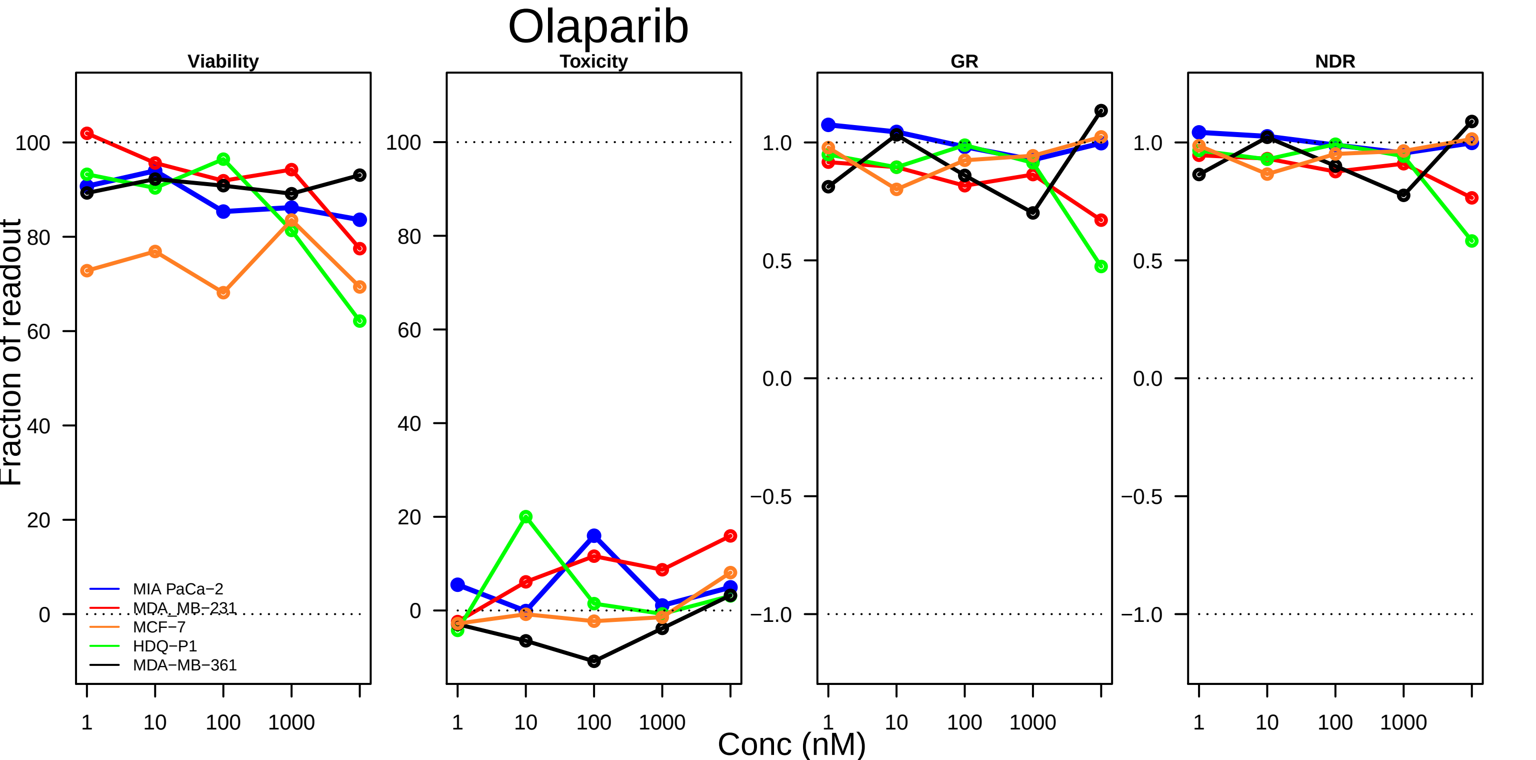
Olaparib
Viability
Toxicity
GR
NDR
Fraction of readout
MIA PaCa−2
MDA_MB−231
MCF−7
HDQ−P1
MDA−MB−361
100
80
60
40
20
0
1.0
0.5
0.0
−0.5
−1.0
1
10
100
1000
Conc (nM)

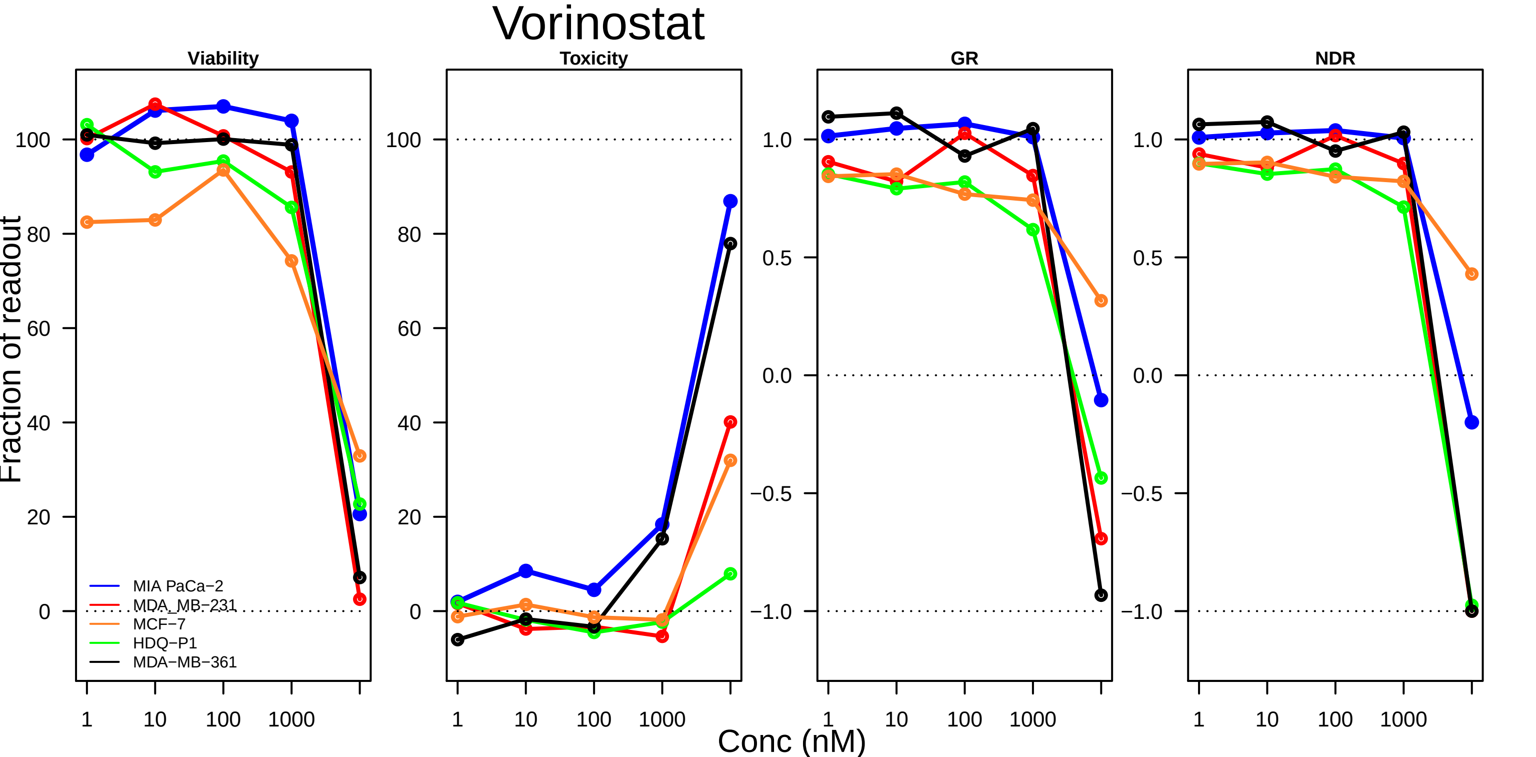

Vorinostat
Viability
Toxicity
GR
NDR
Fraction of readout
100
80
60
40
20
0
1.0
0.5
0.0
−0.5
−1.0
1
10
100
1000
MIA PaCa−2
MDA_MB−231
MCF−7
HDQ−P1
MDA−MB−361
Conc (nM)

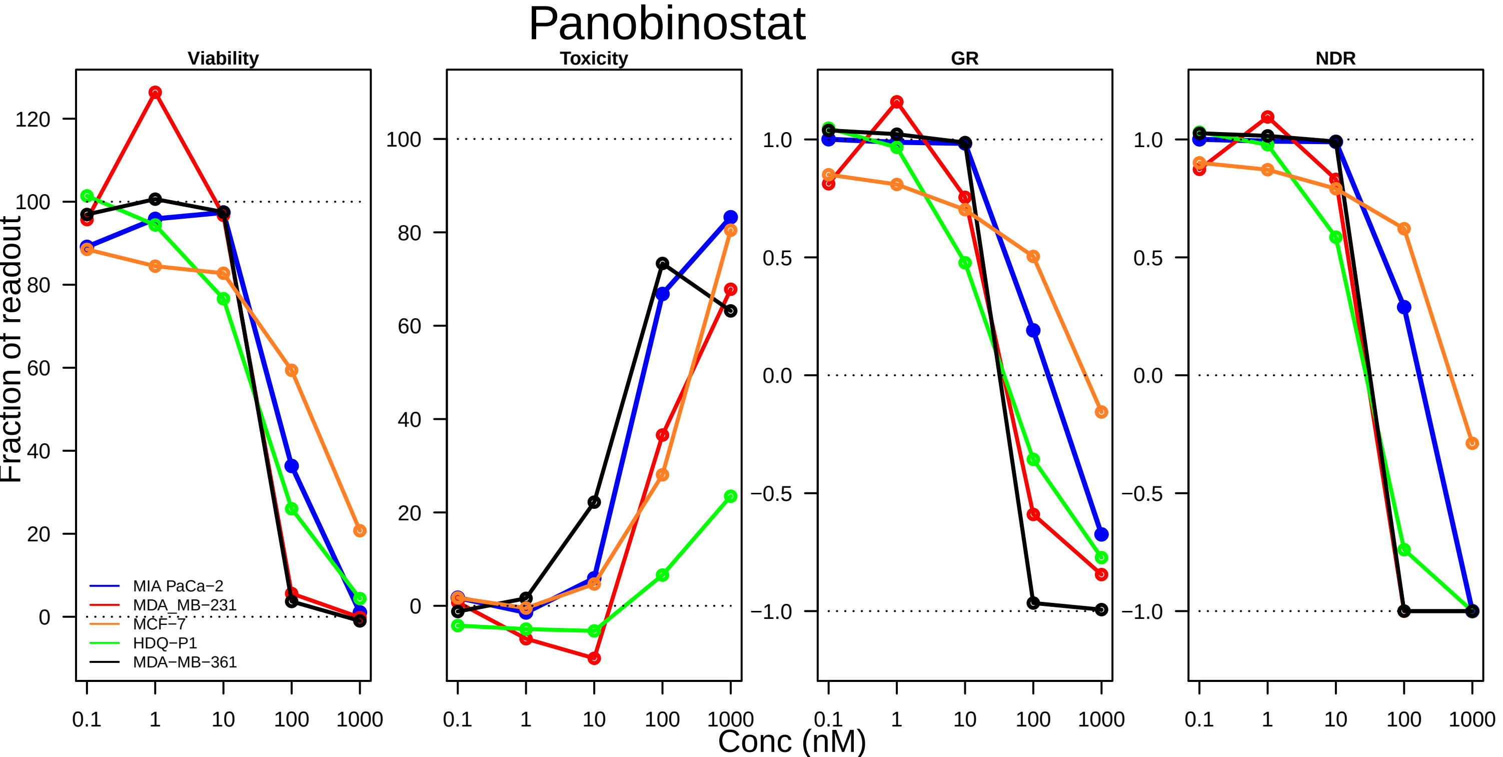
Panobinostat
Viability
Toxicity
GR
NDR
Fraction of readout
Conc (nM)
MIA PaCa−2
MDA_MB−231
MCF−7
HDQ−P1
MDA−MB−361
0.1
1
10
100
1000

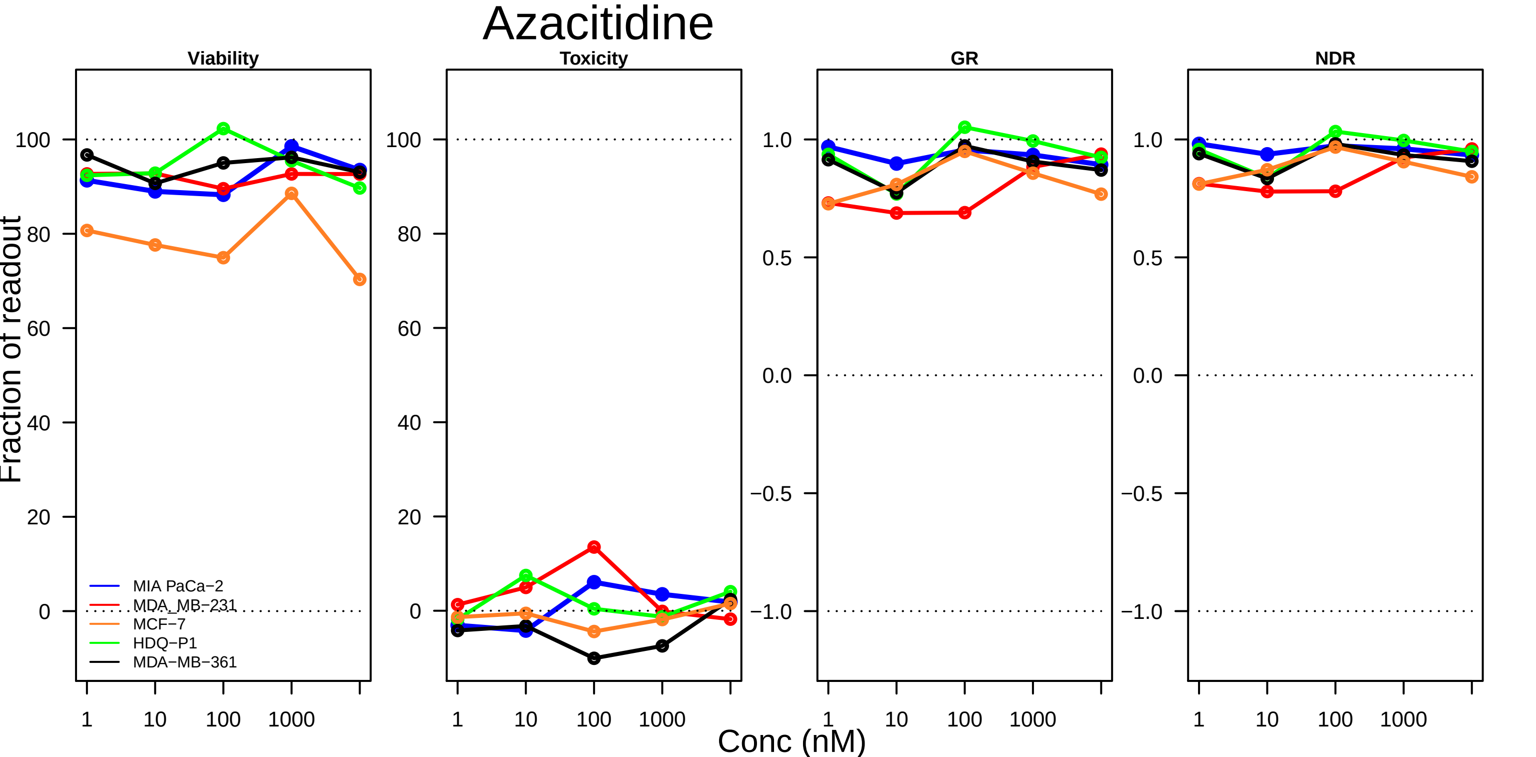

Azacitidine
Viability
Toxicity
GR
NDR
Fraction of readout
Conc (nM)
MIA PaCa−2
MDA_MB−231
MCF−7
HDQ−P1
MDA−MB−361
100
80
60
40
20
0
1.0
0.5
0.0
−0.5
−1.0
1
10
100
1000

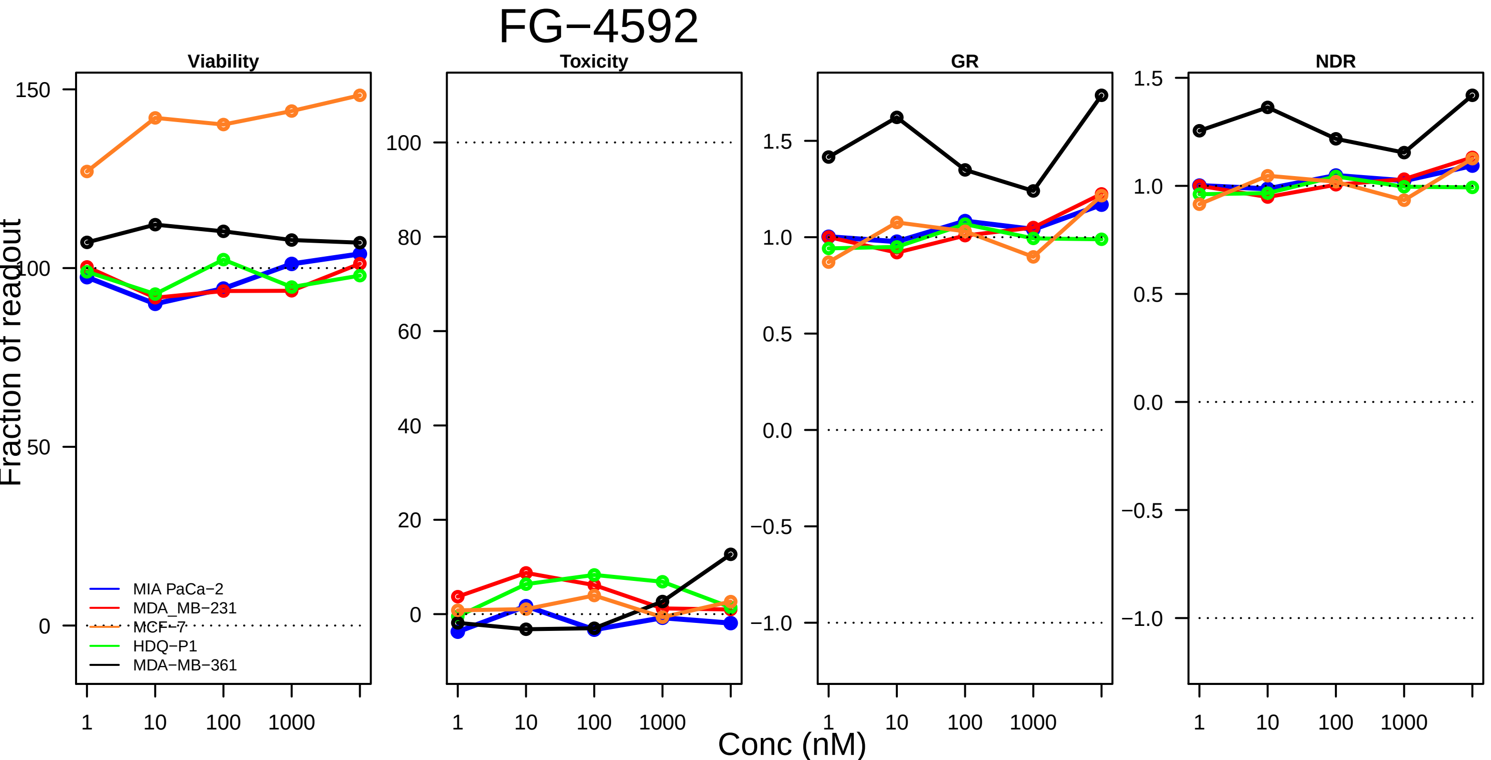
FG−4592
Viability
Toxicity
GR
NDR
Fraction of readout
Conc (nM)
MIA PaCa−2
MDA_MB−231
MCF−7
HDQ−P1
MDA−MB−361

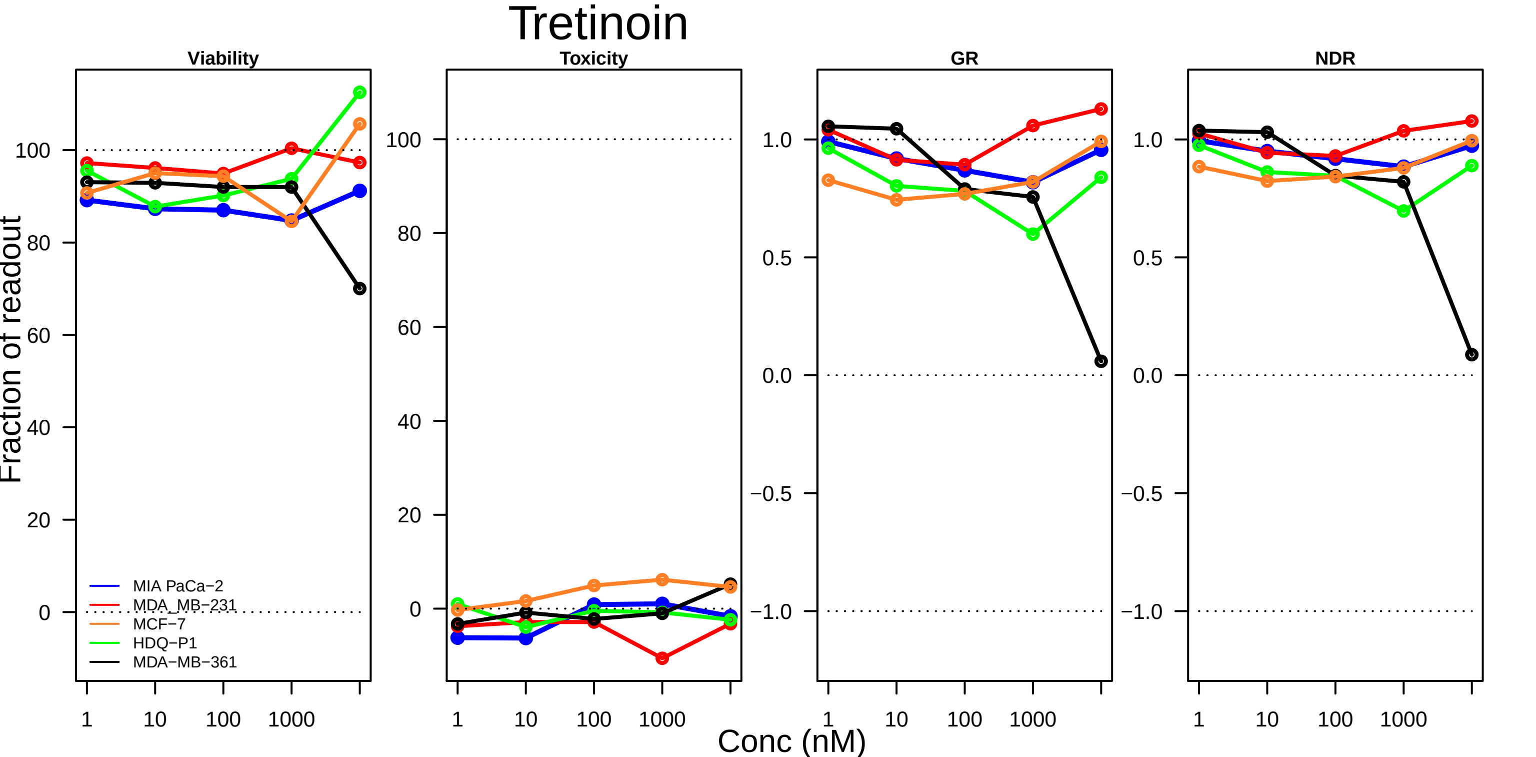

Tretinoin
Viability
Toxicity
GR
NDR
Fraction of readout
Conc (nM)
1
10
100
1000
MIA PaCa−2
MDA_MB−231
MCF−7
HDQ−P1
MDA−MB−361

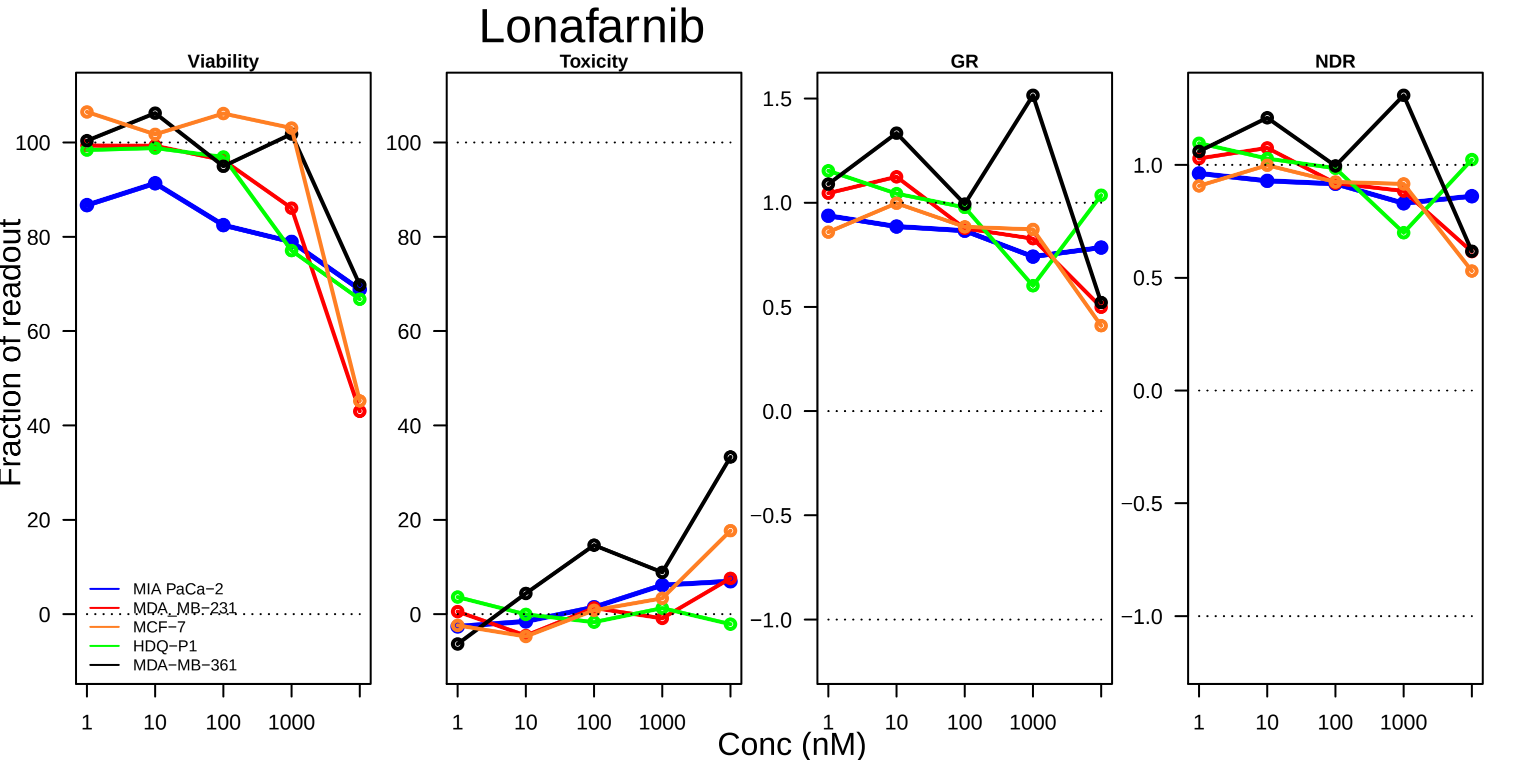

Lonafarnib
Viability
Toxicity
GR
NDR
Fraction of readout
Conc (nM)
1
10
100
1000
0
20
40
60
80
100
1.5
1.0
0.5
0.0
−0.5
−1.0
MIA PaCa−2
MDA_MB−231
MCF−7
HDQ−P1
MDA−MB−361

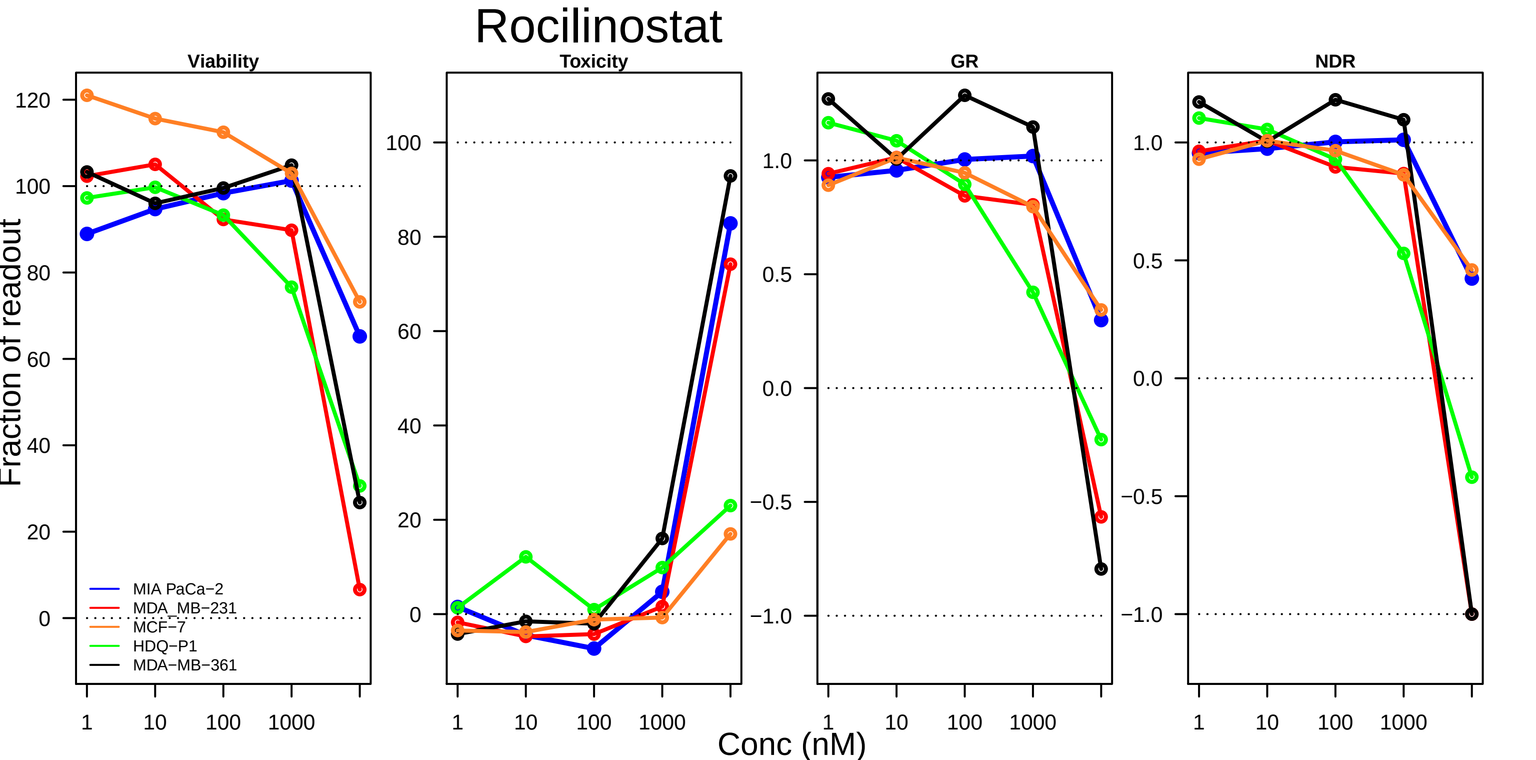
Rocilinostat
Viability
Toxicity
GR
NDR
Fraction of readout
MIA PaCa−2
MDA_MB−231
MCF−7
HDQ−P1
MDA−MB−361
Conc (nM)

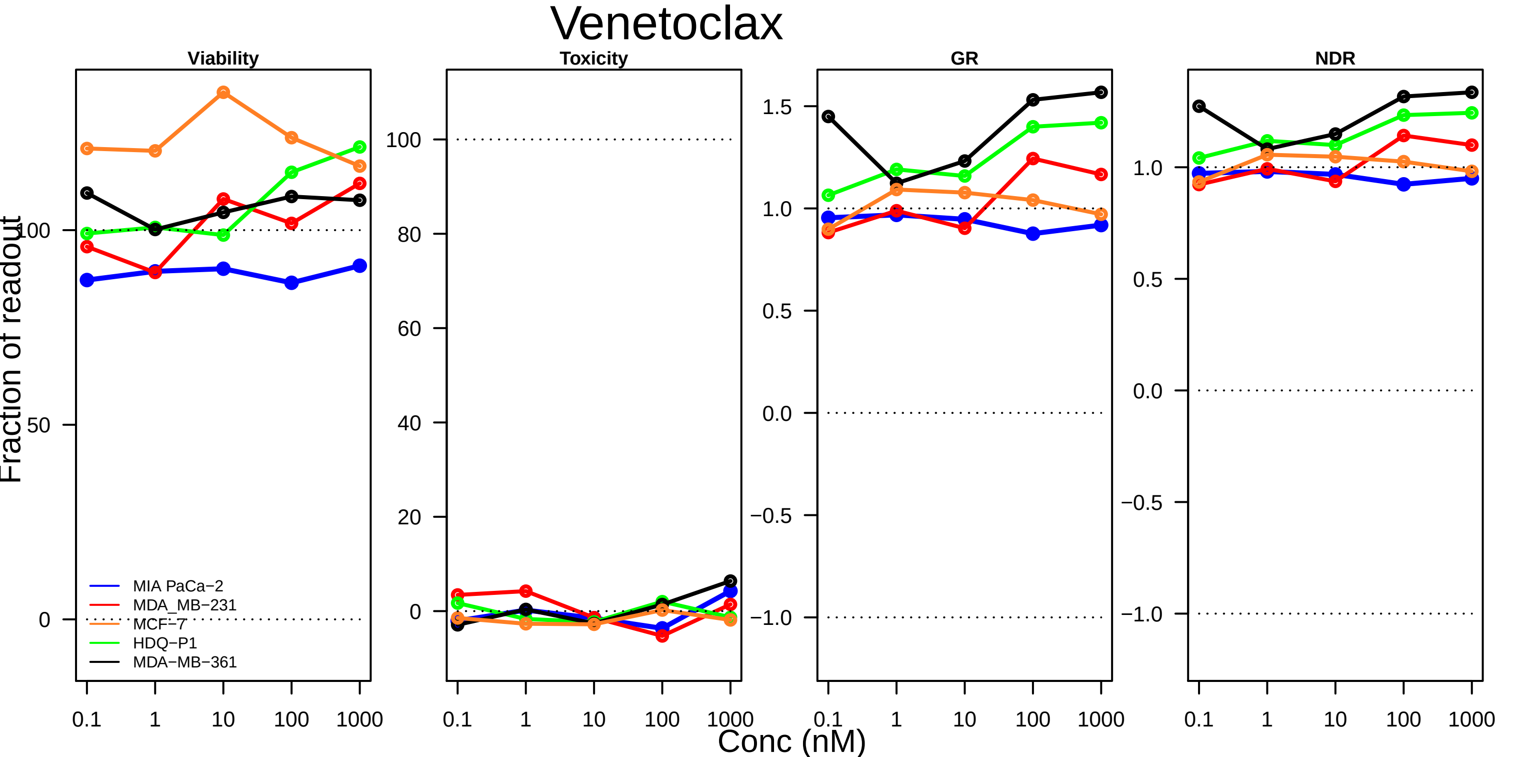

Venetoclax
Viability
Toxicity
GR
NDR
Fraction of readout
Conc (nM)
MIA PaCa−2
MDA_MB−231
MCF−7
HDQ−P1
MDA−MB−361
0.1
1
10
100
1000
0
50
100
20
40
60
80
−1.0
−0.5
0.0
0.5
1.0
1.5

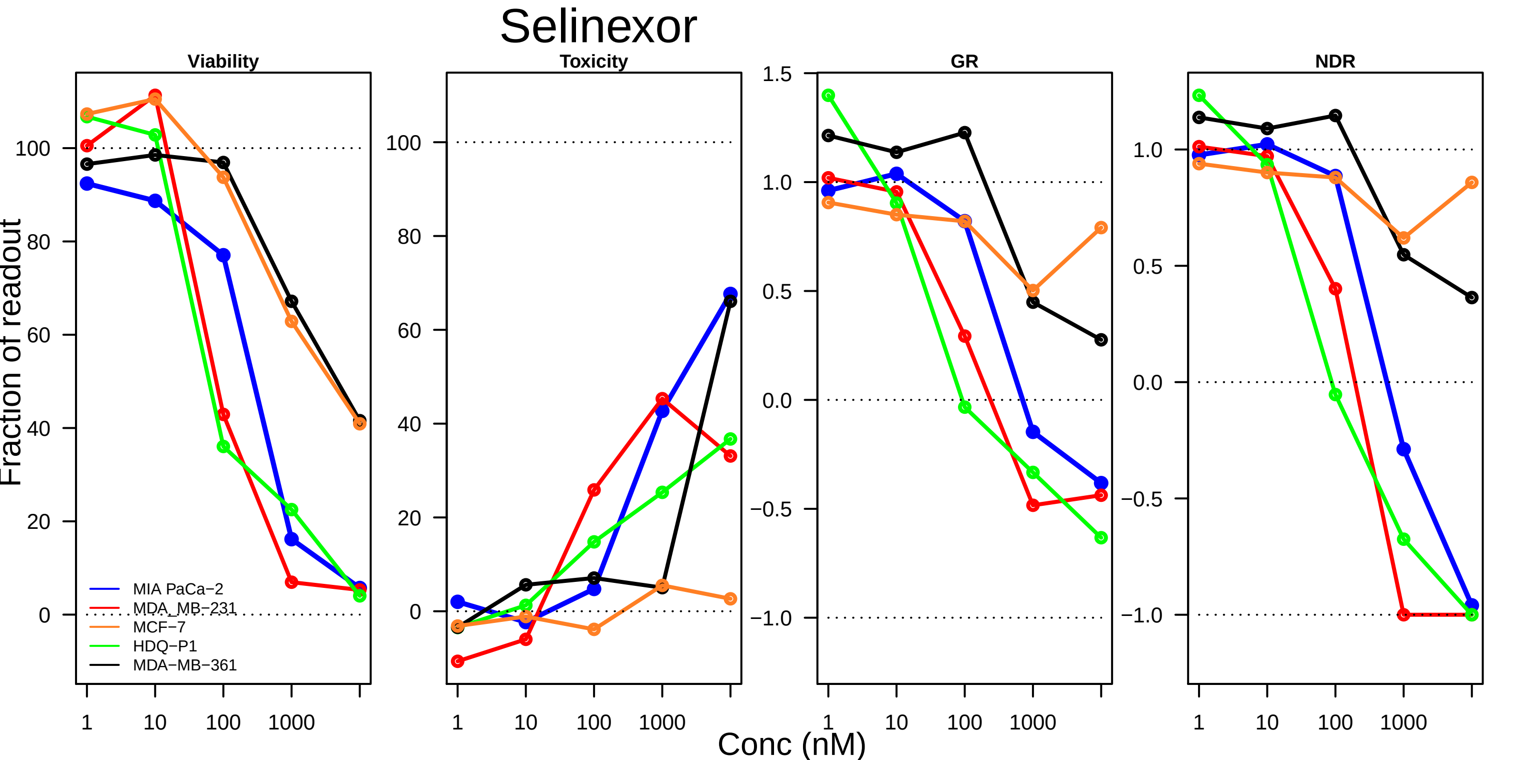

Selinexor
Viability
Toxicity
GR
NDR
Fraction of readout
Conc (nM)
1
10
100
1000
0
20
40
60
80
100
1.5
1.0
0.5
0.0
−0.5
−1.0
MIA PaCa−2
MDA_MB−231
MCF−7
HDQ−P1
MDA−MB−361

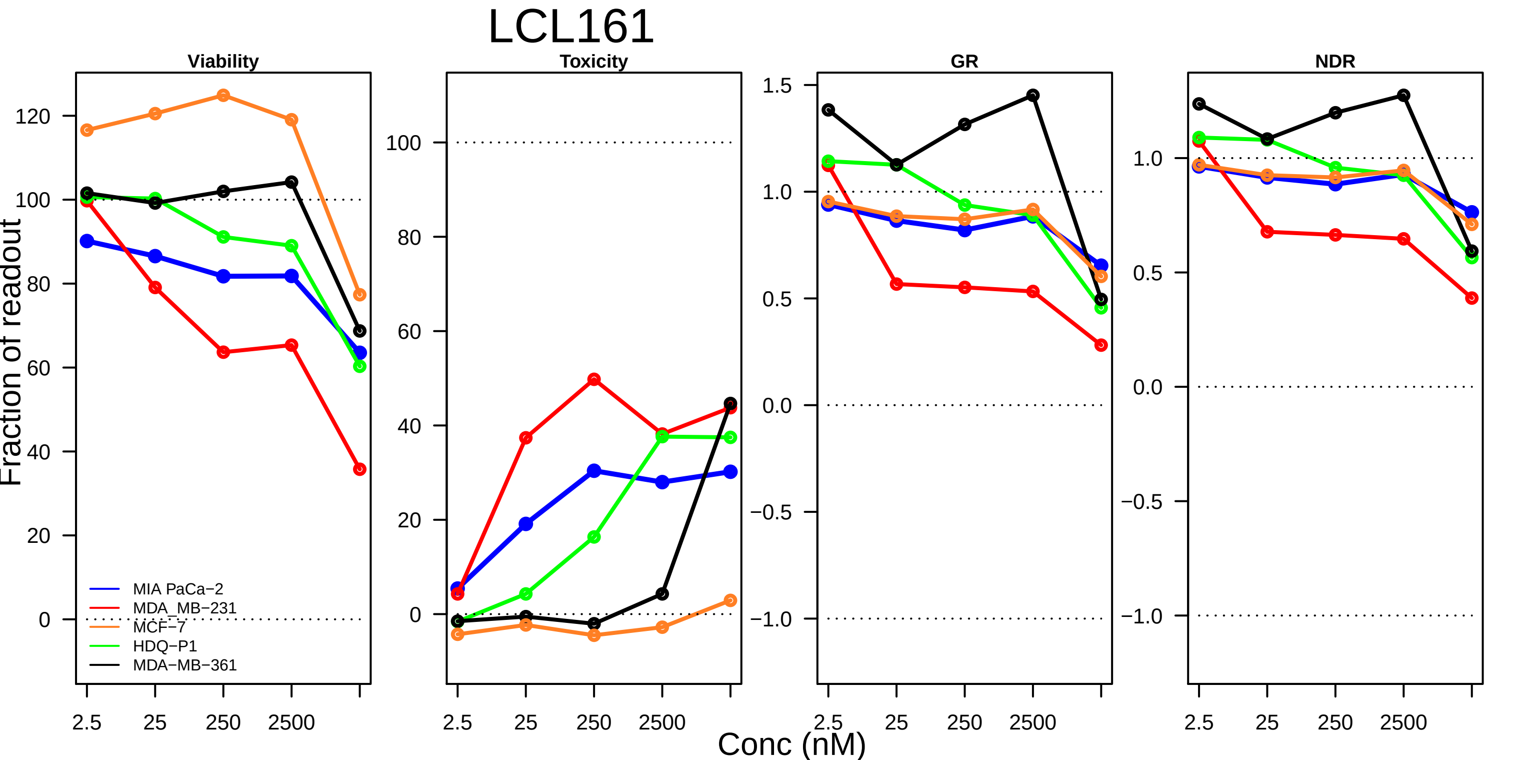
LCL161
Viability
Toxicity
GR
NDR
Fraction of readout
MIA PaCa−2
MDA_MB−231
MCF−7
HDQ−P1
MDA−MB−361
Conc (nM)

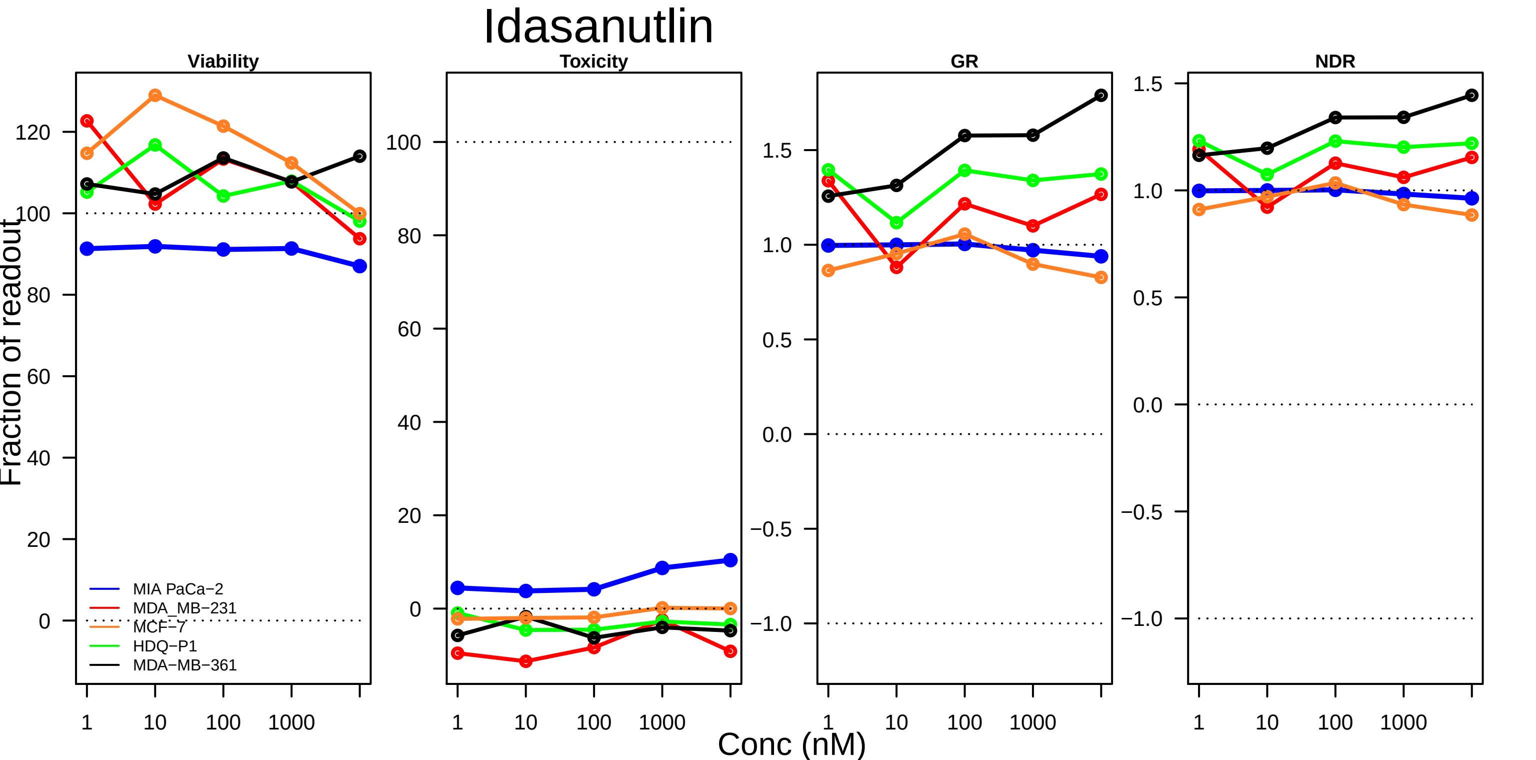
Idasanutlin
Viability
Toxicity
GR
NDR
Fraction of readout
Conc (nM)
MIA PaCa−2
MDA_MB−231
MCF−7
HDQ−P1
MDA−MB−361
1
10
100
1000

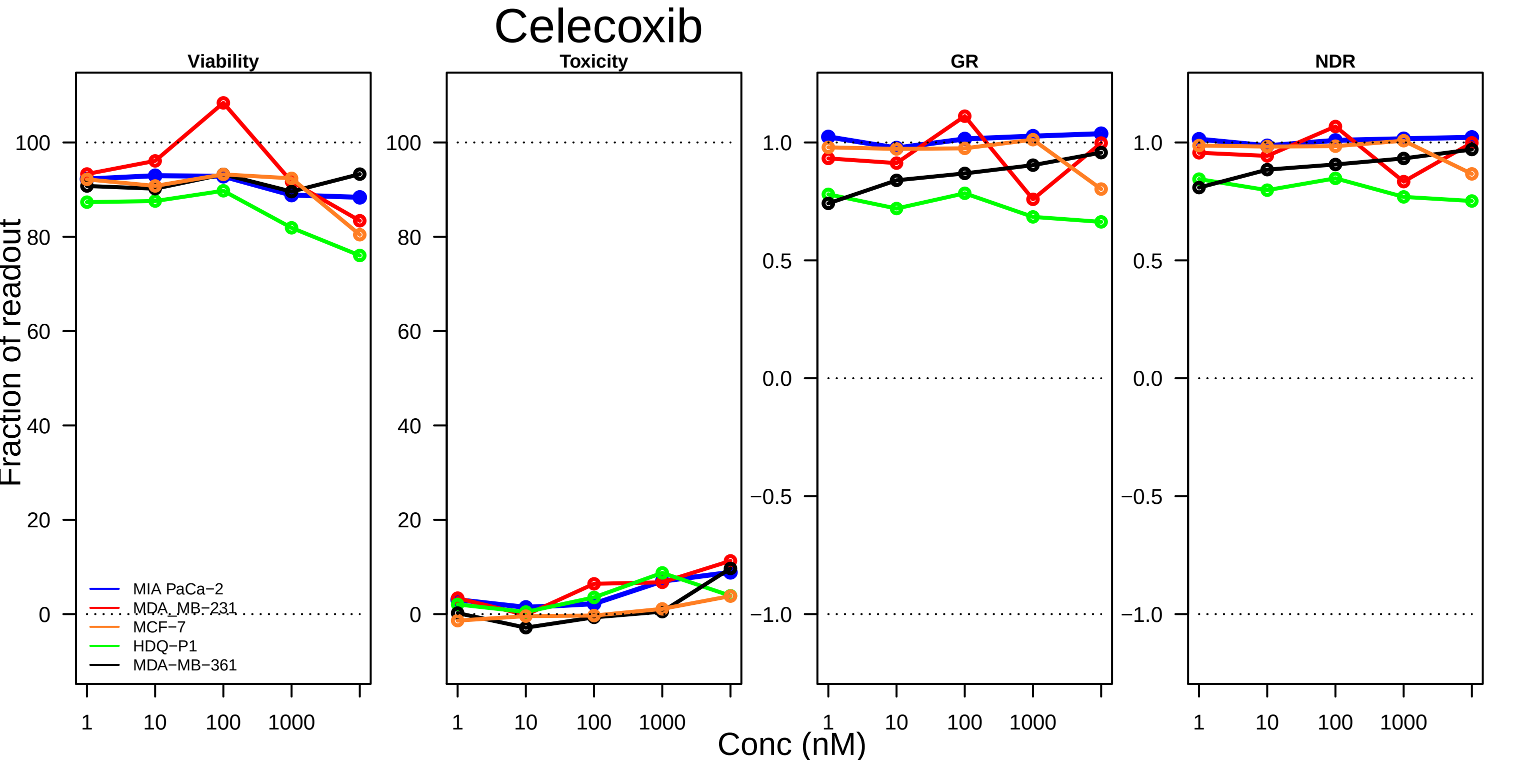

Celecoxib
Viability
Toxicity
GR
NDR
Fraction of readout
Conc (nM)
MIA PaCa−2
MDA_MB−231
MCF−7
HDQ−P1
MDA−MB−361

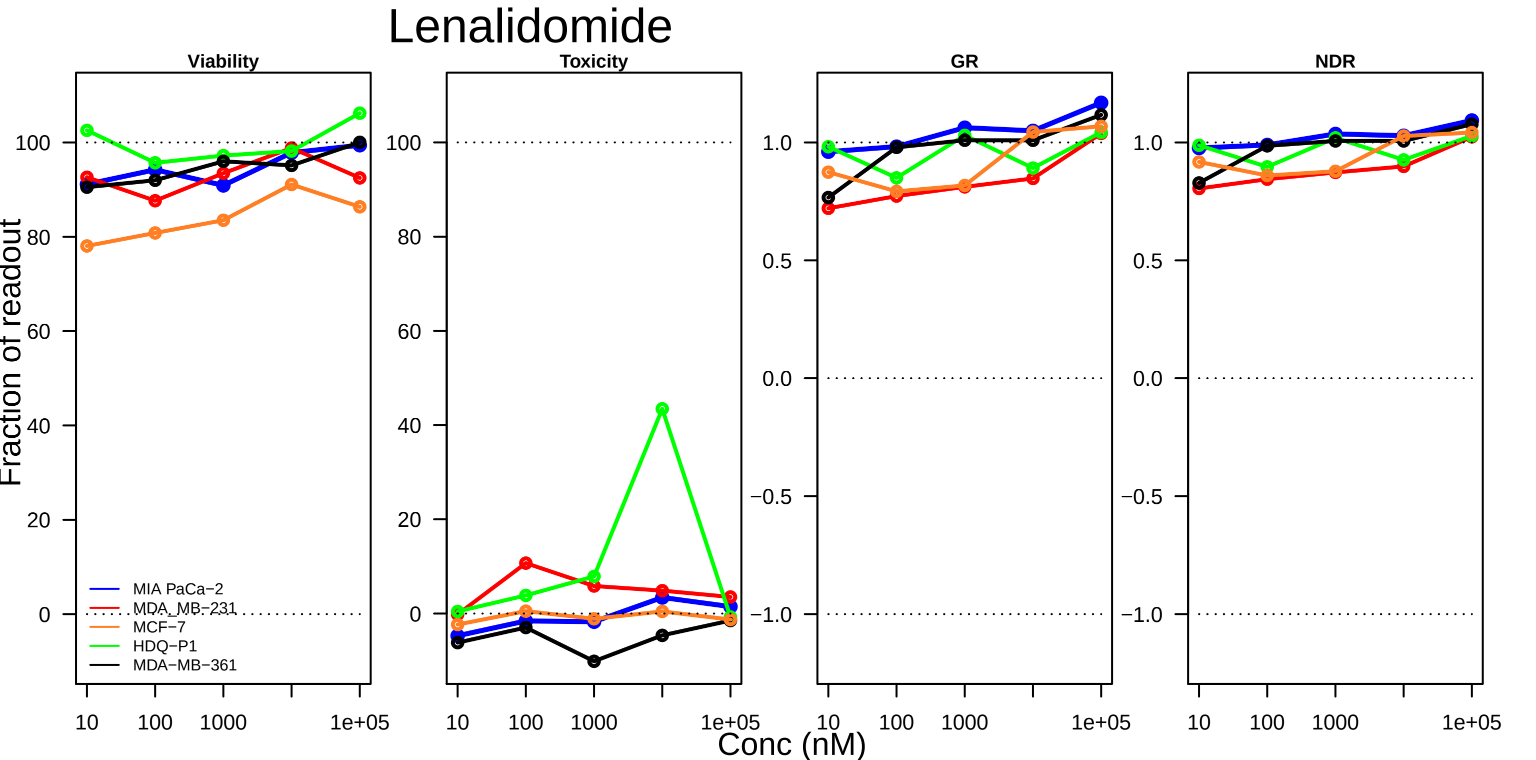

Lenalidomide
Viability
Toxicity
GR
NDR
Fraction of readout
Conc (nM)
MIA PaCa−2
MDA_MB−231
MCF−7
HDQ−P1
MDA−MB−361
0
20
40
60
80
100
−1.0
−0.5
0.0
0.5
1.0
10
100
1000
1e+05

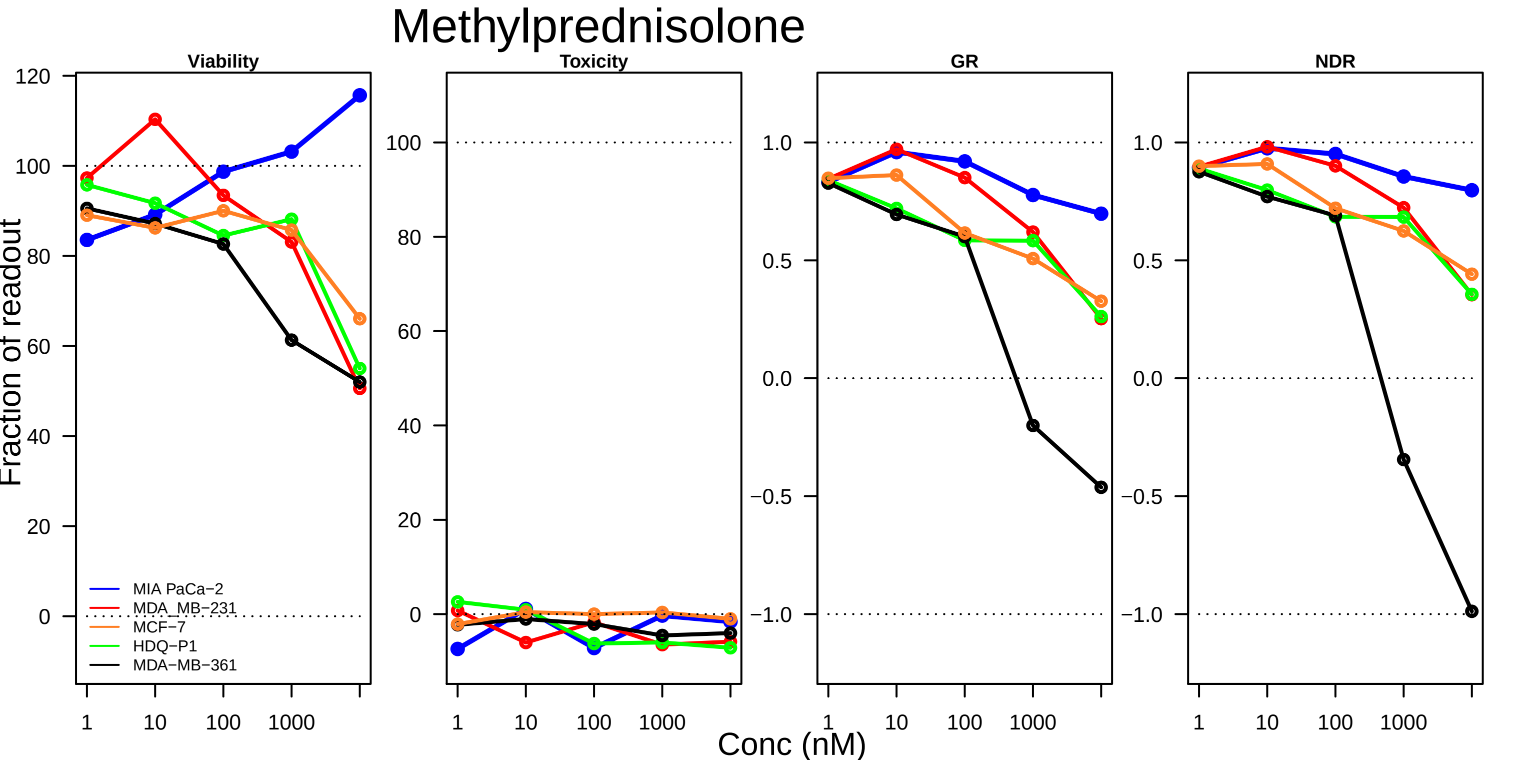

Methylprednisolone
Viability
Toxicity
GR
NDR
Fraction of readout
Conc (nM)
MIA PaCa−2
MDA_MB−231
MCF−7
HDQ−P1
MDA−MB−361
120
100
80
60
40
20
0
1.0
0.5
0.0
−0.5
−1.0
1
10
100
1000

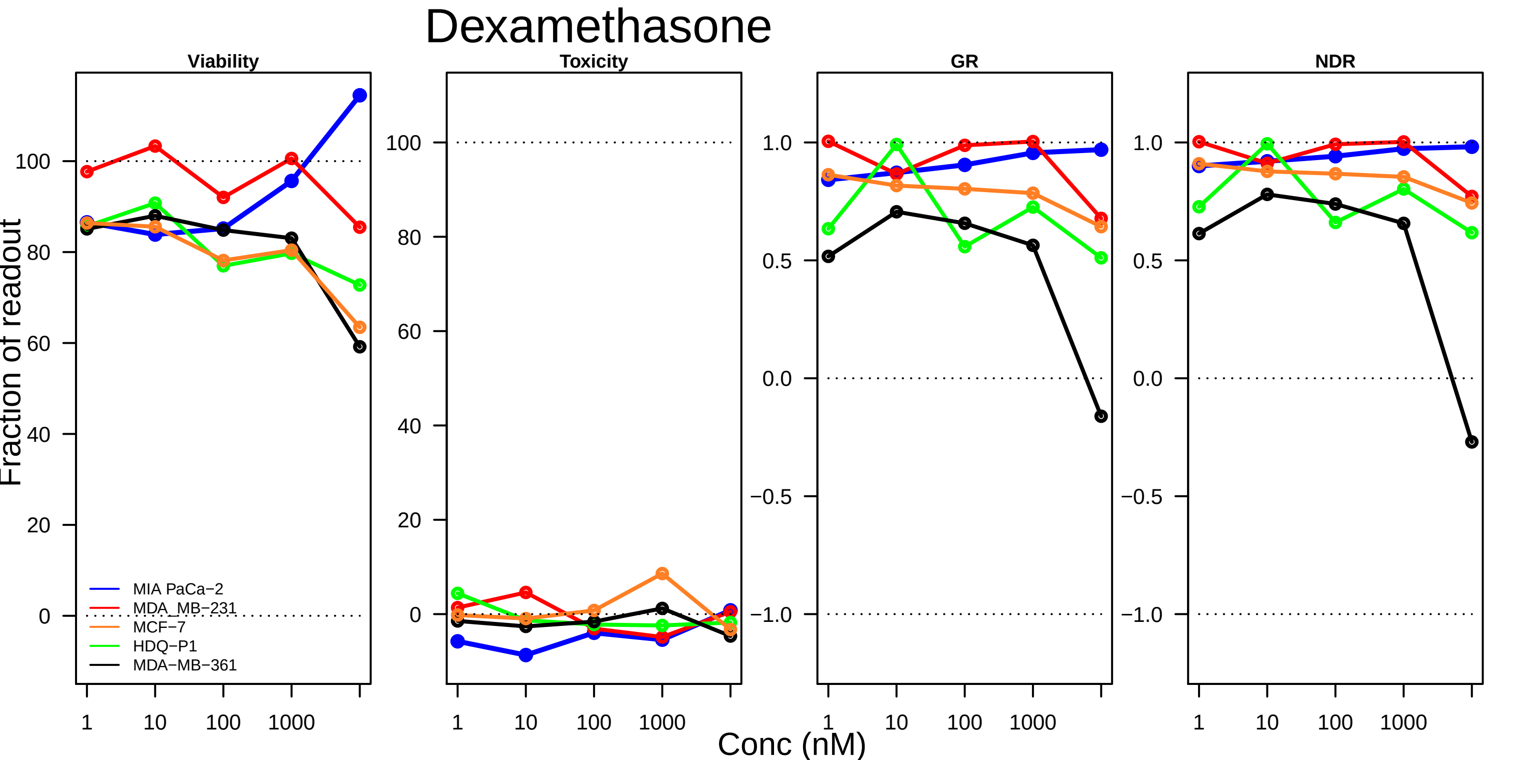
Dexamethasone
Viability
Toxicity
GR
NDR
Fraction of readout
Conc (nM)
1
10
100
1000
100
80
60
40
20
0
1.0
0.5
0.0
−0.5
−1.0
MIA PaCa−2
MDA_MB−231
MCF−7
HDQ−P1
MDA−MB−361

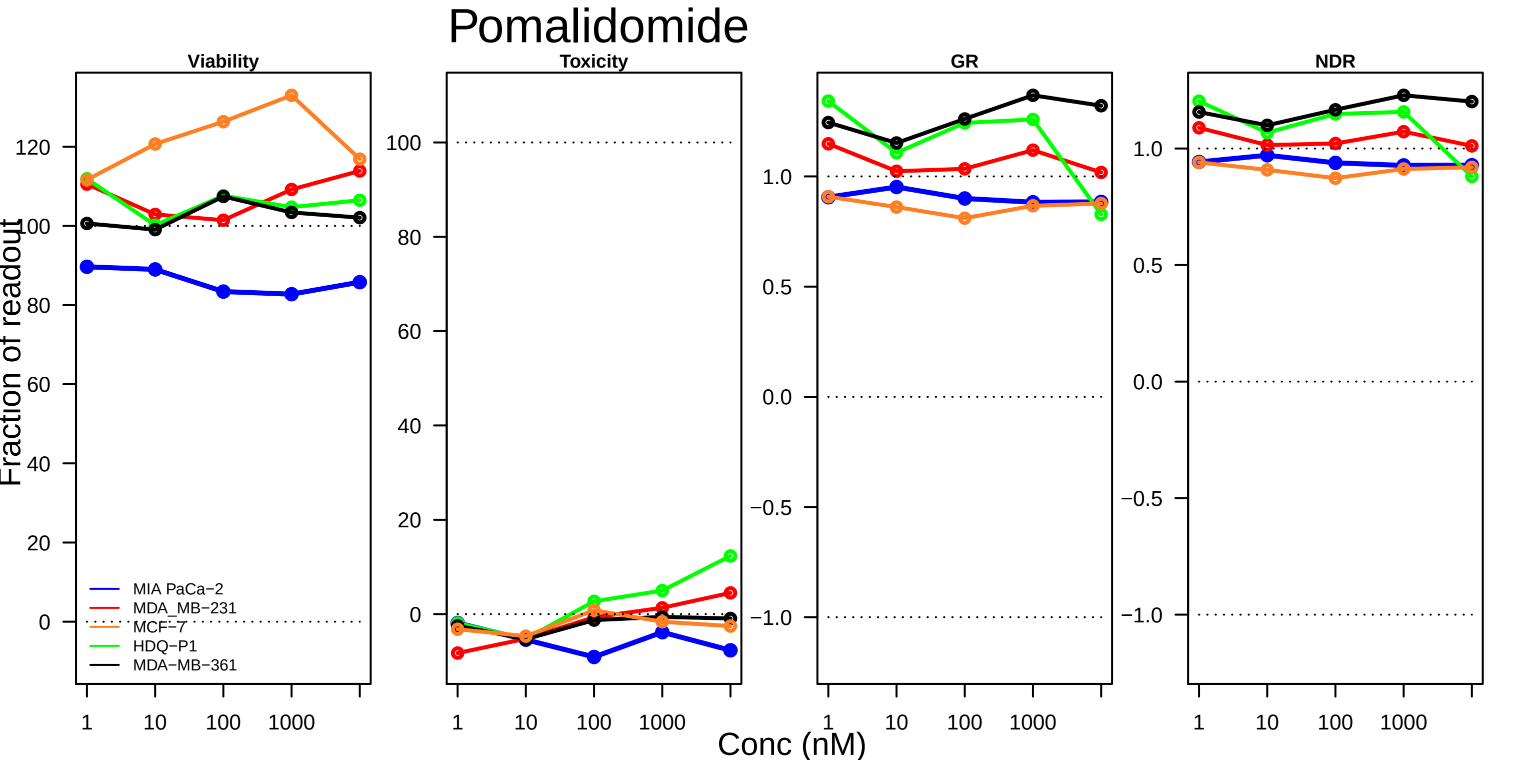
Pomalidomide
Viability
Toxicity
GR
NDR
Fraction of readout
Conc (nM)
1
10
100
1000
MIA PaCa−2
MDA_MB−231
MCF−7
HDQ−P1
MDA−MB−361
120
100
80
60
40
20
0
1.0
0.5
0.0
−0.5
−1.0

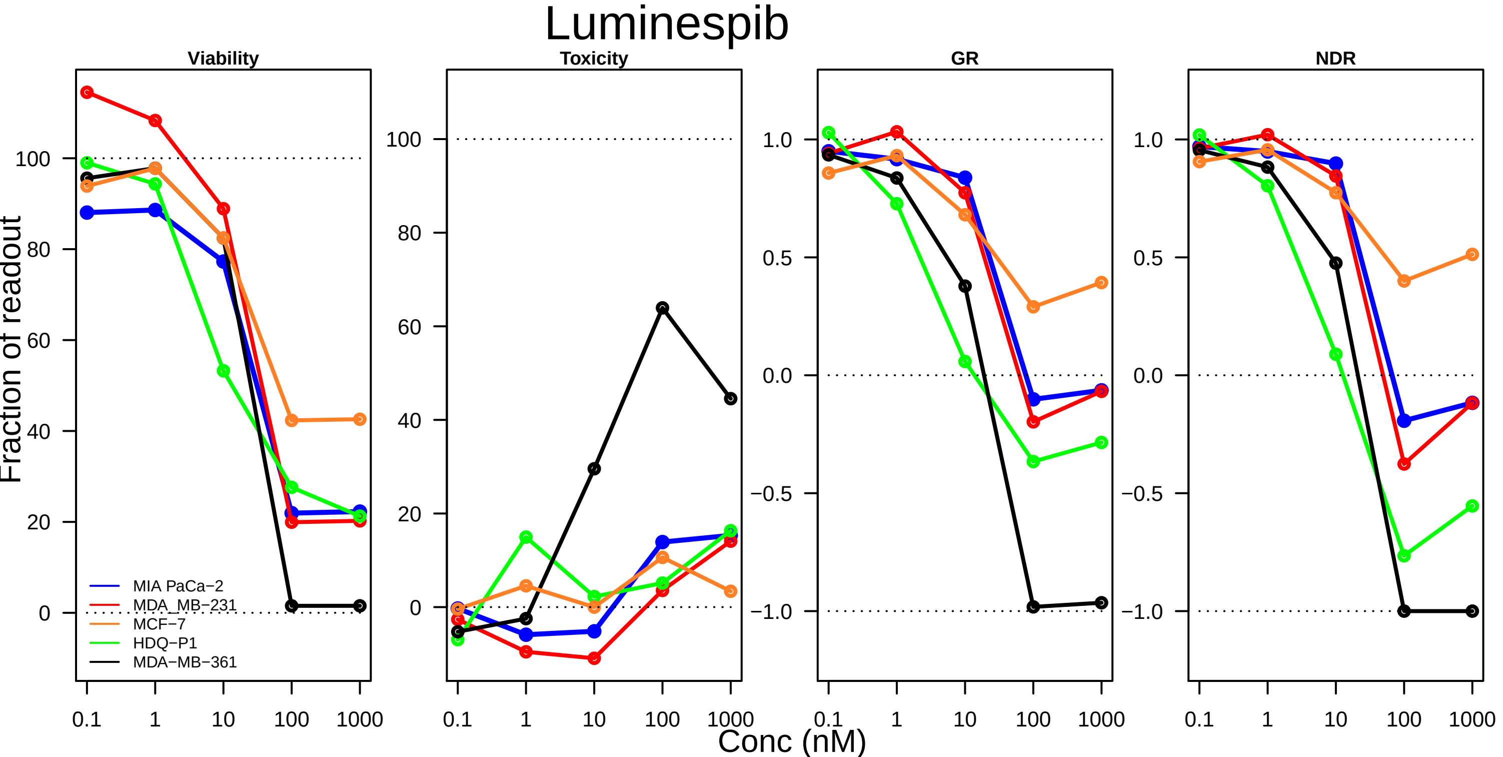

Luminespib
Viability
Toxicity
GR
NDR
Fraction of readout
MIA PaCa−2
MDA_MB−231
MCF−7
HDQ−P1
MDA−MB−361
Conc (nM)

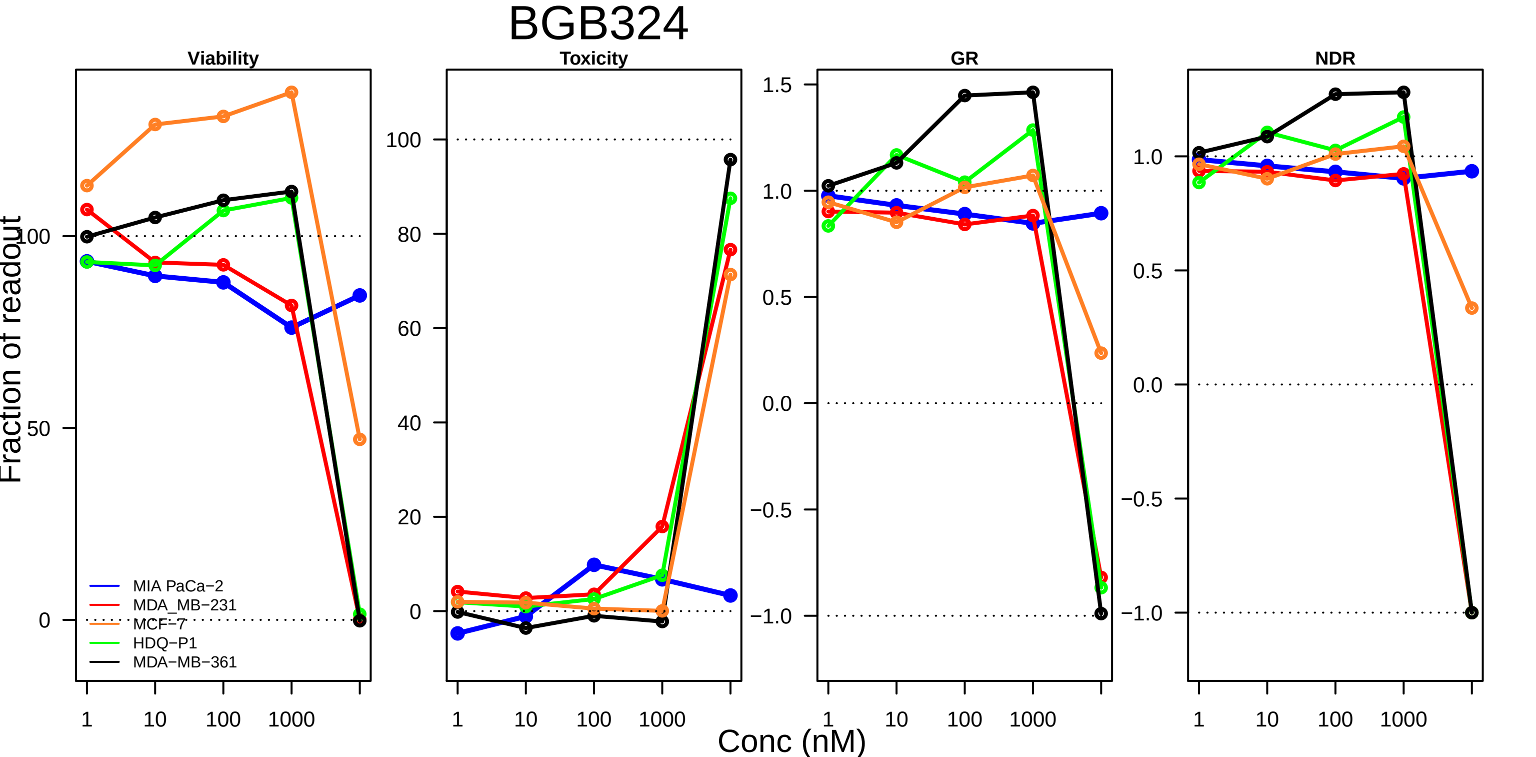
BGB324
Viability
Toxicity
GR
NDR
Fraction of readout
MIA PaCa−2
MDA_MB−231
MCF−7
HDQ−P1
MDA−MB−361
Conc (nM)
1
10
100
1000
0
50
100
20
40
60
80
−1.0
−0.5
0.0
0.5
1.0
1.5

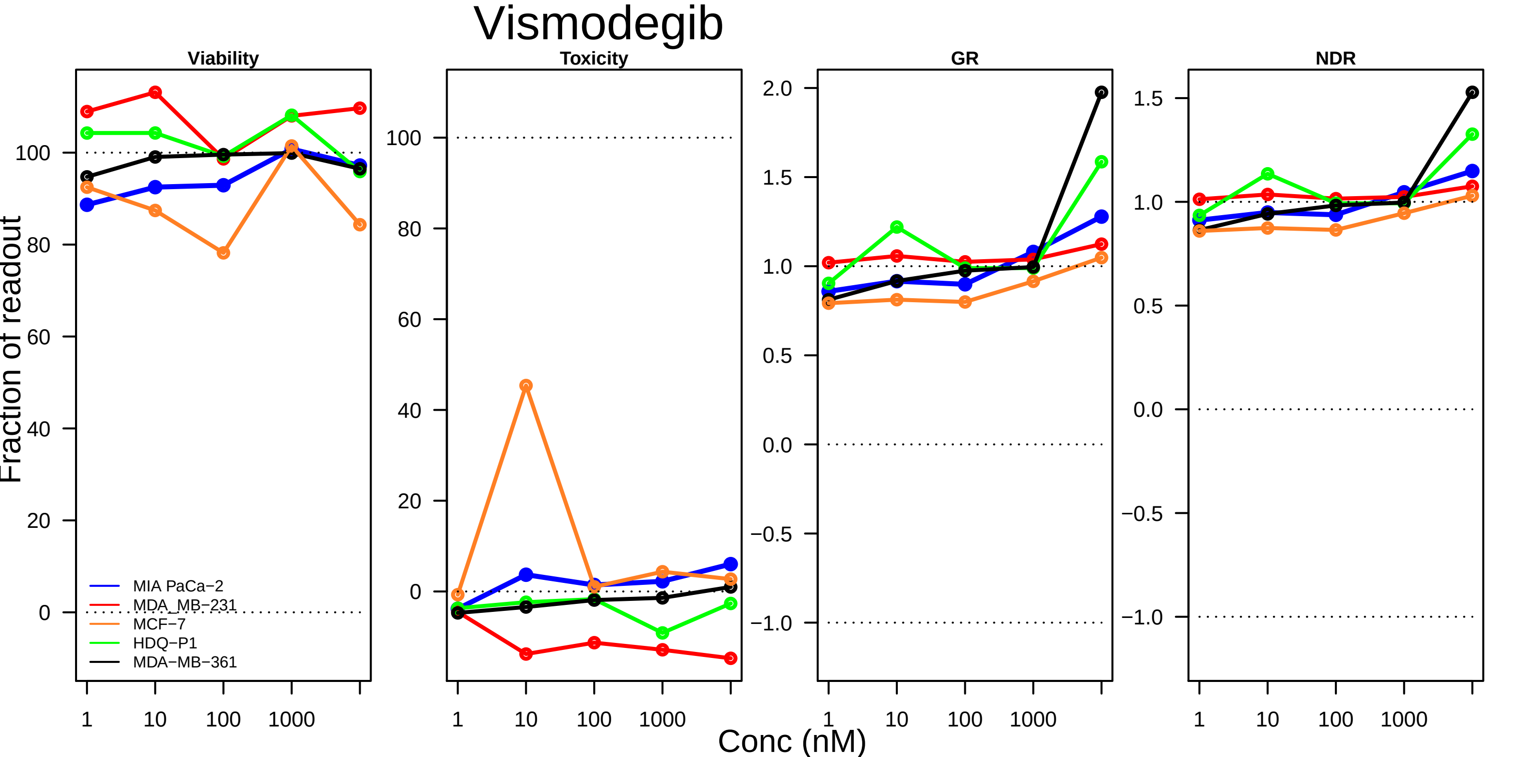

Vismodegib
Viability
Toxicity
GR
NDR
Fraction of readout
Conc (nM)
MIA PaCa−2
MDA_MB−231
MCF−7
HDQ−P1
MDA−MB−361
1
10
100
1000
0
20
40
60
80
100
2.0
1.5
1.0
0.5
0.0
−0.5
−1.0

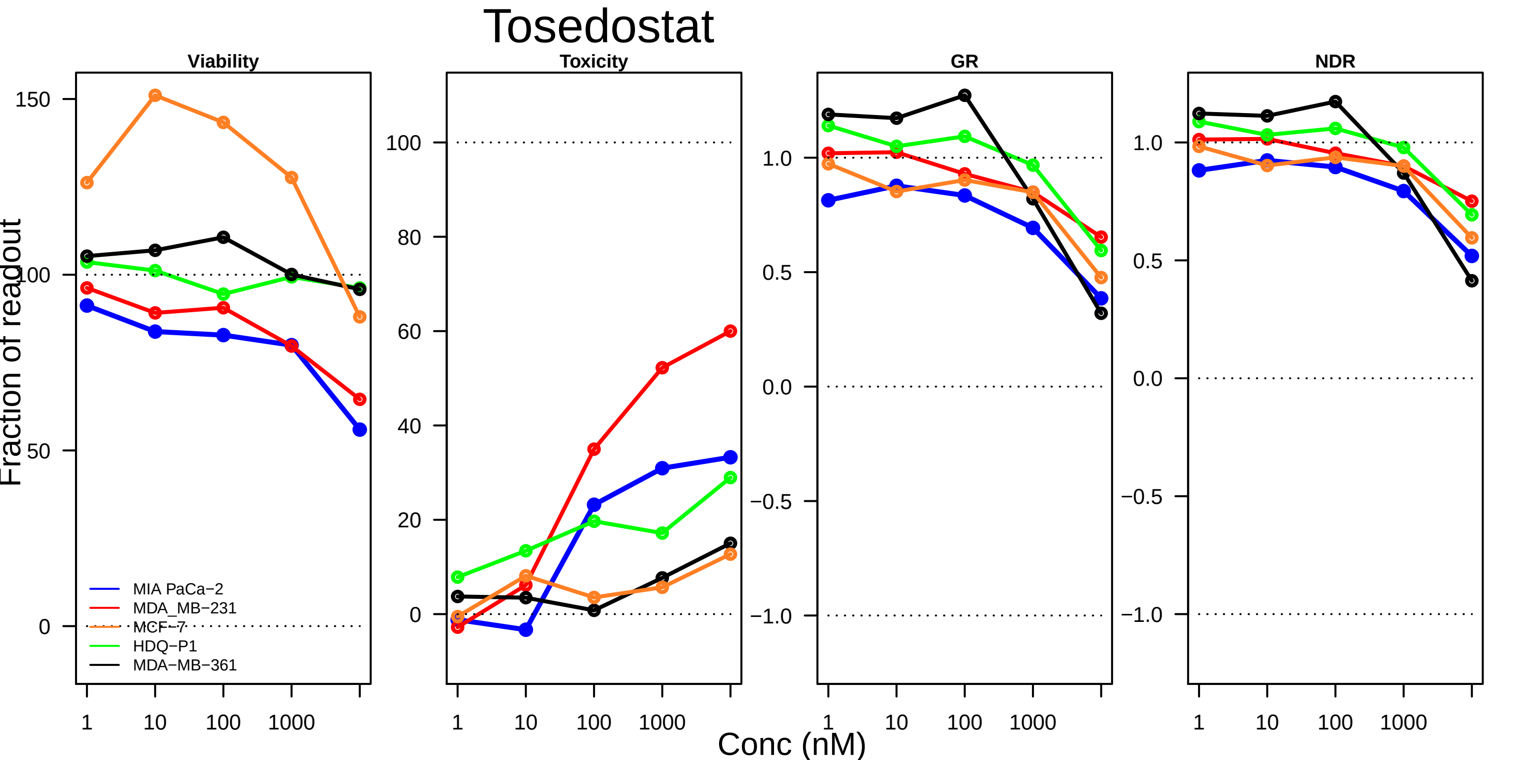
Tosedostat
Viability
Toxicity
GR
NDR
Fraction of readout
150
100
50
0
100
80
60
40
20
0
1.0
0.5
0.0
−0.5
−1.0
MIA PaCa−2
MDA_MB−231
MCF−7
HDQ−P1
MDA−MB−361
1
10
100
1000
Conc (nM)

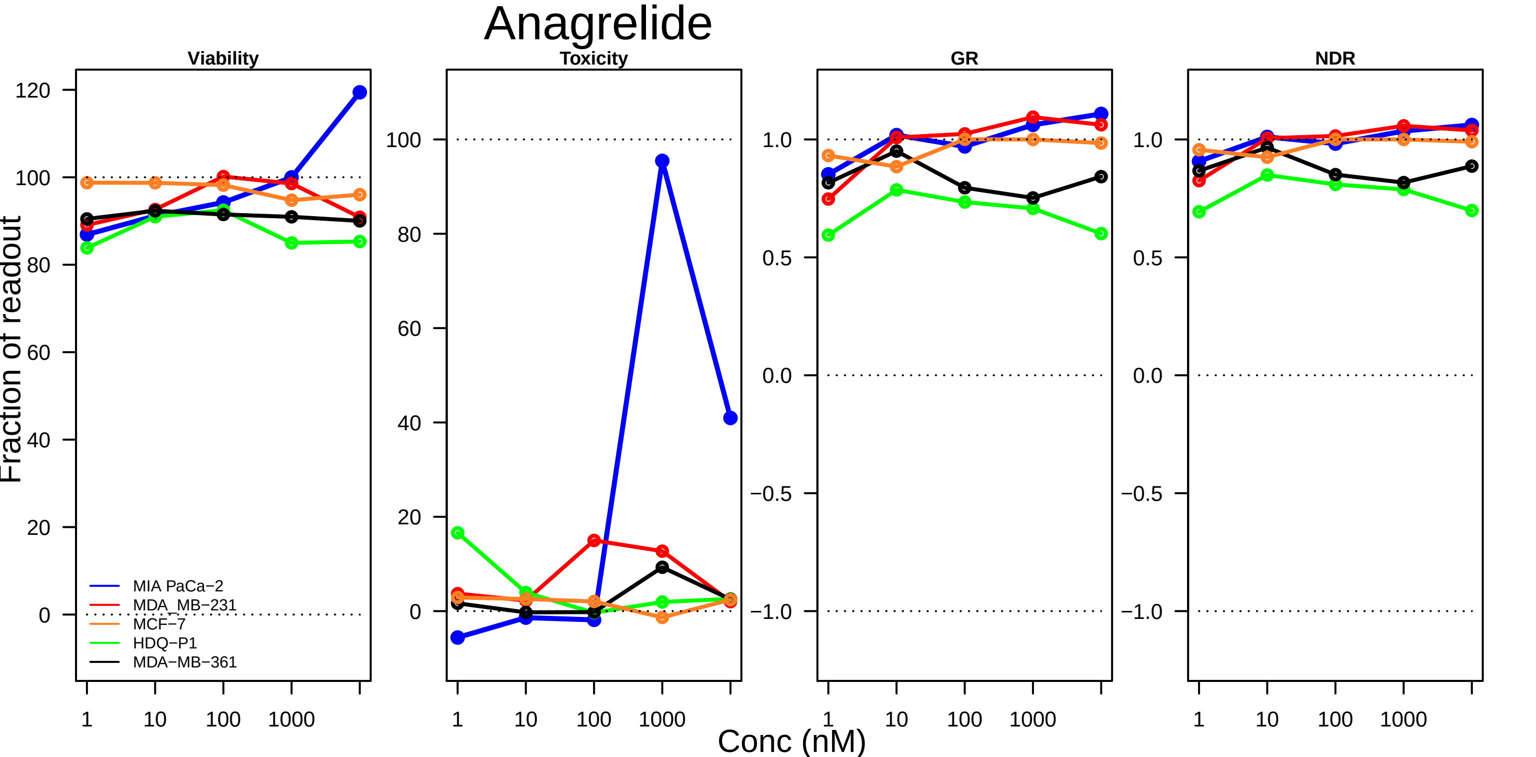

Anagrelide
Viability
Toxicity
GR
NDR
Fraction of readout
Conc (nM)
120
100
80
60
40
20
0
1.0
0.5
0.0
−0.5
−1.0
1
10
100
1000
MIA PaCa−2
MDA_MB−231
MCF−7
HDQ−P1
MDA−MB−361

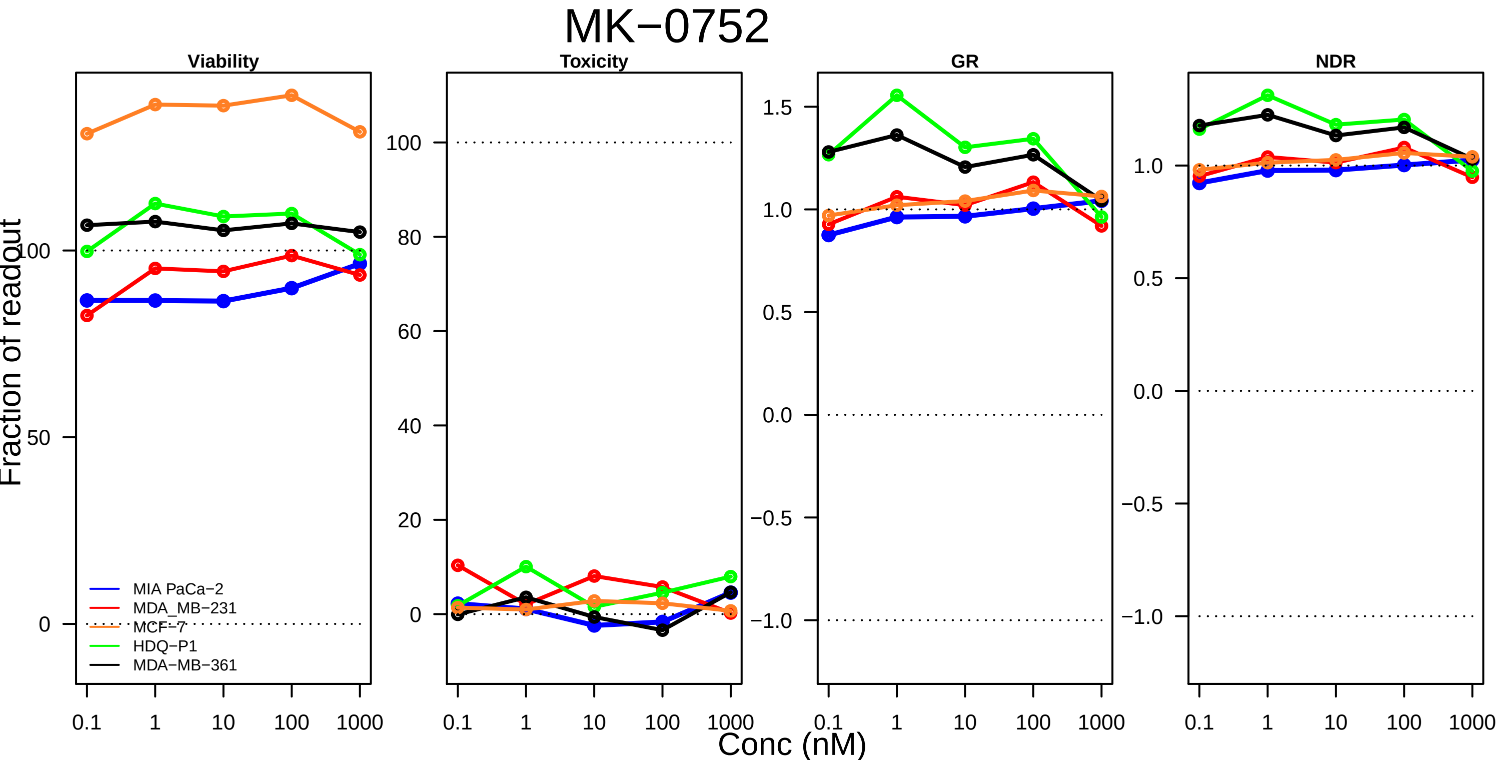
MK−0752
Viability
Toxicity
GR
NDR
Fraction of readout
Conc (nM)
0.1
1
10
100
1000
0
50
100
20
40
60
80
−1.0
−0.5
0.0
0.5
1.0
1.5
MIA PaCa−2
MDA_MB−231
MCF−7
HDQ−P1
MDA−MB−361

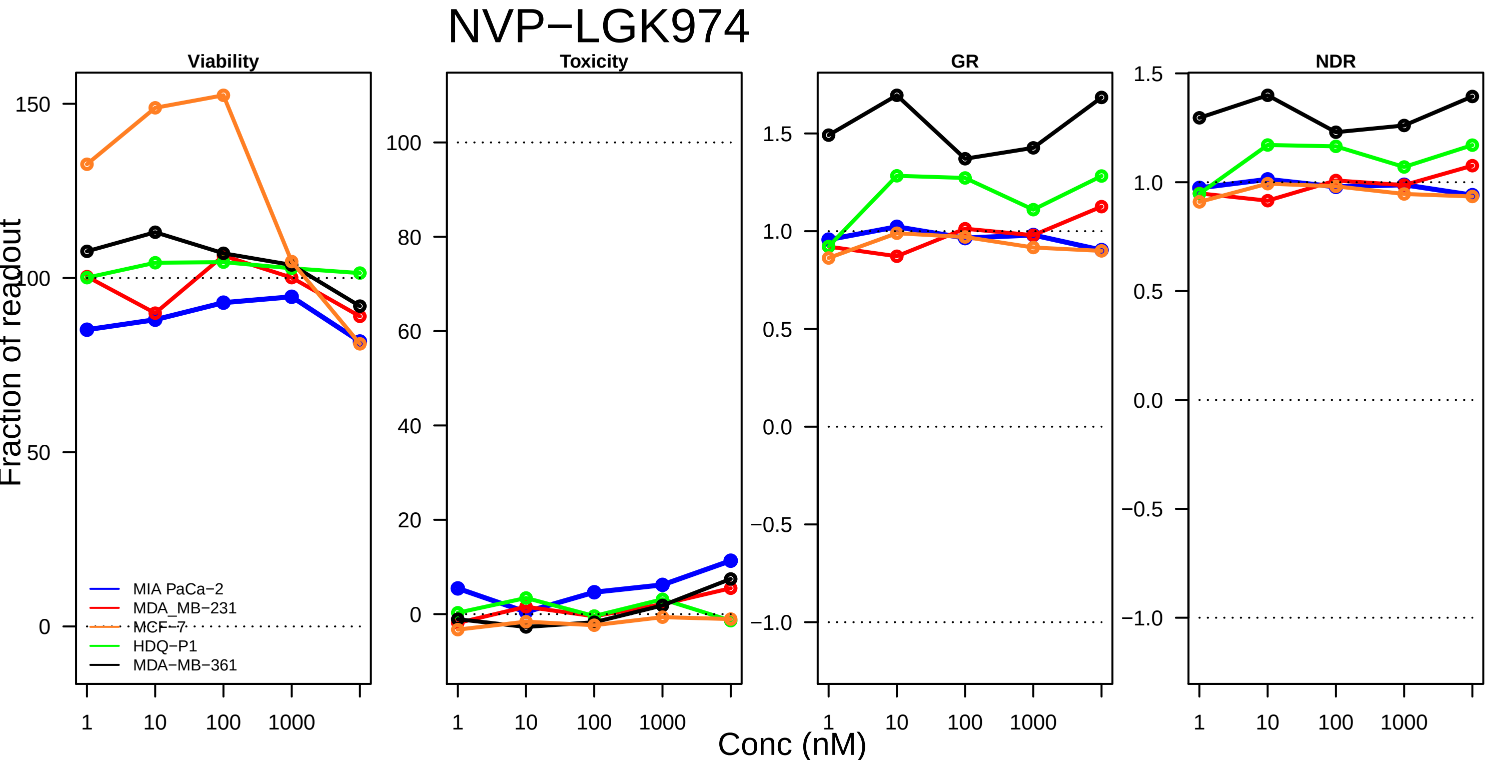

NVP−LGK974
Viability
Toxicity
GR
NDR
Fraction of readout
Conc (nM)
150
100
50
0
80
60
40
20
1.5
1.0
0.5
0.0
−0.5
−1.0
1
10
100
1000
MIA PaCa−2
MDA_MB−231
MCF−7
HDQ−P1
MDA−MB−361

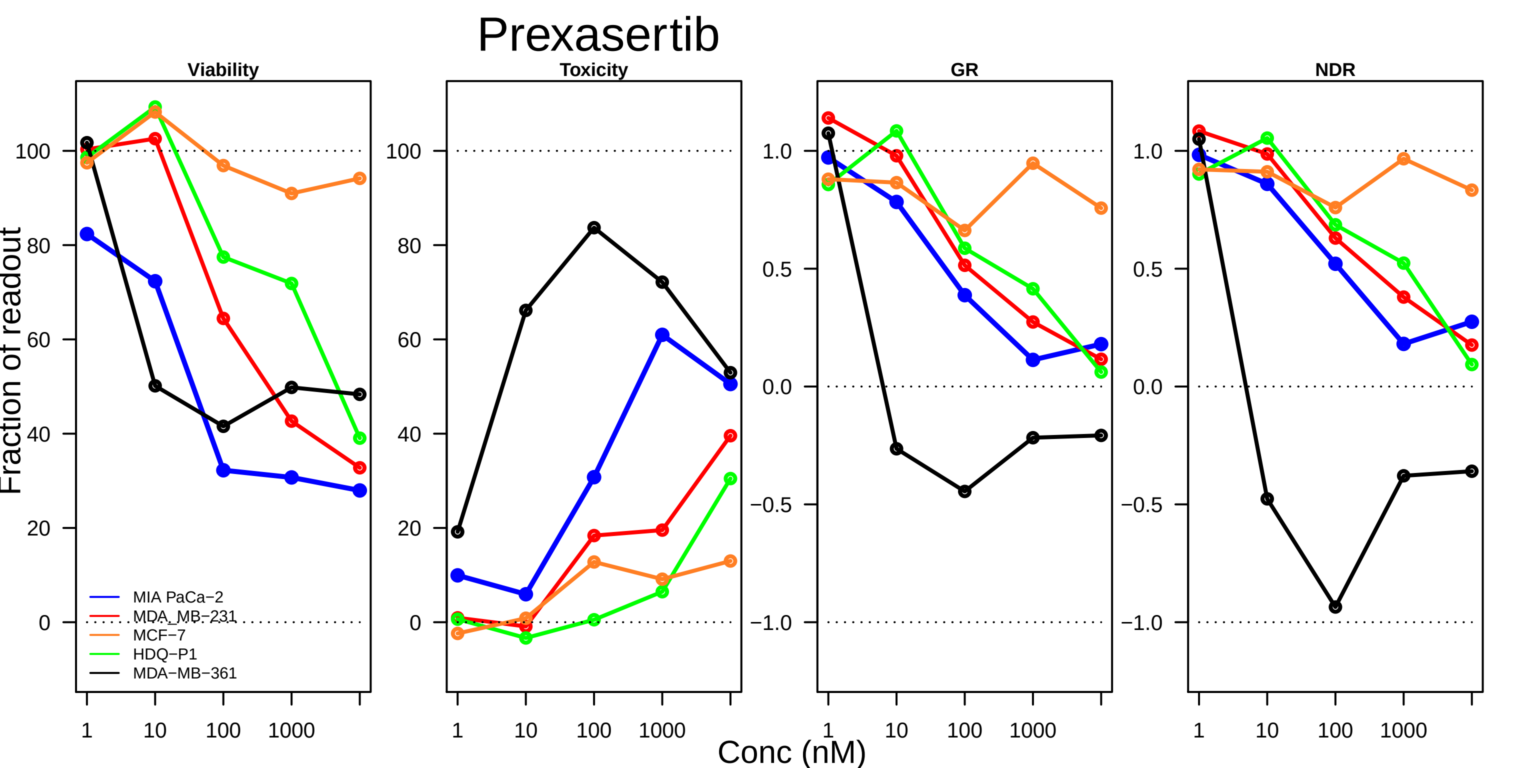

Prexasertib
Viability
Toxicity
GR
NDR
Fraction of readout
Conc (nM)
MIA PaCa−2
MDA_MB−231
MCF−7
HDQ−P1
MDA−MB−361
1
10
100
1000
0
20
40
60
80
100
−1.0
−0.5
0.0
0.5
1.0

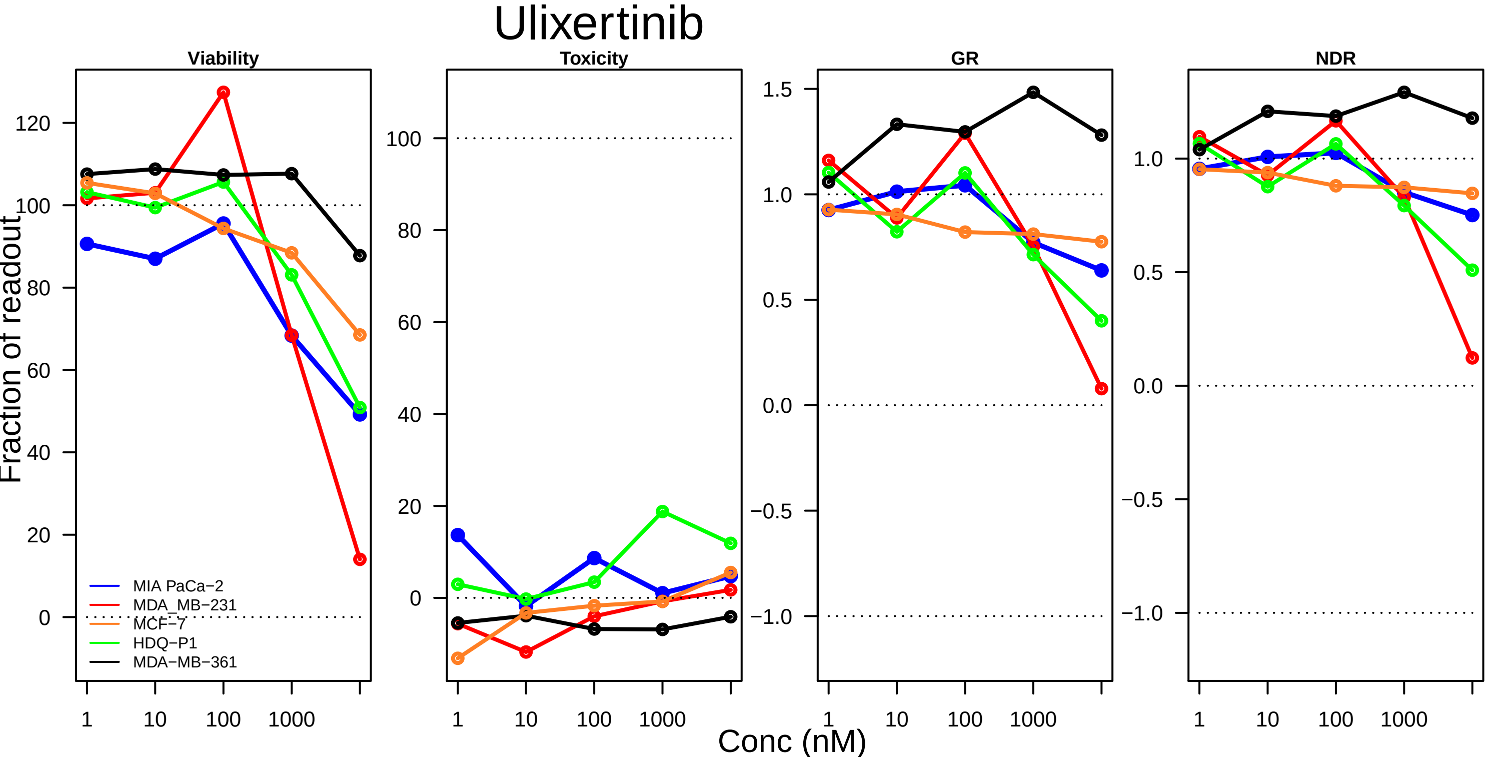

Ulixertinib
Viability
Toxicity
GR
NDR
Fraction of readout
Conc (nM)
MIA PaCa−2
MDA_MB−231
MCF−7
HDQ−P1
MDA−MB−361
1
10
100
1000

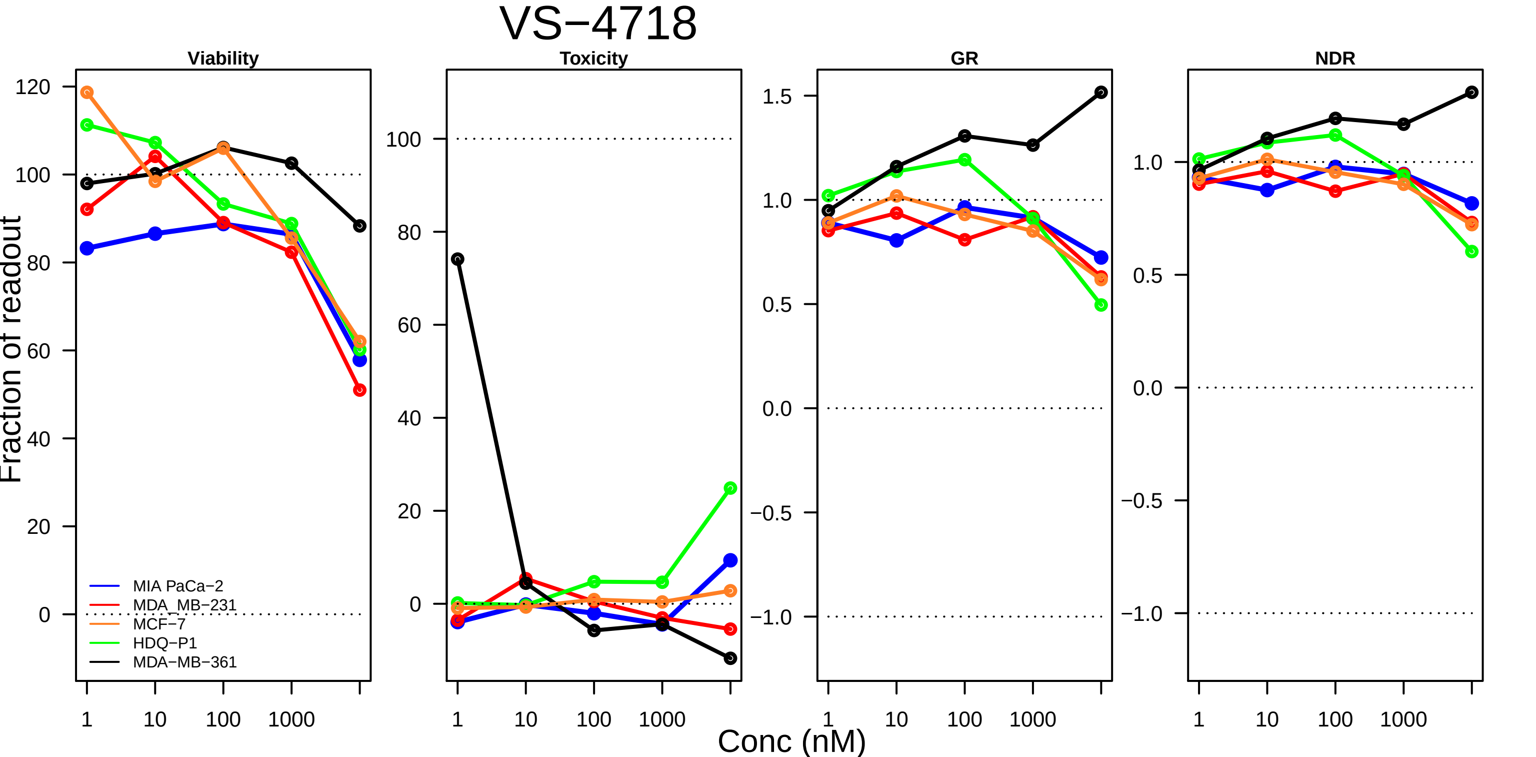

VS−4718
Viability
Toxicity
GR
NDR
Fraction of readout
Conc (nM)
1
10
100
1000
MIA PaCa−2
MDA_MB−231
MCF−7
HDQ−P1
MDA−MB−361

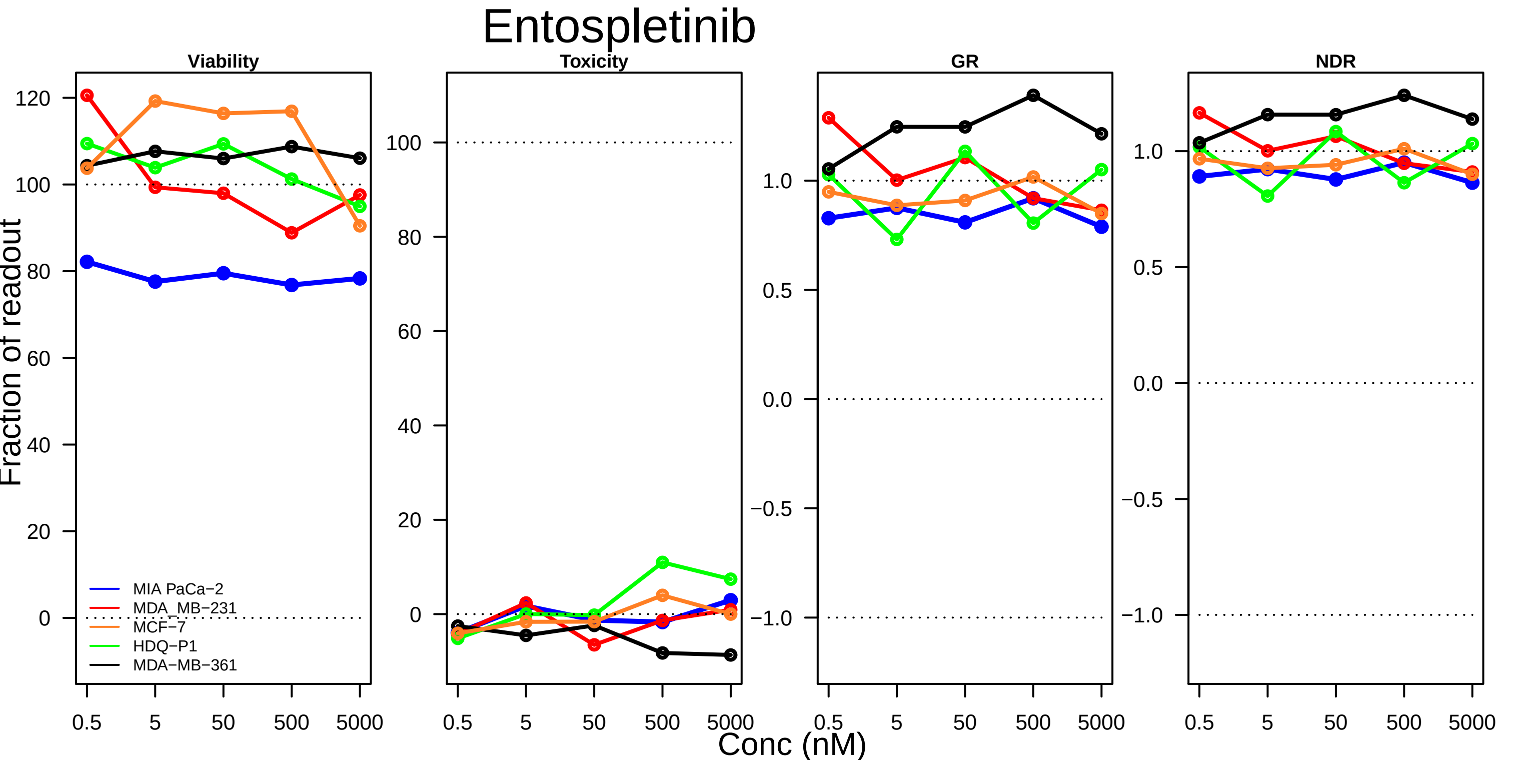

Entospletinib
Viability
Toxicity
GR
NDR
Fraction of readout
Conc (nM)
MIA PaCa−2
MDA_MB−231
MCF−7
HDQ−P1
MDA−MB−361
0.5
5
50
500
5000

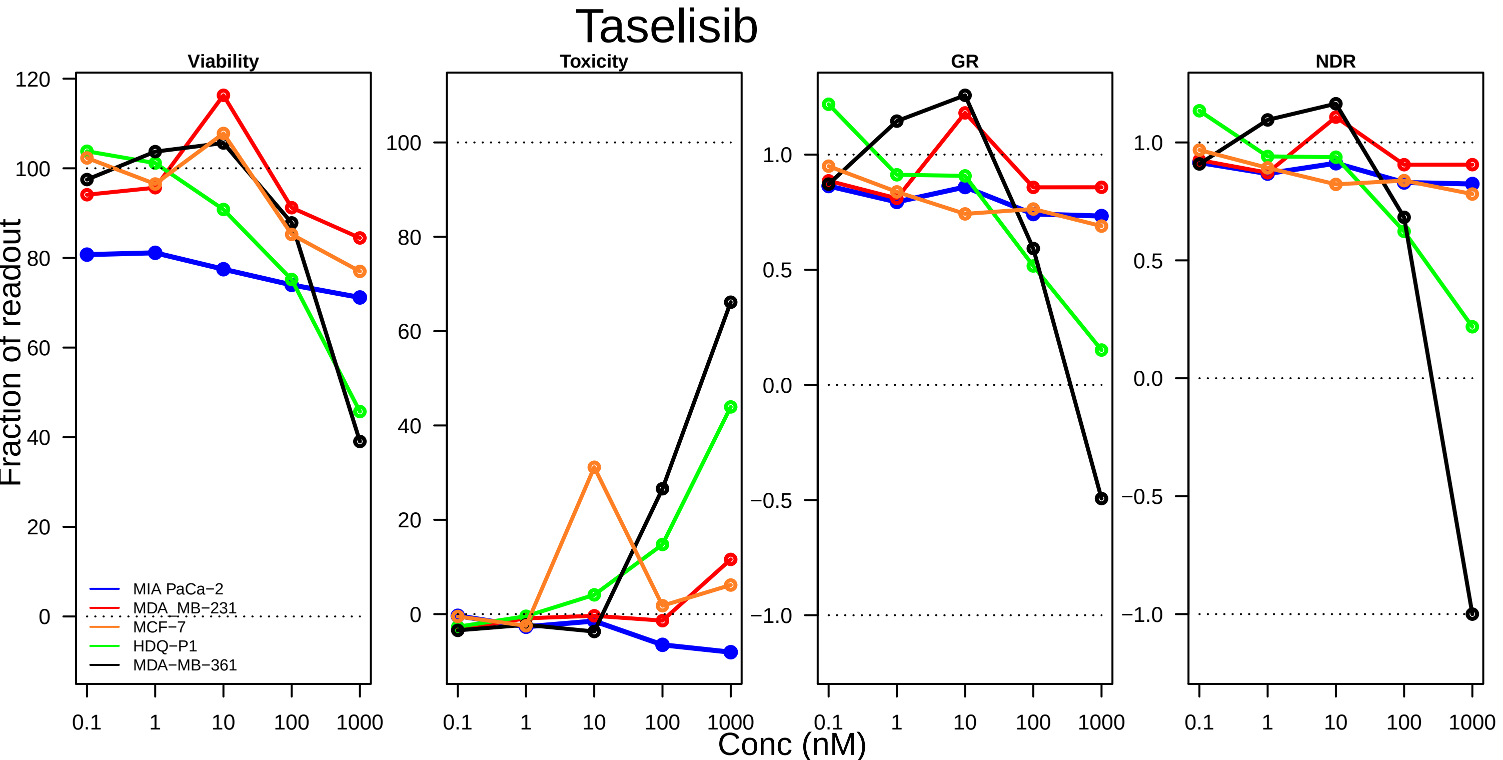
Taselisib
Viability
Toxicity
GR
NDR
Fraction of readout
MIA PaCa−2
MDA_MB−231
MCF−7
HDQ−P1
MDA−MB−361
120
100
80
60
40
20
0
1.0
0.5
0.0
−0.5
−1.0
0.1
1
10
100
1000
Conc (nM)

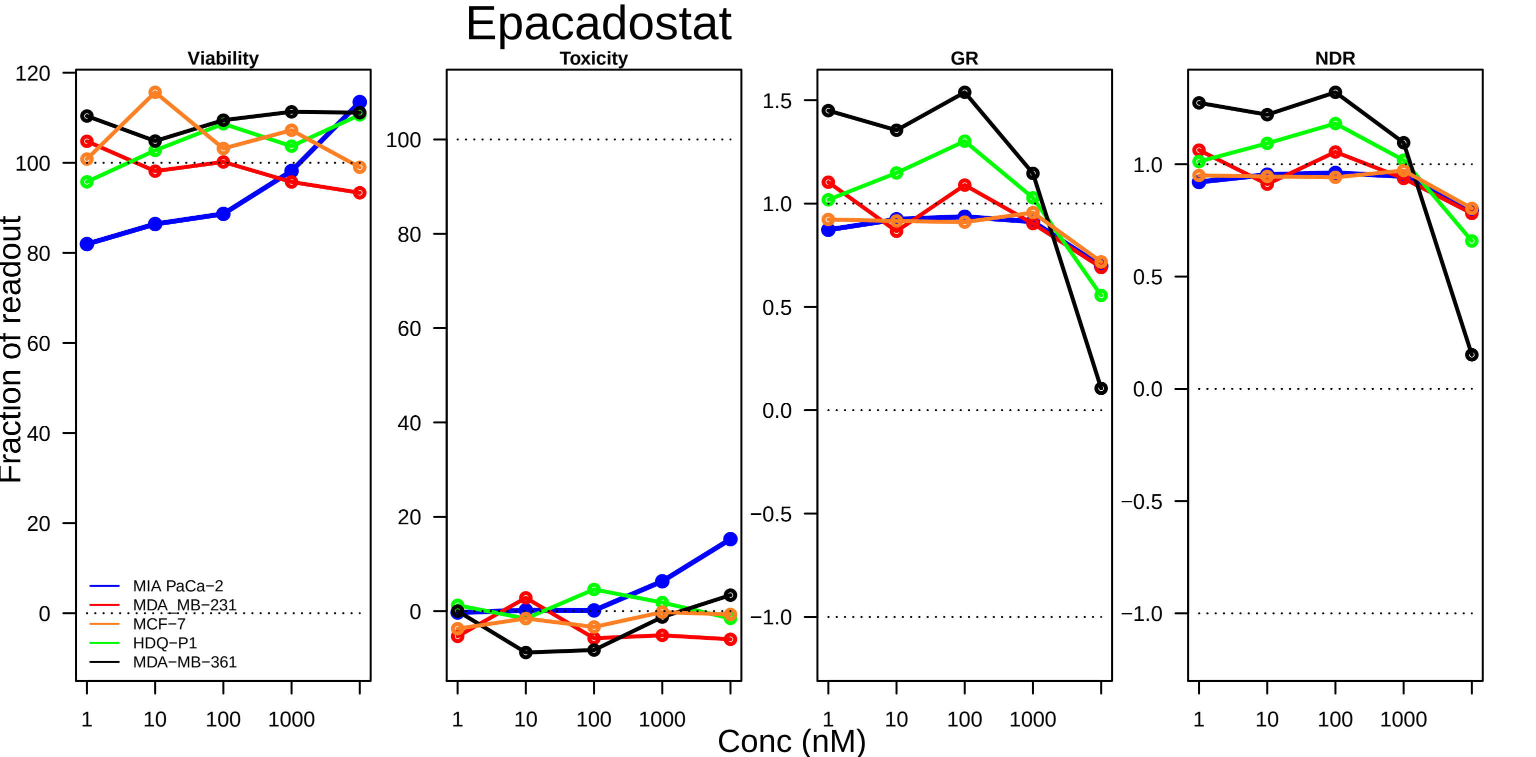
Epacadostat
Viability
Toxicity
GR
NDR
Fraction of readout
Conc (nM)
MIA PaCa−2
MDA_MB−231
MCF−7
HDQ−P1
MDA−MB−361
120
100
80
60
40
20
0
1.5
1.0
0.5
0.0
−0.5
−1.0
1
10
100
1000

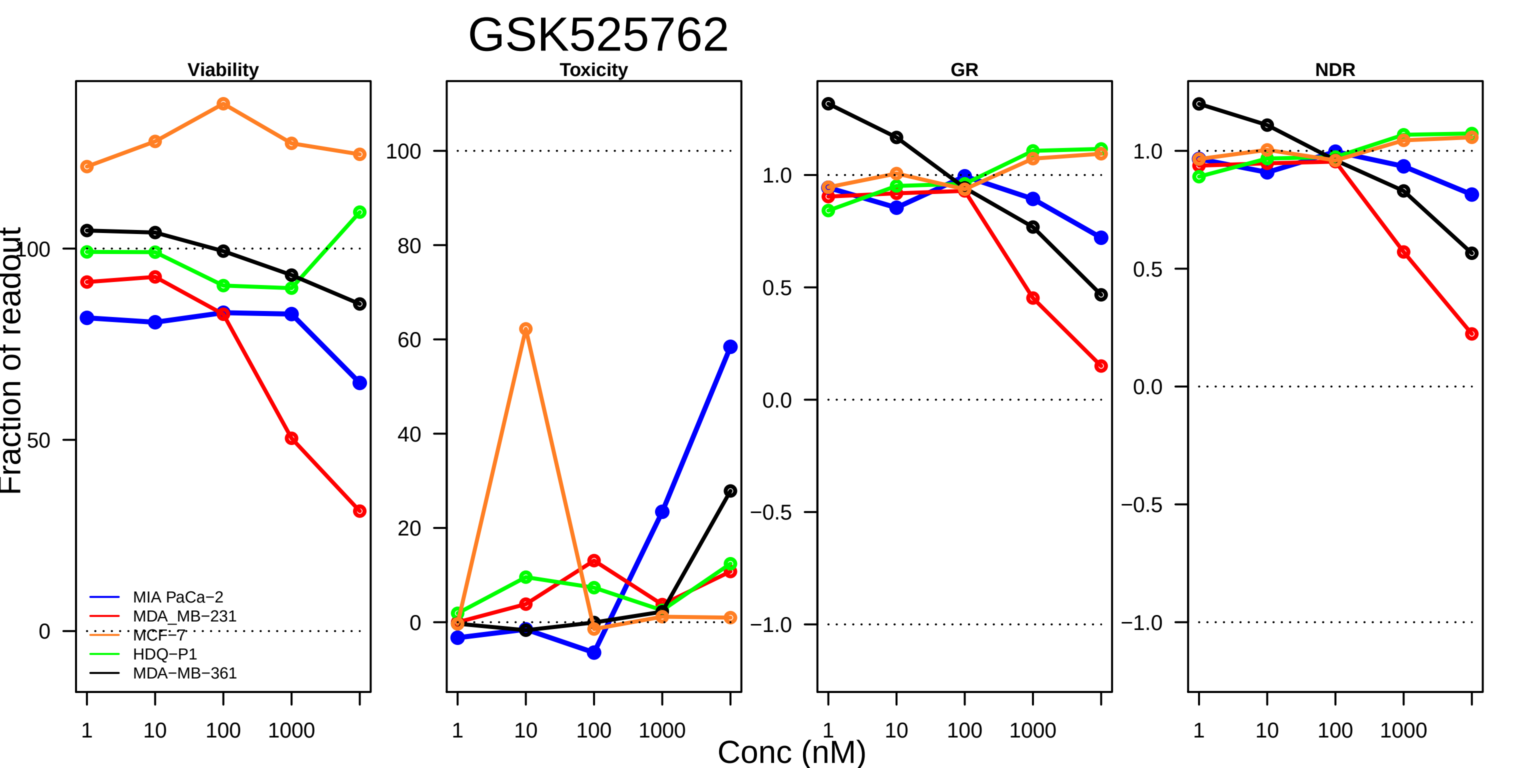

GSK525762
Viability
Toxicity
GR
NDR
Fraction of readout
Conc (nM)
MIA PaCa−2
MDA_MB−231
MCF−7
HDQ−P1
MDA−MB−361

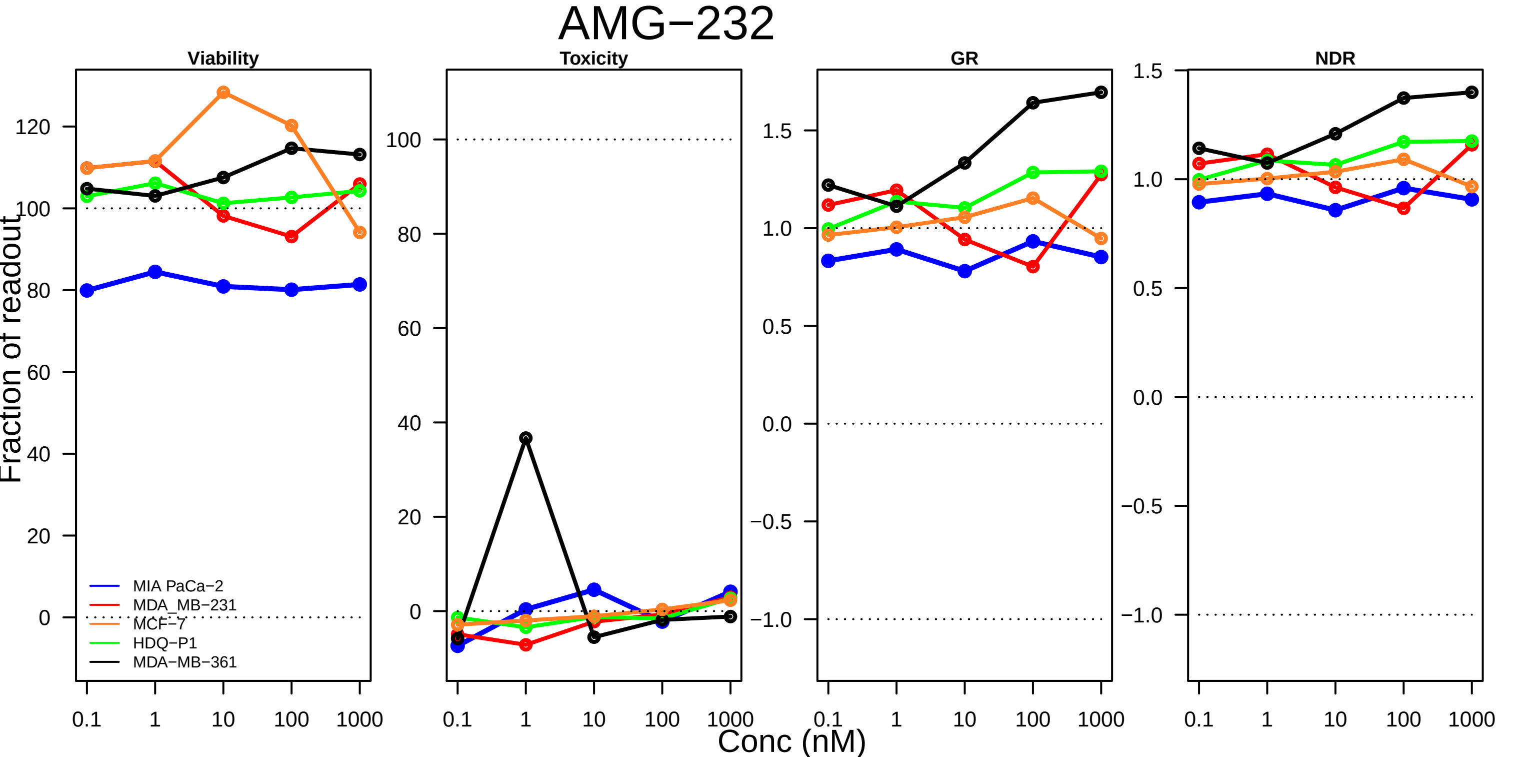

AMG−232
Viability
Toxicity
GR
NDR
Fraction of readout
Conc (nM)
0.1
1
10
100
1000
MIA PaCa−2
MDA_MB−231
MCF−7
HDQ−P1
MDA−MB−361

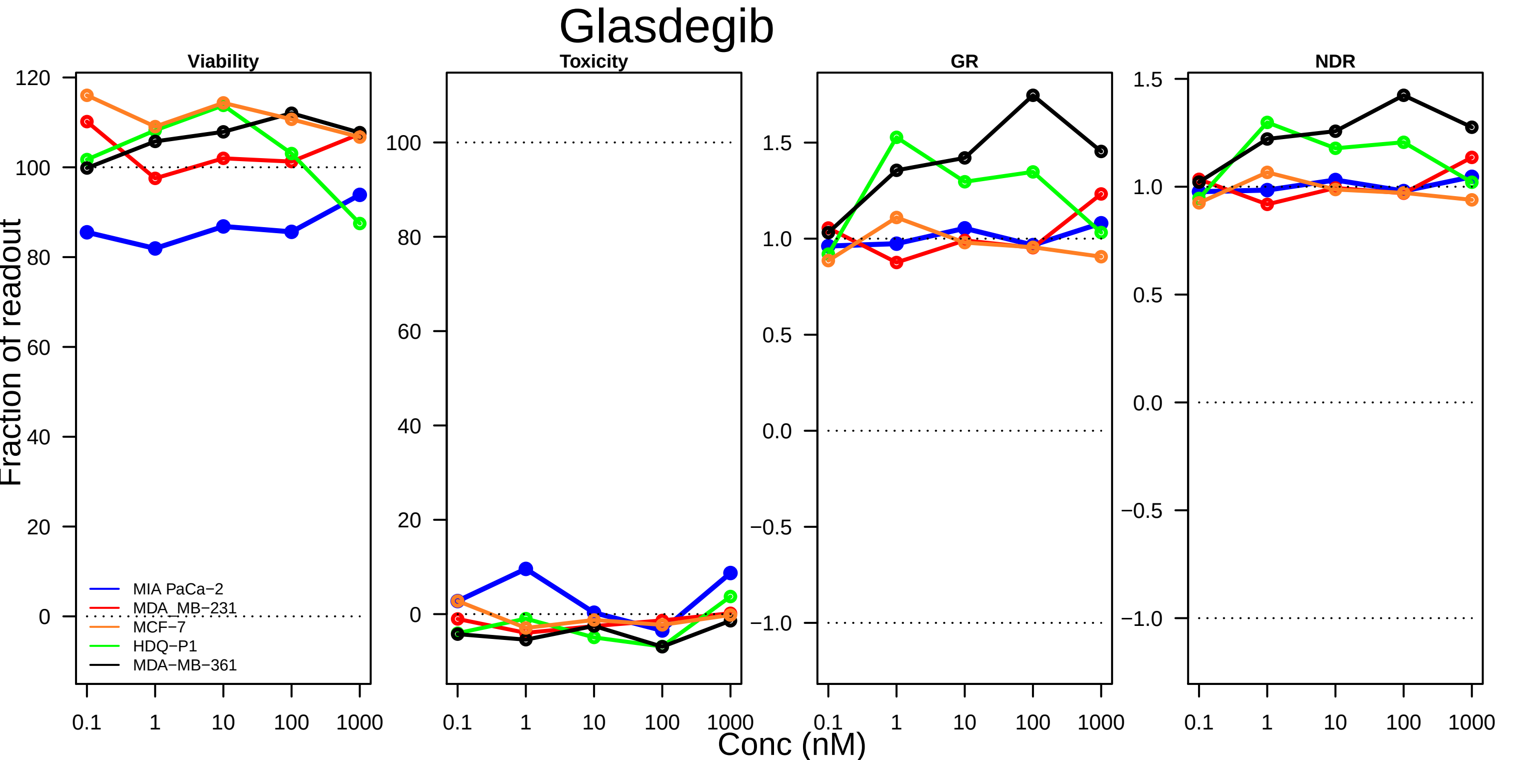

Glasdegib
Viability
Toxicity
GR
NDR
Fraction of readout
Conc (nM)
MIA PaCa−2
MDA_MB−231
MCF−7
HDQ−P1
MDA−MB−361
0.1
1
10
100
1000
